# Functional analysis of Ost3p and Ost6p containing yeast oligosaccharyltransferase*s*

**DOI:** 10.1101/2021.05.17.443621

**Authors:** Julia D. Neuhaus, Rebekka Wild, Jillianne Eyring, Rossitza N. Irobalieva, Julia Kowal, Chia-wei Lin, Kaspar P. Locher, Markus Aebi

## Abstract

The oligosaccharyltransferase (OST) is the central enzyme in the *N*-glycosylation pathway. It transfers a defined oligosaccharide from a lipid-linker onto the asparagine side chain of proteins. The yeast OST consists of eight subunits and exists in two catalytically distinct isoforms that differ in one subunit, Ost3p or Ost6p. The cryo-electron microscopy structure of the Ost6p containing complex was found to be highly similar to the Ost3p containing OST. OST enzymes with altered Ost3p/Ost6p subunits were generated and functionally analysed. The three C-terminal transmembrane helices were responsible for the higher turnover-rate of the Ost3p vs. the Ost6p containing enzyme *in vitro* and the more severe hypoglycosylation in Ost3p lacking strains *in vivo*. Glycosylation of specific OST target sites required the N-terminal thioredoxin domain of Ost3p or Ost6p. This Ost3p/Ost6p dependence was glycosylation site but not protein specific. We concluded that the Ost3p/Ost6p subunits modulate the catalytic activity of OST and provide additional specificity for OST substrate recognition.

## Introduction

*N-*linked protein glycosylation is a highly abundant post-translational modification conserved across all domains of life^1, 2, 3^. The oligosaccharides covalently linked to proteins have important roles in protein folding and trafficking, in transducing signals across membranes and in mediating specific cell-cell interactions^4, 5, 6, 7^. In eukaryotes, *N*-linked glycosylation starts with the stepwise assembly of a lipid-linked oligosaccharide (LLO) dolichyl-PP-GlcNAc_2_Man_9_Glc_3_ at the endoplasmic reticulum (ER) by proteins of the asparagine linked glycosylation (ALG) glycosyltransferase family^8, 9, 10, 11^. In the determining step, the membrane-integral oligosaccharyltransferase (OST) complex catalyses the *en bloc* transfer of the glycan from its dolichylpyrophosphate (Dol-PP) anchor to the asparagine residue of the glycosylation sequon (Asn-Xaa-Ser/Thr) in secretory proteins^12, 13, 14^. X-ray structures of the bacterial single-subunit OST PglB from *Campylobacter jejuni* in complex with a peptide and LLO analogue have revealed mechanistic details of substrate recognition, positioning and activation^15, 16^. Most eukaryotes, however, harbour multi-subunit OST complexes with eight subunits in yeast and sometimes even more subunits in multi-cellular organisms^2, 5, 17^. The PglB homolog Stt3p provides the catalytic center of the eukaryotic enzymes. Multimeric eukaryotic OSTs differ with respect to this catalytic subunit: STT3A containing OSTs associate with the translocon, STT3B complexes glycosylate substrates after their translocation into the ER. It was proposed that the additional subunits evolved to process the higher number of different glycosylation substrates present in these organisms. The yeast OST, an STT3B complex, exists in two isoforms harbouring either Ost3p or Ost6p, which were shown to have different activities and protein substrate specificities^18, 19, 20^. Ost3p/Ost6p are functionally homologous proteins that are composed of a luminal domain harbouring an oxidoreductase activity and four transmembrane helices at their C-terminus. Ost3p/Ost6p are relevant for site-specific glycosylation^18, 21, 22^. As the yeast OST, the mammalian STT3B OST complex incorporates homologues of Ost3p/Ost6p (IAPp/N33p) ^2, 6, 17, 23^. The recently published structures of the Ost3p containing complex and the here presented Ost6p containing complex enable a detailed analysis of the structure of the two isomeric OST complexes in yeast^24, 25^.

To address the origin of the differences found in glycosylation activity and the protein substrate specificities between the two isomers, yeast strains expressing defined Ost3/Ost6p chimeras and truncation variants of the Ost3p and Ost6p proteins were generated for *in vitro* and *in vivo* analysis^20, 26^. The combination of enzyme kinetic studies *in vitro* and glycosylation site occupancy determination *in vivo* reveal how the different domains of Ost3p and Ost6p modulate OST activity.

## Results

### Structure of the Ost6p containing OST complex

To learn more about the functional differences between Ost3p and Ost6p containing OSTs on a molecular level, we aimed for a high-resolution structure of the Ost6p containing complex. OST was purified from a yeast strain that only expressed Ost6p containing complexes24. A short-tailed LLO analogue and an inhibitory peptide containing a 2,4-diaminobutanoic acid (Dab) instead of an asparagine at the glycosylation site were added to the detergent-purified protein prior to cryo-electron microscopy (cryo-EM) grid preparation (see Methods section, Supplementary Figure 1). The peptide had an IC_50_ value in the micromolar range (137 ± 39 μM) suggesting that it can bind with a moderate affinity (Supplementary Figure 1B). We did not, however, observe either of the substrates in the structure.

Processing in Relion 3.0 resulted in a 3D reconstruction at a resolution of 3.46 Å (high-resolution map, Supplementary Figure 2 and 3). The structure shows clear density for all four transmembrane helices of Ost6p with TM1 being slightly less well ordered (Figure 1A and 1B). Similar to the previously published Ost3p structure, the luminal domain of Ost6p was not well resolved. Most importantly, the structures of Ost3p and Ost6p containing OSTs were highly similar (R.M.S.D. of 0.569 Å across 628 atom pairs). The most prominent structural differences were observed in the transmembrane region of Ost3p and Ost6p (R.M.S.D. of 1.025 Å across 84 atom pairs) with TM2 being slightly shifted and Ost6p lacking the short helix between TM3 and TM4 (Figure 1C and 1D).

**Figure 1:**
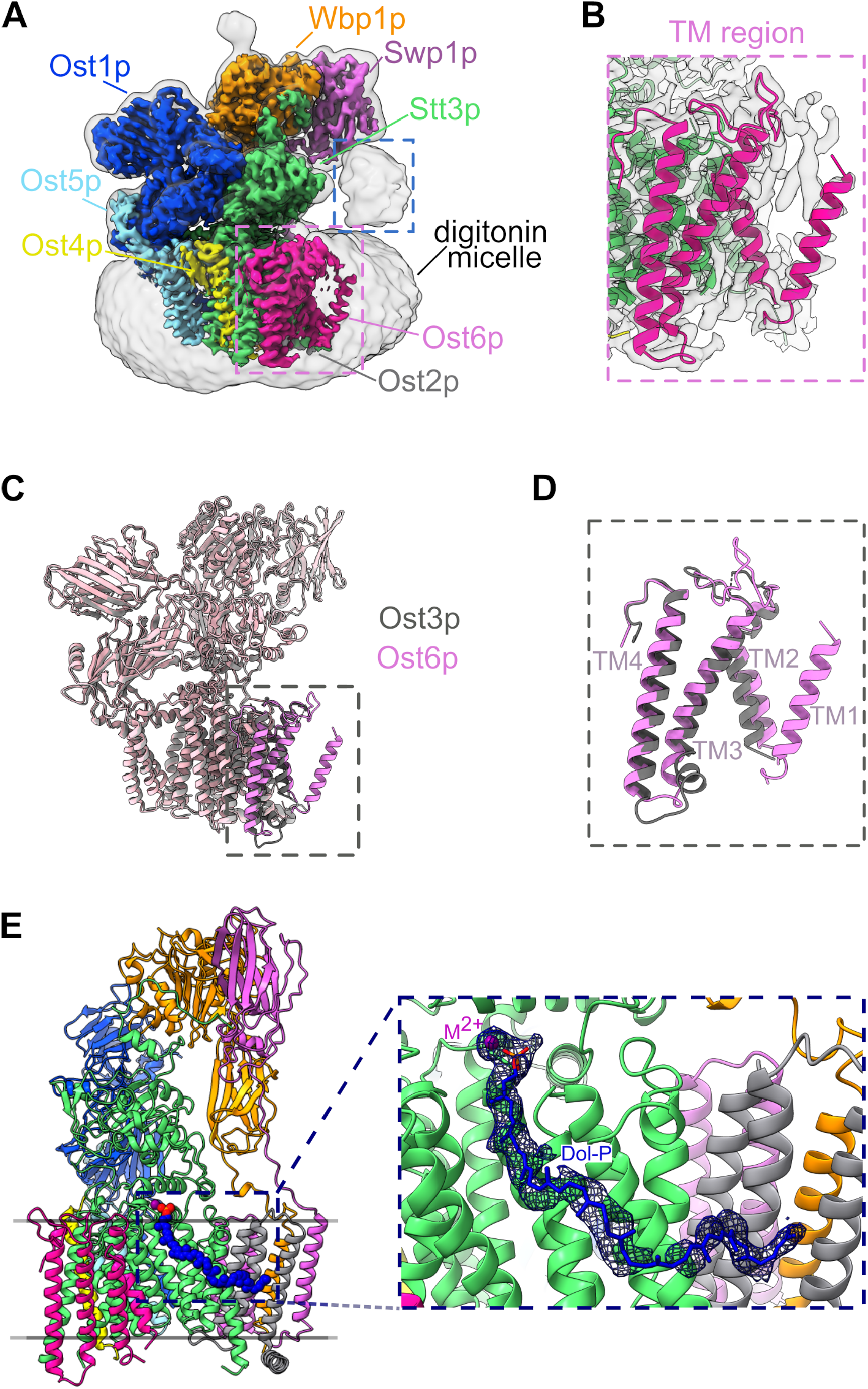
Structure of the Ost6p containing OST and comparison with the Ost3p containing form. **(A)** Local resolution filtered cryo-EM map of the octa-subunit complex with different subunits colored as indicated. An additional low-pass filtered map around the complex shows density for the detergent micelle and the luminal domain of Ost6p, which is indicated with a blue box. A pink box highlights the transmembrane region of Ost6p. **(B)** Zoom-in into the transmembrane region of Ost6p. The structure is shown as ribbon diagram and the surrounding low-pass filtered cryo-EM map is colored in light grey. **(C)** Superposition of the Ost3p (PDB-ID 6EZN) and Ost6 containing complexes colored in grey and pink, respectively. The TM region of Ost3p and Ost6p are colored in darker shades. **(D)** Close-up view onto the superimposed transmembrane region of Ost3p and Ost6p. **(E)** Ribbon representation of the Ost6p containing OST structure with same colors as in (A). The close-up view highlights the binding side of the metal ion (M2+) and a co-purified dolichylphosphate (Dol-P). The blue mesh depicts the cryo-EM map zoned around the substrates with a radius of 2.2 Å.

Similar as reported for the mammalian OST^17^ we found that the LLO binding site was occupied by a density that extends beyond the neryl-citronellyl tail (20 carbon atoms, see Methods) of the supplied LLO analogue and that fitted a phosphate head group rather than the expected oligosaccharide and pyrophosphate moieties (Figure 1E). We thus propose that this structure contains a co-purified dolichylphosphate of which 55 carbon atoms of the isoprenoid lipid tail were resolved.

### Generation of yeast strains with mutant OST complexes

Our structural analysis of the yeast OST complexes revealed that both Ost3p and Ost6p were characterized by a flexible N-terminal part consisting of the thioredoxin fold and TM1 and a C-terminal part formed by TM2-4 that interacted with Stt3p. Based on this high structural similarity, we created, for functional analysis of these two domains, Ost3p and Ost6p chimera, where either the luminal domain or the luminal and TM1 of both proteins were exchanged. Additionally, truncations that lack either the luminal domain or the luminal domain and the TM1 of Ost3p or Ost6p were generated (Figure 2A). The mutant OST3/OST6 genes were placed on a multicopy plasmid and transformed into a yeast strain with a deletion of both OST3 and OST6 (DKO)^28^. The resulting strains expressed a uniform OST and were used as a starting material for the purification of the recombinant enzyme and to study the function of mutant OSTs *in vivo*.

**Figure 2:**
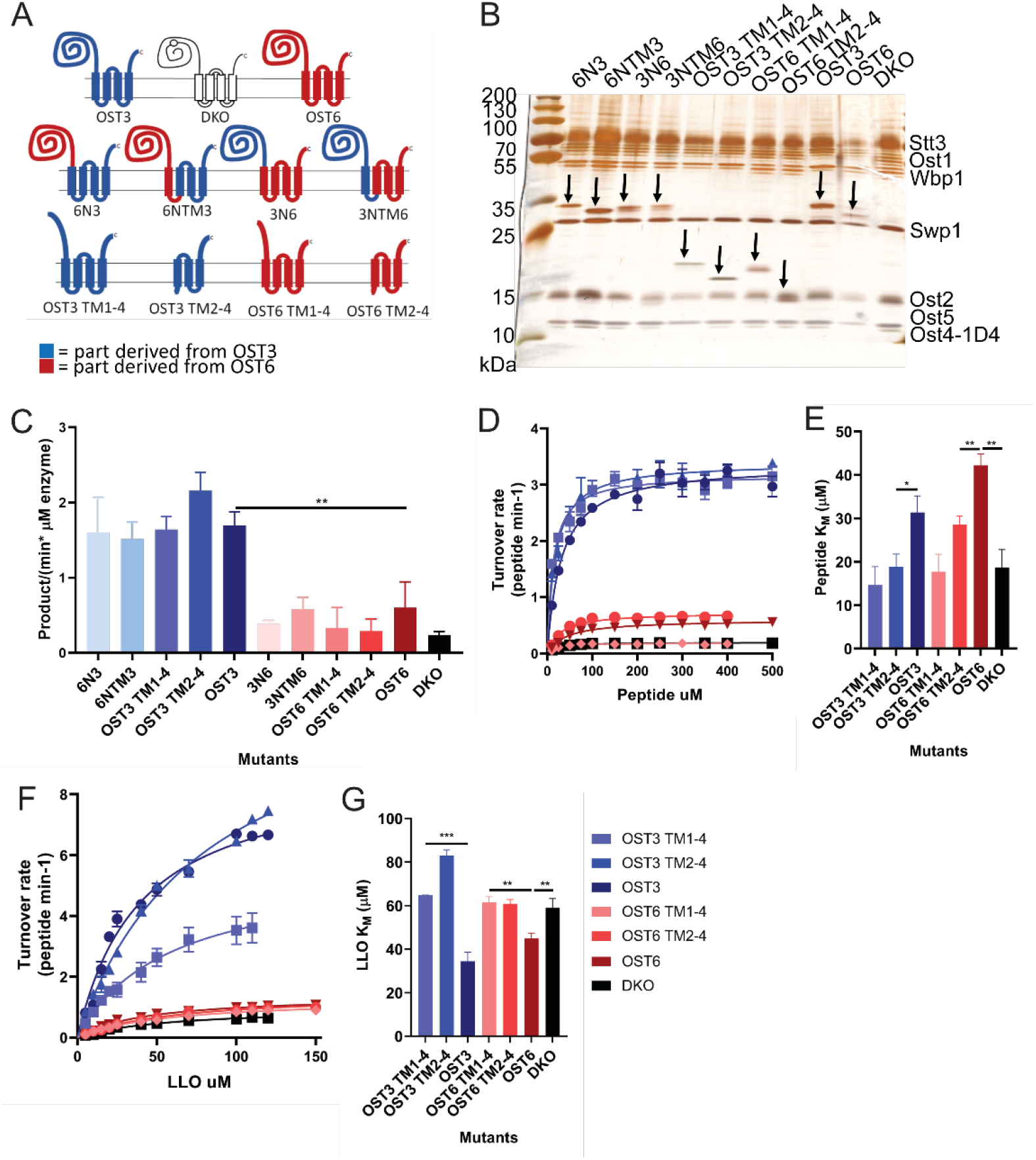
*In vitro* analysis of purified Ost3p and Ost6p derivative containing OST complexes. **(A)** Schematic representation of Ost3p and Ost6p wild-type, chimera and truncation constructs. The mutant OST3/OST6 genes were placed on a multicopy plasmid and transformed into a yeast strain with a deletion of both OST3 and OST6 (DKO) and expressing a 1D4-tagged version of Ost4p. **(B)** Silver-stained SDS-PAGE of the Ost3p and Ost6p derivative containing OST complexes purified via 1D4-tag based affinity enrichment and size exclusion chromatography. **(C)** Initial velocity rates of Ost3p/Ost6p derivative and DKO OST complexes. Reactions were performed using 100 mM TAMRA-YANTAS peptide, 50 mM LLO and varying concentrations between 0,05 and 0,3 mM OST complexes for 2,5-40 minutes. The slopes were determined by using the linear regression of the initial phase of the reaction. **D**) Averaged turnover rates and (**E**) K_M_ of the different Ost3p and Ost6p derivative containing OST complexes at enzyme concentrations of 0,1 mM (Ost3p derivative containing OST complexes) or 0,2 mM (Ost6p derivative and DKO containing OST complexes). Reactions were performed using 10-500 mM TAMRA-YANATS peptide, 100 mM LLO and for 2,5-25 (Ost3p derivative containing OST complexes) or 10-45 (Ost6p derivative and DKO containing OST complexes) minutes. Initial rates of reaction were determined by the linear regression of the linear part of the initial phase of reaction. Enzyme concentrations were taken into account. The resulting data points were fitted by non-linear regression using the Michaelis-Menten formula (K_M_ values). (**F**) Averaged turnover rates and (**G**) K_M_ of the different Ost3p and Ost6p derivative containing OST complexes at enzyme concentrations of 0,1 mM (Ost3p derivative containing OST complexes) or 0,5 mM (Ost6p derivative and DKO containing OST complexes). Reactions were performed using 25 mM TAMRA-YANATS peptide, 5-150 mM LLO and for 2,5-25 minutes. Initial rates of reaction were determined by the linear regression of the linear part of the initial phase of reaction. Enzyme concentrations were taken into account. The resulting data points were fitted by non-linear regression using the Michaelis-Menten formula (K_M_ values).

### *In vitro* analysis of mutant Ost3p and Ost6p containing yeast OST complexes

Purified mutant OST complexes were analysed by SDS-PAGE and the glycosylation status of the different OST subunits was examined using mass spectrometry (Figure 2B, Supplementary Figure 4A and 5A). As some of the OST components are *N-*glycoproteins themselves, alterations of OST activity can lead to hypoglycosylation of the subunits. We observed that all mutant OST complexes showed wild-type glycosylation levels for the sites OST1_N99, STT3_N539 and WBP1_N332, but variably reduced levels for OST1_N217 and WBP1_N60 (Supplementary Figure 4A and 5A). Of note, the WBP1_N60 does not have a clear density for a *N-*linked glycan in our Ost6p containing OST structures. This hypoglycosylation of OST subunits does not affect *in vitro* activity of the purified complexes^20^. The glycan structures of the OST subunits were identical for every mutant OST complex (Supplementary Figure 4B).

*In vitro* glycosylation assays using synthetic substrates confirmed that the Ost3p containing complex showed an approximately three times higher velocity compared to the Ost6p containing complex (Figure 2C)^28^ which in turn was about three times as active as the complex lacking an Ost3p/Ost6p subunit. This regulation of OST velocity depended primarily on the origin of the three C-terminal transmembrane helices: the chimera harbouring an Ost6p luminal domain and the membrane domain of Ost3p, for example, displayed a similar velocity compared to the Ost3p wild-type protein (Figure 2C). The complete absence of the Ost3p/Ost6p subunit (DKO) resulted in the lowest velocity.

We determined the Michaelis-Menten constant K_M_ of different OST complexes for both substrates of the glycosylation reaction, peptide and LLO (Figure 2D-G). Interestingly, the K_M_ values for the peptide were lower for the N-terminal deletion proteins and the DKO when compared to the full-length Ost3p or Ost6p containing OST (Figure 2D and 2E). In contrast, the Ost3p/Ost6p truncations showed a higher K_M_ for the LLO substrate when compared to the full-length proteins (Figure 2F and 2G). We concluded that the Ost3p/Ost6p subunits regulated the activity of OST^20^ with a similar affinity towards peptide and LLO substrate *in vitro*.

### *In vivo* analysis of mutant Ost3p and Ost6p containing yeast OST complexes

*In vivo*, OST glycosylates a multitude of N-X-S/T sequons embedded in different polypeptides. Therefore, we examined the effect of the different OST alterations on the glycosylation state of a defined set of *N-*glycosylation sites *in vivo* (51 glycosylation sites of 41 proteins) (Supplementary Figure 6A) in strains expressing mutant OST complexes. The analysis validated, at the same time, that all OST subunits were expressed at comparable protein levels in all generated strains (Supplementary Figure 5B and 5C).

Our SILAC based quantitative glycoproteomics approach confirmed that individual glycosylation sites were affected differently by Ost3p/Ost6p alterations and that the Ost3p containing complex had a higher *in vivo* glycosylation activity as compared to the Ost6p containing OST. The complete deletion of both proteins (DKO) resulted in the most severe hypoglycosylation phenotype (Figure 3A, Supplementary Figure 6A and 7A) ^18, 19, 20, 22, 26^ but affected only a subset of glycosylation sites. A similar observation was made when comparing the truncated versions of Ost3p and Ost6p harbouring TM2-4 only (Figure 3B).

**Figure 3:**
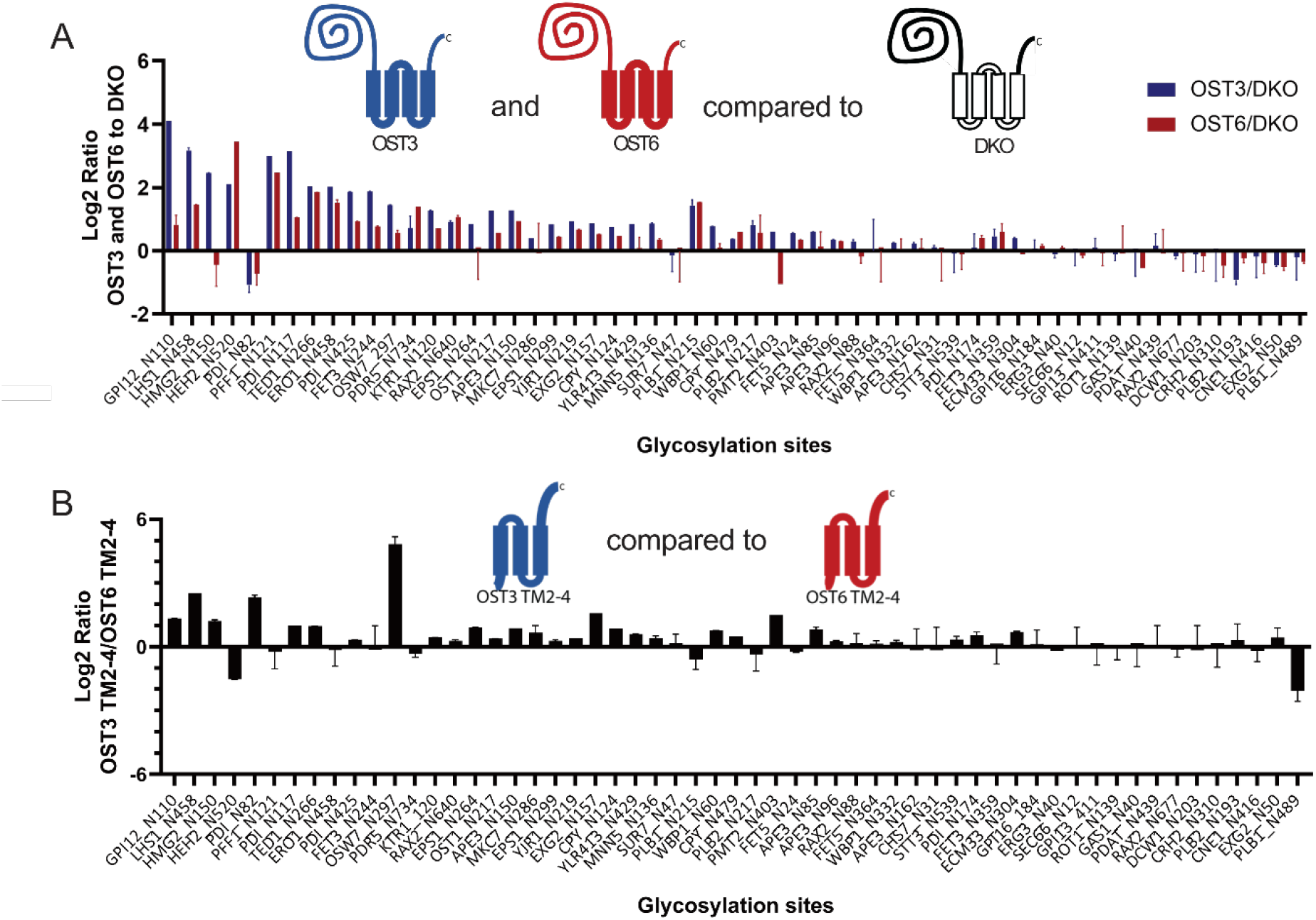
*In vivo* glycosylation occupancy comparison of Ost3p and Ost6p derivative containing OST complexes. Chimera, truncations, Ost3p, Ost6p and DKO containing OST strains were grown in normal medium, mixed in a ratio of 1:1 with the wild-type reference strain grown in ^15^N arginine and lysine medium. The glycosylation occupancy was calculated by the ratio of Ost3p/Ost6p derivate signal to WT signal for each peptide, normalized for protein expression and mixing error. **(A)** Log2 of the glycosylation occupancy ratio of Ost3p or Ost6p containing OST complex relative to the DKO containing. **(B)** Log2 of the glycosylation occupancy ratio of the ratio of OST3 TM2-4 containing OST complex relative to the OST6 TM2-4 containing OST complex are shown. Positive bars represent glycosylation sites with a preference for the presence of the Ost3p derivative subunit.

The different Ost3p and Ost6p chimera and truncation constructs showed intermediate changes in the glycosylation pattern compared to the native Ost3p/Ost6p proteins (Supplementary Figure 6A). A glycosylation occupancy comparison of the native versus the truncated Ost3p/Ost6p and chimera proteins suggested that some glycosylation sites depend on the presence of the Ost3p luminal domain (Supplementary Figure 7B and 7C) while the differences for Ost6p were minor (Supplementary Figure 7D and 7E).

We observed that individual *N-*glycosylation sites responded differently to Ost3p/Ost6p alterations. To visualize this site-specific response, we chose a “star-representation” to facilitate the comparison of individual sites (Figure 4). As a reference, we selected the glycosylation efficiency observed in a strain containing neither Ost3p nor Ost6p (DKO). The majority of the sites were preferably glycosylated by Ost3p OST and its derivatives (e.g. EPS1_N264), some by Ost6p OST (e.g. HEH2_N520), and a few sites were strongly hypoglycosylated in the absence of native Ost3p (e.g. GPI12_N110) (Figure 4B). Interestingly, we noted that some sites showed slightly higher glycosylation site occupancies in the DKO strain (e.g. SEC66_N12) (Figure 4B) ^18, 22, 26^. The four analysed glycosylation sites of Ape3p and the three analysed glycosylation sites of Pdi1p revealed that the efficiency of the different OST derivatives to catalyse glycan transfer was not protein-but site-dependent (Figure 4C and D).

**Figure 4:**
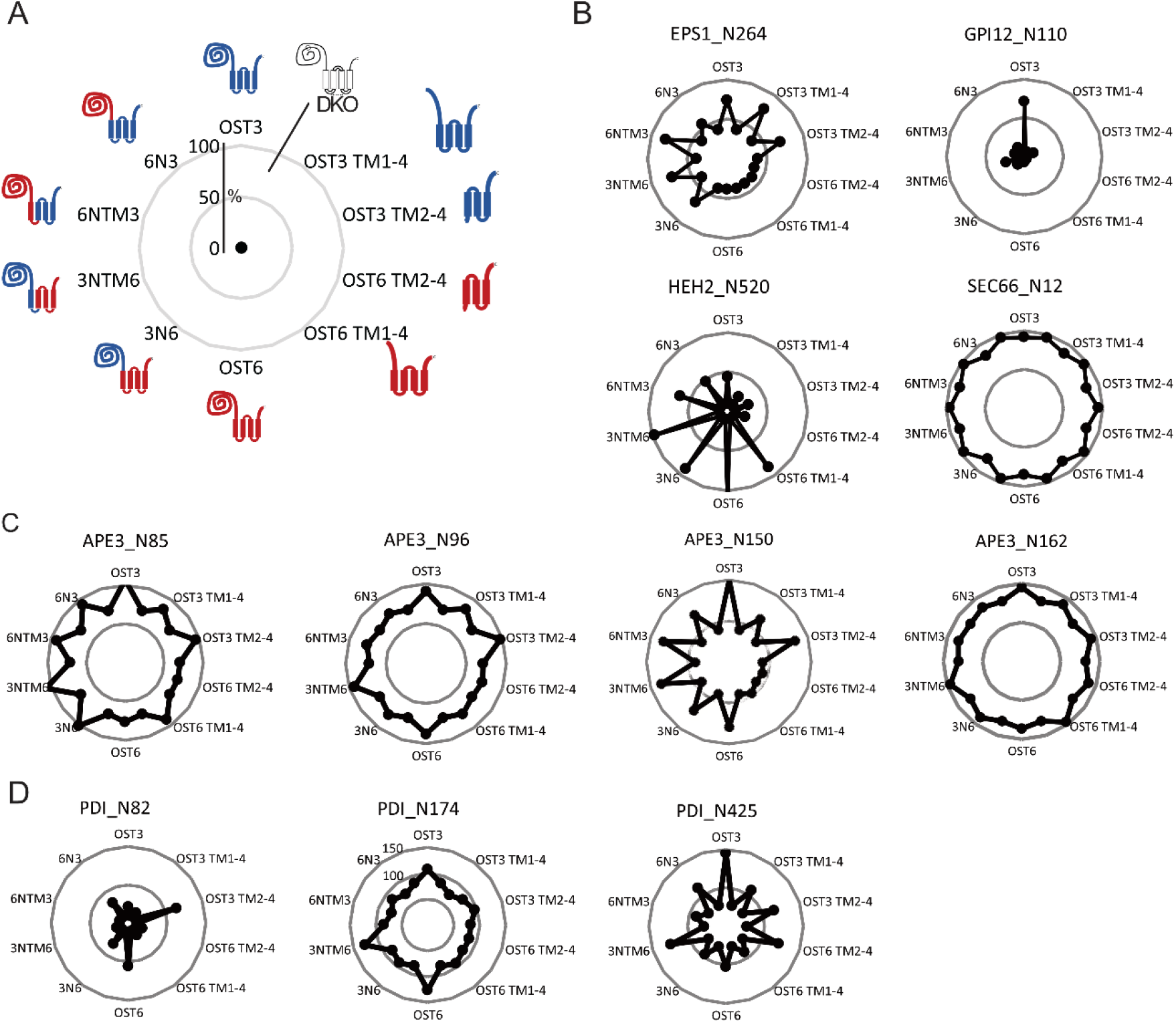
In vivo glycosylation site occupancy analysis of Ost3p and Ost6p complex derivatives relative to WT (in %). **(A)** Example of a star representation. In between the glycosylation occupancy values of the Ost3p/Ost6p derivative complexes the glycosylation occupancy of the OST complex lacking Ost3p and Ost6p (DKO) is shown. n=3. **(B)** Star representation of the glycosylation site occupancy analysis of GPI12_N110, EPS1_N264, HEH2_N520 and Sec66_N12, (**C**) of APE3_N85, APE3_N96, APE3_N150 and APE3_N162 and (**D**) of PDI_N82, PDI_N174 and PDI_N425 for the Ost3p and Ost6p complex derivatives.

## Discussion

Yeast OST complexes exist in two isoforms harbouring either the thioredoxin-containing subunit Ost3p or Ost6p^19^. Previous studies revealed differences in substrate specificity and glycosylation activity between the two isomeric complexes^18, 22, 26, 28^. The molecular basis of these observed differences, however, remained elusive. In this work, we combined *in vitro* and *in vivo* functional analysis with structural studies to pin-point the differences in Ost3p and Ost6p containing OST complexes.

A comparison of the here described Ost6p containing OST structure with a previously reported Ost3p containing structure^24^ suggested that structural changes are limited to the Ost3p/Ost6p subunits. The three TM helices (TM2 – TM4) that form the interaction interphase with the catalytic subunit Stt3p have identical positions in the complex. The major difference between the two homologous subunits was observed in the cytoplasm-exposed loop connecting TM3 and TM4. Within the obtained resolution, the structures of the other OST subunits, in particular that of the directly interacting catalytically active Stt3p, are indistinguishable between the two complexes. The highly similar structures and domain architecture of Ost3p and Ost6p therefore justified the analysis of hybrid subunits and N-terminal deletions as an experimental approach. Indeed, hybrid proteins as well as N-terminal truncations were stably integrated into the OST complex making a functional analysis in the *N-*glycosylation reaction possible.

Our OST structure harboured a co-purified dolichylphosphate in the LLO binding pocket rather than the supplied short-tailed LLO. Notably, the mammalian OST-B complex^17^ and the yeast O-mannosyltransferase Pmt1/Pmt2^29^ also contained a co-purified dolichylphosphate. Whether or not the binding of dolichylphosphate, a reaction product of the dolichylpyrophosphate phosphatase CWH8^30^ and of the Dol-P-Man/Dol-P-Glc-dependent glycosyltransferases^31^, represents a functional state in the *in vivo* reaction cycle of OST remains to be determined.

Our functional *in vitro* studies revealed that Ost3p and Ost6p modulate OST activity: the catalytic activity of the Ost3p containing complex was 2.8 times higher than that of the Ost6p complex, which, in turn was 2.5-times more active than the stable complex lacking the oxidoreductase subunit. Importantly, the binding of both the peptide and the LLO substrate was not different between the Ost3p and the Ost6p complex. The modulatory activity of Ost3p/Ost6p was mediated by the C-terminal transmembrane domain (TM2 – TM4/Figure2C) that interacted with the catalytic subunit Stt3p. We speculate that the interphases between these subunits affect a possible molecular dynamic of the Stt3p subunit differently during the catalytic cycle, resulting in an altered kinetic property of the enzyme. Our study highlights for the first time the importance of the C-terminal part of Ost3p/Ost6p while previous studies attributed functional differences between the two subunits primarily to their N-terminal thioredoxin domain^18, 21, 22, 32, 33^.

Interestingly, the N-terminal part encompassing the luminal oxidoreductase domain and the first transmembrane helix had a slight effect on the binding affinity towards the peptide and the LLO substrate: The presence of the thioredoxin domain and TM1 lowered the affinity toward the peptide and increased it towards the LLO. It is not obvious why the absence of the Ost3p/Ost6p domain (or the subunit altogether (DKO)) should affect the affinity towards the short peptide substrate. A possible explanation could be that the luminal domain of Ost3p/Ost6p influences the flexibility of EL5 of Stt3p. EL5 forms tight interactions with the peptide substrate, but undergoes large structural rearrangements in order to release the bulky glycosylation product^15, 16^. An apparent reduction in affinity for the short peptide substrate in the presence of the Ost3p/Ost6p luminal domain might be compensated for polypeptides of normal length through the possibility of forming additional interactions or mixed-disulphide bonds^18, 21^. Our experimental *in vitro* system using short acceptor peptides (solely bound by the Stt3p subunit^4, 16, 17, 24, 25^) was unable to address this thioredoxin function of the Ost3p and Ost6p.

The *in vivo* analysis of Ost3p/Ost6p N-terminal truncation and chimera mutants was based on the quantification of *N-*glycosylation efficiency for a number of *N-*glycosylation sites of different glycoproteins. We noted that an OST lacking the Ost3p/Ost6p subunit was able to glycosylate many of the sites to a level indistinguishable from the wild-type OST (Figure 3, sites at the right) whereas some sites depended to different degrees on an OST containing either Ost3p or Ost6p (Figure 3, sites at the left).

Our experimental approach allowed us to characterize the Ost3p/Ost6p dependency of site-specific glycosylation in more detail. OST devoid of the N-terminal thioredoxin domain and TM1 of either Ost3p or Ost6p were ideal tools to study the effect of a reduced OST activity on site-specific glycosylation (Figure 3B): for most of the sites, the slower acting OST6 derivative was as efficient as the faster acting OST3 construct. We concluded that for most of the sites analysed, OST activity was not limiting during biogenesis of the corresponding glycoprotein. However, several sites depended, to a different extend, on the presence of an N-terminal thioredoxin domain (supl. Fig. 7) and for most of these sites, the Ost3p-derived domain was more efficient than the Ost6p derivative (supl. Fig. 7E). This observation supports the previously proposed functionality of the Ost3p/Ost6p subunit to extend the substrate specificity of OST by forming mixed disulfides with neighbouring polypeptide sequences^18, 21^. However, the dual functionality of the Ost3p/Ost6p subunit, the modulatory function of the C-terminal transmembrane domain on OST activity and the oxidoreductase activity of the N-terminal thioredoxin domain, makes it difficult to define a specific function of this subunit in the site-specific glycosylation processes.

It is an important finding of our studies that the *N-*glycosylation sites we analysed reacted differently to the alterations of OST activity as reflected in the reactivity diagrams shown in Figure 5. This differential reactivity was site- and not protein specific (Figure 4). It seems unlikely that this site-specific differential response was primarily determined by the -N-X-S/T-sequon recognized by Stt3p. Instead, other processes in *N-*glycoprotein biogenesis might compete in a site-specific manner with *N-*linked protein glycosylation. The folding of protein domains into secondary structure elements can be such a competing event: glycosylated asparagine residues that are located in an alpha helix (e.g. PDI_N425 or KTR1_N120) have to be modified before the terminal folding: the alpha-helical structure is not compatible with substrate binding by OST. Therefore, glycosylation of such sites might become directly affected by OST activity that is modulated by the C-terminal transmembrane domain of Ost3p/Ost6p, or, alternatively, by the thioredoxin domain of Ost3p and Ost6p to keep the respective protein part unfolded via transient mixed disulphide formation^18, 21, 22, 32^ or by a combination of the two functionalities. In a given setting of the glycoprotein processing environment, the extend of these competitive effects define the glycosylation status in our experimental system. For some sites, low levels of such competitive events explain that even a drastically reduced OST activity (as is the case for a complex that lacks the Ost3p/Ost6p subunit) did not affect the steady state of glycosylation (e.g SEC66_N12, Figure 4). Alternatively, sites, such as GPI12_N110 heavily depend on an optimally active OST with a functional thioredoxin domain of Ost3p. Noteworthy, this site was not completely glycosylated in wild-type yeast cells (Figure 4), confirming the rate-limiting nature of *N-*glycosylation in the biogenesis of this glycoprotein.

Within the framework of our analysis, we can postulate a dual function of the thioredoxin domain of the Ost3p/Ost6p subunit: 1) it can delay the oxidative folding of protein domains by mixed disulphide formation and 2) it assigns a fast- or slow-acting OST to defined *N-*glycosylation sites. Both of these activities are site-specific due to the sequence specificity of the respective thioredoxin domains of the Ost3p/Ost6p subunit^34^. It is possible that a site-specific slow acting enzyme that forms a mixed disulphide with its protein substrate might favour the proper folding of an adjacent protein domain. Conversely, the more than two-fold excess of the fast (Ost3p) over the slow (Ost6p) acting OST ensures efficient glycosylation of the thioredoxin-independent sites.

The presence of a thioredoxin subunit is a specific property of the STT3B-containing OST complex. Interestingly, the STT3B containing OST of animals also exists in two forms that contain different thioredoxin subunits^35^. In addition, animals express the STT3A OST that associates with the translocon. This association is mediated by the DC2 protein homologous to the TM2-4 domain of Ost3p/Ost6p and provides the OST enzyme direct access to peptides in “status nascendi”, the optimal, unfolded state for substrate binding and *N-*glycosylation^17^. We therefore view these subunits (thioredoxin subunits; DC2) as activity-modulating OST components that (site-specifically) integrate OST activity at different time points into the folding pathway of *N-*glycoproteins.

Within the framework of our hypothesis, we view eukaryotic OST as one module in ER-localized glycoprotein biogenesis with a regulated and adaptable activity towards different polypeptide substrates. (Regulated) glycosylation per se affects the folding of protein domains but OST itself can modulate folding via auxiliary components such as Ost3p and Ost6p. Individual glycoproteins have evolved in such a specific modification and folding environment, where OST is only one of several functionalities. Accordingly, site-specific glycosylation is not solely dependent of OST activity (mediated by one specific subunit), but rather the result of an interplay of the polypeptide with OST functionalities, different protein modification pathways^35^ and the folding machinery in the endoplasmic reticulum.

## Materials and methods

### Expression and purification of yeast OST for cryo-EM studies

To study the Ost6p containing OST complex a previously described yeast strain (MAT α his3Δ1 leu2Δ0 lys2Δ0 ura3Δ0 arg4Δ0 ost3:LEU2MX6 OST4-1D4::kanMX6 YEp352-OST6) was used^24^. For protein purification, 9 L of yeast culture were grown to OD_600nm_ of 4.0 at 30 °C in standard synthetic dropout medium lacking uracil. All subsequent steps were carried out at 4°C or on ice. Cells were harvested by centrifugation, washed with dH_2_O and resuspended in lysis buffer (50 mM HEPES pH 7.5, 150 mM NaCl and 1 mM MgCl_2_) containing protease inhibitors (2 tablets EDTA-free protease inhibitor cocktail (Roche), 2.6 mg/L aprotinin, 5 mg/L leupeptin, 1 mg/L pepstatin, 2 mM benzamidine HCl, 1 mM phenylmethylsulfonyl fluoride (PMSF)) and 5 mg/L Dnase I. Glass beads were added to the cells and cell lysis was performed by vigorously mixing. After removing unbroken cells by centrifuging for 20 min at 3,000 xg, microsomal membranes were pelleted by centrifuging for 45 min at 142,000 xg. Membranes were resuspended in 10 volumes of membrane solubilization buffer (50 mM HEPES pH 7.5, 500 mM NaCl, 10% (v/v) glycerol, 1% (w/v) N-dodecyl-beta-D-maltopyranoside (DDM, Anatrace), 0.2% (w/v) cholesteryl hemisuccinate (CHS, Anatrace), 1 mM ethylenediaminetetraacetic acid (EDTA), 1 mM ethyleneglycoltetraacetic acid (EGTA)) and incubated for 1.5 hrs while rotating. A second ultracentrifugation step was carried out to remove unsolubilized membranes and the supernatant was applied onto Rho-1D4 antibody-coupled sepharose beads (University of British Columbia). After 3 hrs of incubation, beads were washed five times with 5 column volumes of each wash buffer and 2 column volumes purification buffer: wash buffer I (25 mM HEPES pH 7.5, 500 mM NaCl, 0.05% (w/v) DDM, 0.01% (w/v) CHS, 1 mM EDTA, 1 mM EGTA); wash buffer II (25 mM HEPES pH 7.5, 500 mM NaCl, 0.03% (w/v) DDM, 0.006% (w/v) CHS, 0.1% (w/v) digitonin (EMD Millipore)); wash buffer III (25 mM HEPES pH 7.5, 500 mM NaCl, 0.1%(w/v) digitonin) and purification buffer (25 mM HEPES pH 7.5, 150 mM NaCl, 0.1% (w/v) digitonin, 1 mM MgCl_2_, 1 mM MnCl_2_). The beads were incubated over night with purification buffer containing 500 mg/L 1D4 peptide to elute the protein. The protein was concentrated and loaded onto a Superose 6 column (GE Healthcare) pre-equilibrated in purification buffer and peak fractions were pooled and concentrated to 5-6 mg/mL.

### Cryo-EM sample preparation and data collection

The freshly prepared protein was mixed with 500 μM Dol20-PP-GlcNAc_2_^21^ and 150 μM inhibitory peptide (see below). 3.5 μL of sample were applied onto glow discharged Quantifoil holey carbon grids (R 1.2/1.3, 300 mesh, copper), blotted for 4 s and flash-frozen in a mixture of liquid ethane and propane cooled by liquid nitrogen using a Vitrobot Mark IV (FEI) with the environmental chamber set to 4 °C and a humidity of 100%. Data was collected on a 300 keV Titan Krios (FEI) electron microscope equipped with a Quantum-LS energy filter and a K2 camera (Gatan) using EPU (Thermo Fisher Scientific). Automated data acquisition was performed using super-resolution mode at a nominal magnification of 165,000x, corresponding to a super-resolution pixel size of 0.42 Å. The defocus was varied between -0.5 μm and -2.5 μm and 6 s movies containing 30 frames were collected using a cumulative dose of ∼67 electrons/Å^2^ (Supplementary Table I).

### Image processing

Movies were imported and processed in Relion 3.0^36^ unless stated otherwise. Motion-correction was carried out using the MotionCor2 frame alignment software^37^ and the non-dose-weighted and dose-weighted averages were binned by a factor of 2. GCTF^38^ was used to estimate the CTF of non-dose-weighted images. Automated, template-free particle picking was performed using Gautomatch (K. Zhang, MRC LMB). Several rounds of 2D and 3D classification were performed to sort particles. 3D refinement using soft masks, followed by particle polishing, CTF refinement and another round of 3D refinement finally yielded a 3D reconstruction at a resolution of 3.58 Å. This map contained density of the detergent micelle and the Ost6p luminal domain, which was better visible after low pass filtering using a Gaussian filter (see Gaussian filtered map). A further 3D refinement step using a tight mask, excluding the detergent micelle and the Ost6p luminal domain, was performed and yielded a 3D reconstruction at 3.46 Å resolution. A local resolution-filtered map was generated using a B-factor of -80 Å^2^ (High-resolution map, Supplementary Figure 1).

### Model building and refinement

The previously published Ost3p containing complex structure^24^ (PDB 6EZN) was docked into the high-resolution cryo-EM map using UCSF Chimera^39^. Relion 3.0 mrc files were converted into mtz format using the phenix.map_to_structure_factors command^40^. Model building was performed in COOT ^41^. First, the Ost3p subunit, not present in the Ost6p containing complex, was removed. Then the transmembrane helices TM2-4 of Ost6p (aa214-aa332) were built *de novo* (chain C) and the more flexible TM1 of Ost6p (aa189-aa212) was modeled using a polyAla sequence (chain I). The polyAla residues were renamed to polyUNK for model deposition. The density in the LLO binding site was interpreted as a Dol55-P. Small shifts of the transmembrane helices in relation to the ER luminal part and some loop regions where manually corrected. Several phosphatidylethanolamine and digitonin molecules were built. Phenix.real_space_refine^17^ was used to refine the resulting model against the local resolution filtered map. The corresponding refinement statistics are summarized in Supplementary Table I. Maps and structural models were visualized using UCSF Chimera^39^ and ChimeraX^42^

### Determination of the apparent IC_50_ value

The apparent IC_50_ value of the inhibitory peptide tetramethylrhodamine-YA(Dab)ATS-NH_2_ (Dab: 2,4-diaminobutanoic acid, JPT Peptide Technologies) was determined using an *in vitro* glycosylation assay. 10 μL reaction mixtures contained 0.1 μM OST, 35 μM Dol20-PP-GlcNAc_2_^21^, 1 mM MnCl_2_ 25 μM acceptor peptide (tetrametylrhodamine-DANYTK-NH_2_, JPT Peptide Technologies) and 500, 250, 100, 50, 25, 10, 5, 2.5 or 0 μM inhibitory peptide. Samples were incubated at 30 °C and 1 μL aliquots were taken after 0, 10, 20 and 30 min and mixed with 49 μL of 1x SDS Laemmli buffer. Glycopeptides were analyzed by Tricine SDS-PAGE^43^. Fluorescently labeled peptides were detected using a Typhoon Trio Plus imager (GE Healthcare) and band intensities were quantified using the ImageJ^44^. Initial turnover rates were calculated using linear regression. The resulting rates were then plotted against the inhibitory peptide concentration and data was fitted using nonlinear regression in GraphPad PRISM7.

### Yeast strains, growth conditions and plasmid cloning

All yeast strains and plasmids used in this study are listed in supplementary table III and IV. The yeast strains (YG889 derivatives and WT strain) were grown at 30 °C and 180 rpm^26, 27^. The pRS426 plasmids were created using Ost6p and Ost3p genomic DNA for amplifying the two parts of the Ost3p/Ost6p chimera and truncations using their native promotor and terminator. The two parts of the mutant genes were fused using Fusion-PCR, before inserted via BamHI/ SacI sites into a pRS426 plasmid^45^. The plasmids were transformed into a YG889 yeast strain with an additional deletion in ARG4, expressing a 1D4-tagged version of Ost4p^27, 28, 45^.

### Expression and purification of yeast OST for *in vitro* analysis

The Ost3p, Ost6p, the eight Ost3p/Ost6p derivative and the double knock-out containing strains were cultivated in triplicates of three liters each in a standard synthetic dropout medium lacking uracil to an OD_600_ of 2-4 at 30 °C 180 rpm. The OST complexes were purified as described^20^. In brief the harvested cells were centrifuged at 4000 rpm 10 min 4°C, before once washed with water and resuspended in cold HEPES buffer containing protease inhibitors (50 mM HEPES pH 7.5, 150 mM NaCl, 1 mM MgCl_2_, 1 mM PMSF, 20 tablets/L complete EDTA-free protease inhibitor cocktail (Roche)). Glass beads were added to the cells and cell lysis was performed by vigorously mixing in a bead beaker. After removing unbroken cells by centrifuging for 15 min at 1500x g, microsomal membranes were pelleted by centrifuging for 45 min at 50,000x g. The membranes were resuspended by douncing in 35% glycerol in lysis buffer, afterwards frozen in liquid nitrogen before stored at -80 °C.

The membranes were solubilized in solubilization buffer (50 mM HEPES pH 7.5, 150 mM NaCl, 1 mM MgCl2, 1 mM MnCl2, 10% (v/v) glycerol, 1 mM PMSF, 20 tablets/L complete EDTA-free protease inhibitor cocktail (Roche), 0.05 mg/mL DnaseI (from bovine pancreas), 1% (w/v) (DDM (Anatrace) and 0.2% (w/v) CHS) in a ratio of 1:10 (membranes (g) : buffer (mL)) for 1.5 hrs at 4°C and 1.25 rpm on a rotation wheel. Afterwards the membranes were centrifuged at 50,000 xg for 45 min at 4°C, before the supernatant was mixed with Rho-1D4 antibody (University of British Columbia) coupled beads equilibrated in purification buffer (50 mM HEPES, pH 7.5, 150 mM NaCl, 1 mM MgCl_2_, 1 mM MnCl_2_, 10% (v/v) glycerol, 0.03% (w/v) DDM and 0.006% (w/v) CHS). After 3 hrs of incubation on a rotating wheel, the 1D4 antibody-coupled beads were loaded on a 15 mL protino column and washed two times with purification buffer. To elute the OST complex the beads were incubated for 2 hrs at 4 °C with purification buffer enriched with 0.5 mg/mL 1D4 peptide (GenScript Corp.). The flow-through was collected in a 100 kDa cutoff filter column (Amicon Centrifugal Filter Device), concentrated and further purified via size-exclusion chromatography using a Superose 6 Increase 10/300 GL column (GE Healthcare) pre-equilibrated with purification buffer, and at flow rate of 0.3 mL/min to remove aggregates and unbound 1D4-peptide. The peak eluting around 14 mL was concentrated using a 100 kDa cutoff filter column (Amicon Centrifugal Filter Device) and the protein concentration was measured using BCA assay (Pierce, Thermo Fisher Scientific). The aliquoted OST complexes were shock frozen in liquid nitrogen before stored at -80 °C.

### *In vitro* glycosylation assay for kinetic analysis of the OST mutant complexes

TAMRA (5-Carboxytetramethylrhodamin)-labeled peptide (amino acid sequence: YANATS) with an amide group at the C-terminus and 5-TAMRA at the N-terminus were synthesized by WatsonBio Sciences. Lyophilized peptides were dissolved in dimethyl sulfoxide (DMSO) to obtain a 3 mM stock solution, which was stored at -20 °C and diluted with ddH_2_O to perform reactions. The *in vitro* assay reactions were performed in 10 μL containing 10 mM MnCl_2_, 100 μM peptide-TAMRA-YANATS, 50 μM Dol20-PP-GlcNAc_2_, purified OST complex diluted in purification buffer (50 mM HEPES pH 7.5, 150 mM NaCl, 1 mM MgCl2, 1 mM MnCl2, 10% (v/v) glycerol, 0.03% (w/v) DDM and 0.006% (w/v) CHS) for 2.5 to 40 minutes. The reaction mixture was preincubated for 5 min at 30 °C and the reaction was started by adding 1 μL of peptide. At each time point 1 μL was taken out from the reaction plate and incubated with 50 μL stop solution (0,1% formic acid in 10% acetonitrile, 10 mM KPO_4_ pH 8,0) to stop the reaction. The plates were stored at -20 °C until analyzed by reverse phase chromatography using a UPLC Dionex UltiMate 3000 with an Accucore 150-C18 100 ⨯ 2.1 mm 2.6 μm column (ThermoFisher Scientific). 1 μL sample (2 pmol of peptide) was injected and measured via isocratic elution using a certain proportion of acetonitrile in 10 mM KPO4 pH 8.0 buffer. The glycopeptide and peptide were separated using a flow rate of 0.7 mL/min. The glycopeptide and peptide were detected using the TAMRA-fluorescence (excitation wavelength: 546 nm; emission wavelength: 579 nm).

The UPLC profiles were analyzed using Chromeleon 7 (Dionex) by integrating the glycopeptide and peptide peak. The amount of glycosylation was calculated by dividing the glycopeptide peak area divided by the peak area of both peaks. Time points in the linear phase of the reaction were taken and fitted using GraphPad Prism 8 by linear regression to determine the initial rates of the reaction and by non-linear regression using the Michealis-Menten equation for determination of K_M_ and K_cat_ values.

To compare the Ost3p/Ost6p derivatives including the OST3, OST6 and DKO complex 0,05-0.3 μM purified complex and time points between 2.5 and 40 minutes were used. For all the reactions a concentration of 100 μM TAMRA-YANATS peptide and 50 μM Dol20-PP-GlcNAc_2_ was used. The slope of the linear regression of the initial rates were determined as described above. Statistical analysis of the turnover values was performed using a t-test with Prism GraphPad 8. P-values below 0.05 were considered as significant.

To determine the kinetic parameters of the TAMRA-labelled YANATS peptide, reactions using 0.1 μM (Ost3p derivative OST complexes) or 0.2 μM Ost6p derivative and DKO OST complexes), 100 μM Dol20-PP-GlcNAc_2_ and variable amount of 0-500 μM peptide for reaction time points between 2.5-25 (Ost3p derivative OST complexes) or 10-45 (Ost6p derivative and DKO OST complexes) minutes were performed. To determine the kinetic parameters of the LLO reactions using 0,1 μM (Ost3p derivative OST complexes) or 0,5 μM Ost6p derivative and DKO OST complexes), 25 μM TAMRA-YANATS and variable amount of 0-150 μM Dol20-PP-GlcNAc_2_ for reaction time points between 2.5 - 25 minutes were used. The turnover rates were determined using linear regression of the initial phase of the reaction and enzyme concentration in the reaction was taken into account. The data was fitted by non-linear regression using the Michaelis-Menten formula (for calculation of K_M_ and k_cat_ values) in Graph Prism7.

### Extraction of yeast OST for *in vivo* analysis

For sample preparation 50 OD_600_ units of Ost3p/Ost6p derivative containing YG889 yeast cells grown in standard synthetic dropout medium lacking uracil were mixed with 50 OD_600_ units of a previous described wild-type (WT) yeast strain grown in in standard synthetic dropout medium supplemented with 20 mg/L arginine (^13^C_6_) and lysine (^13^C_6_-^15^N_2_)^26^. Cells were collected and frozen in liquid nitrogen and stored at -80 °C. For MS peptide samples were prepared as described previously^46^. To summarize, cells were lysed with glass beads at 4 °C, followed by pelleting the microsomes by centrifugation. The pellet was dissolved in 0.1 M Na_2_CO_3_, 1 mM EDTA pH 11,3 incubated at 4 °C for 30 min and pelleted again. The membranes were then solubilized in 4% SDS, 50 mM DTT, 0,1 M Tris/HCl pH 7.6 and further processed via filter-assisted sample preparation. The membrane proteins were digested for 16 hrs at RT with endopeptidase LysC, followed by 6 hrs digestion at 37 °C with Trypsin. The peptides were further processed for 40 hrs at 37 °C with Endo-b-N-acetylglucosaminidase H in 50 mM pH 5.5 sodium acetate buffer. Afterwards, the samples were desalted using SepPack C18 columns, dried by Speed Vac centrifugation, diluted with 3% ACN 0,1% formic acid and analyzed via LC-ESI-MS/MS.

### Mass specterometry analysis of the OST mutant complexes

The purified OST complexes were prepared for mass spectrometry measurement as described above. The samples were measured on a Q ExactiveTM mass spectrometer (Thermo Scientific) coupled to a Waters Acquity UPLC M-Class system (Waters, Milford, USA) with a PicoviewTM nanospray source 500 model (New Objective) in DDA mode using an inclusion list^20^.

The tryptic samples were dissolved in 2% acetonitrile/0.1 % formic acid and loaded onto a trap column (Acclaim PepMap 100, 75 μm × 20 mm, 100 Å, 3 μm particle size). The separation was performed on a nanoACQUITY UPLC (BEH130 C18 column, 75 μm × 150 mm, 130 Å, 1.7 μm particle size) using a constant flow rate of 300 nL/min and a column temperature of 50 °C. A linear gradient from 1-35 % acetonitrile/0.1 % formic acid over 42 min, followed by a sharp increase to 98 % acetonitrile in 2 min was used, which was held for another 10 min isocratically. For the DDA analysis, one scan cycle comprised of a full scan MS survey spectrum was done, followed by up to 12 sequential HCD scans based on the intensity. For the HCD MS/MS spectra were recorded in the FT-Orbitrap at a resolution of 35,000 at 400 m/z, while the site-occupancy analysis was acquired by a full-scan MS spectra (400–2000 m/z) in the FT-Orbitrap at a resolution of 70,000 at 400 m/z. The HCD MS/MS spectra were performed with a target value of 1e^5^ by the collision energy setup at a normalized collision energy of 25. For glycosylation profiling analysis the full-scan MS spectra (800–2000 m/z) were acquired in the FT-Orbitrap at a resolution of 70,000 at 400 m/z, while HCD MS/MS spectra were recorded in the FT-Orbitrap at a resolution of 35,000 at 400 m/z with a target value of 5e5 by the collision energy setup at a normalized collision energy of 22^20^.

For the analysis of *in vivo* samples, iRT (indexed Retention Time) standard peptides (Biognosys) were added in a 1:40 ratio, and samples were subjected to a Q Exactive HF mass spectrometer (Thermo Scientific) with a PicoView nanospray source 500 model (New Objective), coupled to an Acquity UPLC M-class System (Waters) operating in trap/elute mode. iRT peptides allowed the determination of the peptide retention times using linear regression and confirmation of optimal operation during the measurement.

The mass spectrometry measurements were done as described^26. In general, 0.5^ μg sample was injected for each measurement and loaded on a 2G C18 trap column (180 mm by 20 mm, 100 Å, 5-mm particle size) and separated on a nanoACQUITY UPLC BEH130 C18 column (75 mm by 250 mm, 130 Å, 1.7 mm particle size) at a constant flow rate of 300 nL/min, with a column temperature of 50 °C and a linear gradient of 1 to 35% acetonitrile/0.1% formic acid over 90 minutes, followed by a sharp increase to 95% acetonitrile in 2 minutes. Afterwards the acetonitrile concentration was decreased with a linear gradient within 10 minutes to 5%. All the samples were analyzed using isolation lists containing iRT peptides and target precursor ions of the light and heavy labeled peptides at an Orbitrap resolution of 60000 as shown in supplementary table V. The Q-Exactive HF performes a MS^1^ scan with a range of 400-1500 m/z, followed by 12 MS^2^ scans. The full MS scan was performed with a target automatic gain control (AGC value) of 3e6 and a maximum injection time of 120 ms. The MS^2^ scans used a resolution for the Orbitrap of 60000 at 200 m/z, an AGC of 2e^5^ and a maximum injection time of 110 ms starting from a mass of 120 m/z. The HCD fragmentation was performed with a normalized collision energy of 28 using a scan window of 10 min for each peptide.

### Data processing and analysis of the obtained mass spectrometry data

The mass spectrometry results of the purified OST complexes the MS and MS/MS data were searched against the Swissprot database (version 201806) through Mascot engine (version 2.2) and analyzed as published previously^20^. For glycan profiling, the identification of each glycoform was confirmed by its corresponding MS/MS spectrum manually and analyzed as published previously^20^.

The PRM raw data from the scheduled PRM runs were imported into Skyline daily and the peaks were manually integrated. For peak identification the calculated retention time, a spectral library (dot product >0,9) and the co-elution of the light and heavy peak was used. The peak area of the light mutant peak (L) was divided by the peak area of the respective heavy wild-type strain peak (H) to obtain the ratio of the mutant strain verses the wild type strain using at least three identical transitions for each peak. To calculate the glycosylation occupancy, the L/H ratio of the glycopeptide was divided by the L/H average ratio of at least two non-sequon-peptides (have no glycosylation sequon) of the same protein. For normalization of the mixing errors between both strains the L/H ratios of all the peptides were divided by the L/H ratio of the stable expressed ribosomal proteins Rpl5 and Rps1a. The glycosylation occupancy is described compared to wild type.

## Supporting information

Supplementary Table VII

## Data availability

The cryo-EM maps have been deposited in the Electron Microscopy Data Bank (EMDB) under accession numbers EMD-XXXX and EMD-XXXX. The atomic structure coordinates of the OST6-containing OST complexes have been deposited in the Protein Data Bank (PDB) under the accession number XXXX. All other data needed to evaluate the conclusions of this paper are provided either in the main text of this manuscript or the supplementary materials.

## Acknowledgements

We thank the electron microscopy facility of ETH Zurich (ScopeM) and the functional genomic center Zurich (FGCZ) for providing equipment for cryo-EM sample preparation, data collection and the mass spectrometry measurements. We also thank Jérémy Boilevin and Jean-Louis Reymond for providing the LLO substrates.

## Funding

This work was supported by the Swiss National Science Foundation Sinergia program GlycoStart (CRSII5_173709). R.W. thanks the ETH postdoctoral fellowship program for support.

## Author contributions

J.N., R.W., K.P.L. and M.A. designed the study. R.W. screened and optimized cryo-EM grids. R.I., J.K. and R.W. collected cryo-EM data. R.W. processed cryo-EM data, built and refined the models. K.P.L. revised the model. J.N. synthesized the Ost3p/Ost6p derivative OST complexes. J.N., R.W. and J.E. performed chemo-enzymatic synthesis reactions, purified the OST complex and carried out *in vitro* glycosylation reactions. L.C. and M.A. analyzed glycopeptides by mass spectrometry. J.N. performed the *in vivo* experiments and analyzed the samples by mass spectrometry. J.N. analyzed the *in vitro* and *in vivo* experiments. J.N., R.W., M.A. and K.P.L. wrote the manuscript; all authors contributed to its revision.

## Abbreviations

ALG: asparagine linked glycosylation
CHS: cholesteryl hemisuccinate
Cryo-EM: cryo-electron microscopy
Dab: 2,4-diaminobutanoic acid
DDM: N-dodecyl-beta-D-maltopyranoside
DKO: double knock-out
Dol-P: dolichyl-phosphate
Dol-PP: dolichyl-pyrophosphate
EDTA: ethylenediaminetetraacetic acid
EGTA: ethyleneglycoltetraacetic acid
ER: endoplasmic reticulum
Glc: glucose
GlcNAc: N-acetylglucoseamine
LLO: lipid-linked oligosaccharide
Man: mannose
OST: oligosaccharyltransferase
ssOST: single-subunit oligosaccharyltransferase
TAMRA: tetramethylrhodamine
TM: transmembrane span

## Supplementary information

– Supplementary Figures 1-7
– Supplementary Tables I-VII

## Notes

### Competing Interest Statement

The authors have declared no competing interest.

