## Supplementary Figures and Tables for "Functional analysis of Ost3p and Ost6p containing yeast oligosaccharyltransferase*s*"

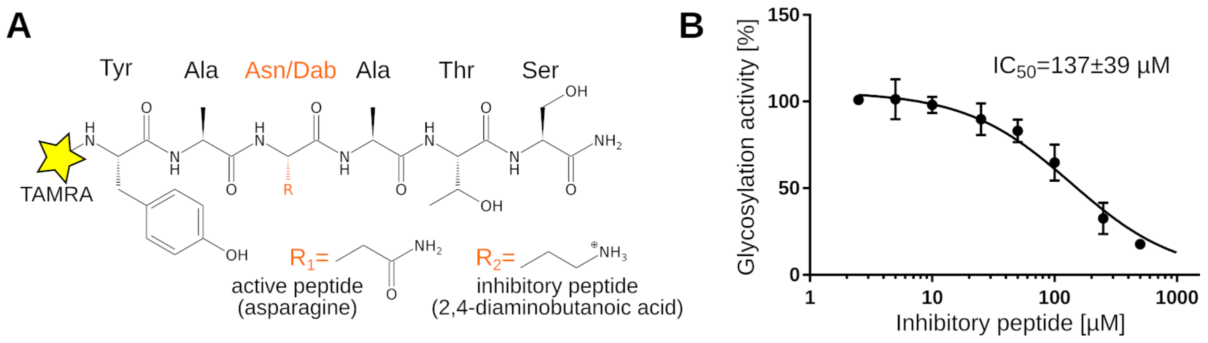

**Supplementary Figure 1:** Characterization of the inhibitory peptide.

(**A**) Structure of the wild-type peptide and an inhibitory version, which was used in the attempt to trap the complex in a ternary state. Peptides carry a N-terminal tetramethylrhodamine fluorophore (TAMRA) and a C-terminal amidation.

**(B)** $IC_{50}$ determination for the inhibitory peptide tetramethylrhodamine-YA(Dab)ATS using Dol20-PP-GlcNAc$_2$ as LLO and tetramethylrhodamine-DANYTK as peptide substrate. Experiments in technical triplicates (error bars indicate s.d.; n=3).

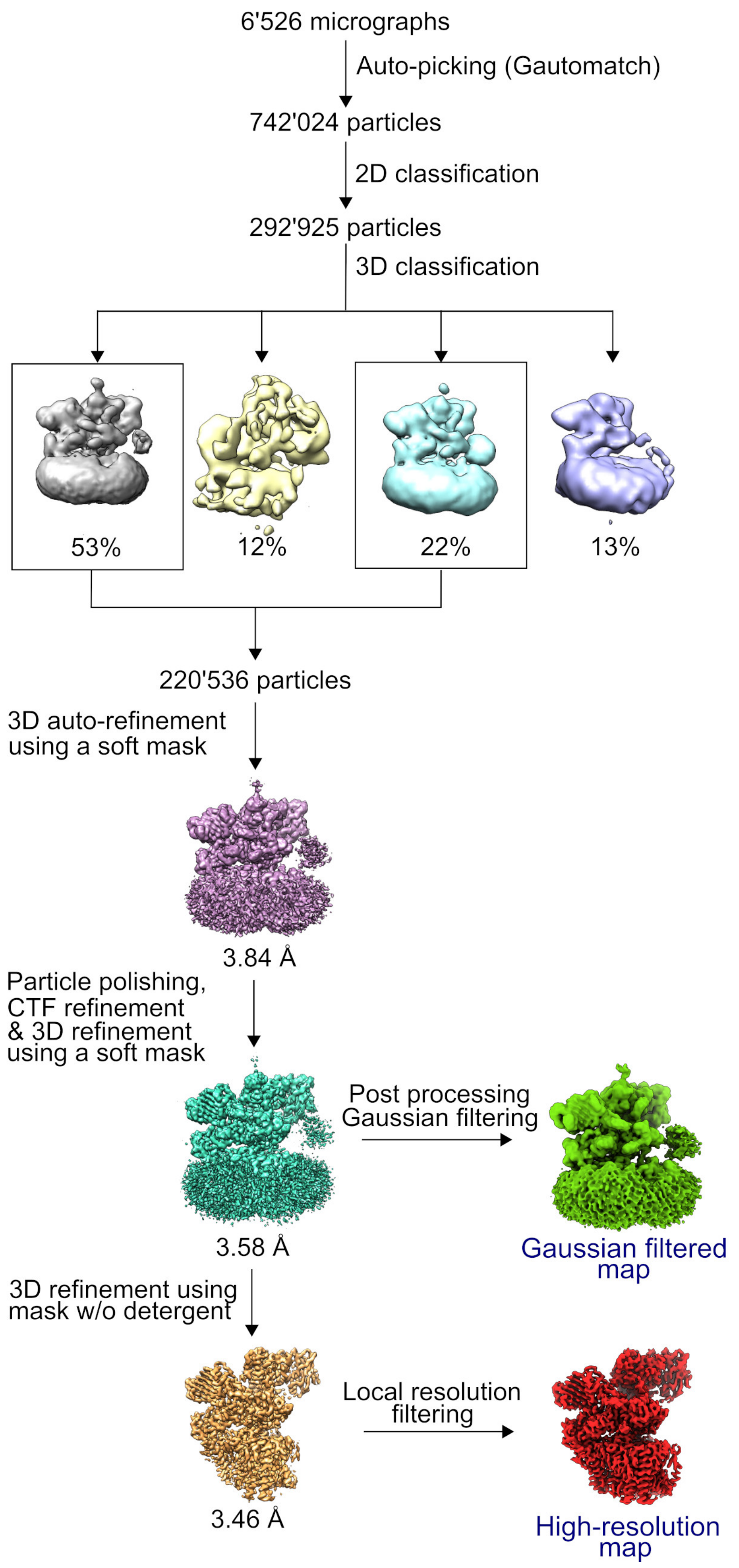

**Supplementary Figure 2:** Flow-chart of cryo-EM data processing procedure using Relion 3.0 resulting into two 3D reconstructions, a Gaussian filtered map containing the Ost6p luminal domain and the detergent micelle and a higher resolution map used for model building.

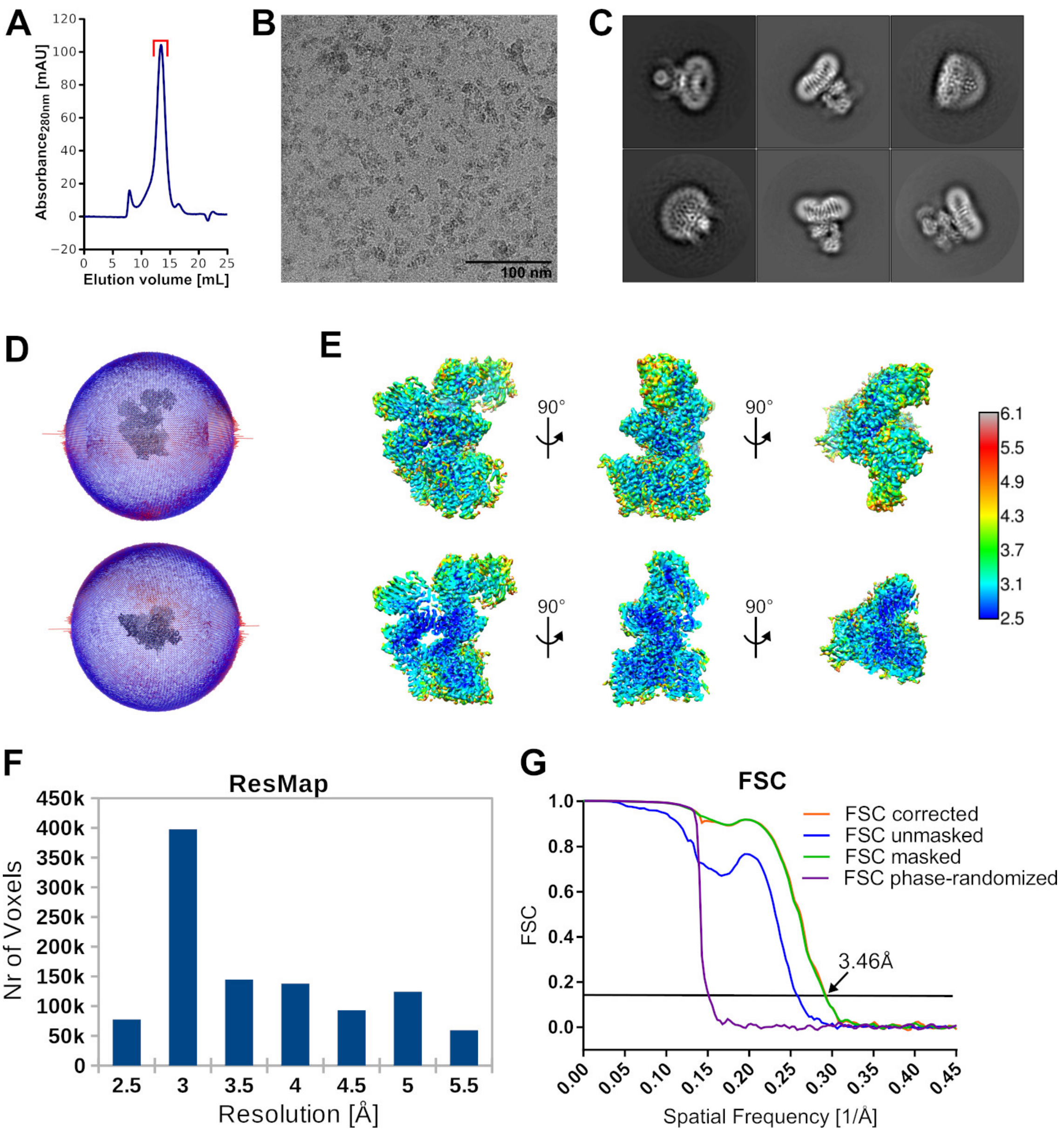

**Supplementary Figure 3:** Cryo-EM data quality assessment and resolution estimation for the high resolution map of yeast OST.

(**A**) Size exclusion chromatogram of the OST complex purification. Peak fractions which were used for EM grid preparation are indicated with a red bracket.

(**B**) Shown is a representative motion corrected and dose weighted micrograph.

(**C**) Selected class averages from 2D classification, sorted by descending number of particles.

(**D**) Plot of angular distribution showing the reconstruction from the side and top.

(**E**) ResMap estimated local resolution illustrated for a full map (top row) and sliced map (bottom row).

(**F**) Bar plot illustrating the number of voxels in the indicated resolution ranges.

(**G**) Relion 3.0 generated fourier shell correlation (FSC) curve indicating estimated resolutions based on the FSC = 0.143 criterion.

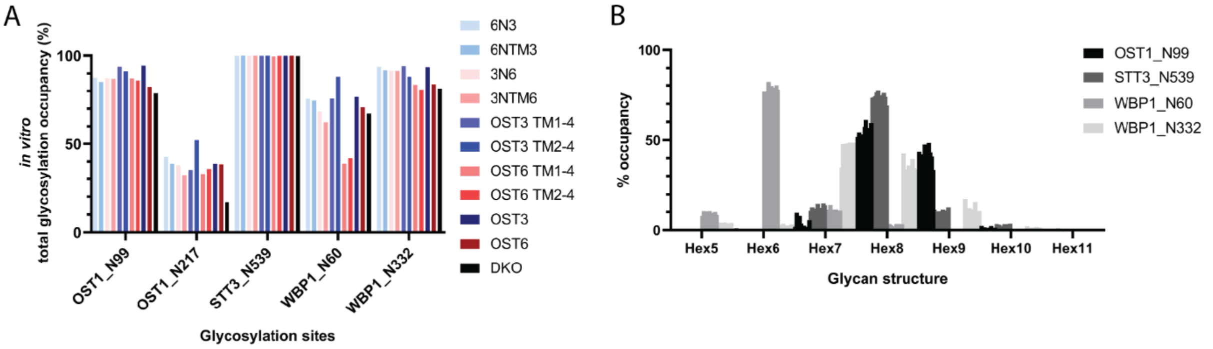

**Supplementary Figure 4:** *In vitro* glycosylation analysis. The Ost3p/Ost6p derivative containing OST complexes were purified via 1D4-tag based affinity enrichment and size exclusion chromatography.

(**A**) Glycosylation occupancy analysis of the three glycoproteins of the OST complex after trypsin digestion. PAbundance of glycosylated versus unglycosylated peptides was measured by Shotgun-MS.

(**B**) Occupancy of the glycan structures present on the glycosylation sites OST1_N99, STT3_N539, WBP1_N60 and WBP1_N332 of the purified derivative complexes. The order of bars is according to the list in (**A**).

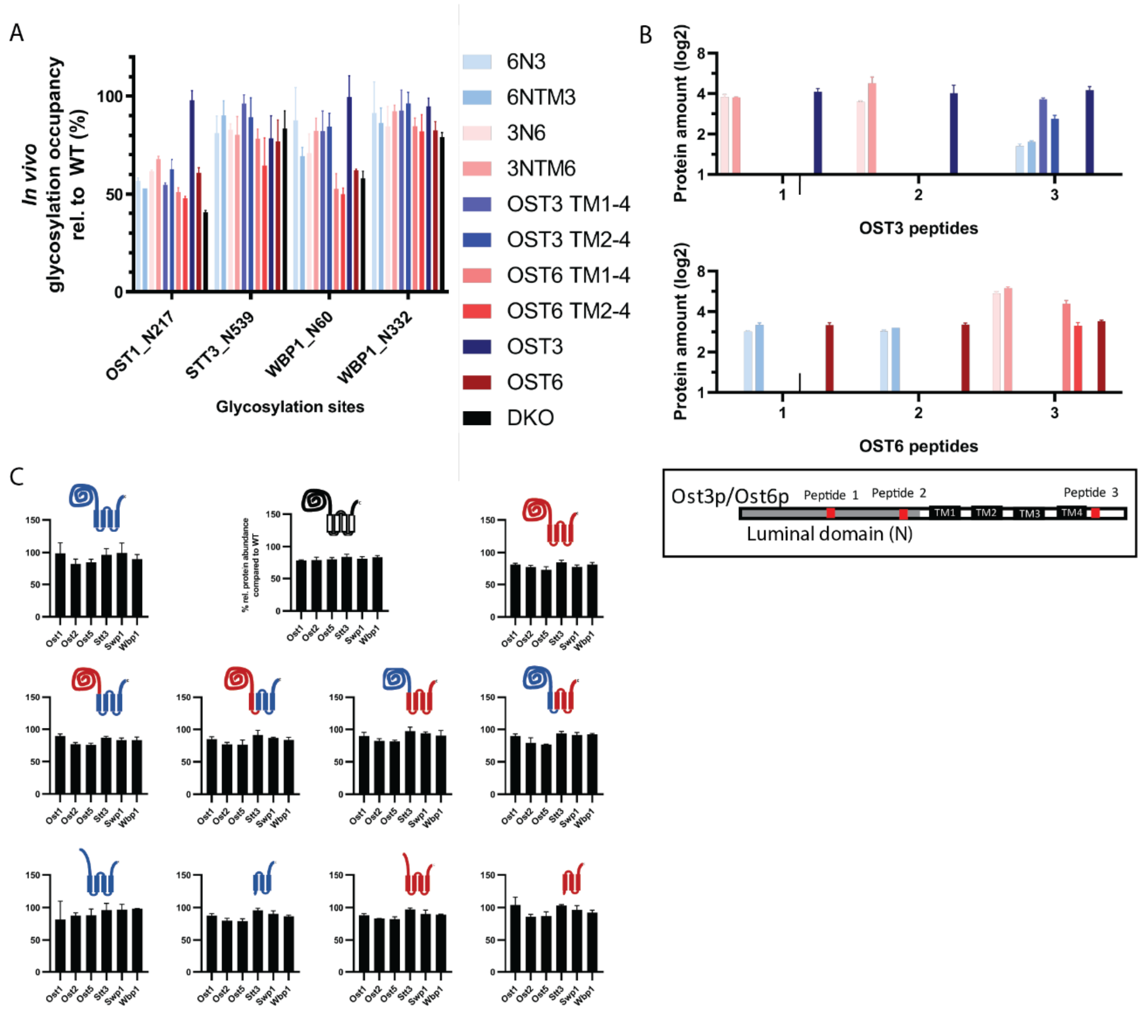

**Supplementary Figure 5:** *In vivo* steady-state analysis of the OST complex and glycosylation site occupancy analysis of the OST complex derivatives. (error bars indicate s.d.; n=3).

(**A**) Glycosylation occupancy analysis (%) of the three glycoproteins of the OST complex measured for the chimera and truncation mutant and OST3, OST6 and DKO strains relative to wild-type.

(**B**) Log2 Steady-state-level analysis of Ost3p and Ost6p derivative and DKO containing OST complex strains relative to wild-type strain. Location of the Ost3p and Ost6p peptides measured by PRM after digestion with trypsin and endoproteinase LysC is shown in the box. The bar diagrams show the detected peptides signals for Ost3p or Ost6p. Depending on which peptide is present in the Ost3p/Ost6p derivatives, the respective signal could be detected.

(**C**) Steady-state level analysis (%) of the six measurable OST proteins of the Ost3p/Ost6p derivative containing OST strains relative to the wild-type strain.

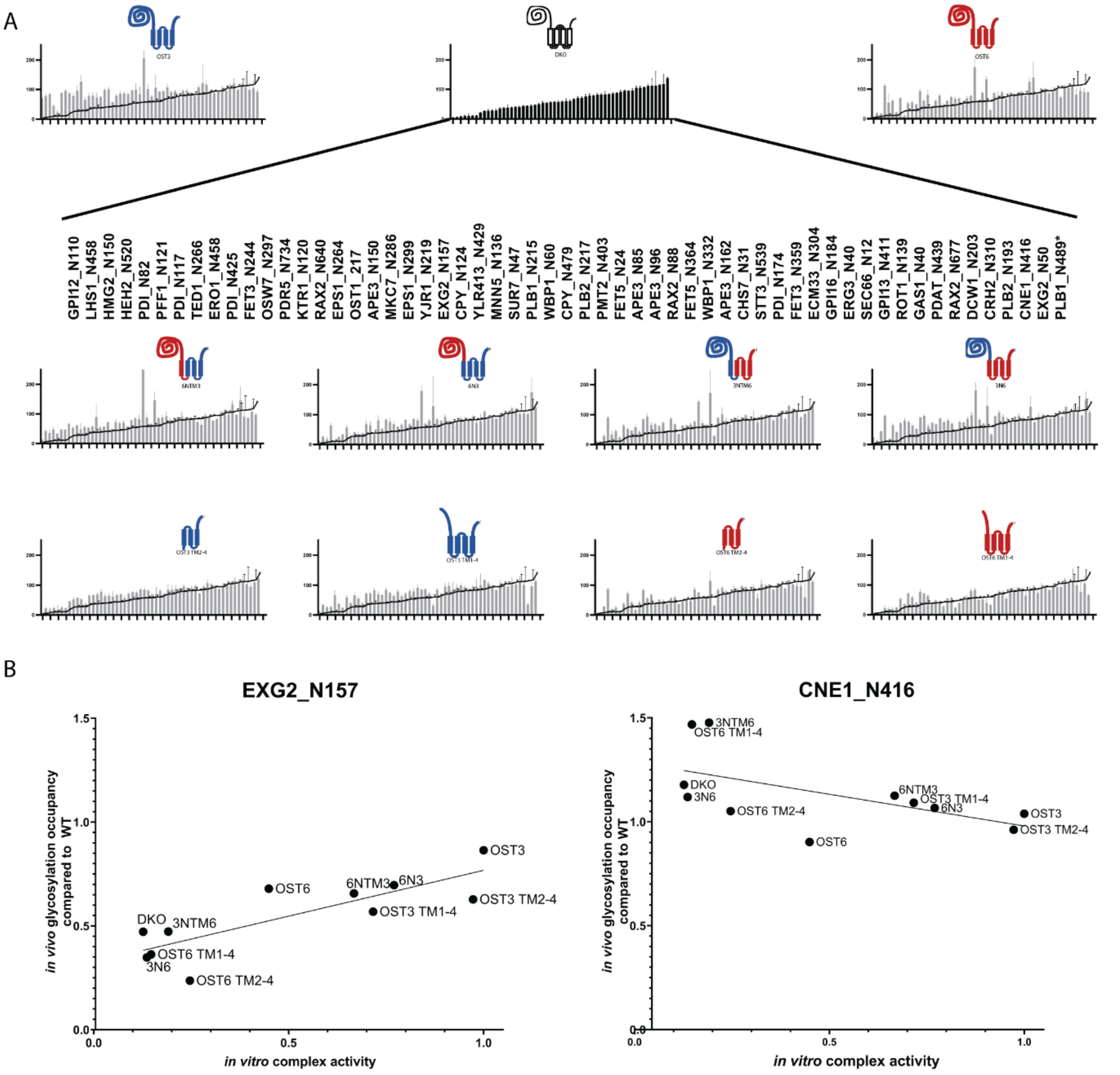

**Supplementary Figure 6:** *In vivo* glycosylation site occupancy analysis of Ost3p and Ost6p complex derivatives. (error bars indicate s.d.; n=3).

**(A)** Glycosylation occupancy analysis at different glycosylation sites of chimera and truncation, Ost3p, Ost6p and DKO containing OST strains y relative to wild-type cells. Glycosylation occupancy of the DKO was added as a line in every graph. Order of the glycosylation sites based on Ost3p/Ost6p dependency.

(**B**) Correlation of *in vitro* activity and *in vivo* glycosylation site occupancy of EXG2_N157 and CNE1_N416. Slope of the nonlinear regression of the glycosylation site occupancies of the Ost3p and Ost6p complex derivatives shows OST activity dependency.

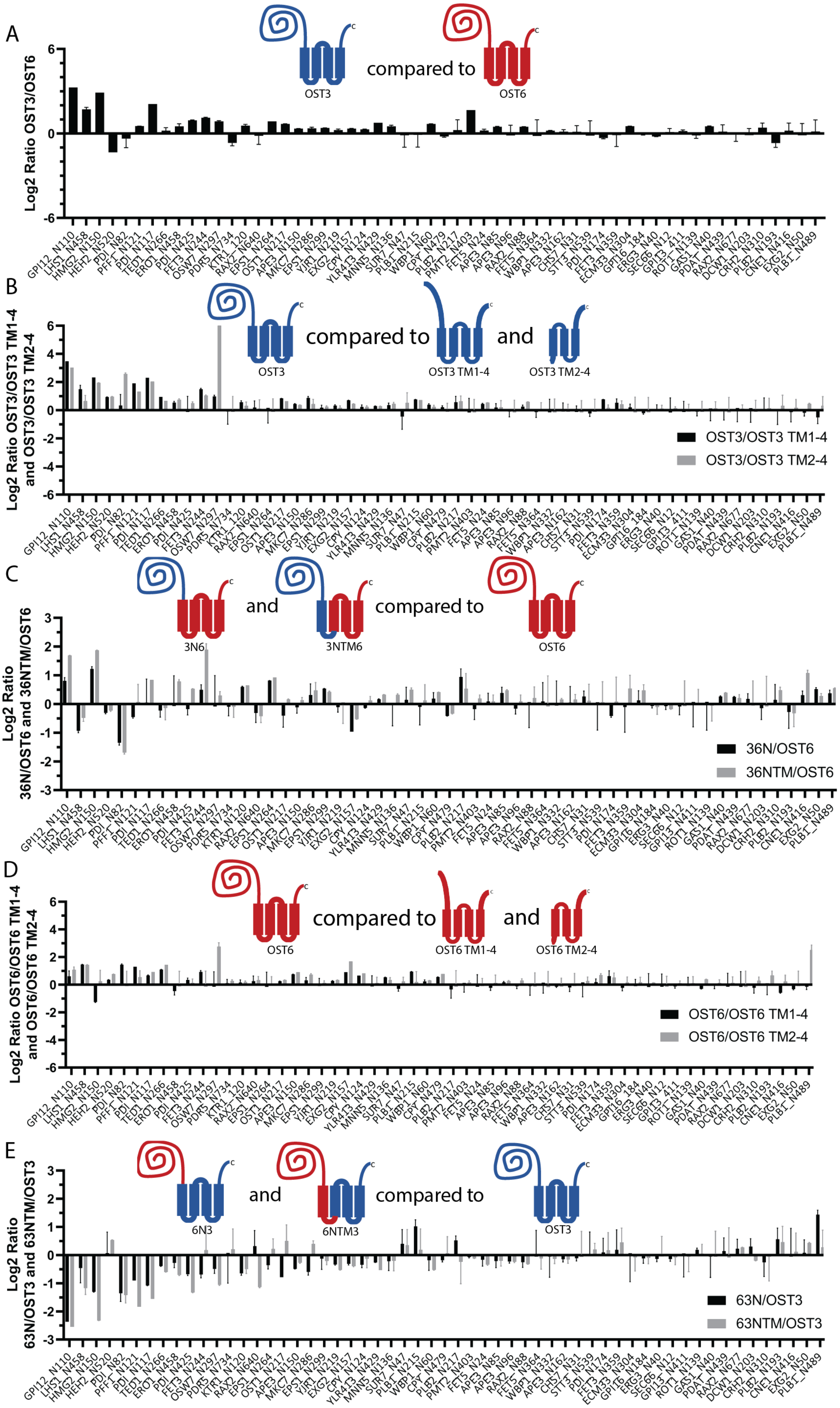
A
OST3
compared to
OST6
Log2 Ratio OST3/OST6
B
OST3
compared to
OST3 TM1-4
and
OST3 TM2-4
Log2 Ratio OST3/OST3 TM1-4 and OST3/OST3 TM2-4
OST3/OST3 TM1-4
OST3/OST3 TM2-4
C
3N6
and
3NTM6
compared to
OST6
Log2 Ratio 36N/OST6 and 36NTM/OST6
36N/OST6
36NTM/OST6
D
OST6
compared to
OST6 TM1-4
and
OST6 TM2-4
Log2 Ratio OST6/OST6 TM1-4 and OST6/OST6 TM2-4
OST6/OST6 TM1-4
OST6/OST6 TM2-4
E
6N3
and
6NTM3
compared to
OST3
Log2 Ratio 63N/OST3 and 63NTM/OST3
63N/OST3
63NTM/OST3
GPI12_N110 LHS1_N458 HMG2_N150 HEH2_N520 PDI_N82 PFF1_N121 PDI_N117 TED1_N266 ERO1_N458 PDI_N425 FET3_N244 OSW7_N297 PDR5_N734 KTR1_120 RAX2_N640 EPS1_N264 OST1_N217 APE3_N150 MKC7_N286 EPS1_N299 YIR1_N219 EXG2_N157 CPY_N124 YLR413_N429 MNN5_N136 SUR7_N47 PLB1_N215 WBP1_N60 CPY_N479 PLB2_N217 PMT2_N403 FET5_N24 APE3_N85 APE3_N96 RAX2_N88 FET5_N364 WBP1_N332 APE3_N162 CHS7_N31 STT3_N539 PDI_N174 FET3_N359 ECM33_N304 GPI16_184 ERG3_N40 SEC66_N12 GPI13_411 ROT1_N139 GAS1_N40 PDAT_N439 RAX2_N677 DCW1_N203 CRH2_N310 PLB2_N193 CNE1_N416 EXG2_N50 PLB1_N489

**Supplement Figure 7:** *In vivo* glycosylation occupancy comparison of Ost3p and Ost6p derivative containing OST complexes. (error bars indicate s.d.; n=3).

(**A**) Log2 of the glycosylation occupancy ratio of Ost3p containing OST complex relative to the Ost6p containing OST complex are shown. Positive bars represent glycosylation sites with a preference for the presence of the Ost3p subunit.

(**B**) and (**D**) Log2 of the glycosylation occupancy ratio of the Ost3 or Ost6 relative to its truncations. Positive bars represent glycosylation sites with a preference for the presence of the (**B**) Ost3p or (**D**) Ost6p thioredoxin domain.

(**C**) and (**E**) Log2 of the glycosylation occupancy ratio of the Ost3 thioredoxin domain containing chimera relative to Ost6p or Ost6 thioredoxin domain containing chimera relative to Ost3p. Positive bars represent glycosylation sites with a preference for the presence of the (**C**) Ost3 or (**E**) Ost6p thioredoxin domain.

**Supplementary Table I:** Cryo-EM data collection, processing and refinement statistics.

| **Data collection and processing** | **EMDB-12808, PDB 7OCI** |
|---|---|
| Microscope | FEI Titan Krios |
| Voltage (kV) | 300 |
| Camera | Gatan K2-Summit |
| Energy Filter | Gatan Quantum-LS |
| Magnification | 165'000 |
| Pixel size (Å) | 0.84 |
| Defocus range (µm) | -0.5 - -2,5 |
| Electron exposure ($e^-$/Å$^2$) | 67 |
| Number of good micrographs | 6’526 |
| Initial number of particles | 724'024 |
| Final number of particles | 220’536 |
| Symmetrie imposed | no |
| Map resolution (Å) FSC threshold | 3.46 |
| Map sharpening B-factor (Å$^2$) | -80 |
| **Coordinate and B-factor refinement** | |
| Initial model used (PDB code) | 6EZN |
| Number of protein atoms (non-H) | 16'538 |
| Number of ligand atoms (non-H) | 539 |
| Mean B-factor protein atoms (Å$^2$) | 63.56 |
| Mean B-factor of non-protein atoms (Å$^2$) | 46.12 |
| RMSD bonds (Å) | 0.003 |
| RMSD bond angles (°) | 0.639 |
| Map CC (whole map) | 0.65 |
| Map CC (around atoms) | 0.87 |
| **Ramachandran plot** | |
| Favored (%) | 97.20 |
| Allowed (%) | 2.75 |
| Disallowed (%) | 0.05 |
| **Validation** | |
| Molprobity score | 1.56 |
| All-atom clashscore | 7.42 |
| Rotamer outliers (%) | 0.00 |

**Supplementary Table II:** Clustering of the *in vivo* measured subset of glycosylation sites according to their Ost3p and Ost6p complex derivatives preference.

Comparison of the *in vitro* velocity and *in vivo* glycosylation occupancy revealed an OST complex activity dependency for every glycosylation site. The cluster "reverse linear" is based on negative slope and an R2 of minimum 0.19. The cluster "no preference" shows no preference for any mutant. The cluster "less than 25% diff." shows less than 25% *in vivo* glycosylation occupancy difference between the OSt3p and Ost6p complex derivatives. The cluster "collinearity" has a slope of minimum 0.27 and a R2 of minimum 0.45. The cluster "OST3 FL" is separated by a minimum 10% higher *in vivo* glycosylation occupancy of the OST3 full-length mutant. The cluster "OST6" shows a preference of the mutants having the OST6 transmembrane spans (TM). APE3_N150 and PLB2_N217 do not fit in any cluster. n=3.

| Glycosylation site | Cluster | Slope of nonlin-ear fit | $R^2$ | OST complex activity depended |
|---|---|---|---|---|
| CNE1_N416 | reverse linear | -0.3053 | 0.3261 | Yes |
| RAX2_N677 | reverse linear | -0.0909 | 0.1901 | Yes |
| ERG3_N40 | reverse linear | -0.1088 | 0.2869 | Yes |
| APE3_N85 | no preference | 0.1231 | 0.1055 | No |
| CPY_N124 | no preference | 0.2652 | 0.3683 | No |
| CPY_N479 | no preference | 0.1472 | 0.2450 | No |
| CRH2_N310 | no preference | 0.0098 | 0.0011 | No |
| DCW1_N203 | no preference | 0.1313 | 0.1421 | No |
| EPS1_N299 | no preference | 0.0914 | 0.0625 | No |
| ERO1_N458 | no preference | 0.0285 | 0.0019 | No |
| EXG2_N50 | no preference | -0.0879 | 0.0207 | No |
| FET5_N24 | no preference | 0.0066 | 0.0001 | No |
| GAS1_N40 | no preference | 0.0941 | 0.1060 | No |
| KTR1_N120 | no preference | 0.0119 | 0.0006 | No |
| MKC7_N286 | no preference | 0.2479 | 0.2633 | No |
| OSW7_N297 | no preference | 0.1084 | 0.1301 | No |
| PDAT_N439 | no preference | -0.0155 | 0.0038 | No |
| PLB2_N193 | no preference | -0.1854 | 0.0793 | No |
| RAX2_N640 | no preference | 0.2043 | 0.1657 | No |
| STT3_N539 | no preference | 0.0979 | 0.1659 | No |
| SUR7_N47 | no preference | 0.1009 | 0.1142 | No |
| TED1_N266 | no preference | 0.2556 | 0.2412 | No |
| APE3_N96 | Less than 25% diff. | 0.1261 | 0.2564 | No |
| APE3_N162 | Less than 25% diff. | 0.0105 | 0.0055 | No |
| CHS7_N31 | Less than 25% diff. | -0.0659 | 0.1810 | No |
| FET5_N364 | Less than 25% diff. | 0.0309 | 0.0296 | No |
| GPI13_N411 | Less than 25% diff. | 0.0776 | 0.3804 | No |
| GPI16_N184 | Less than 25% diff. | 0.0454 | 0.0737 | No |
| ROT1_N139 | Less than 25% diff. | -0.0256 | 0.0337 | No |
| SEC66_N12 | Less than 25% diff. | 0.0109 | 0.0082 | No |

| | | | | |
|---|---|---|---|---|
| WBP1_N332 | Less than 25% diff. | 0.1472 | 0.5837 | No |
| ECM33_N304 | collinearity | 0.2773 | 0.4617 | Yes |
| EXG2_N157 | collinearity | 0.4413 | 0.6746 | Yes |
| MNN5_N136 | collinearity | 0.3400 | 0.5368 | Yes |
| PMT2_N403 | collinearity | 0.6947 | 0.7439 | Yes |
| RAX2_N88 | collinearity | 0.2392 | 0.5321 | Yes |
| WBP1_N60 | collinearity | 0.3614 | 0.5806 | Yes |
| YJR1_N219 | collinearity | 0.2234 | 0.4810 | Yes |
| YLR413_N429 | collinearity | 0.3358 | 0.6603 | Yes |
| EPS1_N264 | Part of OST3 | 0.2846 | 0.2922 | No |
| FET3_N244 | OST3 | 0.4151 | 0.2534 | No |
| FET3_N359 | OST3 | 0.3039 | 0.2980 | No |
| GPI12_N110 | OST3 | 0.2855 | 0.2580 | No |
| HMG2_N150 | OST3 | 0.2826 | 0.2038 | No |
| LHS1_N458 | OST3/ OST3 TM | 0.5345 | 0.7216 | No |
| OST1_N217 | OST3 | 0.2644 | 0.3730 | No |
| PDI_N117 | OST3 | 0.4004 | 0.3496 | No |
| PDI_N425 | OST3 | 0.2823 | 0.2583 | No |
| PFF1_N121 | OST3 | 0.2528 | 0.1390 | No |
| HEH2_N520 | OST6 TM spans | -0.3504 | 0.1183 | No |
| PDR5_N734 | OST6 TM spans | -0.0635 | 0.0165 | No |
| APE3_N150 | OST3 TM or OST6 | 0.3632 | 0.4044 | No |
| PLB2_N217 | All full-length | -0.2614 | 0.0378 | No |

**Supplementary Table III:** Yeast strains used in this study. Related to Experimental Procedures.

| Name | Genotype | Source |
|---|---|---|
| | *MAT α his3Δ1 leu2Δ0 lys2Δ0 ura3Δ0 arg4Δ0 ost3:LEU2MX6 OST4-1D4::kanMX6* YEp352-*OST6* | 18 |
| WT | *MATα his3Δ1 leu2Δ0 lys2Δ0 ura3Δ0 Δarg4Δ0 1D4-ost4:NatMX6* | 26, modified in this study |
| YG889 "DKO" | *MATa ade2-101 his3Δ200 ura3-52 tyr1 Δarg4 ΔOST6::KanMX ΔOST3::HIS3 1D4-ost4:NatMX6* | 27, modified in this study |
| YG889 "36N" | *MATa ade2-101 his3Δ200 ura3-52 tyr1 Δarg4 ΔOST6::KanMX ΔOST3::HIS3 1D4-ost4:NatMX6 pRS426 36N)* | This study |
| YG889 "36NTM" | *MATa ade2-101 his3Δ200 ura3-52 tyr1 Δarg4 ΔOST6::KanMX ΔOST3::HIS3 1D4-ost4:NatMX6 pRS426 (36NTM)* | This study |
| YG889 "63N" | *MATa ade2-101 his3Δ200 ura3-52 tyr1 Δarg4 ΔOST6::KanMX ΔOST3::HIS3 1D4-ost4:NatMX6 pRS426 (63N)* | This study |
| YG889 "63NTM" | *MATa ade2-101 his3Δ200 ura3-52 tyr1 Δarg4 ΔOST6::KanMX ΔOST3::HIS3 1D4-ost4:NatMX6 pRS426 (63NTM)* | This study |
| YG889 "OST3 TM1-4" | *MATa ade2-101 his3Δ200 ura3-52 tyr1 Δarg4 ΔOST6::KanMX ΔOST3::HIS3 1D4-ost4:NatMX6 pRS426 (OST3 TM1-4)* | This study |
| YG889 "OST3 TM2-4" | *MATa ade2-101 his3Δ200 ura3-52 tyr1 Δarg4 ΔOST6::KanMX ΔOST3::HIS3 1D4-ost4:NatMX6 pRS426 (OST3 TM2-4)* | This study |
| YG889 "OST6 TM1-4" | *MATa ade2-101 his3Δ200 ura3-52 tyr1 Δarg4 ΔOST6::KanMX ΔOST3::HIS3 1D4-ost4:NatMX6 pRS426 (OST6 TM1-4)* | This study |
| YG889 "OST6 TM2-4" | *MATa ade2-101 his3Δ200 ura3-52 tyr1 Δarg4 ΔOST6::KanMX ΔOST3::HIS3 1D4-ost4:NatMX6 pRS426 (OST6 TM2-4)* | This study |
| YG889 "OST3" | *MATa ade2-101 his3Δ200 ura3-52 tyr1 Δarg4 ΔOST6::KanMX ΔOST3::HIS3 1D4-ost4:NatMX6 pRS426 (OST3)* | This study |

| | | |
|---|---|---|
| YG889 "OST6" | *MATa ade2-101 his3Δ200 ura3-52 tyr1 Δarg4 ΔOST6::KanMX ΔOST3::HIS3 1D4-ost4:NatMX6 pRS426 (OST6)* | This study |

**Supplementary Table IV:** Plasmids used in this study

| Name | Gene | Souce |
|---|---|---|
| YEp-OST6 | *OST6* | 47 |
| pRS426 36N | *36N* | This study |
| pRS426 36NTM | *36NTM* | This study |
| pRS426 63N | *63N* | This study |
| pRS426 63NTM | *63NTM* | This study |
| pRS426 OST3 TM1-4 | *OST3 TM1-4* | This study |
| pRS426 OST3 TM1-4 | OST3 TM2-4 | This study |
| pRS426 OST6 TM1-4 | OST6 TM1-4 | This study |
| pRS426 OST6 TM2-4 | OST6 TM2-4 | This study |
| pRS426 OST3 | *OST3* | This study |
| pRS426 OST6 | *OST6* | This study |

**Supplementary Table V:** Ost3p/Ost6p derivatives

| OST3 and OST6 derivatives | Promotor | Front part of the construct | Back part of the construct |
|---|---|---|---|
| OST3 | -363 to -1 bp | AA 1-351 of OST3 | - |
| OST3 TM1-4 | -363 to -1 bp | AA 1-43 of OST3 | AA 177-351 of OST3 |
| OST3 TM2-4 | -363 to -1 bp | AA 1-6 of OST3 | AA 204-351 of OST3 |
| OST6 | -369 to -1 bp | AA 1-333 of OST6 | - |
| OST6 TM1-4 | -369 to -1 bp | AA 1-24 | AA 180-333 of OST6 |
| OST6 TM2-4 | -369 to -1 bp | AA 1-6 | AA 209-333 of OST6 |
| 63N | -369 to -1 bp | AA 1-188 of OST6 | AA 186-351 of OST3 |
| 63NTM | -369 to -1 bp | AA 1-214 of OST6 | AA 216-351 of OST3 |
| 36N | -363 to -1 bp | AA 1-185 of OST3 | AA 189-333 of OST6 |
| 36NTM | -363 to -1 bp | AA 1-215 of OST3 | AA 215-333 of OST6 |
| DKO | - | Empty pRS426 plasmid | - |

### Supplementary table VI

PRM MS assay for quantitative profiling of N-linked glycosylation machinery in yeast.

See separate document

### Supplementary table VII

Site-specific N-glycosylation occupancy analysis of various OST mutant strains

See separate document
