## Supplementary Table VI for "Functional analysis of Ost3p and Ost6p containing yeast oligosaccharyltransferase*s*"

**Supplementary Table VI.** PRM MS assay for quantitative profiling of N-linked glycosylation machinery in yeast. PRM assay R=60000

| UniProt ID | Prote-in | Peptide Modified Sequence | Precursor Charge | Isotope Label Type | Pre-cursor Mz | Frag ment Ion | Frag ment Mz | Nor-mali-zed Reten-tion Time |
|---|---|---|---|---|---|---|---|---|
| P236321 | Rpl5 | VFLDIGLQR | 2 | light | 530.8111 | y8 | 961.5465 | 63.75 |
| P236321 | Rpl5 | VFLDIGLQR | 2 | light | 530.8111 | y7 | 814.4781 | 63.75 |
| P236321 | Rpl5 | VFLDIGLQR | 2 | light | 530.8111 | y6 | 701.3941 | 63.75 |
| P236321 | Rpl5 | VFLDIGLQR | 2 | light | 530.8111 | y5 | 586.3671 | 63.75 |
| P236321 | Rpl5 | VFLDIGLQR | 2 | light | 530.8111 | y4 | 473.2831 | 63.75 |
| P236321 | Rpl5 | VFLDIGLQR | 2 | light | 530.8111 | b4 | 475.2551 | 63.75 |
| P236321 | Rpl5 | VFLDIGLQR | 2 | heavy | 533.8212 | y8 | 967.5667 | 63.75 |
| P236321 | Rpl5 | VFLDIGLQR | 2 | heavy | 533.8212 | y7 | 820.4983 | 63.75 |
| P236321 | Rpl5 | VFLDIGLQR | 2 | heavy | 533.8212 | y6 | 707.4142 | 63.75 |
| P236321 | Rpl5 | VFLDIGLQR | 2 | heavy | 533.8212 | y5 | 592.3873 | 63.75 |
| P236321 | Rpl5 | VFLDIGLQR | 2 | heavy | 533.8212 | y4 | 479.3032 | 63.75 |
| P236321 | Rpl5 | VFLDIGLQR | 2 | heavy | 533.8212 | b4 | 475.2551 | 63.75 |
| P236321 | Rpl5 | FPGWDFETEEIDPELLR | 2 | light | 1046.997 | y11 | 1343.669 | 85.09 |
| P236321 | Rpl5 | FPGWDFETEEIDPELLR | 2 | light | 1046.997 | y10 | 1214.626 | 85.09 |
| P236321 | Rpl5 | FPGWDFETEEIDPELLR | 2 | light | 1046.997 | y5 | 627.3824 | 85.09 |
| P236321 | Rpl5 | FPGWDFETEEIDPELLR | 2 | light | 1046.997 | y3 | 401.2871 | 85.09 |
| P236321 | Rpl5 | FPGWDFETEEIDPELLR | 2 | light | 1046.997 | b3 | 302.1499 | 85.09 |
| P236321 | Rpl5 | FPGWDFETEEIDPELLR | 2 | light | 1046.997 | b5 | 603.2562 | 85.09 |
| P236321 | Rpl5 | FPGWDFETEEIDPELLR | 2 | heavy | 1050.007 | y11 | 1349.689 | 85.09 |
| P236321 | Rpl5 | FPGWDFETEEIDPELLR | 2 | heavy | 1050.007 | y10 | 1220.646 | 85.09 |
| P236321 | Rpl5 | FPGWDFETEEIDPELLR | 2 | heavy | 1050.007 | y5 | 633.4026 | 85.09 |
| P236321 | Rpl5 | FPGWDFETEEIDPELLR | 2 | heavy | 1050.007 | y3 | 407.3072 | 85.09 |
| P236321 | Rpl5 | FPGWDFETEEIDPELLR | 2 | heavy | 1050.007 | b3 | 302.1499 | 85.09 |
| P236321 | Rpl5 | FPGWDFETEEIDPELLR | 2 | heavy | 1050.007 | b5 | 603.2562 | 85.09 |
| P33442 | Rps1a | VISEILTK | 2 | light | 451.7815 | y7 | 803.4873 | 42.46 |
| P33442 | Rps1a | VISEILTK | 2 | light | 451.7815 | y6 | 690.4032 | 42.46 |
| P33442 | Rps1a | VISEILTK | 2 | light | 451.7815 | y4 | 474.3286 | 42.46 |

| | | | | | | | | |
|---|---|---|---|---|---|---|---|---|
| P33442 | Rps1a | VISEILTK | 2 | light | 451.7815 | y3 | 361.2445 | 42.46 |
| P33442 | Rps1a | VISEILTK | 2 | light | 451.7815 | y2 | 248.1605 | 42.46 |
| P33442 | Rps1a | VISEILTK | 2 | heavy | 455.7886 | y7 | 811.5015 | 42.46 |
| P33442 | Rps1a | VISEILTK | 2 | heavy | 455.7886 | y6 | 698.4174 | 42.46 |
| P33442 | Rps1a | VISEILTK | 2 | heavy | 455.7886 | y4 | 482.3428 | 42.46 |
| P33442 | Rps1a | VISEILTK | 2 | heavy | 455.7886 | y3 | 369.2587 | 42.46 |
| P33442 | Rps1a | VISEILTK | 2 | heavy | 455.7886 | y2 | 256.1747 | 42.46 |
| P33442 | Rps1a | EVQGSTLAQLTSK | 2 | light | 681.3672 | y11 | 1133.616 | 42.84 |
| P33442 | Rps1a | EVQGSTLAQLTSK | 2 | light | 681.3672 | y10 | 1005.558 | 42.84 |
| P33442 | Rps1a | EVQGSTLAQLTSK | 2 | light | 681.3672 | y9 | 948.536 | 42.84 |
| P33442 | Rps1a | EVQGSTLAQLTSK | 2 | light | 681.3672 | y8 | 861.504 | 42.84 |
| P33442 | Rps1a | EVQGSTLAQLTSK | 2 | light | 681.3672 | y7 | 760.4563 | 42.84 |
| P33442 | Rps1a | EVQGSTLAQLTSK | 2 | heavy | 685.3743 | y11 | 1141.63 | 42.84 |
| P33442 | Rps1a | EVQGSTLAQLTSK | 2 | heavy | 685.3743 | y10 | 1013.572 | 42.84 |
| P33442 | Rps1a | EVQGSTLAQLTSK | 2 | heavy | 685.3743 | y9 | 956.5502 | 42.84 |
| P33442 | Rps1a | EVQGSTLAQLTSK | 2 | heavy | 685.3743 | y8 | 869.5182 | 42.84 |
| P33442 | Rps1a | EVQGSTLAQLTSK | 2 | heavy | 685.3743 | y7 | 768.4705 | 42.84 |
| P22146 | GAS1 | FFYSNN[+203.079373]GSQFYIR | 2 | light | 923.4258 | y10 | 1388.644 | 59.6 |
| P22146 | GAS1 | FFYSNN[+203.079373]GSQFYIR | 2 | light | 923.4258 | y9 | 1301.612 | 59.6 |
| P22146 | GAS1 | FFYSNN[+203.079373]GSQFYIR | 2 | light | 923.4258 | y7 | 870.4468 | 59.6 |
| P22146 | GAS1 | FFYSNN[+203.079373]GSQFYIR | 2 | light | 923.4258 | y6 | 813.4254 | 59.6 |
| P22146 | GAS1 | FFYSNN[+203.079373]GSQFYIR | 2 | light | 923.4258 | y5 | 726.3933 | 59.6 |
| P22146 | GAS1 | FFYSNN[+203.079373]GSQFYIR | 2 | light | 923.4258 | y4 | 598.3348 | 59.6 |
| P22146 | GAS1 | FFYSNN[+203.079373]GSQFYIR | 2 | light | 923.4258 | y3 | 451.2663 | 59.6 |
| P22146 | GAS1 | FFYSNN[+203.079373]GSQFYIR | 2 | heavy | 926.4358 | y10 | 1394.664 | 59.6 |
| P22146 | GAS1 | FFYSNN[+203.079373]GSQFYIR | 2 | heavy | 926.4358 | y9 | 1307.632 | 59.6 |
| P22146 | GAS1 | FFYSNN[+203.079373]GSQFYIR | 2 | heavy | 926.4358 | y7 | 876.467 | 59.6 |
| P22146 | GAS1 | FFYSNN[+203.079373]GSQFYIR | 2 | heavy | 926.4358 | y6 | 819.4455 | 59.6 |
| P22146 | GAS1 | FFYSNN[+203.079373]GSQFYIR | 2 | heavy | 926.4358 | y5 | 732.4135 | 59.6 |
| P22146 | GAS1 | FFYSNN[+203.079373]GSQFYIR | 2 | heavy | 926.4358 | y4 | 604.3549 | 59.6 |
| P22146 | GAS1 | FFYSNN[+203.079373]GSQFYIR | 2 | heavy | 926.4358 | y3 | 457.2865 | 59.6 |
| P22146 | GAS1 | DDPTWTVDLFNSYK | 2 | light | 850.8938 | y10 | 1272.626 | 76.22 |

| | | | | | | | | |
|---|---|---|---|---|---|---|---|---|
| P22146 | GAS1 | DDPTWTVDLFNSYK | 2 | light | 850.8938 | y9 | 1086.547 | 76.22 |
| P22146 | GAS1 | DDPTWTVDLFNSYK | 2 | light | 850.8938 | y8 | 985.4989 | 76.22 |
| P22146 | GAS1 | DDPTWTVDLFNSYK | 2 | light | 850.8938 | y7 | 886.4305 | 76.22 |
| P22146 | GAS1 | DDPTWTVDLFNSYK | 2 | light | 850.8938 | y6 | 771.4036 | 76.22 |
| P22146 | GAS1 | DDPTWTVDLFNSYK | 2 | light | 850.8938 | y5 | 658.3195 | 76.22 |
| P22146 | GAS1 | DDPTWTVDLFNSYK | 2 | light | 850.8938 | y4 | 511.2511 | 76.22 |
| P22146 | GAS1 | DDPTWTVDLFNSYK | 2 | heavy | 854.9009 | y10 | 1280.64 | 76.22 |
| P22146 | GAS1 | DDPTWTVDLFNSYK | 2 | heavy | 854.9009 | y9 | 1094.561 | 76.22 |
| P22146 | GAS1 | DDPTWTVDLFNSYK | 2 | heavy | 854.9009 | y8 | 993.5131 | 76.22 |
| P22146 | GAS1 | DDPTWTVDLFNSYK | 2 | heavy | 854.9009 | y7 | 894.4447 | 76.22 |
| P22146 | GAS1 | DDPTWTVDLFNSYK | 2 | heavy | 854.9009 | y6 | 779.4178 | 76.22 |
| P22146 | GAS1 | DDPTWTVDLFNSYK | 2 | heavy | 854.9009 | y5 | 666.3337 | 76.22 |
| P22146 | GAS1 | DDPTWTVDLFNSYK | 2 | heavy | 854.9009 | y4 | 519.2653 | 76.22 |
| P22146 | GAS1 | MTDYFAC[+57.021464]GDDDVK | 2 | light | 768.8027 | y11 | 1304.51 | 45.38 |
| P22146 | GAS1 | MTDYFAC[+57.021464]GDDDVK | 2 | light | 768.8027 | y10 | 1189.483 | 45.38 |
| P22146 | GAS1 | MTDYFAC[+57.021464]GDDDVK | 2 | light | 768.8027 | y9 | 1026.42 | 45.38 |
| P22146 | GAS1 | MTDYFAC[+57.021464]GDDDVK | 2 | light | 768.8027 | y8 | 879.3513 | 45.38 |
| P22146 | GAS1 | MTDYFAC[+57.021464]GDDDVK | 2 | light | 768.8027 | y7 | 808.3142 | 45.38 |
| P22146 | GAS1 | MTDYFAC[+57.021464]GDDDVK | 2 | light | 768.8027 | y6 | 648.2835 | 45.38 |
| P22146 | GAS1 | MTDYFAC[+57.021464]GDDDVK | 2 | light | 768.8027 | y5 | 591.262 | 45.38 |
| P22146 | GAS1 | MTDYFAC[+57.021464]GDDDVK | 2 | light | 768.8027 | y4 | 476.2351 | 45.38 |
| P22146 | GAS1 | MTDYFAC[+57.021464]GDDDVK | 2 | heavy | 772.8098 | y11 | 1312.524 | 45.38 |
| P22146 | GAS1 | MTDYFAC[+57.021464]GDDDVK | 2 | heavy | 772.8098 | y10 | 1197.497 | 45.38 |
| P22146 | GAS1 | MTDYFAC[+57.021464]GDDDVK | 2 | heavy | 772.8098 | y9 | 1034.434 | 45.38 |
| P22146 | GAS1 | MTDYFAC[+57.021464]GDDDVK | 2 | heavy | 772.8098 | y8 | 887.3655 | 45.38 |
| P22146 | GAS1 | MTDYFAC[+57.021464]GDDDVK | 2 | heavy | 772.8098 | y7 | 816.3284 | 45.38 |
| P22146 | GAS1 | MTDYFAC[+57.021464]GDDDVK | 2 | heavy | 772.8098 | y6 | 656.2977 | 45.38 |
| P22146 | GAS1 | MTDYFAC[+57.021464]GDDDVK | 2 | heavy | 772.8098 | y5 | 599.2762 | 45.38 |
| P22146 | GAS1 | MTDYFAC[+57.021464]GDDDVK | 2 | heavy | 772.8098 | y4 | 484.2493 | 45.38 |
| P46982 | MNN5 | LN[+203.079373]FSIPQR | 2 | light | 589.3142 | y7 | 861.4577 | 49.93 |
| P46982 | MNN5 | LN[+203.079373]FSIPQR | 2 | light | 589.3142 | y6 | 747.4148 | 49.93 |
| P46982 | MNN5 | LN[+203.079373]FSIPQR | 2 | light | 589.3142 | y5 | 600.3464 | 49.93 |

| | | | | | | | | |
|---|---|---|---|---|---|---|---|---|
| P46982 | MNN5 | LN[+203.079373]FSIPQR | 2 | light | 589.3142 | y4 | 513.3144 | 49.93 |
| P46982 | MNN5 | LN[+203.079373]FSIPQR | 2 | light | 589.3142 | y3 | 400.2303 | 49.93 |
| P46982 | MNN5 | LN[+203.079373]FSIPQR | 2 | light | 589.3142 | b3 | 375.2027 | 49.93 |
| P46982 | MNN5 | LN[+203.079373]FSIPQR | 2 | heavy | 592.3243 | y7 | 867.4779 | 49.93 |
| P46982 | MNN5 | LN[+203.079373]FSIPQR | 2 | heavy | 592.3243 | y6 | 753.4349 | 49.93 |
| P46982 | MNN5 | LN[+203.079373]FSIPQR | 2 | heavy | 592.3243 | y5 | 606.3665 | 49.93 |
| P46982 | MNN5 | LN[+203.079373]FSIPQR | 2 | heavy | 592.3243 | y4 | 519.3345 | 49.93 |
| P46982 | MNN5 | LN[+203.079373]FSIPQR | 2 | heavy | 592.3243 | y3 | 406.2504 | 49.93 |
| P46982 | MNN5 | LN[+203.079373]FSIPQR | 2 | heavy | 592.3243 | b3 | 375.2027 | 49.93 |
| P46982 | MNN5 | ADPWTLYHENR | 2 | light | 701.3309 | y7 | 932.4585 | 44.89 |
| P46982 | MNN5 | ADPWTLYHENR | 2 | light | 701.3309 | y5 | 718.3267 | 44.89 |
| P46982 | MNN5 | ADPWTLYHENR | 2 | light | 701.3309 | y4 | 555.2634 | 44.89 |
| P46982 | MNN5 | ADPWTLYHENR | 2 | light | 701.3309 | y9 | 608.2989 | 44.89 |
| P46982 | MNN5 | ADPWTLYHENR | 2 | heavy | 704.341 | y7 | 938.4786 | 44.89 |
| P46982 | MNN5 | ADPWTLYHENR | 2 | heavy | 704.341 | y5 | 724.3468 | 44.89 |
| P46982 | MNN5 | ADPWTLYHENR | 2 | heavy | 704.341 | y4 | 561.2835 | 44.89 |
| P46982 | MNN5 | ADPWTLYHENR | 2 | heavy | 704.341 | y9 | 611.309 | 44.89 |
| P46982 | MNN5 | EALFSGSEGIVTIGGGK | 2 | light | 811.4252 | y13 | 1161.611 | 62.43 |
| P46982 | MNN5 | EALFSGSEGIVTIGGGK | 2 | light | 811.4252 | y12 | 1074.579 | 62.43 |
| P46982 | MNN5 | EALFSGSEGIVTIGGGK | 2 | light | 811.4252 | y10 | 930.5255 | 62.43 |
| P46982 | MNN5 | EALFSGSEGIVTIGGGK | 2 | light | 811.4252 | y9 | 801.4829 | 62.43 |
| P46982 | MNN5 | EALFSGSEGIVTIGGGK | 2 | light | 811.4252 | y5 | 431.2613 | 62.43 |
| P46982 | MNN5 | EALFSGSEGIVTIGGGK | 2 | heavy | 815.4323 | y13 | 1169.625 | 62.43 |
| P46982 | MNN5 | EALFSGSEGIVTIGGGK | 2 | heavy | 815.4323 | y12 | 1082.593 | 62.43 |
| P46982 | MNN5 | EALFSGSEGIVTIGGGK | 2 | heavy | 815.4323 | y10 | 938.5397 | 62.43 |
| P46982 | MNN5 | EALFSGSEGIVTIGGGK | 2 | heavy | 815.4323 | y9 | 809.4971 | 62.43 |
| P46982 | MNN5 | EALFSGSEGIVTIGGGK | 2 | heavy | 815.4323 | y5 | 439.2755 | 62.43 |
| P37302 | APE3 | IISFN[+203.079373]LSDAETGK | 2 | light | 799.4014 | y9 | 934.4476 | 56.91 |
| P37302 | APE3 | IISFN[+203.079373]LSDAETGK | 2 | light | 799.4014 | y8 | 820.4047 | 56.91 |
| P37302 | APE3 | IISFN[+203.079373]LSDAETGK | 2 | light | 799.4014 | y7 | 707.3206 | 56.91 |
| P37302 | APE3 | IISFN[+203.079373]LSDAETGK | 2 | light | 799.4014 | y5 | 505.2617 | 56.91 |
| P37302 | APE3 | IISFN[+203.079373]LSDAETGK | 2 | light | 799.4014 | y3 | 305.1819 | 56.91 |

| | | | | | | | | |
|---|---|---|---|---|---|---|---|---|
| P37302 | APE3 | IISFN[+203.079373]LSDAETGK | 2 | heavy | 803.4085 | y9 | 942.4618 | 56.91 |
| P37302 | APE3 | IISFN[+203.079373]LSDAETGK | 2 | heavy | 803.4085 | y8 | 828.4189 | 56.91 |
| P37302 | APE3 | IISFN[+203.079373]LSDAETGK | 2 | heavy | 803.4085 | y7 | 715.3348 | 56.91 |
| P37302 | APE3 | IISFN[+203.079373]LSDAETGK | 2 | heavy | 803.4085 | y5 | 513.2759 | 56.91 |
| P37302 | APE3 | IISFN[+203.079373]LSDAETGK | 2 | heavy | 803.4085 | y3 | 313.1961 | 56.91 |
| P37302 | APE3 | IISFN[+203.079373]LSDAETGK | 3 | light | 598.948 | y10 | 1130.523 | 30.37 |
| P37302 | APE3 | IISFN[+203.079373]LSDAETGK | 3 | light | 598.948 | y6 | 730.3631 | 30.37 |
| P37302 | APE3 | IISFN[+203.079373]LSDAETGK | 3 | light | 598.948 | y5 | 567.2998 | 30.37 |
| P37302 | APE3 | IISFN[+203.079373]LSDAETGK | 3 | light | 598.948 | y3 | 373.2194 | 30.37 |
| P37302 | APE3 | IISFN[+203.079373]LSDAETGK | 3 | light | 598.948 | y8 | 471.725 | 30.37 |
| P37302 | APE3 | IISFN[+203.079373]LSDAETGK | 3 | heavy | 600.9547 | y10 | 1136.543 | 30.37 |
| P37302 | APE3 | IISFN[+203.079373]LSDAETGK | 3 | heavy | 600.9547 | y6 | 736.3832 | 30.37 |
| P37302 | APE3 | IISFN[+203.079373]LSDAETGK | 3 | heavy | 600.9547 | y5 | 573.3199 | 30.37 |
| P37302 | APE3 | IISFN[+203.079373]LSDAETGK | 3 | heavy | 600.9547 | y3 | 379.2395 | 30.37 |
| P37302 | APE3 | IISFN[+203.079373]LSDAETGK | 3 | heavy | 600.9547 | y8 | 474.7351 | 30.37 |
| P37302 | APE3 | IKVDDLN[+203.079373]ATAWDL-YR | 3 | light | 666.0056 | y11 | 1337.648 | 62.13 |
| P37302 | APE3 | IKVDDLN[+203.079373]ATAWDL-YR | 3 | light | 666.0056 | y9 | 1312.617 | 62.13 |
| P37302 | APE3 | IKVDDLN[+203.079373]ATAWDL-YR | 3 | light | 666.0056 | y9 | 1109.537 | 62.13 |
| P37302 | APE3 | IKVDDLN[+203.079373]ATAWDL-YR | 3 | light | 666.0056 | y8 | 995.4945 | 62.13 |
| P37302 | APE3 | IKVDDLN[+203.079373]ATAWDL-YR | 3 | light | 666.0056 | y7 | 924.4574 | 62.13 |
| P37302 | APE3 | IKVDDLN[+203.079373]ATAWDL-YR | 3 | light | 666.0056 | y6 | 823.4097 | 62.13 |
| P37302 | APE3 | IKVDDLN[+203.079373]ATAWDL-YR | 3 | light | 666.0056 | y5 | 752.3726 | 62.13 |
| P37302 | APE3 | IKVDDLN[+203.079373]ATAWDL-YR | 3 | heavy | 670.6837 | y11 | 1343.669 | 62.13 |
| P37302 | APE3 | IKVDDLN[+203.079373]ATAWDL-YR | 3 | heavy | 670.6837 | y9 | 1318.637 | 62.13 |
| P37302 | APE3 | IKVDDLN[+203.079373]ATAWDL-YR | 3 | heavy | 670.6837 | y9 | 1115.558 | 62.13 |
| P37302 | APE3 | IKVDDLN[+203.079373]ATAWDL-YR | 3 | heavy | 670.6837 | y8 | 1001.515 | 62.13 |
| P37302 | APE3 | IKVDDLN[+203.079373]ATAWDL-YR | 3 | heavy | 670.6837 | y7 | 930.4775 | 62.13 |
| P37302 | APE3 | IKVDDLN[+203.079373]ATAWDL-YR | 3 | heavy | 670.6837 | y6 | 829.4298 | 62.13 |
| P37302 | APE3 | IKVDDLN[+203.079373]ATAWDL-YR | 3 | heavy | 670.6837 | y5 | 758.3927 | 62.13 |
| P37302 | APE3 | SFAN[+203.079373]TTA-FALSPPVDGFVGK | 2 | light | 1115.057 | y12 | 1186.647 | 73.93 |
| P37302 | APE3 | SFAN[+203.079373]TTA-FALSPPVDGFVGK | 2 | light | 1115.057 | y10 | 1002.525 | 73.93 |
| P37302 | APE3 | SFAN[+203.079373]TTA-FALSPPVDGFVGK | 2 | light | 1115.057 | y9 | 915.4934 | 73.93 |

| | | | | | | | | |
|---|---|---|---|---|---|---|---|---|
| P37302 | APE3 | SFAN[+203.079373]TTA-FALSPPVDGFVGK | 2 | light | 1115.057 | y5 | 507.2926 | 73.93 |
| P37302 | APE3 | SFAN[+203.079373]TTA-FALSPPVDGFVGK | 2 | heavy | 1119.064 | y12 | 1194.661 | 73.93 |
| P37302 | APE3 | SFAN[+203.079373]TTA-FALSPPVDGFVGK | 2 | heavy | 1119.064 | y10 | 1010.54 | 73.93 |
| P37302 | APE3 | SFAN[+203.079373]TTA-FALSPPVDGFVGK | 2 | heavy | 1119.064 | y9 | 923.5076 | 73.93 |
| P37302 | APE3 | SFAN[+203.079373]TTA-FALSPPVDGFVGK | 2 | heavy | 1119.064 | y5 | 515.3068 | 73.93 |
| P37302 | APE3 | HTVATVGVPYK | 2 | light | 586.3271 | y10 | 1034.588 | 32.77 |
| P37302 | APE3 | HTVATVGVPYK | 2 | light | 586.3271 | y9 | 933.5404 | 32.77 |
| P37302 | APE3 | HTVATVGVPYK | 2 | light | 586.3271 | y8 | 834.472 | 32.77 |
| P37302 | APE3 | HTVATVGVPYK | 2 | light | 586.3271 | y5 | 563.3188 | 32.77 |
| P37302 | APE3 | HTVATVGVPYK | 2 | light | 586.3271 | y3 | 407.2289 | 32.77 |
| P37302 | APE3 | HTVATVGVPYK | 2 | heavy | 590.3342 | y10 | 1042.602 | 32.77 |
| P37302 | APE3 | HTVATVGVPYK | 2 | heavy | 590.3342 | y9 | 941.5546 | 32.77 |
| P37302 | APE3 | HTVATVGVPYK | 2 | heavy | 590.3342 | y8 | 842.4862 | 32.77 |
| P37302 | APE3 | HTVATVGVPYK | 2 | heavy | 590.3342 | y5 | 571.333 | 32.77 |
| P37302 | APE3 | HTVATVGVPYK | 2 | heavy | 590.3342 | y3 | 415.2431 | 32.77 |
| P33767 | WBP1 | LEYLDIN[+203.079373]STSTTVDLYDK | 2 | light | 1147.052 | y11 | 1229.59 | 63.58 |
| P33767 | WBP1 | LEYLDIN[+203.079373]STSTTVDLYDK | 2 | light | 1147.052 | y9 | 1041.51 | 63.58 |
| P33767 | WBP1 | LEYLDIN[+203.079373]STSTTVDLYDK | 2 | light | 1147.052 | y8 | 954.4779 | 63.58 |
| P33767 | WBP1 | LEYLDIN[+203.079373]STSTTVDLYDK | 2 | light | 1147.052 | y7 | 853.4302 | 63.58 |
| P33767 | WBP1 | LEYLDIN[+203.079373]STSTTVDLYDK | 2 | light | 1147.052 | y6 | 752.3825 | 63.58 |
| P33767 | WBP1 | LEYLDIN[+203.079373]STSTTVDLYDK | 2 | light | 1147.052 | y5 | 653.3141 | 63.58 |
| P33767 | WBP1 | LEYLDIN[+203.079373]STSTTVDLYDK | 2 | light | 1147.052 | y4 | 538.2871 | 63.58 |
| P33767 | WBP1 | LEYLDIN[+203.079373]STSTTVDLYDK | 2 | light | 1147.052 | y3 | 425.2031 | 63.58 |
| P33767 | WBP1 | LEYLDIN[+203.079373]STSTTVDLYDK | 2 | light | 1147.052 | b3 | 406.1973 | 63.58 |
| P33767 | WBP1 | LEYLDIN[+203.079373]STSTTVDLYDK | 2 | light | 1147.052 | b5 | 634.3083 | 63.58 |
| P33767 | WBP1 | LEYLDIN[+203.079373]STSTTVDLYDK | 2 | heavy | 1151.059 | y11 | 1237.604 | 63.58 |
| P33767 | WBP1 | LEYLDIN[+203.079373]STSTTVDLYDK | 2 | heavy | 1151.059 | y9 | 1049.524 | 63.58 |
| P33767 | WBP1 | LEYLDIN[+203.079373]STSTTVDLYDK | 2 | heavy | 1151.059 | y8 | 962.4921 | 63.58 |
| P33767 | WBP1 | LEYLDIN[+203.079373]STSTTVDLYDK | 2 | heavy | 1151.059 | y7 | 861.4444 | 63.58 |
| P33767 | WBP1 | LEYLDIN[+203.079373]STSTTVDLYDK | 2 | heavy | 1151.059 | y6 | 760.3967 | 63.58 |
| P33767 | WBP1 | LEYLDIN[+203.079373]STSTTVDLYDK | 2 | heavy | 1151.059 | y5 | 661.3283 | 63.58 |
| P33767 | WBP1 | LEYLDIN[+203.079373]STSTTVDLYDK | 2 | heavy | 1151.059 | y4 | 546.3013 | 63.58 |

| | | | | | | | | |
|---|---|---|---|---|---|---|---|---|
| P33767 | WBP1 | LEYLDIN[+203.079373]STSTTVDLYDK | 2 | heavy | 1151.059 | y3 | 433.2173 | 63.58 |
| P33767 | WBP1 | LEYLDIN[+203.079373]STSTTVDLYDK | 2 | heavy | 1151.059 | b3 | 406.1973 | 63.58 |
| P33767 | WBP1 | LEYLDIN[+203.079373]STSTTVDLYDK | 2 | heavy | 1151.059 | b5 | 634.3083 | 63.58 |
| P33767 | WBP1 | LTLSPSGN[+203.079373]DSET-QYYTTGEFILPDR | 3 | light | 1003.14 | y11 | 1311.658 | 69.69 |
| P33767 | WBP1 | LTLSPSGN[+203.079373]DSET-QYYTTGEFILPDR | 3 | light | 1003.14 | y10 | 1148.595 | 69.69 |
| P33767 | WBP1 | LTLSPSGN[+203.079373]DSET-QYYTTGEFILPDR | 3 | light | 1003.14 | y9 | 1047.547 | 69.69 |
| P33767 | WBP1 | LTLSPSGN[+203.079373]DSET-QYYTTGEFILPDR | 3 | light | 1003.14 | y8 | 946.4993 | 69.69 |
| P33767 | WBP1 | LTLSPSGN[+203.079373]DSET-QYYTTGEFILPDR | 3 | light | 1003.14 | y6 | 760.4352 | 69.69 |
| P33767 | WBP1 | LTLSPSGN[+203.079373]DSET-QYYTTGEFILPDR | 3 | light | 1003.14 | y4 | 500.2827 | 69.69 |
| P33767 | WBP1 | LTLSPSGN[+203.079373]DSET-QYYTTGEFILPDR | 3 | light | 1003.14 | y3 | 387.1987 | 69.69 |
| P33767 | WBP1 | LTLSPSGN[+203.079373]DSET-QYYTTGEFILPDR | 3 | heavy | 1005.147 | y11 | 1317.678 | 69.69 |
| P33767 | WBP1 | LTLSPSGN[+203.079373]DSET-QYYTTGEFILPDR | 3 | heavy | 1005.147 | y10 | 1154.615 | 69.69 |
| P33767 | WBP1 | LTLSPSGN[+203.079373]DSET-QYYTTGEFILPDR | 3 | heavy | 1005.147 | y9 | 1053.567 | 69.69 |
| P33767 | WBP1 | LTLSPSGN[+203.079373]DSET-QYYTTGEFILPDR | 3 | heavy | 1005.147 | y8 | 952.5194 | 69.69 |
| P33767 | WBP1 | LTLSPSGN[+203.079373]DSET-QYYTTGEFILPDR | 3 | heavy | 1005.147 | y6 | 766.4553 | 69.69 |
| P33767 | WBP1 | LTLSPSGN[+203.079373]DSET-QYYTTGEFILPDR | 3 | heavy | 1005.147 | y4 | 506.3029 | 69.69 |
| P33767 | WBP1 | LTLSPSGN[+203.079373]DSET-QYYTTGEFILPDR | 3 | heavy | 1005.147 | y3 | 393.2188 | 69.69 |
| P33767 | WBP1 | LFLNELGIYPSPK | 2 | light | 745.9163 | y8 | 874.5033 | 74.48 |
| P33767 | WBP1 | LFLNELGIYPSPK | 2 | light | 745.9163 | y7 | 761.4192 | 74.48 |
| P33767 | WBP1 | LFLNELGIYPSPK | 2 | light | 745.9163 | y5 | 591.3137 | 74.48 |
| P33767 | WBP1 | LFLNELGIYPSPK | 2 | light | 745.9163 | y4 | 428.2504 | 74.48 |
| P33767 | WBP1 | LFLNELGIYPSPK | 2 | heavy | 749.9234 | y8 | 882.5175 | 74.48 |
| P33767 | WBP1 | LFLNELGIYPSPK | 2 | heavy | 749.9234 | y7 | 769.4334 | 74.48 |
| P33767 | WBP1 | LFLNELGIYPSPK | 2 | heavy | 749.9234 | y5 | 599.3279 | 74.48 |
| P33767 | WBP1 | LFLNELGIYPSPK | 2 | heavy | 749.9234 | y4 | 436.2646 | 74.48 |
| P33767 | WBP1 | EQIVPILNAPR | 2 | light | 625.3668 | y9 | 992.6251 | 58.71 |
| P33767 | WBP1 | EQIVPILNAPR | 2 | light | 625.3668 | y8 | 879.5411 | 58.71 |
| P33767 | WBP1 | EQIVPILNAPR | 2 | light | 625.3668 | y7 | 780.4726 | 58.71 |
| P33767 | WBP1 | EQIVPILNAPR | 2 | light | 625.3668 | y6 | 683.4199 | 58.71 |
| P33767 | WBP1 | EQIVPILNAPR | 2 | light | 625.3668 | y5 | 570.3358 | 58.71 |
| P33767 | WBP1 | EQIVPILNAPR | 2 | heavy | 628.3769 | y9 | 998.6453 | 58.71 |
| P33767 | WBP1 | EQIVPILNAPR | 2 | heavy | 628.3769 | y8 | 885.5612 | 58.71 |

| | | | | | | | | |
|---|---|---|---|---|---|---|---|---|
| P33767 | WBP1 | EQIVPILNAPR | 2 | heavy | 628.3769 | y7 | 786.4928 | 58.71 |
| P33767 | WBP1 | EQIVPILNAPR | 2 | heavy | 628.3769 | y6 | 689.44 | 58.71 |
| P33767 | WBP1 | EQIVPILNAPR | 2 | heavy | 628.3769 | y5 | 576.356 | 58.71 |
| P33767 | WBP1 | LFDNIIVFPTK | 2 | light | 653.8739 | y9 | 1046.588 | 77.19 |
| P33767 | WBP1 | LFDNIIVFPTK | 2 | light | 653.8739 | y8 | 931.5611 | 77.19 |
| P33767 | WBP1 | LFDNIIVFPTK | 2 | light | 653.8739 | y6 | 704.4341 | 77.19 |
| P33767 | WBP1 | LFDNIIVFPTK | 2 | light | 653.8739 | y5 | 591.3501 | 77.19 |
| P33767 | WBP1 | LFDNIIVFPTK | 2 | light | 653.8739 | y4 | 492.2817 | 77.19 |
| P33767 | WBP1 | LFDNIIVFPTK | 2 | heavy | 657.881 | y9 | 1054.602 | 77.19 |
| P33767 | WBP1 | LFDNIIVFPTK | 2 | heavy | 657.881 | y8 | 939.5753 | 77.19 |
| P33767 | WBP1 | LFDNIIVFPTK | 2 | heavy | 657.881 | y6 | 712.4483 | 77.19 |
| P33767 | WBP1 | LFDNIIVFPTK | 2 | heavy | 657.881 | y5 | 599.3643 | 77.19 |
| P33767 | WBP1 | LFDNIIVFPTK | 2 | heavy | 657.881 | y4 | 500.2959 | 77.19 |
| P33767 | WBP1 | LVWIGSSDFLK | 2 | light | 632.8504 | y9 | 1052.541 | 74.48 |
| P33767 | WBP1 | LVWIGSSDFLK | 2 | light | 632.8504 | y8 | 866.4618 | 74.48 |
| P33767 | WBP1 | LVWIGSSDFLK | 2 | light | 632.8504 | y7 | 753.3777 | 74.48 |
| P33767 | WBP1 | LVWIGSSDFLK | 2 | light | 632.8504 | y6 | 696.3563 | 74.48 |
| P33767 | WBP1 | LVWIGSSDFLK | 2 | light | 632.8504 | y5 | 609.3243 | 74.48 |
| P33767 | WBP1 | LVWIGSSDFLK | 2 | light | 632.8504 | y4 | 522.2922 | 74.48 |
| P33767 | WBP1 | LVWIGSSDFLK | 2 | light | 632.8504 | y3 | 407.2653 | 74.48 |
| P33767 | WBP1 | LVWIGSSDFLK | 2 | heavy | 636.8575 | y9 | 1060.555 | 74.48 |
| P33767 | WBP1 | LVWIGSSDFLK | 2 | heavy | 636.8575 | y8 | 874.476 | 74.48 |
| P33767 | WBP1 | LVWIGSSDFLK | 2 | heavy | 636.8575 | y7 | 761.3919 | 74.48 |
| P33767 | WBP1 | LVWIGSSDFLK | 2 | heavy | 636.8575 | y6 | 704.3705 | 74.48 |
| P33767 | WBP1 | LVWIGSSDFLK | 2 | heavy | 636.8575 | y5 | 617.3385 | 74.48 |
| P33767 | WBP1 | LVWIGSSDFLK | 2 | heavy | 636.8575 | y4 | 530.3064 | 74.48 |
| P33767 | WBP1 | LVWIGSSDFLK | 2 | heavy | 636.8575 | y3 | 415.2795 | 74.48 |
| P33767 | WBP1 | IGLSFTTDK | 2 | light | 491.2662 | y8 | 868.4411 | 48.83 |
| P33767 | WBP1 | IGLSFTTDK | 2 | light | 491.2662 | y7 | 811.4196 | 48.83 |
| P33767 | WBP1 | IGLSFTTDK | 2 | light | 491.2662 | y6 | 698.3355 | 48.83 |
| P33767 | WBP1 | IGLSFTTDK | 2 | light | 491.2662 | y5 | 611.3035 | 48.83 |
| P33767 | WBP1 | IGLSFTTDK | 2 | light | 491.2662 | y4 | 464.2351 | 48.83 |

| | | | | | | | | |
|---|---|---|---|---|---|---|---|---|
| P33767 | WBP1 | IGLSFTTDK | 2 | light | 491.2662 | y3 | 363.1874 | 48.83 |
| P33767 | WBP1 | IGLSFTTDK | 2 | heavy | 495.2733 | y8 | 876.4553 | 48.83 |
| P33767 | WBP1 | IGLSFTTDK | 2 | heavy | 495.2733 | y7 | 819.4338 | 48.83 |
| P33767 | WBP1 | IGLSFTTDK | 2 | heavy | 495.2733 | y6 | 706.3497 | 48.83 |
| P33767 | WBP1 | IGLSFTTDK | 2 | heavy | 495.2733 | y5 | 619.3177 | 48.83 |
| P33767 | WBP1 | IGLSFTTDK | 2 | heavy | 495.2733 | y4 | 472.2493 | 48.83 |
| P33767 | WBP1 | IGLSFTTDK | 2 | heavy | 495.2733 | y3 | 371.2016 | 48.83 |
| P41543 | OST1 | FSSN[+203.079373]ETLAIVYSH-NAPLNQVVNLR | 3 | light | 963.8296 | y12 | 1374.76 | 71.31 |
| P41543 | OST1 | FSSN[+203.079373]ETLAIVYSH-NAPLNQVVNLR | 3 | light | 963.8296 | y11 | 1237.701 | 71.31 |
| P41543 | OST1 | FSSN[+203.079373]ETLAIVYSH-NAPLNQVVNLR | 3 | light | 963.8296 | y10 | 1123.658 | 71.31 |
| P41543 | OST1 | FSSN[+203.079373]ETLAIVYSH-NAPLNQVVNLR | 3 | light | 963.8296 | y9 | 1052.621 | 71.31 |
| P41543 | OST1 | FSSN[+203.079373]ETLAIVYSH-NAPLNQVVNLR | 3 | light | 963.8296 | y7 | 842.4843 | 71.31 |
| P41543 | OST1 | FSSN[+203.079373]ETLAIVYSH-NAPLNQVVNLR | 3 | light | 963.8296 | y5 | 600.3828 | 71.31 |
| P41543 | OST1 | FSSN[+203.079373]ETLAIVYSH-NAPLNQVVNLR | 3 | light | 963.8296 | y3 | 402.2459 | 71.31 |
| P41543 | OST1 | FSSN[+203.079373]ETLAIVYSH-NAPLNQVVNLR | 3 | light | 963.8296 | b2 | 235.1077 | 71.31 |
| P41543 | OST1 | FSSN[+203.079373]ETLAIVYSH-NAPLNQVVNLR | 3 | light | 963.8296 | b8 | 850.3941 | 71.31 |
| P41543 | OST1 | FSSN[+203.079373]ETLAIVYSH-NAPLNQVVNLR | 3 | heavy | 965.8363 | y12 | 1380.78 | 71.31 |
| P41543 | OST1 | FSSN[+203.079373]ETLAIVYSH-NAPLNQVVNLR | 3 | heavy | 965.8363 | y11 | 1243.721 | 71.31 |
| P41543 | OST1 | FSSN[+203.079373]ETLAIVYSH-NAPLNQVVNLR | 3 | heavy | 965.8363 | y10 | 1129.678 | 71.31 |
| P41543 | OST1 | FSSN[+203.079373]ETLAIVYSH-NAPLNQVVNLR | 3 | heavy | 965.8363 | y9 | 1058.641 | 71.31 |
| P41543 | OST1 | FSSN[+203.079373]ETLAIVYSH-NAPLNQVVNLR | 3 | heavy | 965.8363 | y7 | 848.5044 | 71.31 |
| P41543 | OST1 | FSSN[+203.079373]ETLAIVYSH-NAPLNQVVNLR | 3 | heavy | 965.8363 | y5 | 606.4029 | 71.31 |
| P41543 | OST1 | FSSN[+203.079373]ETLAIVYSH-NAPLNQVVNLR | 3 | heavy | 965.8363 | y3 | 408.2661 | 71.31 |
| P41543 | OST1 | FSSN[+203.079373]ETLAIVYSH-NAPLNQVVNLR | 3 | heavy | 965.8363 | b2 | 235.1077 | 71.31 |
| P41543 | OST1 | FSSN[+203.079373]ETLAIVYSH-NAPLNQVVNLR | 3 | heavy | 965.8363 | b8 | 850.3941 | 71.31 |
| P41543 | OST1 | LSDFLHVSSGSDEK | 3 | light | 507.5791 | y9 | 945.4272 | 44.46 |
| P41543 | OST1 | LSDFLHVSSGSDEK | 3 | light | 507.5791 | y8 | 808.3683 | 44.46 |
| P41543 | OST1 | LSDFLHVSSGSDEK | 3 | light | 507.5791 | y7 | 709.2999 | 44.46 |
| P41543 | OST1 | LSDFLHVSSGSDEK | 3 | light | 507.5791 | y6 | 622.2679 | 44.46 |
| P41543 | OST1 | LSDFLHVSSGSDEK | 3 | light | 507.5791 | y2 | 276.1554 | 44.46 |
| P41543 | OST1 | LSDFLHVSSGSDEK | 3 | heavy | 510.2505 | y9 | 953.4414 | 44.46 |
| P41543 | OST1 | LSDFLHVSSGSDEK | 3 | heavy | 510.2505 | y8 | 816.3825 | 44.46 |

| | | | | | | | | |
|---|---|---|---|---|---|---|---|---|
| P41543 | OST1 | LSDFLHVSSGSDEK | 3 | heavy | 510.2505 | y7 | 717.3141 | 44.46 |
| P41543 | OST1 | LSDFLHVSSGSDEK | 3 | heavy | 510.2505 | y6 | 630.2821 | 44.46 |
| P41543 | OST1 | LSDFLHVSSGSDEK | 3 | heavy | 510.2505 | y2 | 284.1696 | 44.46 |
| P41543 | OST1 | NLISQVANGQVLIK | 2 | light | 748.9434 | y11 | 1156.668 | 65.78 |
| P41543 | OST1 | NLISQVANGQVLIK | 2 | light | 748.9434 | y10 | 1069.636 | 65.78 |
| P41543 | OST1 | NLISQVANGQVLIK | 2 | light | 748.9434 | y9 | 941.5778 | 65.78 |
| P41543 | OST1 | NLISQVANGQVLIK | 2 | light | 748.9434 | y8 | 842.5094 | 65.78 |
| P41543 | OST1 | NLISQVANGQVLIK | 2 | light | 748.9434 | y7 | 771.4723 | 65.78 |
| P41543 | OST1 | NLISQVANGQVLIK | 2 | heavy | 752.9505 | y11 | 1164.683 | 65.78 |
| P41543 | OST1 | NLISQVANGQVLIK | 2 | heavy | 752.9505 | y10 | 1077.651 | 65.78 |
| P41543 | OST1 | NLISQVANGQVLIK | 2 | heavy | 752.9505 | y9 | 949.592 | 65.78 |
| P41543 | OST1 | NLISQVANGQVLIK | 2 | heavy | 752.9505 | y8 | 850.5236 | 65.78 |
| P41543 | OST1 | NLISQVANGQVLIK | 2 | heavy | 752.9505 | y7 | 779.4865 | 65.78 |
| P41543 | OST1 | LTFSYR | 2 | light | 393.7109 | y5 | 673.3304 | 40.67 |
| P41543 | OST1 | LTFSYR | 2 | light | 393.7109 | y4 | 572.2827 | 40.67 |
| P41543 | OST1 | LTFSYR | 2 | light | 393.7109 | y3 | 425.2143 | 40.67 |
| P41543 | OST1 | LTFSYR | 2 | heavy | 396.7209 | y5 | 679.3505 | 40.67 |
| P41543 | OST1 | LTFSYR | 2 | heavy | 396.7209 | y4 | 578.3029 | 40.67 |
| P41543 | OST1 | LTFSYR | 2 | heavy | 396.7209 | y3 | 431.2344 | 40.67 |
| P41543 | OST1 | ANGNSFEFGPWEDIPR | 2 | light | 918.421 | y9 | 1116.547 | 74.43 |
| P41543 | OST1 | ANGNSFEFGPWEDIPR | 2 | light | 918.421 | y8 | 969.4789 | 74.43 |
| P41543 | OST1 | ANGNSFEFGPWEDIPR | 2 | light | 918.421 | y7 | 912.4574 | 74.43 |
| P41543 | OST1 | ANGNSFEFGPWEDIPR | 2 | light | 918.421 | y3 | 385.2558 | 74.43 |
| P41543 | OST1 | ANGNSFEFGPWEDIPR | 2 | light | 918.421 | y2 | 272.1717 | 74.43 |
| P41543 | OST1 | ANGNSFEFGPWEDIPR | 2 | heavy | 921.4311 | y9 | 1122.567 | 74.43 |
| P41543 | OST1 | ANGNSFEFGPWEDIPR | 2 | heavy | 921.4311 | y8 | 975.499 | 74.43 |
| P41543 | OST1 | ANGNSFEFGPWEDIPR | 2 | heavy | 921.4311 | y7 | 918.4775 | 74.43 |
| P41543 | OST1 | ANGNSFEFGPWEDIPR | 2 | heavy | 921.4311 | y3 | 391.2759 | 74.43 |
| P41543 | OST1 | ANGNSFEFGPWEDIPR | 2 | heavy | 921.4311 | y2 | 278.1918 | 74.43 |
| OST5 | YEAST | TYEQLYK | 2 | light | 472.7398 | y6 | 843.4247 | 30.95 |
| OST5 | YEAST | TYEQLYK | 2 | light | 472.7398 | y5 | 680.3614 | 30.95 |
| OST5 | YEAST | TYEQLYK | 2 | light | 472.7398 | y4 | 551.3188 | 30.95 |

| | | | | | | | | |
|---|---|---|---|---|---|---|---|---|
| OST5 | YEAST | TYEQLYK | 2 | light | 472.7398 | y2 | 310.1761 | 30.95 |
| OST5 | YEAST | TYEQLYK | 2 | heavy | 476.7469 | y6 | 851.4389 | 30.95 |
| OST5 | YEAST | TYEQLYK | 2 | heavy | 476.7469 | y5 | 688.3756 | 30.95 |
| OST5 | YEAST | TYEQLYK | 2 | heavy | 476.7469 | y4 | 559.333 | 30.95 |
| OST5 | YEAST | TYEQLYK | 2 | heavy | 476.7469 | y2 | 318.1903 | 30.95 |
| OST6 | YEAST | FVEMNAIPFIAR | 2 | light | 704.3763 | y10 | 1161.608 | 75.68 |
| OST6 | YEAST | FVEMNAIPFIAR | 2 | light | 704.3763 | y9 | 1032.566 | 75.68 |
| OST6 | YEAST | FVEMNAIPFIAR | 2 | light | 704.3763 | y8 | 901.5254 | 75.68 |
| OST6 | YEAST | FVEMNAIPFIAR | 2 | light | 704.3763 | y7 | 787.4825 | 75.68 |
| OST6 | YEAST | FVEMNAIPFIAR | 2 | light | 704.3763 | y5 | 603.3613 | 75.68 |
| OST6 | YEAST | FVEMNAIPFIAR | 2 | heavy | 707.3864 | y10 | 1167.629 | 75.68 |
| OST6 | YEAST | FVEMNAIPFIAR | 2 | heavy | 707.3864 | y9 | 1038.586 | 75.68 |
| OST6 | YEAST | FVEMNAIPFIAR | 2 | heavy | 707.3864 | y8 | 907.5455 | 75.68 |
| OST6 | YEAST | FVEMNAIPFIAR | 2 | heavy | 707.3864 | y7 | 793.5026 | 75.68 |
| OST6 | YEAST | FVEMNAIPFIAR | 2 | heavy | 707.3864 | y5 | 609.3814 | 75.68 |
| OST6 | YEAST | LQNVPHLVVYPPAESNK | 2 | light | 953.0151 | y8 | 905.4363 | 52.42 |
| OST6 | YEAST | LQNVPHLVVYPPAESNK | 2 | light | 953.0151 | y7 | 742.373 | 52.42 |
| OST6 | YEAST | LQNVPHLVVYPPAESNK | 2 | light | 953.0151 | y6 | 645.3202 | 52.42 |
| OST6 | YEAST | LQNVPHLVVYPPAESNK | 2 | light | 953.0151 | y3 | 348.1878 | 52.42 |
| OST6 | YEAST | LQNVPHLVVYPPAESNK | 2 | light | 953.0151 | b2 | 242.1499 | 52.42 |
| OST6 | YEAST | LQNVPHLVVYPPAESNK | 2 | light | 953.0151 | b3 | 356.1928 | 52.42 |
| OST6 | YEAST | LQNVPHLVVYPPAESNK | 2 | light | 953.0151 | b4 | 455.2613 | 52.42 |
| OST6 | YEAST | LQNVPHLVVYPPAESNK | 2 | light | 953.0151 | b10 | 1163.657 | 52.42 |
| OST6 | YEAST | LQNVPHLVVYPPAESNK | 2 | heavy | 957.0222 | y8 | 913.4505 | 52.42 |
| OST6 | YEAST | LQNVPHLVVYPPAESNK | 2 | heavy | 957.0222 | y7 | 750.3872 | 52.42 |
| OST6 | YEAST | LQNVPHLVVYPPAESNK | 2 | heavy | 957.0222 | y6 | 653.3344 | 52.42 |
| OST6 | YEAST | LQNVPHLVVYPPAESNK | 2 | heavy | 957.0222 | y3 | 356.202 | 52.42 |
| OST6 | YEAST | LQNVPHLVVYPPAESNK | 2 | heavy | 957.0222 | b2 | 242.1499 | 52.42 |
| OST6 | YEAST | LQNVPHLVVYPPAESNK | 2 | heavy | 957.0222 | b3 | 356.1928 | 52.42 |
| OST6 | YEAST | LQNVPHLVVYPPAESNK | 2 | heavy | 957.0222 | b4 | 455.2613 | 52.42 |
| OST6 | YEAST | LQNVPHLVVYPPAESNK | 2 | heavy | 957.0222 | b10 | 1163.657 | 52.42 |
| OST6 | YEAST | TYHAVADVIR | 2 | light | 572.8091 | y8 | 880.4999 | 34.74 |

| | | | | | | | | |
|---|---|---|---|---|---|---|---|---|
| OST6 | YEAST | TYHAVADVIR | 2 | light | 572.8091 | y3 | 387.2714 | 34.74 |
| OST6 | YEAST | TYHAVADVIR | 2 | light | 572.8091 | y8 | 440.7536 | 34.74 |
| OST6 | YEAST | TYHAVADVIR | 2 | heavy | 575.8192 | y8 | 886.5201 | 34.74 |
| OST6 | YEAST | TYHAVADVIR | 2 | heavy | 575.8192 | y3 | 393.2916 | 34.74 |
| OST6 | YEAST | TYHAVADVIR | 2 | heavy | 575.8192 | y8 | 443.7637 | 34.74 |
| OST2 | YEAST | VTSTSSAVLTDFQETFK | 2 | light | 930.9649 | y10 | 1227.626 | 69.6 |
| OST2 | YEAST | VTSTSSAVLTDFQETFK | 2 | light | 930.9649 | y9 | 1128.557 | 69.6 |
| OST2 | YEAST | VTSTSSAVLTDFQETFK | 2 | light | 930.9649 | y8 | 1015.473 | 69.6 |
| OST2 | YEAST | VTSTSSAVLTDFQETFK | 2 | light | 930.9649 | y7 | 914.4254 | 69.6 |
| OST2 | YEAST | VTSTSSAVLTDFQETFK | 2 | light | 930.9649 | y6 | 799.3985 | 69.6 |
| OST2 | YEAST | VTSTSSAVLTDFQETFK | 2 | heavy | 934.972 | y10 | 1235.64 | 69.6 |
| OST2 | YEAST | VTSTSSAVLTDFQETFK | 2 | heavy | 934.972 | y9 | 1136.571 | 69.6 |
| OST2 | YEAST | VTSTSSAVLTDFQETFK | 2 | heavy | 934.972 | y8 | 1023.487 | 69.6 |
| OST2 | YEAST | VTSTSSAVLTDFQETFK | 2 | heavy | 934.972 | y7 | 922.4396 | 69.6 |
| OST2 | YEAST | VTSTSSAVLTDFQETFK | 2 | heavy | 934.972 | y6 | 807.4127 | 69.6 |
| OST2 | YEAST | RAYFAQIEK | 2 | light | 563.3062 | b5 | 609.3144 | 34.06 |
| OST2 | YEAST | RAYFAQIEK | 2 | light | 563.3062 | b6 | 737.3729 | 34.06 |
| OST2 | YEAST | RAYFAQIEK | 2 | light | 563.3062 | b7 | 850.457 | 34.06 |
| OST2 | YEAST | RAYFAQIEK | 2 | light | 563.3062 | b8 | 979.4996 | 34.06 |
| OST2 | YEAST | RAYFAQIEK | 2 | heavy | 570.3234 | b5 | 615.3345 | 34.06 |
| OST2 | YEAST | RAYFAQIEK | 2 | heavy | 570.3234 | b6 | 743.3931 | 34.06 |
| OST2 | YEAST | RAYFAQIEK | 2 | heavy | 570.3234 | b7 | 856.4771 | 34.06 |
| OST2 | YEAST | RAYFAQIEK | 2 | heavy | 570.3234 | b8 | 985.5197 | 34.06 |
| OSTD | YEAST.SWP1 | NLEMAFEPEIK | 2 | light | 660.8288 | y9 | 1093.523 | 61.67 |
| OSTD | YEAST.SWP1 | NLEMAFEPEIK | 2 | light | 660.8288 | y8 | 964.4808 | 61.67 |
| OSTD | YEAST.SWP1 | NLEMAFEPEIK | 2 | light | 660.8288 | y7 | 833.4403 | 61.67 |
| OSTD | YEAST.SWP1 | NLEMAFEPEIK | 2 | light | 660.8288 | y5 | 615.3348 | 61.67 |
| OSTD | YEAST.SWP1 | NLEMAFEPEIK | 2 | light | 660.8288 | y4 | 486.2922 | 61.67 |
| OSTD | YEAST.SWP1 | NLEMAFEPEIK | 2 | heavy | 664.8359 | y9 | 1101.538 | 61.67 |
| OSTD | YEAST.SWP1 | NLEMAFEPEIK | 2 | heavy | 664.8359 | y8 | 972.495 | 61.67 |
| OSTD | YEAST.SWP1 | NLEMAFEPEIK | 2 | heavy | 664.8359 | y7 | 841.4545 | 61.67 |
| OSTD | YEAST.SWP1 | NLEMAFEPEIK | 2 | heavy | 664.8359 | y5 | 623.349 | 61.67 |

| | | | | | | | | |
|---|---|---|---|---|---|---|---|---|
| OSTD | YEAST.SWP1 | NLEMAFEPEIK | 2 | heavy | 664.8359 | y4 | 494.3064 | 61.67 |
| OSTD | YEAST.SWP1 | YRIDLAK | 2 | light | 439.7584 | y3 | 331.234 | 32.67 |
| OSTD | YEAST.SWP1 | YRIDLAK | 2 | light | 439.7584 | y2 | 218.1499 | 32.67 |
| OSTD | YEAST.SWP1 | YRIDLAK | 2 | light | 439.7584 | b4 | 548.2827 | 32.67 |
| OSTD | YEAST.SWP1 | YRIDLAK | 2 | light | 439.7584 | b5 | 661.3668 | 32.67 |
| OSTD | YEAST.SWP1 | YRIDLAK | 2 | light | 439.7584 | b6 | 732.4039 | 32.67 |
| OSTD | YEAST.SWP1 | YRIDLAK | 2 | heavy | 446.7755 | y3 | 339.2482 | 32.67 |
| OSTD | YEAST.SWP1 | YRIDLAK | 2 | heavy | 446.7755 | y2 | 226.1641 | 32.67 |
| OSTD | YEAST.SWP1 | YRIDLAK | 2 | heavy | 446.7755 | b4 | 554.3029 | 32.67 |
| OSTD | YEAST.SWP1 | YRIDLAK | 2 | heavy | 446.7755 | b5 | 667.3869 | 32.67 |
| OSTD | YEAST.SWP1 | YRIDLAK | 2 | heavy | 446.7755 | b6 | 738.424 | 32.67 |
| OST3 | YEAST | SPAYPFPLLR | 2 | light | 580.8268 | y7 | 905.5244 | 70.53 |
| OST3 | YEAST | SPAYPFPLLR | 2 | light | 580.8268 | y6 | 742.461 | 70.53 |
| OST3 | YEAST | SPAYPFPLLR | 2 | light | 580.8268 | y5 | 645.4083 | 70.53 |
| OST3 | YEAST | SPAYPFPLLR | 2 | light | 580.8268 | b4 | 419.1925 | 70.53 |
| OST3 | YEAST | SPAYPFPLLR | 2 | heavy | 583.8368 | y7 | 911.5445 | 70.53 |
| OST3 | YEAST | SPAYPFPLLR | 2 | heavy | 583.8368 | y6 | 748.4812 | 70.53 |
| OST3 | YEAST | SPAYPFPLLR | 2 | heavy | 583.8368 | y5 | 651.4284 | 70.53 |
| OST3 | YEAST | SPAYPFPLLR | 2 | heavy | 583.8368 | b4 | 419.1925 | 70.53 |
| OST3 | YEAST | SSNSDTSIFFTK | 2 | light | 667.3172 | y10 | 1159.563 | 50.77 |
| OST3 | YEAST | SSNSDTSIFFTK | 2 | light | 667.3172 | y7 | 843.4611 | 50.77 |
| OST3 | YEAST | SSNSDTSIFFTK | 2 | light | 667.3172 | y6 | 742.4134 | 50.77 |
| OST3 | YEAST | SSNSDTSIFFTK | 2 | light | 667.3172 | y4 | 542.2973 | 50.77 |
| OST3 | YEAST | SSNSDTSIFFTK | 2 | heavy | 671.3243 | y10 | 1167.577 | 50.77 |
| OST3 | YEAST | SSNSDTSIFFTK | 2 | heavy | 671.3243 | y7 | 851.4753 | 50.77 |
| OST3 | YEAST | SSNSDTSIFFTK | 2 | heavy | 671.3243 | y6 | 750.4276 | 50.77 |
| OST3 | YEAST | SSNSDTSIFFTK | 2 | heavy | 671.3243 | y4 | 550.3115 | 50.77 |
| OST3 | YEAST | QIIQAIK | 2 | light | 407.2633 | y6 | 685.4607 | 35.89 |
| OST3 | YEAST | QIIQAIK | 2 | light | 407.2633 | y5 | 572.3766 | 35.89 |
| OST3 | YEAST | QIIQAIK | 2 | light | 407.2633 | y4 | 459.2926 | 35.89 |
| OST3 | YEAST | QIIQAIK | 2 | light | 407.2633 | y3 | 331.234 | 35.89 |
| OST3 | YEAST | QIIQAIK | 2 | light | 407.2633 | y5 | 286.692 | 35.89 |

| | | | | | | | | |
|---|---|---|---|---|---|---|---|---|
| OST3 | YEAST | QIIQAIK | 2 | heavy | 411.2704 | y6 | 693.4749 | 35.89 |
| OST3 | YEAST | QIIQAIK | 2 | heavy | 411.2704 | y5 | 580.3908 | 35.89 |
| OST3 | YEAST | QIIQAIK | 2 | heavy | 411.2704 | y4 | 467.3068 | 35.89 |
| OST3 | YEAST | QIIQAIK | 2 | heavy | 411.2704 | y3 | 339.2482 | 35.89 |
| OST3 | YEAST | QIIQAIK | 2 | heavy | 411.2704 | y5 | 290.699 | 35.89 |
| P39007 | STT3 | TTLVDNNTWN[+203.079373]NTHIAIVGK | 3 | light | 772.0606 | y12 | 1353.727 | 54.42 |
| P39007 | STT3 | TTLVDNNTWN[+203.079373]NTHIAIVGK | 3 | light | 772.0606 | y11 | 1252.68 | 54.42 |
| P39007 | STT3 | TTLVDNNTWN[+203.079373]NTHIAIVGK | 3 | light | 772.0606 | y10 | 1269.68 | 54.42 |
| P39007 | STT3 | TTLVDNNTWN[+203.079373]NTHIAIVGK | 3 | light | 772.0606 | y10 | 1066.6 | 54.42 |
| P39007 | STT3 | TTLVDNNTWN[+203.079373]NTHIAIVGK | 3 | light | 772.0606 | y9 | 952.5574 | 54.42 |
| P39007 | STT3 | TTLVDNNTWN[+203.079373]NTHIAIVGK | 3 | light | 772.0606 | y8 | 838.5145 | 54.42 |
| P39007 | STT3 | TTLVDNNTWN[+203.079373]NTHIAIVGK | 3 | light | 772.0606 | y7 | 737.4668 | 54.42 |
| P39007 | STT3 | TTLVDNNTWN[+203.079373]NTHIAIVGK | 3 | light | 772.0606 | y6 | 600.4079 | 54.42 |
| P39007 | STT3 | TTLVDNNTWN[+203.079373]NTHIAIVGK | 3 | light | 772.0606 | y5 | 487.3239 | 54.42 |
| P39007 | STT3 | TTLVDNNTWN[+203.079373]NTHIAIVGK | 3 | light | 772.0606 | y4 | 416.2867 | 54.42 |
| P39007 | STT3 | TTLVDNNTWN[+203.079373]NTHIAIVGK | 3 | light | 772.0606 | b4 | 415.2551 | 54.42 |
| P39007 | STT3 | TTLVDNNTWN[+203.079373]NTHIAIVGK | 3 | heavy | 774.732 | y12 | 1361.742 | 54.42 |
| P39007 | STT3 | TTLVDNNTWN[+203.079373]NTHIAIVGK | 3 | heavy | 774.732 | y11 | 1260.694 | 54.42 |
| P39007 | STT3 | TTLVDNNTWN[+203.079373]NTHIAIVGK | 3 | heavy | 774.732 | y10 | 1277.694 | 54.42 |
| P39007 | STT3 | TTLVDNNTWN[+203.079373]NTHIAIVGK | 3 | heavy | 774.732 | y10 | 1074.615 | 54.42 |
| P39007 | STT3 | TTLVDNNTWN[+203.079373]NTHIAIVGK | 3 | heavy | 774.732 | y9 | 960.5716 | 54.42 |
| P39007 | STT3 | TTLVDNNTWN[+203.079373]NTHIAIVGK | 3 | heavy | 774.732 | y8 | 846.5287 | 54.42 |
| P39007 | STT3 | TTLVDNNTWN[+203.079373]NTHIAIVGK | 3 | heavy | 774.732 | y7 | 745.481 | 54.42 |
| P39007 | STT3 | TTLVDNNTWN[+203.079373]NTHIAIVGK | 3 | heavy | 774.732 | y6 | 608.4221 | 54.42 |
| P39007 | STT3 | TTLVDNNTWN[+203.079373]NTHIAIVGK | 3 | heavy | 774.732 | y5 | 495.3381 | 54.42 |
| P39007 | STT3 | TTLVDNNTWN[+203.079373]NTHIAIVGK | 3 | heavy | 774.732 | y4 | 424.3009 | 54.42 |
| P39007 | STT3 | TTLVDNNTWN[+203.079373]NTHIAIVGK | 3 | heavy | 774.732 | b4 | 415.2551 | 54.42 |
| P39007 | STT3 | TAYSSPSVVLPSQTPDGK | 2 | light | 917.4651 | y10 | 1041.558 | 47.16 |
| P39007 | STT3 | TAYSSPSVVLPSQTPDGK | 2 | light | 917.4651 | y9 | 942.4891 | 47.16 |
| P39007 | STT3 | TAYSSPSVVLPSQTPDGK | 2 | light | 917.4651 | y8 | 829.405 | 47.16 |
| P39007 | STT3 | TAYSSPSVVLPSQTPDGK | 2 | light | 917.4651 | y7 | 732.3523 | 47.16 |
| P39007 | STT3 | TAYSSPSVVLPSQTPDGK | 2 | light | 917.4651 | y6 | 645.3202 | 47.16 |

| | | | | | | | | |
|---|---|---|---|---|---|---|---|---|
| P39007 | STT3 | TAYSSPSVVLPSQTPDGK | 2 | light | 917.4651 | y5 | 517.2617 | 47.16 |
| P39007 | STT3 | TAYSSPSVVLPSQTPDGK | 2 | light | 917.4651 | y4 | 416.214 | 47.16 |
| P39007 | STT3 | TAYSSPSVVLPSQTPDGK | 2 | heavy | 921.4722 | y10 | 1049.572 | 47.16 |
| P39007 | STT3 | TAYSSPSVVLPSQTPDGK | 2 | heavy | 921.4722 | y9 | 950.5033 | 47.16 |
| P39007 | STT3 | TAYSSPSVVLPSQTPDGK | 2 | heavy | 921.4722 | y8 | 837.4192 | 47.16 |
| P39007 | STT3 | TAYSSPSVVLPSQTPDGK | 2 | heavy | 921.4722 | y7 | 740.3665 | 47.16 |
| P39007 | STT3 | TAYSSPSVVLPSQTPDGK | 2 | heavy | 921.4722 | y6 | 653.3344 | 47.16 |
| P39007 | STT3 | TAYSSPSVVLPSQTPDGK | 2 | heavy | 921.4722 | y5 | 525.2759 | 47.16 |
| P39007 | STT3 | TAYSSPSVVLPSQTPDGK | 2 | heavy | 921.4722 | y4 | 424.2282 | 47.16 |
| P39007 | STT3 | ISEGIWPEEIK | 2 | light | 650.8428 | y10 | 1187.594 | 58.54 |
| P39007 | STT3 | ISEGIWPEEIK | 2 | light | 650.8428 | y9 | 1100.562 | 58.54 |
| P39007 | STT3 | ISEGIWPEEIK | 2 | light | 650.8428 | y8 | 971.5197 | 58.54 |
| P39007 | STT3 | ISEGIWPEEIK | 2 | light | 650.8428 | y6 | 801.4141 | 58.54 |
| P39007 | STT3 | ISEGIWPEEIK | 2 | light | 650.8428 | y5 | 615.3348 | 58.54 |
| P39007 | STT3 | ISEGIWPEEIK | 2 | heavy | 654.8499 | y10 | 1195.608 | 58.54 |
| P39007 | STT3 | ISEGIWPEEIK | 2 | heavy | 654.8499 | y9 | 1108.576 | 58.54 |
| P39007 | STT3 | ISEGIWPEEIK | 2 | heavy | 654.8499 | y8 | 979.5339 | 58.54 |
| P39007 | STT3 | ISEGIWPEEIK | 2 | heavy | 654.8499 | y6 | 809.4283 | 58.54 |
| P39007 | STT3 | ISEGIWPEEIK | 2 | heavy | 654.8499 | y5 | 623.349 | 58.54 |
| Q07830 | GPI13 | N[+203.079373]ISNTPPTSDPEK | 2 | light | 801.8783 | y12 | 1285.627 | 24.26 |
| Q07830 | GPI13 | N[+203.079373]ISNTPPTSDPEK | 2 | light | 801.8783 | y11 | 1172.543 | 24.26 |
| Q07830 | GPI13 | N[+203.079373]ISNTPPTSDPEK | 2 | light | 801.8783 | y10 | 1085.511 | 24.26 |
| Q07830 | GPI13 | N[+203.079373]ISNTPPTSDPEK | 2 | light | 801.8783 | y9 | 971.468 | 24.26 |
| Q07830 | GPI13 | N[+203.079373]ISNTPPTSDPEK | 2 | light | 801.8783 | y8 | 870.4203 | 24.26 |
| Q07830 | GPI13 | N[+203.079373]ISNTPPTSDPEK | 2 | light | 801.8783 | y7 | 773.3676 | 24.26 |
| Q07830 | GPI13 | N[+203.079373]ISNTPPTSDPEK | 2 | light | 801.8783 | y8 | 435.7138 | 24.26 |
| Q07830 | GPI13 | N[+203.079373]ISNTPPTSDPEK | 2 | heavy | 805.8854 | y12 | 1293.641 | 24.26 |
| Q07830 | GPI13 | N[+203.079373]ISNTPPTSDPEK | 2 | heavy | 805.8854 | y11 | 1180.557 | 24.26 |
| Q07830 | GPI13 | N[+203.079373]ISNTPPTSDPEK | 2 | heavy | 805.8854 | y10 | 1093.525 | 24.26 |
| Q07830 | GPI13 | N[+203.079373]ISNTPPTSDPEK | 2 | heavy | 805.8854 | y9 | 979.4822 | 24.26 |
| Q07830 | GPI13 | N[+203.079373]ISNTPPTSDPEK | 2 | heavy | 805.8854 | y8 | 878.4345 | 24.26 |
| Q07830 | GPI13 | N[+203.079373]ISNTPPTSDPEK | 2 | heavy | 805.8854 | y7 | 781.3818 | 24.26 |

| | | | | | | | | |
|---|---|---|---|---|---|---|---|---|
| Q07830 | GPI13 | N[+203.079373]ISNTPPTSDPEK | 2 | heavy | 805.8854 | y8 | 439.7209 | 24.26 |
| Q07830 | GPI13 | ETSNYNIDNLGHDYR | 2 | light | 905.9032 | y10 | 1216.571 | 41.14 |
| Q07830 | GPI13 | ETSNYNIDNLGHDYR | 2 | light | 905.9032 | y9 | 1102.528 | 41.14 |
| Q07830 | GPI13 | ETSNYNIDNLGHDYR | 2 | light | 905.9032 | y8 | 989.4435 | 41.14 |
| Q07830 | GPI13 | ETSNYNIDNLGHDYR | 2 | light | 905.9032 | y6 | 760.3737 | 41.14 |
| Q07830 | GPI13 | ETSNYNIDNLGHDYR | 2 | light | 905.9032 | y5 | 647.2896 | 41.14 |
| Q07830 | GPI13 | ETSNYNIDNLGHDYR | 2 | light | 905.9032 | y2 | 338.1823 | 41.14 |
| Q07830 | GPI13 | ETSNYNIDNLGHDYR | 2 | heavy | 908.9132 | y10 | 1222.591 | 41.14 |
| Q07830 | GPI13 | ETSNYNIDNLGHDYR | 2 | heavy | 908.9132 | y9 | 1108.548 | 41.14 |
| Q07830 | GPI13 | ETSNYNIDNLGHDYR | 2 | heavy | 908.9132 | y8 | 995.4637 | 41.14 |
| Q07830 | GPI13 | ETSNYNIDNLGHDYR | 2 | heavy | 908.9132 | y6 | 766.3938 | 41.14 |
| Q07830 | GPI13 | ETSNYNIDNLGHDYR | 2 | heavy | 908.9132 | y5 | 653.3097 | 41.14 |
| Q07830 | GPI13 | ETSNYNIDNLGHDYR | 2 | heavy | 908.9132 | y2 | 344.2024 | 41.14 |
| P36016 | LHS1 | LSN[+203.079373]ESELYDVFTR | 2 | light | 888.4203 | y10 | 1258.595 | 62.29 |
| P36016 | LHS1 | LSN[+203.079373]ESELYDVFTR | 2 | light | 888.4203 | y9 | 1129.552 | 62.29 |
| P36016 | LHS1 | LSN[+203.079373]ESELYDVFTR | 2 | light | 888.4203 | y8 | 1042.52 | 62.29 |
| P36016 | LHS1 | LSN[+203.079373]ESELYDVFTR | 2 | light | 888.4203 | y7 | 913.4778 | 62.29 |
| P36016 | LHS1 | LSN[+203.079373]ESELYDVFTR | 2 | light | 888.4203 | y6 | 800.3937 | 62.29 |
| P36016 | LHS1 | LSN[+203.079373]ESELYDVFTR | 2 | light | 888.4203 | y5 | 637.3304 | 62.29 |
| P36016 | LHS1 | LSN[+203.079373]ESELYDVFTR | 2 | light | 888.4203 | y4 | 522.3035 | 62.29 |
| P36016 | LHS1 | LSN[+203.079373]ESELYDVFTR | 2 | light | 888.4203 | y3 | 423.235 | 62.29 |
| P36016 | LHS1 | LSN[+203.079373]ESELYDVFTR | 2 | heavy | 891.4304 | y10 | 1264.615 | 62.29 |
| P36016 | LHS1 | LSN[+203.079373]ESELYDVFTR | 2 | heavy | 891.4304 | y9 | 1135.573 | 62.29 |
| P36016 | LHS1 | LSN[+203.079373]ESELYDVFTR | 2 | heavy | 891.4304 | y8 | 1048.541 | 62.29 |
| P36016 | LHS1 | LSN[+203.079373]ESELYDVFTR | 2 | heavy | 891.4304 | y7 | 919.4979 | 62.29 |
| P36016 | LHS1 | LSN[+203.079373]ESELYDVFTR | 2 | heavy | 891.4304 | y6 | 806.4139 | 62.29 |
| P36016 | LHS1 | LSN[+203.079373]ESELYDVFTR | 2 | heavy | 891.4304 | y5 | 643.3505 | 62.29 |
| P36016 | LHS1 | LSN[+203.079373]ESELYDVFTR | 2 | heavy | 891.4304 | y4 | 528.3236 | 62.29 |
| P36016 | LHS1 | LSN[+203.079373]ESELYDVFTR | 2 | heavy | 891.4304 | y3 | 429.2552 | 62.29 |
| P36016 | LHS1 | AIVVSPQAPLELVLTPEAK | 3 | light | 659.054 | y11 | 1209.709 | 82.57 |
| P36016 | LHS1 | AIVVSPQAPLELVLTPEAK | 3 | light | 659.054 | y8 | 870.5295 | 82.57 |
| P36016 | LHS1 | AIVVSPQAPLELVLTPEAK | 3 | light | 659.054 | y7 | 757.4454 | 82.57 |

| | | | | | | | | |
|---|---|---|---|---|---|---|---|---|
| P36016 | LHS1 | AIVVSPQAPLELVLTPEAK | 3 | light | 659.054 | y6 | 658.377 | 82.57 |
| P36016 | LHS1 | AIVVSPQAPLELVLTPEAK | 3 | light | 659.054 | y5 | 545.293 | 82.57 |
| P36016 | LHS1 | AIVVSPQAPLELVLTPEAK | 3 | heavy | 661.7254 | y11 | 1217.723 | 82.57 |
| P36016 | LHS1 | AIVVSPQAPLELVLTPEAK | 3 | heavy | 661.7254 | y8 | 878.5437 | 82.57 |
| P36016 | LHS1 | AIVVSPQAPLELVLTPEAK | 3 | heavy | 661.7254 | y7 | 765.4596 | 82.57 |
| P36016 | LHS1 | AIVVSPQAPLELVLTPEAK | 3 | heavy | 661.7254 | y6 | 666.3912 | 82.57 |
| P36016 | LHS1 | AIVVSPQAPLELVLTPEAK | 3 | heavy | 661.7254 | y5 | 553.3072 | 82.57 |
| P36016 | LHS1 | ALSTWEETLTSFK | 2 | light | 756.8827 | y9 | 1140.557 | 72.23 |
| P36016 | LHS1 | ALSTWEETLTSFK | 2 | light | 756.8827 | y8 | 954.4779 | 72.23 |
| P36016 | LHS1 | ALSTWEETLTSFK | 2 | light | 756.8827 | y7 | 825.4353 | 72.23 |
| P36016 | LHS1 | ALSTWEETLTSFK | 2 | light | 756.8827 | y6 | 696.3927 | 72.23 |
| P36016 | LHS1 | ALSTWEETLTSFK | 2 | light | 756.8827 | y5 | 595.345 | 72.23 |
| P36016 | LHS1 | ALSTWEETLTSFK | 2 | light | 756.8827 | y4 | 482.2609 | 72.23 |
| P36016 | LHS1 | ALSTWEETLTSFK | 2 | heavy | 760.8898 | y9 | 1148.571 | 72.23 |
| P36016 | LHS1 | ALSTWEETLTSFK | 2 | heavy | 760.8898 | y8 | 962.4921 | 72.23 |
| P36016 | LHS1 | ALSTWEETLTSFK | 2 | heavy | 760.8898 | y7 | 833.4495 | 72.23 |
| P36016 | LHS1 | ALSTWEETLTSFK | 2 | heavy | 760.8898 | y6 | 704.4069 | 72.23 |
| P36016 | LHS1 | ALSTWEETLTSFK | 2 | heavy | 760.8898 | y5 | 603.3592 | 72.23 |
| P36016 | LHS1 | ALSTWEETLTSFK | 2 | heavy | 760.8898 | y4 | 490.2751 | 72.23 |
| P39105 | PLB1 | DAGFN[+203.079373]IS-LADVWGR | 2 | light | 862.4179 | y9 | 1016.552 | 78.75 |
| P39105 | PLB1 | DAGFN[+203.079373]IS-LADVWGR | 2 | light | 862.4179 | y8 | 903.4683 | 78.75 |
| P39105 | PLB1 | DAGFN[+203.079373]IS-LADVWGR | 2 | light | 862.4179 | y7 | 816.4363 | 78.75 |
| P39105 | PLB1 | DAGFN[+203.079373]IS-LADVWGR | 2 | light | 862.4179 | y6 | 703.3522 | 78.75 |
| P39105 | PLB1 | DAGFN[+203.079373]IS-LADVWGR | 2 | light | 862.4179 | y4 | 517.2881 | 78.75 |
| P39105 | PLB1 | DAGFN[+203.079373]IS-LADVWGR | 2 | heavy | 865.428 | y9 | 1022.572 | 78.75 |
| P39105 | PLB1 | DAGFN[+203.079373]IS-LADVWGR | 2 | heavy | 865.428 | y8 | 909.4884 | 78.75 |
| P39105 | PLB1 | DAGFN[+203.079373]IS-LADVWGR | 2 | heavy | 865.428 | y7 | 822.4564 | 78.75 |
| P39105 | PLB1 | DAGFN[+203.079373]IS-LADVWGR | 2 | heavy | 865.428 | y6 | 709.3723 | 78.75 |
| P39105 | PLB1 | DAGFN[+203.079373]IS-LADVWGR | 2 | heavy | 865.428 | y4 | 523.3083 | 78.75 |
| P39105 | PLB1 | N[+203.079373]LTDLEYIPPLI-VYIPNSR | 2 | light | 1217.149 | y12 | 1381.82 | 91.73 |
| P39105 | PLB1 | N[+203.079373]LTDLEYIPPLI-VYIPNSR | 2 | light | 1217.149 | y10 | 1171.683 | 91.73 |
| P39105 | PLB1 | N[+203.079373]LTDLEYIPPLI-VYIPNSR | 2 | light | 1217.149 | y9 | 1074.631 | 91.73 |

| | | | | | | | | |
|---|---|---|---|---|---|---|---|---|
| P39105 | PLB1 | N[+203.079373]LTDLEYIPPLI-VYIPNSR | 2 | light | 1217.149 | y8 | 961.5465 | 91.73 |
| P39105 | PLB1 | N[+203.079373]LTDLEYIPPLI-VYIPNSR | 2 | light | 1217.149 | y7 | 848.4625 | 91.73 |
| P39105 | PLB1 | N[+203.079373]LTDLEYIPPLI-VYIPNSR | 2 | light | 1217.149 | y6 | 749.3941 | 91.73 |
| P39105 | PLB1 | N[+203.079373]LTDLEYIPPLI-VYIPNSR | 2 | light | 1217.149 | y5 | 586.3307 | 91.73 |
| P39105 | PLB1 | N[+203.079373]LTDLEYIPPLI-VYIPNSR | 2 | light | 1217.149 | y4 | 473.2467 | 91.73 |
| P39105 | PLB1 | N[+203.079373]LTDLEYIPPLI-VYIPNSR | 2 | heavy | 1220.159 | y12 | 1387.84 | 91.73 |
| P39105 | PLB1 | N[+203.079373]LTDLEYIPPLI-VYIPNSR | 2 | heavy | 1220.159 | y10 | 1177.703 | 91.73 |
| P39105 | PLB1 | N[+203.079373]LTDLEYIPPLI-VYIPNSR | 2 | heavy | 1220.159 | y9 | 1080.651 | 91.73 |
| P39105 | PLB1 | N[+203.079373]LTDLEYIPPLI-VYIPNSR | 2 | heavy | 1220.159 | y8 | 967.5667 | 91.73 |
| P39105 | PLB1 | N[+203.079373]LTDLEYIPPLI-VYIPNSR | 2 | heavy | 1220.159 | y7 | 854.4826 | 91.73 |
| P39105 | PLB1 | N[+203.079373]LTDLEYIPPLI-VYIPNSR | 2 | heavy | 1220.159 | y6 | 755.4142 | 91.73 |
| P39105 | PLB1 | N[+203.079373]LTDLEYIPPLI-VYIPNSR | 2 | heavy | 1220.159 | y5 | 592.3509 | 91.73 |
| P39105 | PLB1 | N[+203.079373]LTDLEYIPPLI-VYIPNSR | 2 | heavy | 1220.159 | y4 | 479.2668 | 91.73 |
| P39105 | PLB1 | IPLVPLLQK | 2 | light | 510.8444 | y7 | 810.5448 | 70.7 |
| P39105 | PLB1 | IPLVPLLQK | 2 | light | 510.8444 | y6 | 697.4607 | 70.7 |
| P39105 | PLB1 | IPLVPLLQK | 2 | light | 510.8444 | y5 | 598.3923 | 70.7 |
| P39105 | PLB1 | IPLVPLLQK | 2 | light | 510.8444 | y7 | 405.776 | 70.7 |
| P39105 | PLB1 | IPLVPLLQK | 2 | heavy | 514.8515 | y7 | 818.559 | 70.7 |
| P39105 | PLB1 | IPLVPLLQK | 2 | heavy | 514.8515 | y6 | 705.4749 | 70.7 |
| P39105 | PLB1 | IPLVPLLQK | 2 | heavy | 514.8515 | y5 | 606.4065 | 70.7 |
| P39105 | PLB1 | IPLVPLLQK | 2 | heavy | 514.8515 | y7 | 409.7831 | 70.7 |
| P39105 | PLB1 | IAVAC[+57.021464]SGGGYR | 2 | light | 555.7717 | y10 | 997.452 | 28.11 |
| P39105 | PLB1 | IAVAC[+57.021464]SGGGYR | 2 | light | 555.7717 | y9 | 926.4149 | 28.11 |
| P39105 | PLB1 | IAVAC[+57.021464]SGGGYR | 2 | light | 555.7717 | y8 | 827.3465 | 28.11 |
| P39105 | PLB1 | IAVAC[+57.021464]SGGGYR | 2 | light | 555.7717 | y7 | 756.3093 | 28.11 |
| P39105 | PLB1 | IAVAC[+57.021464]SGGGYR | 2 | light | 555.7717 | y6 | 596.2787 | 28.11 |
| P39105 | PLB1 | IAVAC[+57.021464]SGGGYR | 2 | light | 555.7717 | y5 | 509.2467 | 28.11 |
| P39105 | PLB1 | IAVAC[+57.021464]SGGGYR | 2 | light | 555.7717 | b3 | 284.1969 | 28.11 |
| P39105 | PLB1 | IAVAC[+57.021464]SGGGYR | 2 | heavy | 558.7817 | y10 | 1003.472 | 28.11 |
| P39105 | PLB1 | IAVAC[+57.021464]SGGGYR | 2 | heavy | 558.7817 | y9 | 932.435 | 28.11 |
| P39105 | PLB1 | IAVAC[+57.021464]SGGGYR | 2 | heavy | 558.7817 | y8 | 833.3666 | 28.11 |
| P39105 | PLB1 | IAVAC[+57.021464]SGGGYR | 2 | heavy | 558.7817 | y7 | 762.3295 | 28.11 |

| | | | | | | | | |
|---|---|---|---|---|---|---|---|---|
| P39105 | PLB1 | IAVAC[+57.021464]SGGGYR | 2 | heavy | 558.7817 | y6 | 602.2988 | 28.11 |
| P39105 | PLB1 | IAVAC[+57.021464]SGGGYR | 2 | heavy | 558.7817 | y5 | 515.2668 | 28.11 |
| P39105 | PLB1 | IAVAC[+57.021464]SGGGYR | 2 | heavy | 558.7817 | b3 | 284.1969 | 28.11 |
| Q03674 | PLB2 | SIVNPGGSN[+203.079373]LTY-TIER | 2 | light | 962.4865 | y10 | 1356.664 | 51.85 |
| Q03674 | PLB2 | SIVNPGGSN[+203.079373]LTY-TIER | 2 | light | 962.4865 | y9 | 1299.643 | 51.85 |
| Q03674 | PLB2 | SIVNPGGSN[+203.079373]LTY-TIER | 2 | light | 962.4865 | y8 | 1212.611 | 51.85 |
| Q03674 | PLB2 | SIVNPGGSN[+203.079373]LTY-TIER | 2 | light | 962.4865 | y6 | 782.4043 | 51.85 |
| Q03674 | PLB2 | SIVNPGGSN[+203.079373]LTY-TIER | 2 | light | 962.4865 | y5 | 681.3566 | 51.85 |
| Q03674 | PLB2 | SIVNPGGSN[+203.079373]LTY-TIER | 2 | light | 962.4865 | y4 | 518.2933 | 51.85 |
| Q03674 | PLB2 | SIVNPGGSN[+203.079373]LTY-TIER | 2 | heavy | 965.4966 | y10 | 1362.684 | 51.85 |
| Q03674 | PLB2 | SIVNPGGSN[+203.079373]LTY-TIER | 2 | heavy | 965.4966 | y9 | 1305.663 | 51.85 |
| Q03674 | PLB2 | SIVNPGGSN[+203.079373]LTY-TIER | 2 | heavy | 965.4966 | y8 | 1218.631 | 51.85 |
| Q03674 | PLB2 | SIVNPGGSN[+203.079373]LTY-TIER | 2 | heavy | 965.4966 | y6 | 788.4244 | 51.85 |
| Q03674 | PLB2 | SIVNPGGSN[+203.079373]LTY-TIER | 2 | heavy | 965.4966 | y5 | 687.3767 | 51.85 |
| Q03674 | PLB2 | SIVNPGGSN[+203.079373]LTY-TIER | 2 | heavy | 965.4966 | y4 | 524.3134 | 51.85 |
| Q03674 | PLB2 | SDAGFN[+203.079373]ISLSDL-WAR | 2 | light | 927.9471 | y8 | 947.4945 | 79.73 |
| Q03674 | PLB2 | SDAGFN[+203.079373]ISLSDL-WAR | 2 | light | 927.9471 | y6 | 747.3784 | 79.73 |
| Q03674 | PLB2 | SDAGFN[+203.079373]ISLSDL-WAR | 2 | light | 927.9471 | y4 | 545.3194 | 79.73 |
| Q03674 | PLB2 | SDAGFN[+203.079373]ISLSDL-WAR | 2 | light | 927.9471 | b2 | 203.0662 | 79.73 |
| Q03674 | PLB2 | SDAGFN[+203.079373]ISLSDL-WAR | 2 | light | 927.9471 | b3 | 137.5553 | 79.73 |
| Q03674 | PLB2 | SDAGFN[+203.079373]ISLSDL-WAR | 2 | heavy | 930.9571 | y8 | 953.5146 | 79.73 |
| Q03674 | PLB2 | SDAGFN[+203.079373]ISLSDL-WAR | 2 | heavy | 930.9571 | y6 | 753.3985 | 79.73 |
| Q03674 | PLB2 | SDAGFN[+203.079373]ISLSDL-WAR | 2 | heavy | 930.9571 | y4 | 551.3396 | 79.73 |
| Q03674 | PLB2 | SDAGFN[+203.079373]ISLSDL-WAR | 2 | heavy | 930.9571 | b2 | 203.0662 | 79.73 |
| Q03674 | PLB2 | SDAGFN[+203.079373]ISLSDL-WAR | 2 | heavy | 930.9571 | b3 | 137.5553 | 79.73 |
| Q03674 | PLB2 | IGIAC[+57.021464]SGGGYR | 2 | light | 555.7717 | y10 | 997.452 | 31.5 |
| Q03674 | PLB2 | IGIAC[+57.021464]SGGGYR | 2 | light | 555.7717 | y8 | 827.3465 | 31.5 |
| Q03674 | PLB2 | IGIAC[+57.021464]SGGGYR | 2 | light | 555.7717 | y7 | 756.3093 | 31.5 |
| Q03674 | PLB2 | IGIAC[+57.021464]SGGGYR | 2 | light | 555.7717 | y6 | 596.2787 | 31.5 |
| Q03674 | PLB2 | IGIAC[+57.021464]SGGGYR | 2 | light | 555.7717 | y5 | 509.2467 | 31.5 |
| Q03674 | PLB2 | IGIAC[+57.021464]SGGGYR | 2 | light | 555.7717 | b3 | 284.1969 | 31.5 |
| Q03674 | PLB2 | IGIAC[+57.021464]SGGGYR | 2 | heavy | 558.7817 | y10 | 1003.472 | 31.5 |

| | | | | | | | | |
|---|---|---|---|---|---|---|---|---|
| Q03674 | PLB2 | IGIAC[+57.021464]SGGGYR | 2 | heavy | 558.7817 | y8 | 833.3666 | 31.5 |
| Q03674 | PLB2 | IGIAC[+57.021464]SGGGYR | 2 | heavy | 558.7817 | y7 | 762.3295 | 31.5 |
| Q03674 | PLB2 | IGIAC[+57.021464]SGGGYR | 2 | heavy | 558.7817 | y6 | 602.2988 | 31.5 |
| Q03674 | PLB2 | IGIAC[+57.021464]SGGGYR | 2 | heavy | 558.7817 | y5 | 515.2668 | 31.5 |
| Q03674 | PLB2 | IGIAC[+57.021464]SGGGYR | 2 | heavy | 558.7817 | b3 | 284.1969 | 31.5 |
| Q03674 | PLB2 | EALHSFLSR | 2 | light | 530.2827 | y7 | 859.4785 | 40.84 |
| Q03674 | PLB2 | EALHSFLSR | 2 | light | 530.2827 | y6 | 746.3944 | 40.84 |
| Q03674 | PLB2 | EALHSFLSR | 2 | light | 530.2827 | y5 | 609.3355 | 40.84 |
| Q03674 | PLB2 | EALHSFLSR | 2 | light | 530.2827 | y4 | 522.3035 | 40.84 |
| Q03674 | PLB2 | EALHSFLSR | 2 | light | 530.2827 | y3 | 375.235 | 40.84 |
| Q03674 | PLB2 | EALHSFLSR | 2 | light | 530.2827 | y6 | 373.7008 | 40.84 |
| Q03674 | PLB2 | EALHSFLSR | 2 | heavy | 533.2928 | y7 | 865.4986 | 40.84 |
| Q03674 | PLB2 | EALHSFLSR | 2 | heavy | 533.2928 | y6 | 752.4145 | 40.84 |
| Q03674 | PLB2 | EALHSFLSR | 2 | heavy | 533.2928 | y5 | 615.3556 | 40.84 |
| Q03674 | PLB2 | EALHSFLSR | 2 | heavy | 533.2928 | y4 | 528.3236 | 40.84 |
| Q03674 | PLB2 | EALHSFLSR | 2 | heavy | 533.2928 | y3 | 381.2552 | 40.84 |
| Q03674 | PLB2 | EALHSFLSR | 2 | heavy | 533.2928 | y6 | 376.7109 | 40.84 |
| P32623 | CRH2 | N[+203.079373]GTSAYVYTSSSEF-LAK | 2 | light | 1014.476 | y11 | 1231.62 | 53.26 |
| P32623 | CRH2 | N[+203.079373]GTSAYVYTSSSEF-LAK | 2 | light | 1014.476 | y10 | 1132.552 | 53.26 |
| P32623 | CRH2 | N[+203.079373]GTSAYVYTSSSEF-LAK | 2 | light | 1014.476 | y9 | 969.4888 | 53.26 |
| P32623 | CRH2 | N[+203.079373]GTSAYVYTSSSEF-LAK | 2 | light | 1014.476 | y7 | 781.409 | 53.26 |
| P32623 | CRH2 | N[+203.079373]GTSAYVYTSSSEF-LAK | 2 | light | 1014.476 | y4 | 478.3024 | 53.26 |
| P32623 | CRH2 | N[+203.079373]GTSAYVYTSSSEF-LAK | 2 | light | 1014.476 | y3 | 331.234 | 53.26 |
| P32623 | CRH2 | N[+203.079373]GTSAYVYTSSSEF-LAK | 2 | heavy | 1018.483 | y11 | 1239.635 | 53.26 |
| P32623 | CRH2 | N[+203.079373]GTSAYVYTSSSEF-LAK | 2 | heavy | 1018.483 | y10 | 1140.566 | 53.26 |
| P32623 | CRH2 | N[+203.079373]GTSAYVYTSSSEF-LAK | 2 | heavy | 1018.483 | y9 | 977.503 | 53.26 |
| P32623 | CRH2 | N[+203.079373]GTSAYVYTSSSEF-LAK | 2 | heavy | 1018.483 | y7 | 789.4232 | 53.26 |
| P32623 | CRH2 | N[+203.079373]GTSAYVYTSSSEF-LAK | 2 | heavy | 1018.483 | y4 | 486.3166 | 53.26 |
| P32623 | CRH2 | N[+203.079373]GTSAYVYTSSSEF-LAK | 2 | heavy | 1018.483 | y3 | 339.2482 | 53.26 |
| P32623 | CRH2 | NSGGTVLSSTR | 2 | light | 539.778 | y10 | 964.5058 | 22.43 |
| P32623 | CRH2 | NSGGTVLSSTR | 2 | light | 539.778 | y9 | 877.4738 | 22.43 |
| P32623 | CRH2 | NSGGTVLSSTR | 2 | light | 539.778 | y8 | 820.4523 | 22.43 |

| | | | | | | | | |
|---|---|---|---|---|---|---|---|---|
| P32623 | CRH2 | NSGGTVLSSTR | 2 | light | 539.778 | y7 | 763.4308 | 22.43 |
| P32623 | CRH2 | NSGGTVLSSTR | 2 | light | 539.778 | y6 | 662.3832 | 22.43 |
| P32623 | CRH2 | NSGGTVLSSTR | 2 | light | 539.778 | y5 | 563.3148 | 22.43 |
| P32623 | CRH2 | NSGGTVLSSTR | 2 | light | 539.778 | y4 | 450.2307 | 22.43 |
| P32623 | CRH2 | NSGGTVLSSTR | 2 | heavy | 542.7881 | y10 | 970.5259 | 22.43 |
| P32623 | CRH2 | NSGGTVLSSTR | 2 | heavy | 542.7881 | y9 | 883.4939 | 22.43 |
| P32623 | CRH2 | NSGGTVLSSTR | 2 | heavy | 542.7881 | y8 | 826.4724 | 22.43 |
| P32623 | CRH2 | NSGGTVLSSTR | 2 | heavy | 542.7881 | y7 | 769.451 | 22.43 |
| P32623 | CRH2 | NSGGTVLSSTR | 2 | heavy | 542.7881 | y6 | 668.4033 | 22.43 |
| P32623 | CRH2 | NSGGTVLSSTR | 2 | heavy | 542.7881 | y5 | 569.3349 | 22.43 |
| P32623 | CRH2 | NSGGTVLSSTR | 2 | heavy | 542.7881 | y4 | 456.2508 | 22.43 |
| P32623 | CRH2 | YQYPQTPSK | 2 | light | 556.2746 | y7 | 820.4199 | 26.64 |
| P32623 | CRH2 | YQYPQTPSK | 2 | light | 556.2746 | y6 | 657.3566 | 26.64 |
| P32623 | CRH2 | YQYPQTPSK | 2 | light | 556.2746 | y5 | 560.3039 | 26.64 |
| P32623 | CRH2 | YQYPQTPSK | 2 | light | 556.2746 | y4 | 432.2453 | 26.64 |
| P32623 | CRH2 | YQYPQTPSK | 2 | light | 556.2746 | y3 | 331.1976 | 26.64 |
| P32623 | CRH2 | YQYPQTPSK | 2 | light | 556.2746 | y2 | 234.1448 | 26.64 |
| P32623 | CRH2 | YQYPQTPSK | 2 | heavy | 560.2817 | y7 | 828.4341 | 26.64 |
| P32623 | CRH2 | YQYPQTPSK | 2 | heavy | 560.2817 | y6 | 665.3708 | 26.64 |
| P32623 | CRH2 | YQYPQTPSK | 2 | heavy | 560.2817 | y5 | 568.3181 | 26.64 |
| P32623 | CRH2 | YQYPQTPSK | 2 | heavy | 560.2817 | y4 | 440.2595 | 26.64 |
| P32623 | CRH2 | YQYPQTPSK | 2 | heavy | 560.2817 | y3 | 339.2118 | 26.64 |
| P32623 | CRH2 | YQYPQTPSK | 2 | heavy | 560.2817 | y2 | 242.159 | 26.64 |
| P54003 | SUR7 | FYWVQGN[+203.079373]TTGIP-NAGDETR | 2 | light | 1165.04 | y11 | 1130.544 | 57.45 |
| P54003 | SUR7 | FYWVQGN[+203.079373]TTGIP-NAGDETR | 2 | light | 1165.04 | y10 | 1029.496 | 57.45 |
| P54003 | SUR7 | FYWVQGN[+203.079373]TTGIP-NAGDETR | 2 | light | 1165.04 | y9 | 972.4745 | 57.45 |
| P54003 | SUR7 | FYWVQGN[+203.079373]TTGIP-NAGDETR | 2 | light | 1165.04 | y5 | 577.2576 | 57.45 |
| P54003 | SUR7 | FYWVQGN[+203.079373]TTGIP-NAGDETR | 2 | light | 1165.04 | y4 | 520.2362 | 57.45 |
| P54003 | SUR7 | FYWVQGN[+203.079373]TTGIP-NAGDETR | 2 | light | 1165.04 | y3 | 405.2092 | 57.45 |
| P54003 | SUR7 | FYWVQGN[+203.079373]TTGIP-NAGDETR | 2 | light | 1165.04 | y2 | 276.1666 | 57.45 |
| P54003 | SUR7 | FYWVQGN[+203.079373]TTGIP-NAGDETR | 2 | light | 1165.04 | b2 | 311.139 | 57.45 |
| P54003 | SUR7 | FYWVQGN[+203.079373]TTGIP-NAGDETR | 2 | light | 1165.04 | b3 | 497.2183 | 57.45 |

| | | | | | | | | |
|---|---|---|---|---|---|---|---|---|
| P54003 | SUR7 | FYWVQGN[+203.079373]TTGIP-NAGDETR | 2 | heavy | 1168.05 | y11 | 1136.564 | 57.45 |
| P54003 | SUR7 | FYWVQGN[+203.079373]TTGIP-NAGDETR | 2 | heavy | 1168.05 | y10 | 1035.516 | 57.45 |
| P54003 | SUR7 | FYWVQGN[+203.079373]TTGIP-NAGDETR | 2 | heavy | 1168.05 | y9 | 978.4946 | 57.45 |
| P54003 | SUR7 | FYWVQGN[+203.079373]TTGIP-NAGDETR | 2 | heavy | 1168.05 | y5 | 583.2778 | 57.45 |
| P54003 | SUR7 | FYWVQGN[+203.079373]TTGIP-NAGDETR | 2 | heavy | 1168.05 | y4 | 526.2563 | 57.45 |
| P54003 | SUR7 | FYWVQGN[+203.079373]TTGIP-NAGDETR | 2 | heavy | 1168.05 | y3 | 411.2294 | 57.45 |
| P54003 | SUR7 | FYWVQGN[+203.079373]TTGIP-NAGDETR | 2 | heavy | 1168.05 | y2 | 282.1868 | 57.45 |
| P54003 | SUR7 | FYWVQGN[+203.079373]TTGIP-NAGDETR | 2 | heavy | 1168.05 | b2 | 311.139 | 57.45 |
| P54003 | SUR7 | FYWVQGN[+203.079373]TTGIP-NAGDETR | 2 | heavy | 1168.05 | b3 | 497.2183 | 57.45 |
| P54003 | SUR7 | LASTYSIDNSR | 2 | light | 613.8042 | y10 | 1113.517 | 31.62 |
| P54003 | SUR7 | LASTYSIDNSR | 2 | light | 613.8042 | y9 | 1042.48 | 31.62 |
| P54003 | SUR7 | LASTYSIDNSR | 2 | light | 613.8042 | y8 | 955.448 | 31.62 |
| P54003 | SUR7 | LASTYSIDNSR | 2 | light | 613.8042 | y7 | 854.4003 | 31.62 |
| P54003 | SUR7 | LASTYSIDNSR | 2 | light | 613.8042 | y6 | 691.3369 | 31.62 |
| P54003 | SUR7 | LASTYSIDNSR | 2 | light | 613.8042 | y5 | 604.3049 | 31.62 |
| P54003 | SUR7 | LASTYSIDNSR | 2 | light | 613.8042 | y4 | 491.2209 | 31.62 |
| P54003 | SUR7 | LASTYSIDNSR | 2 | light | 613.8042 | y3 | 376.1939 | 31.62 |
| P54003 | SUR7 | LASTYSIDNSR | 2 | heavy | 616.8143 | y10 | 1119.537 | 31.62 |
| P54003 | SUR7 | LASTYSIDNSR | 2 | heavy | 616.8143 | y9 | 1048.5 | 31.62 |
| P54003 | SUR7 | LASTYSIDNSR | 2 | heavy | 616.8143 | y8 | 961.4681 | 31.62 |
| P54003 | SUR7 | LASTYSIDNSR | 2 | heavy | 616.8143 | y7 | 860.4204 | 31.62 |
| P54003 | SUR7 | LASTYSIDNSR | 2 | heavy | 616.8143 | y6 | 697.3571 | 31.62 |
| P54003 | SUR7 | LASTYSIDNSR | 2 | heavy | 616.8143 | y5 | 610.325 | 31.62 |
| P54003 | SUR7 | LASTYSIDNSR | 2 | heavy | 616.8143 | y4 | 497.241 | 31.62 |
| P54003 | SUR7 | LASTYSIDNSR | 2 | heavy | 616.8143 | y3 | 382.214 | 31.62 |
| P54003 | SUR7 | WTFWGAC[+57.021464]LQDK | 2 | light | 706.3268 | y9 | 1124.519 | 70.48 |
| P54003 | SUR7 | WTFWGAC[+57.021464]LQDK | 2 | light | 706.3268 | y8 | 977.4509 | 70.48 |
| P54003 | SUR7 | WTFWGAC[+57.021464]LQDK | 2 | light | 706.3268 | y7 | 791.3716 | 70.48 |
| P54003 | SUR7 | WTFWGAC[+57.021464]LQDK | 2 | light | 706.3268 | y6 | 734.3502 | 70.48 |
| P54003 | SUR7 | WTFWGAC[+57.021464]LQDK | 2 | light | 706.3268 | y5 | 663.313 | 70.48 |
| P54003 | SUR7 | WTFWGAC[+57.021464]LQDK | 2 | light | 706.3268 | y3 | 390.1983 | 70.48 |
| P54003 | SUR7 | WTFWGAC[+57.021464]LQDK | 2 | light | 706.3268 | y2 | 262.1397 | 70.48 |

| | | | | | | | | |
|---|---|---|---|---|---|---|---|---|
| P54003 | SUR7 | WTFWGAC[+57.021464]LQDK | 2 | light | 706.3268 | b2 | 288.1343 | 70.48 |
| P54003 | SUR7 | WTFWGAC[+57.021464]LQDK | 2 | heavy | 710.3339 | y9 | 1132.534 | 70.48 |
| P54003 | SUR7 | WTFWGAC[+57.021464]LQDK | 2 | heavy | 710.3339 | y8 | 985.4651 | 70.48 |
| P54003 | SUR7 | WTFWGAC[+57.021464]LQDK | 2 | heavy | 710.3339 | y7 | 799.3858 | 70.48 |
| P54003 | SUR7 | WTFWGAC[+57.021464]LQDK | 2 | heavy | 710.3339 | y6 | 742.3644 | 70.48 |
| P54003 | SUR7 | WTFWGAC[+57.021464]LQDK | 2 | heavy | 710.3339 | y5 | 671.3272 | 70.48 |
| P54003 | SUR7 | WTFWGAC[+57.021464]LQDK | 2 | heavy | 710.3339 | y3 | 398.2125 | 70.48 |
| P54003 | SUR7 | WTFWGAC[+57.021464]LQDK | 2 | heavy | 710.3339 | y2 | 270.1539 | 70.48 |
| P54003 | SUR7 | WTFWGAC[+57.021464]LQDK | 2 | heavy | 710.3339 | b2 | 288.1343 | 70.48 |
| P33754 | SEC66 | FSNN[+203.079373]GTFFETEEPI-VETK | 2 | light | 1146.531 | y11 | 1321.652 | 60.28 |
| P33754 | SEC66 | FSNN[+203.079373]GTFFETEEPI-VETK | 2 | light | 1146.531 | y10 | 1174.584 | 60.28 |
| P33754 | SEC66 | FSNN[+203.079373]GTFFETEEPI-VETK | 2 | light | 1146.531 | y9 | 1045.541 | 60.28 |
| P33754 | SEC66 | FSNN[+203.079373]GTFFETEEPI-VETK | 2 | light | 1146.531 | y8 | 944.4935 | 60.28 |
| P33754 | SEC66 | FSNN[+203.079373]GTFFETEEPI-VETK | 2 | light | 1146.531 | y6 | 686.4083 | 60.28 |
| P33754 | SEC66 | FSNN[+203.079373]GTFFETEEPI-VETK | 2 | light | 1146.531 | y4 | 476.2715 | 60.28 |
| P33754 | SEC66 | FSNN[+203.079373]GTFFETEEPI-VETK | 2 | light | 1146.531 | y2 | 248.1605 | 60.28 |
| P33754 | SEC66 | FSNN[+203.079373]GTFFETEEPI-VETK | 2 | light | 1146.531 | b4 | 463.1936 | 60.28 |
| P33754 | SEC66 | FSNN[+203.079373]GTFFETEEPI-VETK | 2 | light | 1146.531 | b5 | 520.215 | 60.28 |
| P33754 | SEC66 | FSNN[+203.079373]GTFFETEEPI-VETK | 2 | heavy | 1150.538 | y11 | 1329.666 | 60.28 |
| P33754 | SEC66 | FSNN[+203.079373]GTFFETEEPI-VETK | 2 | heavy | 1150.538 | y10 | 1182.598 | 60.28 |
| P33754 | SEC66 | FSNN[+203.079373]GTFFETEEPI-VETK | 2 | heavy | 1150.538 | y9 | 1053.555 | 60.28 |
| P33754 | SEC66 | FSNN[+203.079373]GTFFETEEPI-VETK | 2 | heavy | 1150.538 | y8 | 952.5077 | 60.28 |
| P33754 | SEC66 | FSNN[+203.079373]GTFFETEEPI-VETK | 2 | heavy | 1150.538 | y6 | 694.4225 | 60.28 |
| P33754 | SEC66 | FSNN[+203.079373]GTFFETEEPI-VETK | 2 | heavy | 1150.538 | y4 | 484.2857 | 60.28 |
| P33754 | SEC66 | FSNN[+203.079373]GTFFETEEPI-VETK | 2 | heavy | 1150.538 | y2 | 256.1747 | 60.28 |
| P33754 | SEC66 | FSNN[+203.079373]GTFFETEEPI-VETK | 2 | heavy | 1150.538 | b4 | 463.1936 | 60.28 |
| P33754 | SEC66 | FSNN[+203.079373]GTFFETEEPI-VETK | 2 | heavy | 1150.538 | b5 | 520.215 | 60.28 |
| P33754 | SEC66 | EIC[+57.021464]FNQALSR | 2 | light | 619.3033 | y8 | 995.4727 | 47.65 |
| P33754 | SEC66 | EIC[+57.021464]FNQALSR | 2 | light | 619.3033 | y7 | 835.4421 | 47.65 |
| P33754 | SEC66 | EIC[+57.021464]FNQALSR | 2 | light | 619.3033 | y6 | 688.3737 | 47.65 |
| P33754 | SEC66 | EIC[+57.021464]FNQALSR | 2 | light | 619.3033 | y5 | 574.3307 | 47.65 |
| P33754 | SEC66 | EIC[+57.021464]FNQALSR | 2 | light | 619.3033 | y4 | 446.2722 | 47.65 |

| | | | | | | | | |
|---|---|---|---|---|---|---|---|---|
| P33754 | SEC66 | EIC[+57.021464]FNQALSR | 2 | light | 619.3033 | y3 | 375.235 | 47.65 |
| P33754 | SEC66 | EIC[+57.021464]FNQALSR | 2 | heavy | 622.3134 | y8 | 1001.493 | 47.65 |
| P33754 | SEC66 | EIC[+57.021464]FNQALSR | 2 | heavy | 622.3134 | y7 | 841.4622 | 47.65 |
| P33754 | SEC66 | EIC[+57.021464]FNQALSR | 2 | heavy | 622.3134 | y6 | 694.3938 | 47.65 |
| P33754 | SEC66 | EIC[+57.021464]FNQALSR | 2 | heavy | 622.3134 | y5 | 580.3509 | 47.65 |
| P33754 | SEC66 | EIC[+57.021464]FNQALSR | 2 | heavy | 622.3134 | y4 | 452.2923 | 47.65 |
| P33754 | SEC66 | EIC[+57.021464]FNQALSR | 2 | heavy | 622.3134 | y3 | 381.2552 | 47.65 |
| P33754 | SEC66 | LIELEFK | 2 | light | 446.2629 | y6 | 778.4345 | 58.54 |
| P33754 | SEC66 | LIELEFK | 2 | light | 446.2629 | y5 | 665.3505 | 58.54 |
| P33754 | SEC66 | LIELEFK | 2 | light | 446.2629 | y4 | 536.3079 | 58.54 |
| P33754 | SEC66 | LIELEFK | 2 | light | 446.2629 | y3 | 423.2238 | 58.54 |
| P33754 | SEC66 | LIELEFK | 2 | light | 446.2629 | y2 | 294.1812 | 58.54 |
| P33754 | SEC66 | LIELEFK | 2 | light | 446.2629 | y6 | 389.7209 | 58.54 |
| P33754 | SEC66 | LIELEFK | 2 | heavy | 450.27 | y6 | 786.4487 | 58.54 |
| P33754 | SEC66 | LIELEFK | 2 | heavy | 450.27 | y5 | 673.3647 | 58.54 |
| P33754 | SEC66 | LIELEFK | 2 | heavy | 450.27 | y4 | 544.3221 | 58.54 |
| P33754 | SEC66 | LIELEFK | 2 | heavy | 450.27 | y3 | 431.238 | 58.54 |
| P33754 | SEC66 | LIELEFK | 2 | heavy | 450.27 | y2 | 302.1954 | 58.54 |
| P33754 | SEC66 | LIELEFK | 2 | heavy | 450.27 | y6 | 393.728 | 58.54 |
| P32353 | ERG3 | LLGLNSGFSN[+203.079373]STIL-QETLNSK | 2 | light | 1220.134 | y10 | 1146.636 | 74.93 |
| P32353 | ERG3 | LLGLNSGFSN[+203.079373]STIL-QETLNSK | 2 | light | 1220.134 | y9 | 1045.589 | 74.93 |
| P32353 | ERG3 | LLGLNSGFSN[+203.079373]STIL-QETLNSK | 2 | light | 1220.134 | y8 | 932.5047 | 74.93 |
| P32353 | ERG3 | LLGLNSGFSN[+203.079373]STIL-QETLNSK | 2 | light | 1220.134 | y7 | 819.4207 | 74.93 |
| P32353 | ERG3 | LLGLNSGFSN[+203.079373]STIL-QETLNSK | 2 | light | 1220.134 | y6 | 691.3621 | 74.93 |
| P32353 | ERG3 | LLGLNSGFSN[+203.079373]STIL-QETLNSK | 2 | light | 1220.134 | y5 | 562.3195 | 74.93 |
| P32353 | ERG3 | LLGLNSGFSN[+203.079373]STIL-QETLNSK | 2 | light | 1220.134 | y4 | 461.2718 | 74.93 |
| P32353 | ERG3 | LLGLNSGFSN[+203.079373]STIL-QETLNSK | 2 | light | 1220.134 | y3 | 348.1878 | 74.93 |
| P32353 | ERG3 | LLGLNSGFSN[+203.079373]STIL-QETLNSK | 2 | heavy | 1224.141 | y10 | 1154.651 | 74.93 |
| P32353 | ERG3 | LLGLNSGFSN[+203.079373]STIL-QETLNSK | 2 | heavy | 1224.141 | y9 | 1053.603 | 74.93 |
| P32353 | ERG3 | LLGLNSGFSN[+203.079373]STIL-QETLNSK | 2 | heavy | 1224.141 | y8 | 940.5189 | 74.93 |
| P32353 | ERG3 | LLGLNSGFSN[+203.079373]STIL-QETLNSK | 2 | heavy | 1224.141 | y7 | 827.4349 | 74.93 |
| P32353 | ERG3 | LLGLNSGFSN[+203.079373]STIL-QETLNSK | 2 | heavy | 1224.141 | y6 | 699.3763 | 74.93 |

| | | | | | | | | |
|---|---|---|---|---|---|---|---|---|
| P32353 | ERG3 | LLGLNSGFSN[+203.079373]STIL-QETLNSK | 2 | heavy | 1224.141 | y5 | 570.3337 | 74.93 |
| P32353 | ERG3 | LLGLNSGFSN[+203.079373]STIL-QETLNSK | 2 | heavy | 1224.141 | y4 | 469.286 | 74.93 |
| P32353 | ERG3 | LLGLNSGFSN[+203.079373]STIL-QETLNSK | 2 | heavy | 1224.141 | y3 | 356.202 | 74.93 |
| P32353 | ERG3 | VLPASLAANIPVK | 2 | light | 646.9005 | y11 | 1080.641 | 62.01 |
| P32353 | ERG3 | VLPASLAANIPVK | 2 | light | 646.9005 | y10 | 983.5884 | 62.01 |
| P32353 | ERG3 | VLPASLAANIPVK | 2 | light | 646.9005 | y9 | 912.5513 | 62.01 |
| P32353 | ERG3 | VLPASLAANIPVK | 2 | light | 646.9005 | y7 | 712.4352 | 62.01 |
| P32353 | ERG3 | VLPASLAANIPVK | 2 | light | 646.9005 | y6 | 641.3981 | 62.01 |
| P32353 | ERG3 | VLPASLAANIPVK | 2 | light | 646.9005 | y3 | 343.234 | 62.01 |
| P32353 | ERG3 | VLPASLAANIPVK | 2 | light | 646.9005 | y11 | 540.8242 | 62.01 |
| P32353 | ERG3 | VLPASLAANIPVK | 2 | heavy | 650.9076 | y11 | 1088.655 | 62.01 |
| P32353 | ERG3 | VLPASLAANIPVK | 2 | heavy | 650.9076 | y10 | 991.6026 | 62.01 |
| P32353 | ERG3 | VLPASLAANIPVK | 2 | heavy | 650.9076 | y9 | 920.5655 | 62.01 |
| P32353 | ERG3 | VLPASLAANIPVK | 2 | heavy | 650.9076 | y7 | 720.4494 | 62.01 |
| P32353 | ERG3 | VLPASLAANIPVK | 2 | heavy | 650.9076 | y6 | 649.4123 | 62.01 |
| P32353 | ERG3 | VLPASLAANIPVK | 2 | heavy | 650.9076 | y3 | 351.2482 | 62.01 |
| P32353 | ERG3 | VLPASLAANIPVK | 2 | heavy | 650.9076 | y11 | 544.8313 | 62.01 |
| P38248 | ECM33 | VQTVGGAIEVTGN[+203.079373]FSTLDLSSLK | 2 | light | 1270.161 | y9 | 963.5357 | 74.18 |
| P38248 | ECM33 | VQTVGGAIEVTGN[+203.079373]FSTLDLSSLK | 2 | light | 1270.161 | y7 | 775.456 | 74.18 |
| P38248 | ECM33 | VQTVGGAIEVTGN[+203.079373]FSTLDLSSLK | 2 | light | 1270.161 | y6 | 662.3719 | 74.18 |
| P38248 | ECM33 | VQTVGGAIEVTGN[+203.079373]FSTLDLSSLK | 2 | light | 1270.161 | y5 | 547.345 | 74.18 |
| P38248 | ECM33 | VQTVGGAIEVTGN[+203.079373]FSTLDLSSLK | 2 | light | 1270.161 | y4 | 434.2609 | 74.18 |
| P38248 | ECM33 | VQTVGGAIEVTGN[+203.079373]FSTLDLSSLK | 2 | light | 1270.161 | y3 | 347.2289 | 74.18 |
| P38248 | ECM33 | VQTVGGAIEVTGN[+203.079373]FSTLDLSSLK | 2 | heavy | 1274.168 | y9 | 971.5499 | 74.18 |
| P38248 | ECM33 | VQTVGGAIEVTGN[+203.079373]FSTLDLSSLK | 2 | heavy | 1274.168 | y7 | 783.4702 | 74.18 |
| P38248 | ECM33 | VQTVGGAIEVTGN[+203.079373]FSTLDLSSLK | 2 | heavy | 1274.168 | y6 | 670.3861 | 74.18 |
| P38248 | ECM33 | VQTVGGAIEVTGN[+203.079373]FSTLDLSSLK | 2 | heavy | 1274.168 | y5 | 555.3592 | 74.18 |
| P38248 | ECM33 | VQTVGGAIEVTGN[+203.079373]FSTLDLSSLK | 2 | heavy | 1274.168 | y4 | 442.2751 | 74.18 |
| P38248 | ECM33 | VQTVGGAIEVTGN[+203.079373]FSTLDLSSLK | 2 | heavy | 1274.168 | y3 | 355.2431 | 74.18 |
| P38248 | ECM33 | VNVFNINNNR | 2 | light | 602.3151 | y8 | 990.5116 | 45.88 |
| P38248 | ECM33 | VNVFNINNNR | 2 | light | 602.3151 | y7 | 891.4431 | 45.88 |
| P38248 | ECM33 | VNVFNINNNR | 2 | light | 602.3151 | y6 | 744.3747 | 45.88 |

| | | | | | | | | |
|---|---|---|---|---|---|---|---|---|
| P38248 | ECM33 | VNVFNINNNR | 2 | light | 602.3151 | y5 | 630.3318 | 45.88 |
| P38248 | ECM33 | VNVFNINNNR | 2 | light | 602.3151 | y4 | 517.2477 | 45.88 |
| P38248 | ECM33 | VNVFNINNNR | 2 | light | 602.3151 | b2 | 214.1186 | 45.88 |
| P38248 | ECM33 | VNVFNINNNR | 2 | light | 602.3151 | b3 | 313.187 | 45.88 |
| P38248 | ECM33 | VNVFNINNNR | 2 | heavy | 605.3252 | y8 | 996.5317 | 45.88 |
| P38248 | ECM33 | VNVFNINNNR | 2 | heavy | 605.3252 | y7 | 897.4633 | 45.88 |
| P38248 | ECM33 | VNVFNINNNR | 2 | heavy | 605.3252 | y6 | 750.3949 | 45.88 |
| P38248 | ECM33 | VNVFNINNNR | 2 | heavy | 605.3252 | y5 | 636.3519 | 45.88 |
| P38248 | ECM33 | VNVFNINNNR | 2 | heavy | 605.3252 | y4 | 523.2679 | 45.88 |
| P38248 | ECM33 | VNVFNINNNR | 2 | heavy | 605.3252 | b2 | 214.1186 | 45.88 |
| P38248 | ECM33 | VNVFNINNNR | 2 | heavy | 605.3252 | b3 | 313.187 | 45.88 |
| P38248 | ECM33 | VGQSLSIVSNDELSK | 2 | light | 788.4149 | y10 | 1091.558 | 49.13 |
| P38248 | ECM33 | VGQSLSIVSNDELSK | 2 | light | 788.4149 | y8 | 891.4418 | 49.13 |
| P38248 | ECM33 | VGQSLSIVSNDELSK | 2 | light | 788.4149 | y7 | 792.3734 | 49.13 |
| P38248 | ECM33 | VGQSLSIVSNDELSK | 2 | light | 788.4149 | y5 | 591.2984 | 49.13 |
| P38248 | ECM33 | VGQSLSIVSNDELSK | 2 | light | 788.4149 | y4 | 476.2715 | 49.13 |
| P38248 | ECM33 | VGQSLSIVSNDELSK | 2 | light | 788.4149 | b3 | 285.1557 | 49.13 |
| P38248 | ECM33 | VGQSLSIVSNDELSK | 2 | heavy | 792.422 | y10 | 1099.572 | 49.13 |
| P38248 | ECM33 | VGQSLSIVSNDELSK | 2 | heavy | 792.422 | y8 | 899.456 | 49.13 |
| P38248 | ECM33 | VGQSLSIVSNDELSK | 2 | heavy | 792.422 | y7 | 800.3876 | 49.13 |
| P38248 | ECM33 | VGQSLSIVSNDELSK | 2 | heavy | 792.422 | y5 | 599.3126 | 49.13 |
| P38248 | ECM33 | VGQSLSIVSNDELSK | 2 | heavy | 792.422 | y4 | 484.2857 | 49.13 |
| P38248 | ECM33 | VGQSLSIVSNDELSK | 2 | heavy | 792.422 | b3 | 285.1557 | 49.13 |
| P33302 | PDR5 | GPAYAN[+203.079373]ISSTESVC[+57.021464]TVVGAVPGQ-DYVLGDDFIR | 3 | light | 1220.917 | y9 | 1097.563 | 78.91 |
| P33302 | PDR5 | GPAYAN[+203.079373]ISSTESVC[+57.021464]TVVGAVPGQ-DYVLGDDFIR | 3 | light | 1220.917 | y8 | 934.4993 | 78.91 |
| P33302 | PDR5 | GPAYAN[+203.079373]ISSTESVC[+57.021464]TVVGAVPGQ-DYVLGDDFIR | 3 | light | 1220.917 | y7 | 835.4308 | 78.91 |
| P33302 | PDR5 | GPAYAN[+203.079373]ISSTESVC[+57.021464]TVVGAVPGQ-DYVLGDDFIR | 3 | light | 1220.917 | y6 | 722.3468 | 78.91 |
| P33302 | PDR5 | GPAYAN[+203.079373]ISSTESVC[+57.021464]TVVGAVPGQ-DYVLGDDFIR | 3 | light | 1220.917 | y5 | 665.3253 | 78.91 |
| P33302 | PDR5 | GPAYAN[+203.079373]ISSTESVC[+57.021464]TVVGAVPGQ-DYVLGDDFIR | 3 | light | 1220.917 | y3 | 435.2714 | 78.91 |

| | | | | | | | | |
|---|---|---|---|---|---|---|---|---|
| P33302 | PDR5 | GPAYAN[+203.079373]ISSTESVC[+57.021464]TVVGAVPGQ-DYVLGDDFIR | 3 | light | 1220.917 | b2 | 155.0815 | 78.91 |
| P33302 | PDR5 | GPAYAN[+203.079373]ISSTESVC[+57.021464]TVVGAVPGQ-DYVLGDDFIR | 3 | heavy | 1222.924 | y9 | 1103.583 | 78.91 |
| P33302 | PDR5 | GPAYAN[+203.079373]ISSTESVC[+57.021464]TVVGAVPGQ-DYVLGDDFIR | 3 | heavy | 1222.924 | y8 | 940.5194 | 78.91 |
| P33302 | PDR5 | GPAYAN[+203.079373]ISSTESVC[+57.021464]TVVGAVPGQ-DYVLGDDFIR | 3 | heavy | 1222.924 | y7 | 841.451 | 78.91 |
| P33302 | PDR5 | GPAYAN[+203.079373]ISSTESVC[+57.021464]TVVGAVPGQ-DYVLGDDFIR | 3 | heavy | 1222.924 | y6 | 728.3669 | 78.91 |
| P33302 | PDR5 | GPAYAN[+203.079373]ISSTESVC[+57.021464]TVVGAVPGQ-DYVLGDDFIR | 3 | heavy | 1222.924 | y5 | 671.3454 | 78.91 |
| P33302 | PDR5 | GPAYAN[+203.079373]ISSTESVC[+57.021464]TVVGAVPGQ-DYVLGDDFIR | 3 | heavy | 1222.924 | y3 | 441.2916 | 78.91 |
| P33302 | PDR5 | GPAYAN[+203.079373]ISSTESVC[+57.021464]TVVGAVPGQ-DYVLGDDFIR | 3 | heavy | 1222.924 | b2 | 155.0815 | 78.91 |
| P33302 | PDR5 | AVQSELDWMER | 2 | light | 682.3192 | y9 | 1193.526 | 56 |
| P33302 | PDR5 | AVQSELDWMER | 2 | light | 682.3192 | y8 | 1065.467 | 56 |
| P33302 | PDR5 | AVQSELDWMER | 2 | light | 682.3192 | y7 | 978.4349 | 56 |
| P33302 | PDR5 | AVQSELDWMER | 2 | light | 682.3192 | y6 | 849.3924 | 56 |
| P33302 | PDR5 | AVQSELDWMER | 2 | light | 682.3192 | y5 | 736.3083 | 56 |
| P33302 | PDR5 | AVQSELDWMER | 2 | light | 682.3192 | y4 | 621.2813 | 56 |
| P33302 | PDR5 | AVQSELDWMER | 2 | heavy | 685.3292 | y9 | 1199.546 | 56 |
| P33302 | PDR5 | AVQSELDWMER | 2 | heavy | 685.3292 | y8 | 1071.487 | 56 |
| P33302 | PDR5 | AVQSELDWMER | 2 | heavy | 685.3292 | y7 | 984.4551 | 56 |
| P33302 | PDR5 | AVQSELDWMER | 2 | heavy | 685.3292 | y6 | 855.4125 | 56 |
| P33302 | PDR5 | AVQSELDWMER | 2 | heavy | 685.3292 | y5 | 742.3284 | 56 |
| P33302 | PDR5 | AVQSELDWMER | 2 | heavy | 685.3292 | y4 | 627.3015 | 56 |
| P33302 | PDR5 | QTTADFLTSVTSPSER | 2 | light | 870.426 | y10 | 1076.558 | 57.52 |
| P33302 | PDR5 | QTTADFLTSVTSPSER | 2 | light | 870.426 | y9 | 963.4742 | 57.52 |
| P33302 | PDR5 | QTTADFLTSVTSPSER | 2 | light | 870.426 | y8 | 862.4265 | 57.52 |
| P33302 | PDR5 | QTTADFLTSVTSPSER | 2 | light | 870.426 | y6 | 676.326 | 57.52 |
| P33302 | PDR5 | QTTADFLTSVTSPSER | 2 | light | 870.426 | y4 | 488.2463 | 57.52 |
| P33302 | PDR5 | QTTADFLTSVTSPSER | 2 | heavy | 873.436 | y10 | 1082.578 | 57.52 |
| P33302 | PDR5 | QTTADFLTSVTSPSER | 2 | heavy | 873.436 | y9 | 969.4943 | 57.52 |
| P33302 | PDR5 | QTTADFLTSVTSPSER | 2 | heavy | 873.436 | y8 | 868.4466 | 57.52 |

| | | | | | | | | |
|---|---|---|---|---|---|---|---|---|
| P33302 | PDR5 | QTTADFLTSVTSPSER | 2 | heavy | 873.436 | y6 | 682.3462 | 57.52 |
| P33302 | PDR5 | QTTADFLTSVTSPSER | 2 | heavy | 873.436 | y4 | 494.2665 | 57.52 |
| P31382 | PMT2 | GLPSWSEN[+203.079373]ET-DIEYLKPGTSYR | 3 | light | 915.7679 | y11 | 1326.705 | 60.71 |
| P31382 | PMT2 | GLPSWSEN[+203.079373]ET-DIEYLKPGTSYR | 3 | light | 915.7679 | y10 | 1213.621 | 60.71 |
| P31382 | PMT2 | GLPSWSEN[+203.079373]ET-DIEYLKPGTSYR | 3 | light | 915.7679 | y9 | 1084.579 | 60.71 |
| P31382 | PMT2 | GLPSWSEN[+203.079373]ET-DIEYLKPGTSYR | 3 | light | 915.7679 | y8 | 921.5152 | 60.71 |
| P31382 | PMT2 | GLPSWSEN[+203.079373]ET-DIEYLKPGTSYR | 3 | light | 915.7679 | y7 | 808.4312 | 60.71 |
| P31382 | PMT2 | GLPSWSEN[+203.079373]ET-DIEYLKPGTSYR | 3 | light | 915.7679 | y6 | 680.3362 | 60.71 |
| P31382 | PMT2 | GLPSWSEN[+203.079373]ET-DIEYLKPGTSYR | 3 | light | 915.7679 | y3 | 425.2143 | 60.71 |
| P31382 | PMT2 | GLPSWSEN[+203.079373]ET-DIEYLKPGTSYR | 3 | heavy | 920.446 | y11 | 1340.74 | 60.71 |
| P31382 | PMT2 | GLPSWSEN[+203.079373]ET-DIEYLKPGTSYR | 3 | heavy | 920.446 | y10 | 1227.655 | 60.71 |
| P31382 | PMT2 | GLPSWSEN[+203.079373]ET-DIEYLKPGTSYR | 3 | heavy | 920.446 | y9 | 1098.613 | 60.71 |
| P31382 | PMT2 | GLPSWSEN[+203.079373]ET-DIEYLKPGTSYR | 3 | heavy | 920.446 | y8 | 935.5496 | 60.71 |
| P31382 | PMT2 | GLPSWSEN[+203.079373]ET-DIEYLKPGTSYR | 3 | heavy | 920.446 | y7 | 822.4655 | 60.71 |
| P31382 | PMT2 | GLPSWSEN[+203.079373]ET-DIEYLKPGTSYR | 3 | heavy | 920.446 | y6 | 686.3563 | 60.71 |
| P31382 | PMT2 | GLPSWSEN[+203.079373]ET-DIEYLKPGTSYR | 3 | heavy | 920.446 | y3 | 431.2344 | 60.71 |
| P31382 | PMT2 | EKPAAQSSLLR | 2 | light | 600.3408 | y9 | 942.5367 | 26.71 |
| P31382 | PMT2 | EKPAAQSSLLR | 2 | light | 600.3408 | y8 | 845.4839 | 26.71 |
| P31382 | PMT2 | EKPAAQSSLLR | 2 | light | 600.3408 | y7 | 774.4468 | 26.71 |
| P31382 | PMT2 | EKPAAQSSLLR | 2 | light | 600.3408 | y6 | 703.4097 | 26.71 |
| P31382 | PMT2 | EKPAAQSSLLR | 2 | light | 600.3408 | y5 | 575.3511 | 26.71 |
| P31382 | PMT2 | EKPAAQSSLLR | 2 | heavy | 607.3579 | y9 | 948.5568 | 26.71 |
| P31382 | PMT2 | EKPAAQSSLLR | 2 | heavy | 607.3579 | y8 | 851.5041 | 26.71 |
| P31382 | PMT2 | EKPAAQSSLLR | 2 | heavy | 607.3579 | y7 | 780.467 | 26.71 |
| P31382 | PMT2 | EKPAAQSSLLR | 2 | heavy | 607.3579 | y6 | 709.4298 | 26.71 |
| P31382 | PMT2 | EKPAAQSSLLR | 2 | heavy | 607.3579 | y5 | 581.3713 | 26.71 |
| P31382 | PMT2 | NLHTHPVAAPVSK | 2 | light | 685.8806 | y9 | 905.5203 | 21.38 |
| P31382 | PMT2 | NLHTHPVAAPVSK | 2 | light | 685.8806 | y8 | 768.4614 | 21.38 |
| P31382 | PMT2 | NLHTHPVAAPVSK | 2 | light | 685.8806 | y4 | 430.266 | 21.38 |
| P31382 | PMT2 | NLHTHPVAAPVSK | 2 | light | 685.8806 | y2 | 234.1448 | 21.38 |
| P31382 | PMT2 | NLHTHPVAAPVSK | 2 | light | 685.8806 | y4 | 215.6366 | 21.38 |
| P31382 | PMT2 | NLHTHPVAAPVSK | 2 | light | 685.8806 | b3 | 365.1932 | 21.38 |

| | | | | | | | | |
|---|---|---|---|---|---|---|---|---|
| P31382 | PMT2 | NLHTHPVAAPVSK | 2 | light | 685.8806 | b5 | 603.2998 | 21.38 |
| P31382 | PMT2 | NLHTHPVAAPVSK | 2 | light | 685.8806 | b9 | 941.4952 | 21.38 |
| P31382 | PMT2 | NLHTHPVAAPVSK | 2 | heavy | 689.8877 | y9 | 913.5345 | 21.38 |
| P31382 | PMT2 | NLHTHPVAAPVSK | 2 | heavy | 689.8877 | y8 | 776.4756 | 21.38 |
| P31382 | PMT2 | NLHTHPVAAPVSK | 2 | heavy | 689.8877 | y4 | 438.2802 | 21.38 |
| P31382 | PMT2 | NLHTHPVAAPVSK | 2 | heavy | 689.8877 | y2 | 242.159 | 21.38 |
| P31382 | PMT2 | NLHTHPVAAPVSK | 2 | heavy | 689.8877 | y4 | 219.6437 | 21.38 |
| P31382 | PMT2 | NLHTHPVAAPVSK | 2 | heavy | 689.8877 | b3 | 365.1932 | 21.38 |
| P31382 | PMT2 | NLHTHPVAAPVSK | 2 | heavy | 689.8877 | b5 | 603.2998 | 21.38 |
| P31382 | PMT2 | NLHTHPVAAPVSK | 2 | heavy | 689.8877 | b9 | 941.4952 | 21.38 |
| P36091 | DCW1 | YTGN[+203.079373]QTYVDWAEK | 2 | light | 889.3994 | y8 | 1011.478 | 44.41 |
| P36091 | DCW1 | YTGN[+203.079373]QTYVDWAEK | 2 | light | 889.3994 | y6 | 747.3672 | 44.41 |
| P36091 | DCW1 | YTGN[+203.079373]QTYVDWAEK | 2 | light | 889.3994 | y5 | 648.2988 | 44.41 |
| P36091 | DCW1 | YTGN[+203.079373]QTYVDWAEK | 2 | light | 889.3994 | y4 | 533.2718 | 44.41 |
| P36091 | DCW1 | YTGN[+203.079373]QTYVDWAEK | 2 | light | 889.3994 | y3 | 347.1925 | 44.41 |
| P36091 | DCW1 | YTGN[+203.079373]QTYVDWAEK | 2 | light | 889.3994 | b2 | 265.1183 | 44.41 |
| P36091 | DCW1 | YTGN[+203.079373]QTYVDWAEK | 2 | light | 889.3994 | b5 | 564.2413 | 44.41 |
| P36091 | DCW1 | YTGN[+203.079373]QTYVDWAEK | 2 | heavy | 893.4065 | y8 | 1019.492 | 44.41 |
| P36091 | DCW1 | YTGN[+203.079373]QTYVDWAEK | 2 | heavy | 893.4065 | y6 | 755.3814 | 44.41 |
| P36091 | DCW1 | YTGN[+203.079373]QTYVDWAEK | 2 | heavy | 893.4065 | y5 | 656.313 | 44.41 |
| P36091 | DCW1 | YTGN[+203.079373]QTYVDWAEK | 2 | heavy | 893.4065 | y4 | 541.286 | 44.41 |
| P36091 | DCW1 | YTGN[+203.079373]QTYVDWAEK | 2 | heavy | 893.4065 | y3 | 355.2067 | 44.41 |
| P36091 | DCW1 | YTGN[+203.079373]QTYVDWAEK | 2 | heavy | 893.4065 | b2 | 265.1183 | 44.41 |
| P36091 | DCW1 | YTGN[+203.079373]QTYVDWAEK | 2 | heavy | 893.4065 | b5 | 564.2413 | 44.41 |
| P36091 | DCW1 | NTVSNGALFHIAAR | 3 | light | 490.9319 | y6 | 714.4046 | 51.42 |
| P36091 | DCW1 | NTVSNGALFHIAAR | 3 | light | 490.9319 | y5 | 567.3362 | 51.42 |
| P36091 | DCW1 | NTVSNGALFHIAAR | 3 | light | 490.9319 | y4 | 430.2772 | 51.42 |
| P36091 | DCW1 | NTVSNGALFHIAAR | 3 | light | 490.9319 | y3 | 317.1932 | 51.42 |
| P36091 | DCW1 | NTVSNGALFHIAAR | 3 | light | 490.9319 | y12 | 628.3489 | 51.42 |
| P36091 | DCW1 | NTVSNGALFHIAAR | 3 | heavy | 492.9386 | y6 | 720.4247 | 51.42 |
| P36091 | DCW1 | NTVSNGALFHIAAR | 3 | heavy | 492.9386 | y5 | 573.3563 | 51.42 |
| P36091 | DCW1 | NTVSNGALFHIAAR | 3 | heavy | 492.9386 | y4 | 436.2974 | 51.42 |

| | | | | | | | | |
|---|---|---|---|---|---|---|---|---|
| P36091 | DCW1 | NTVSNGALFHIAAR | 3 | heavy | 492.9386 | y3 | 323.2133 | 51.42 |
| P36091 | DCW1 | NTVSNGALFHIAAR | 3 | heavy | 492.9386 | y12 | 631.359 | 51.42 |
| Q06689 | YL413 | ILNSAVN[+203.079373]MTTIT-PEQLK | 2 | light | 1038.548 | y11 | 1275.661 | 60.52 |
| Q06689 | YL413 | ILNSAVN[+203.079373]MTTIT-PEQLK | 2 | light | 1038.548 | y9 | 1030.578 | 60.52 |
| Q06689 | YL413 | ILNSAVN[+203.079373]MTTIT-PEQLK | 2 | light | 1038.548 | y8 | 929.5302 | 60.52 |
| Q06689 | YL413 | ILNSAVN[+203.079373]MTTIT-PEQLK | 2 | light | 1038.548 | y7 | 828.4825 | 60.52 |
| Q06689 | YL413 | ILNSAVN[+203.079373]MTTIT-PEQLK | 2 | light | 1038.548 | y6 | 715.3985 | 60.52 |
| Q06689 | YL413 | ILNSAVN[+203.079373]MTTIT-PEQLK | 2 | light | 1038.548 | y5 | 614.3508 | 60.52 |
| Q06689 | YL413 | ILNSAVN[+203.079373]MTTIT-PEQLK | 2 | light | 1038.548 | b5 | 499.2875 | 60.52 |
| Q06689 | YL413 | ILNSAVN[+203.079373]MTTIT-PEQLK | 2 | light | 1038.548 | b6 | 598.3559 | 60.52 |
| Q06689 | YL413 | ILNSAVN[+203.079373]MTTIT-PEQLK | 2 | heavy | 1042.555 | y11 | 1283.676 | 60.52 |
| Q06689 | YL413 | ILNSAVN[+203.079373]MTTIT-PEQLK | 2 | heavy | 1042.555 | y9 | 1038.592 | 60.52 |
| Q06689 | YL413 | ILNSAVN[+203.079373]MTTIT-PEQLK | 2 | heavy | 1042.555 | y8 | 937.5444 | 60.52 |
| Q06689 | YL413 | ILNSAVN[+203.079373]MTTIT-PEQLK | 2 | heavy | 1042.555 | y7 | 836.4967 | 60.52 |
| Q06689 | YL413 | ILNSAVN[+203.079373]MTTIT-PEQLK | 2 | heavy | 1042.555 | y6 | 723.4127 | 60.52 |
| Q06689 | YL413 | ILNSAVN[+203.079373]MTTIT-PEQLK | 2 | heavy | 1042.555 | y5 | 622.365 | 60.52 |
| Q06689 | YL413 | ILNSAVN[+203.079373]MTTIT-PEQLK | 2 | heavy | 1042.555 | b5 | 499.2875 | 60.52 |
| Q06689 | YL413 | ILNSAVN[+203.079373]MTTIT-PEQLK | 2 | heavy | 1042.555 | b6 | 598.3559 | 60.52 |
| Q06689 | YL413 | SHAVQNMDFR | 2 | light | 602.78 | y8 | 980.4618 | 28.23 |
| Q06689 | YL413 | SHAVQNMDFR | 2 | light | 602.78 | y7 | 909.4247 | 28.23 |
| Q06689 | YL413 | SHAVQNMDFR | 2 | light | 602.78 | y6 | 810.3563 | 28.23 |
| Q06689 | YL413 | SHAVQNMDFR | 2 | light | 602.78 | y2 | 322.1874 | 28.23 |
| Q06689 | YL413 | SHAVQNMDFR | 2 | light | 602.78 | b2 | 225.0982 | 28.23 |
| Q06689 | YL413 | SHAVQNMDFR | 2 | light | 602.78 | b3 | 296.1353 | 28.23 |
| Q06689 | YL413 | SHAVQNMDFR | 2 | light | 602.78 | b4 | 395.2037 | 28.23 |
| Q06689 | YL413 | SHAVQNMDFR | 2 | heavy | 605.7901 | y8 | 986.482 | 28.23 |
| Q06689 | YL413 | SHAVQNMDFR | 2 | heavy | 605.7901 | y7 | 915.4448 | 28.23 |
| Q06689 | YL413 | SHAVQNMDFR | 2 | heavy | 605.7901 | y6 | 816.3764 | 28.23 |
| Q06689 | YL413 | SHAVQNMDFR | 2 | heavy | 605.7901 | y2 | 328.2075 | 28.23 |
| Q06689 | YL413 | SHAVQNMDFR | 2 | heavy | 605.7901 | b2 | 225.0982 | 28.23 |
| Q06689 | YL413 | SHAVQNMDFR | 2 | heavy | 605.7901 | b3 | 296.1353 | 28.23 |
| Q06689 | YL413 | SHAVQNMDFR | 2 | heavy | 605.7901 | b4 | 395.2037 | 28.23 |

| | | | | | | | | |
|---|---|---|---|---|---|---|---|---|
| P46992 | YJR1 | N[+203.079373]SSSIGYYDL-PAIWLLNDHIAR | 3 | light | 907.7888 | y9 | 1137.616 | 88.27 |
| P46992 | YJR1 | N[+203.079373]SSSIGYYDL-PAIWLLNDHIAR | 3 | light | 907.7888 | y8 | 951.537 | 88.27 |
| P46992 | YJR1 | N[+203.079373]SSSIGYYDL-PAIWLLNDHIAR | 3 | light | 907.7888 | y7 | 838.453 | 88.27 |
| P46992 | YJR1 | N[+203.079373]SSSIGYYDL-PAIWLLNDHIAR | 3 | light | 907.7888 | y4 | 496.299 | 88.27 |
| P46992 | YJR1 | N[+203.079373]SSSIGYYDL-PAIWLLNDHIAR | 3 | light | 907.7888 | b2 | 202.0822 | 88.27 |
| P46992 | YJR1 | N[+203.079373]SSSIGYYDL-PAIWLLNDHIAR | 3 | heavy | 909.7955 | y9 | 1143.636 | 88.27 |
| P46992 | YJR1 | N[+203.079373]SSSIGYYDL-PAIWLLNDHIAR | 3 | heavy | 909.7955 | y8 | 957.5572 | 88.27 |
| P46992 | YJR1 | N[+203.079373]SSSIGYYDL-PAIWLLNDHIAR | 3 | heavy | 909.7955 | y7 | 844.4731 | 88.27 |
| P46992 | YJR1 | N[+203.079373]SSSIGYYDL-PAIWLLNDHIAR | 3 | heavy | 909.7955 | y4 | 502.3192 | 88.27 |
| P46992 | YJR1 | N[+203.079373]SSSIGYYDL-PAIWLLNDHIAR | 3 | heavy | 909.7955 | b2 | 202.0822 | 88.27 |
| P46992 | YJR1 | SGIPAYYGYGGTTK | 2 | light | 717.8486 | y11 | 1177.552 | 44.44 |
| P46992 | YJR1 | SGIPAYYGYGGTTK | 2 | light | 717.8486 | y10 | 1080.5 | 44.44 |
| P46992 | YJR1 | SGIPAYYGYGGTTK | 2 | light | 717.8486 | y9 | 1009.463 | 44.44 |
| P46992 | YJR1 | SGIPAYYGYGGTTK | 2 | light | 717.8486 | y8 | 846.3992 | 44.44 |
| P46992 | YJR1 | SGIPAYYGYGGTTK | 2 | light | 717.8486 | y7 | 683.3359 | 44.44 |
| P46992 | YJR1 | SGIPAYYGYGGTTK | 2 | light | 717.8486 | y5 | 463.2511 | 44.44 |
| P46992 | YJR1 | SGIPAYYGYGGTTK | 2 | light | 717.8486 | y11 | 589.2798 | 44.44 |
| P46992 | YJR1 | SGIPAYYGYGGTTK | 2 | heavy | 721.8557 | y11 | 1185.567 | 44.44 |
| P46992 | YJR1 | SGIPAYYGYGGTTK | 2 | heavy | 721.8557 | y10 | 1088.514 | 44.44 |
| P46992 | YJR1 | SGIPAYYGYGGTTK | 2 | heavy | 721.8557 | y9 | 1017.477 | 44.44 |
| P46992 | YJR1 | SGIPAYYGYGGTTK | 2 | heavy | 721.8557 | y8 | 854.4134 | 44.44 |
| P46992 | YJR1 | SGIPAYYGYGGTTK | 2 | heavy | 721.8557 | y7 | 691.3501 | 44.44 |
| P46992 | YJR1 | SGIPAYYGYGGTTK | 2 | heavy | 721.8557 | y5 | 471.2653 | 44.44 |
| P46992 | YJR1 | SGIPAYYGYGGTTK | 2 | heavy | 721.8557 | y11 | 593.2869 | 44.44 |
| P12684 | HMDH2 | IPTELVSEN[+203.079373]GTK | 2 | light | 745.8829 | y10 | 1280.622 | 36.27 |
| P12684 | HMDH2 | IPTELVSEN[+203.079373]GTK | 2 | light | 745.8829 | y9 | 1179.574 | 36.27 |
| P12684 | HMDH2 | IPTELVSEN[+203.079373]GTK | 2 | light | 745.8829 | y8 | 1050.531 | 36.27 |
| P12684 | HMDH2 | IPTELVSEN[+203.079373]GTK | 2 | light | 745.8829 | y7 | 937.4473 | 36.27 |
| P12684 | HMDH2 | IPTELVSEN[+203.079373]GTK | 2 | light | 745.8829 | y6 | 838.3789 | 36.27 |
| P12684 | HMDH2 | IPTELVSEN[+203.079373]GTK | 2 | light | 745.8829 | y5 | 751.3468 | 36.27 |
| P12684 | HMDH2 | IPTELVSEN[+203.079373]GTK | 2 | heavy | 749.89 | y10 | 1288.636 | 36.27 |
| P12684 | HMDH2 | IPTELVSEN[+203.079373]GTK | 2 | heavy | 749.89 | y9 | 1187.588 | 36.27 |

| | | | | | | | | |
|---|---|---|---|---|---|---|---|---|
| P12684 | HMDH2 | IPTELVSEN[+203.079373]GTK | 2 | heavy | 749.89 | y8 | 1058.546 | 36.27 |
| P12684 | HMDH2 | IPTELVSEN[+203.079373]GTK | 2 | heavy | 749.89 | y7 | 945.4615 | 36.27 |
| P12684 | HMDH2 | IPTELVSEN[+203.079373]GTK | 2 | heavy | 749.89 | y6 | 846.3931 | 36.27 |
| P12684 | HMDH2 | IPTELVSEN[+203.079373]GTK | 2 | heavy | 749.89 | y5 | 759.361 | 36.27 |
| P12684 | HMDH2 | ALSTLAESPILVSEK | 2 | light | 779.4403 | y11 | 1185.673 | 60.36 |
| P12684 | HMDH2 | ALSTLAESPILVSEK | 2 | light | 779.4403 | y10 | 1072.588 | 60.36 |
| P12684 | HMDH2 | ALSTLAESPILVSEK | 2 | light | 779.4403 | y9 | 1001.551 | 60.36 |
| P12684 | HMDH2 | ALSTLAESPILVSEK | 2 | light | 779.4403 | y8 | 872.5088 | 60.36 |
| P12684 | HMDH2 | ALSTLAESPILVSEK | 2 | light | 779.4403 | y7 | 785.4767 | 60.36 |
| P12684 | HMDH2 | ALSTLAESPILVSEK | 2 | light | 779.4403 | y5 | 575.3399 | 60.36 |
| P12684 | HMDH2 | ALSTLAESPILVSEK | 2 | light | 779.4403 | y4 | 462.2558 | 60.36 |
| P12684 | HMDH2 | ALSTLAESPILVSEK | 2 | light | 779.4403 | y3 | 363.1874 | 60.36 |
| P12684 | HMDH2 | ALSTLAESPILVSEK | 2 | heavy | 783.4474 | y11 | 1193.687 | 60.36 |
| P12684 | HMDH2 | ALSTLAESPILVSEK | 2 | heavy | 783.4474 | y10 | 1080.603 | 60.36 |
| P12684 | HMDH2 | ALSTLAESPILVSEK | 2 | heavy | 783.4474 | y9 | 1009.566 | 60.36 |
| P12684 | HMDH2 | ALSTLAESPILVSEK | 2 | heavy | 783.4474 | y8 | 880.523 | 60.36 |
| P12684 | HMDH2 | ALSTLAESPILVSEK | 2 | heavy | 783.4474 | y7 | 793.4909 | 60.36 |
| P12684 | HMDH2 | ALSTLAESPILVSEK | 2 | heavy | 783.4474 | y5 | 583.3541 | 60.36 |
| P12684 | HMDH2 | ALSTLAESPILVSEK | 2 | heavy | 783.4474 | y4 | 470.27 | 60.36 |
| P12684 | HMDH2 | ALSTLAESPILVSEK | 2 | heavy | 783.4474 | y3 | 371.2016 | 60.36 |
| P53379 | MKC7 | STAYSLFAN[+203.079373]DSDSK | 2 | light | 854.881 | y9 | 1199.543 | 44.96 |
| P53379 | MKC7 | STAYSLFAN[+203.079373]DSDSK | 2 | light | 854.881 | y8 | 1086.459 | 44.96 |
| P53379 | MKC7 | STAYSLFAN[+203.079373]DSDSK | 2 | light | 854.881 | y7 | 939.3902 | 44.96 |
| P53379 | MKC7 | STAYSLFAN[+203.079373]DSDSK | 2 | light | 854.881 | y5 | 551.2307 | 44.96 |
| P53379 | MKC7 | STAYSLFAN[+203.079373]DSDSK | 2 | light | 854.881 | y4 | 436.2038 | 44.96 |
| P53379 | MKC7 | STAYSLFAN[+203.079373]DSDSK | 2 | light | 854.881 | b3 | 260.1241 | 44.96 |
| P53379 | MKC7 | STAYSLFAN[+203.079373]DSDSK | 2 | heavy | 858.8881 | y9 | 1207.557 | 44.96 |
| P53379 | MKC7 | STAYSLFAN[+203.079373]DSDSK | 2 | heavy | 858.8881 | y8 | 1094.473 | 44.96 |
| P53379 | MKC7 | STAYSLFAN[+203.079373]DSDSK | 2 | heavy | 858.8881 | y7 | 947.4044 | 44.96 |
| P53379 | MKC7 | STAYSLFAN[+203.079373]DSDSK | 2 | heavy | 858.8881 | y5 | 559.2449 | 44.96 |
| P53379 | MKC7 | STAYSLFAN[+203.079373]DSDSK | 2 | heavy | 858.8881 | y4 | 444.218 | 44.96 |
| P53379 | MKC7 | STAYSLFAN[+203.079373]DSDSK | 2 | heavy | 858.8881 | b3 | 260.1241 | 44.96 |

| | | | | | | | | |
|---|---|---|---|---|---|---|---|---|
| P53379 | MKC7 | HGTILFGAVDHGK | 3 | light | 451.2421 | y7 | 683.3471 | 36.85 |
| P53379 | MKC7 | HGTILFGAVDHGK | 3 | light | 451.2421 | y5 | 555.2885 | 36.85 |
| P53379 | MKC7 | HGTILFGAVDHGK | 3 | light | 451.2421 | y4 | 456.2201 | 36.85 |
| P53379 | MKC7 | HGTILFGAVDHGK | 3 | light | 451.2421 | y3 | 341.1932 | 36.85 |
| P53379 | MKC7 | HGTILFGAVDHGK | 3 | light | 451.2421 | b3 | 296.1353 | 36.85 |
| P53379 | MKC7 | HGTILFGAVDHGK | 3 | light | 451.2421 | b4 | 409.2194 | 36.85 |
| P53379 | MKC7 | HGTILFGAVDHGK | 3 | light | 451.2421 | b5 | 522.3035 | 36.85 |
| P53379 | MKC7 | HGTILFGAVDHGK | 3 | heavy | 453.9135 | y7 | 691.3613 | 36.85 |
| P53379 | MKC7 | HGTILFGAVDHGK | 3 | heavy | 453.9135 | y5 | 563.3027 | 36.85 |
| P53379 | MKC7 | HGTILFGAVDHGK | 3 | heavy | 453.9135 | y4 | 464.2343 | 36.85 |
| P53379 | MKC7 | HGTILFGAVDHGK | 3 | heavy | 453.9135 | y3 | 349.2074 | 36.85 |
| P53379 | MKC7 | HGTILFGAVDHGK | 3 | heavy | 453.9135 | b3 | 296.1353 | 36.85 |
| P53379 | MKC7 | HGTILFGAVDHGK | 3 | heavy | 453.9135 | b4 | 409.2194 | 36.85 |
| P53379 | MKC7 | HGTILFGAVDHGK | 3 | heavy | 453.9135 | b5 | 522.3035 | 36.85 |
| P53379 | MKC7 | YAGDLYTIPIINTLQHR | 2 | light | 994.5336 | y11 | 1305.764 | 74.83 |
| P53379 | MKC7 | YAGDLYTIPIINTLQHR | 2 | light | 994.5336 | y10 | 1204.716 | 74.83 |
| P53379 | MKC7 | YAGDLYTIPIINTLQHR | 2 | light | 994.5336 | y9 | 1091.632 | 74.83 |
| P53379 | MKC7 | YAGDLYTIPIINTLQHR | 2 | light | 994.5336 | b4 | 407.1561 | 74.83 |
| P53379 | MKC7 | YAGDLYTIPIINTLQHR | 2 | heavy | 997.5437 | y11 | 1311.784 | 74.83 |
| P53379 | MKC7 | YAGDLYTIPIINTLQHR | 2 | heavy | 997.5437 | y10 | 1210.736 | 74.83 |
| P53379 | MKC7 | YAGDLYTIPIINTLQHR | 2 | heavy | 997.5437 | y9 | 1097.652 | 74.83 |
| P53379 | MKC7 | YAGDLYTIPIINTLQHR | 2 | heavy | 997.5437 | b4 | 407.1561 | 74.83 |
| Q12465 | RAX2 | EIGPETSSHGLVYYSN[+203.079373]NTYIQLEDASDDTR | 3 | light | 1193.208 | y11 | 1262.586 | 58.75 |
| Q12465 | RAX2 | EIGPETSSHGLVYYSN[+203.079373]NTYIQLEDASDDTR | 3 | light | 1193.208 | y10 | 1149.502 | 58.75 |
| Q12465 | RAX2 | EIGPETSSHGLVYYSN[+203.079373]NTYIQLEDASDDTR | 3 | light | 1193.208 | y9 | 1021.443 | 58.75 |
| Q12465 | RAX2 | EIGPETSSHGLVYYSN[+203.079373]NTYIQLEDASDDTR | 3 | light | 1193.208 | y8 | 908.3592 | 58.75 |
| Q12465 | RAX2 | EIGPETSSHGLVYYSN[+203.079373]NTYIQLEDASDDTR | 3 | light | 1193.208 | y7 | 779.3166 | 58.75 |
| Q12465 | RAX2 | EIGPETSSHGLVYYSN[+203.079373]NTYIQLEDASDDTR | 3 | light | 1193.208 | y6 | 664.2897 | 58.75 |
| Q12465 | RAX2 | EIGPETSSHGLVYYSN[+203.079373]NTYIQLEDASDDTR | 3 | light | 1193.208 | y5 | 593.2525 | 58.75 |
| Q12465 | RAX2 | EIGPETSSHGLVYYSN[+203.079373]NTYIQLEDASDDTR | 3 | light | 1193.208 | y4 | 506.2205 | 58.75 |
| Q12465 | RAX2 | EIGPETSSHGLVYYSN[+203.079373]NTYIQLEDASDDTR | 3 | heavy | 1195.215 | y11 | 1268.606 | 58.75 |
| Q12465 | RAX2 | EIGPETSSHGLVYYSN[+203.079373]NTYIQLEDASDDTR | 3 | heavy | 1195.215 | y10 | 1155.522 | 58.75 |

| | | | | | | | | |
|---|---|---|---|---|---|---|---|---|
| Q12465 | RAX2 | EIGPETSSHGLVYYSN[+203.079373]NTYIQLEDASDDTR | 3 | heavy | 1195.215 | y9 | 1027.463 | 58.75 |
| Q12465 | RAX2 | EIGPETSSHGLVYYSN[+203.079373]NTYIQLEDASDDTR | 3 | heavy | 1195.215 | y8 | 914.3793 | 58.75 |
| Q12465 | RAX2 | EIGPETSSHGLVYYSN[+203.079373]NTYIQLEDASDDTR | 3 | heavy | 1195.215 | y7 | 785.3367 | 58.75 |
| Q12465 | RAX2 | EIGPETSSHGLVYYSN[+203.079373]NTYIQLEDASDDTR | 3 | heavy | 1195.215 | y6 | 670.3098 | 58.75 |
| Q12465 | RAX2 | EIGPETSSHGLVYYSN[+203.079373]NTYIQLEDASDDTR | 3 | heavy | 1195.215 | y5 | 599.2727 | 58.75 |
| Q12465 | RAX2 | EIGPETSSHGLVYYSN[+203.079373]NTYIQLEDASDDTR | 3 | heavy | 1195.215 | y4 | 512.2406 | 58.75 |
| Q12465 | RAX2 | N[+203.079373]SSLYADIYDNK | 2 | light | 803.3676 | y8 | 1001.457 | 44.95 |
| Q12465 | RAX2 | N[+203.079373]SSLYADIYDNK | 2 | light | 803.3676 | y6 | 767.357 | 44.95 |
| Q12465 | RAX2 | N[+203.079373]SSLYADIYDNK | 2 | light | 803.3676 | y5 | 652.3301 | 44.95 |
| Q12465 | RAX2 | N[+203.079373]SSLYADIYDNK | 2 | light | 803.3676 | y4 | 539.246 | 44.95 |
| Q12465 | RAX2 | N[+203.079373]SSLYADIYDNK | 2 | heavy | 807.3747 | y8 | 1009.472 | 44.95 |
| Q12465 | RAX2 | N[+203.079373]SSLYADIYDNK | 2 | heavy | 807.3747 | y6 | 775.3712 | 44.95 |
| Q12465 | RAX2 | N[+203.079373]SSLYADIYDNK | 2 | heavy | 807.3747 | y5 | 660.3443 | 44.95 |
| Q12465 | RAX2 | N[+203.079373]SSLYADIYDNK | 2 | heavy | 807.3747 | y4 | 547.2602 | 44.95 |
| Q12465 | RAX2 | N[+203.079373]QTIQGDVHGITK | 2 | light | 807.4101 | y11 | 1168.632 | 27.89 |
| Q12465 | RAX2 | N[+203.079373]QTIQGDVHGITK | 2 | light | 807.4101 | y10 | 1067.584 | 27.89 |
| Q12465 | RAX2 | N[+203.079373]QTIQGDVHGITK | 2 | light | 807.4101 | y9 | 954.5003 | 27.89 |
| Q12465 | RAX2 | N[+203.079373]QTIQGDVHGITK | 2 | light | 807.4101 | y8 | 826.4417 | 27.89 |
| Q12465 | RAX2 | N[+203.079373]QTIQGDVHGITK | 2 | light | 807.4101 | y6 | 654.3933 | 27.89 |
| Q12465 | RAX2 | N[+203.079373]QTIQGDVHGITK | 2 | light | 807.4101 | y5 | 555.3249 | 27.89 |
| Q12465 | RAX2 | N[+203.079373]QTIQGDVHGITK | 2 | light | 807.4101 | y4 | 418.266 | 27.89 |
| Q12465 | RAX2 | N[+203.079373]QTIQGDVHGITK | 2 | light | 807.4101 | b2 | 243.1088 | 27.89 |
| Q12465 | RAX2 | N[+203.079373]QTIQGDVHGITK | 2 | light | 807.4101 | b3 | 344.1565 | 27.89 |
| Q12465 | RAX2 | N[+203.079373]QTIQGDVHGITK | 2 | heavy | 811.4172 | y11 | 1176.646 | 27.89 |
| Q12465 | RAX2 | N[+203.079373]QTIQGDVHGITK | 2 | heavy | 811.4172 | y10 | 1075.599 | 27.89 |
| Q12465 | RAX2 | N[+203.079373]QTIQGDVHGITK | 2 | heavy | 811.4172 | y9 | 962.5145 | 27.89 |
| Q12465 | RAX2 | N[+203.079373]QTIQGDVHGITK | 2 | heavy | 811.4172 | y8 | 834.4559 | 27.89 |
| Q12465 | RAX2 | N[+203.079373]QTIQGDVHGITK | 2 | heavy | 811.4172 | y6 | 662.4075 | 27.89 |
| Q12465 | RAX2 | N[+203.079373]QTIQGDVHGITK | 2 | heavy | 811.4172 | y5 | 563.3391 | 27.89 |
| Q12465 | RAX2 | N[+203.079373]QTIQGDVHGITK | 2 | heavy | 811.4172 | y4 | 426.2802 | 27.89 |
| Q12465 | RAX2 | N[+203.079373]QTIQGDVHGITK | 2 | heavy | 811.4172 | b2 | 243.1088 | 27.89 |
| Q12465 | RAX2 | N[+203.079373]QTIQGDVHGITK | 2 | heavy | 811.4172 | b3 | 344.1565 | 27.89 |

| | | | | | | | | |
|---|---|---|---|---|---|---|---|---|
| Q12465 | RAX2 | IPVLLDSGTTISYMPTELVK | 2 | light | 1089.092 | y9 | 1067.544 | 81.96 |
| Q12465 | RAX2 | IPVLLDSGTTISYMPTELVK | 2 | light | 1089.092 | y8 | 980.5121 | 81.96 |
| Q12465 | RAX2 | IPVLLDSGTTISYMPTELVK | 2 | light | 1089.092 | y7 | 817.4488 | 81.96 |
| Q12465 | RAX2 | IPVLLDSGTTISYMPTELVK | 2 | light | 1089.092 | y6 | 686.4083 | 81.96 |
| Q12465 | RAX2 | IPVLLDSGTTISYMPTELVK | 2 | light | 1089.092 | y5 | 589.3556 | 81.96 |
| Q12465 | RAX2 | IPVLLDSGTTISYMPTELVK | 2 | light | 1089.092 | y3 | 359.2653 | 81.96 |
| Q12465 | RAX2 | IPVLLDSGTTISYMPTELVK | 2 | heavy | 1093.099 | y9 | 1075.558 | 81.96 |
| Q12465 | RAX2 | IPVLLDSGTTISYMPTELVK | 2 | heavy | 1093.099 | y8 | 988.5263 | 81.96 |
| Q12465 | RAX2 | IPVLLDSGTTISYMPTELVK | 2 | heavy | 1093.099 | y7 | 825.463 | 81.96 |
| Q12465 | RAX2 | IPVLLDSGTTISYMPTELVK | 2 | heavy | 1093.099 | y6 | 694.4225 | 81.96 |
| Q12465 | RAX2 | IPVLLDSGTTISYMPTELVK | 2 | heavy | 1093.099 | y5 | 597.3698 | 81.96 |
| Q12465 | RAX2 | IPVLLDSGTTISYMPTELVK | 2 | heavy | 1093.099 | y3 | 367.2795 | 81.96 |
| Q12465 | RAX2 | YVPDQNEPIPR | 2 | light | 664.3357 | y9 | 1065.532 | 34.96 |
| Q12465 | RAX2 | YVPDQNEPIPR | 2 | light | 664.3357 | y8 | 968.4796 | 34.96 |
| Q12465 | RAX2 | YVPDQNEPIPR | 2 | light | 664.3357 | y7 | 853.4526 | 34.96 |
| Q12465 | RAX2 | YVPDQNEPIPR | 2 | light | 664.3357 | y6 | 725.3941 | 34.96 |
| Q12465 | RAX2 | YVPDQNEPIPR | 2 | light | 664.3357 | y5 | 611.3511 | 34.96 |
| Q12465 | RAX2 | YVPDQNEPIPR | 2 | light | 664.3357 | y4 | 482.3085 | 34.96 |
| Q12465 | RAX2 | YVPDQNEPIPR | 2 | heavy | 667.3457 | y9 | 1071.552 | 34.96 |
| Q12465 | RAX2 | YVPDQNEPIPR | 2 | heavy | 667.3457 | y8 | 974.4997 | 34.96 |
| Q12465 | RAX2 | YVPDQNEPIPR | 2 | heavy | 667.3457 | y7 | 859.4728 | 34.96 |
| Q12465 | RAX2 | YVPDQNEPIPR | 2 | heavy | 667.3457 | y6 | 731.4142 | 34.96 |
| Q12465 | RAX2 | YVPDQNEPIPR | 2 | heavy | 667.3457 | y5 | 617.3713 | 34.96 |
| Q12465 | RAX2 | YVPDQNEPIPR | 2 | heavy | 667.3457 | y4 | 488.3287 | 34.96 |
| P27825 | CAL-XorC-NE1 | N[+203.079373]VTEAQIIGN[+203.079373]K | 2 | light | 796.9043 | y10 | 1275.679 | 32.58 |
| P27825 | CAL-XorC-NE1 | N[+203.079373]VTEAQIIGN[+203.079373]K | 2 | light | 796.9043 | y9 | 1176.611 | 32.58 |
| P27825 | CAL-XorC-NE1 | N[+203.079373]VTEAQIIGN[+203.079373]K | 2 | light | 796.9043 | y8 | 1075.563 | 32.58 |
| P27825 | CAL-XorC-NE1 | N[+203.079373]VTEAQIIGN[+203.079373]K | 2 | light | 796.9043 | y7 | 946.5204 | 32.58 |
| P27825 | CAL-XorC-NE1 | N[+203.079373]VTEAQIIGN[+203.079373]K | 2 | light | 796.9043 | y6 | 875.4833 | 32.58 |

| | | | | | | | | |
|---|---|---|---|---|---|---|---|---|
| P27825 | CAL-XorC-NE1 | N[+203.079373]VTEAQIIGN[+203.079373]K | 2 | light | 796.9043 | y5 | 747.4247 | 32.58 |
| P27825 | CAL-XorC-NE1 | N[+203.079373]VTEAQIIGN[+203.079373]K | 2 | light | 796.9043 | y4 | 634.3406 | 32.58 |
| P27825 | CAL-XorC-NE1 | N[+203.079373]VTEAQIIGN[+203.079373]K | 2 | light | 796.9043 | y3 | 521.2566 | 32.58 |
| P27825 | CAL-XorC-NE1 | N[+203.079373]VTEAQIIGN[+203.079373]K | 2 | heavy | 800.9114 | y10 | 1283.693 | 32.58 |
| P27825 | CAL-XorC-NE1 | N[+203.079373]VTEAQIIGN[+203.079373]K | 2 | heavy | 800.9114 | y9 | 1184.625 | 32.58 |
| P27825 | CAL-XorC-NE1 | N[+203.079373]VTEAQIIGN[+203.079373]K | 2 | heavy | 800.9114 | y8 | 1083.577 | 32.58 |
| P27825 | CAL-XorC-NE1 | N[+203.079373]VTEAQIIGN[+203.079373]K | 2 | heavy | 800.9114 | y7 | 954.5346 | 32.58 |
| P27825 | CAL-XorC-NE1 | N[+203.079373]VTEAQIIGN[+203.079373]K | 2 | heavy | 800.9114 | y6 | 883.4975 | 32.58 |
| P27825 | CAL-XorC-NE1 | N[+203.079373]VTEAQIIGN[+203.079373]K | 2 | heavy | 800.9114 | y5 | 755.4389 | 32.58 |
| P27825 | CAL-XorC-NE1 | N[+203.079373]VTEAQIIGN[+203.079373]K | 2 | heavy | 800.9114 | y4 | 642.3548 | 32.58 |
| P27825 | CAL-XorC-NE1 | N[+203.079373]VTEAQIIGN[+203.079373]K | 2 | heavy | 800.9114 | y3 | 529.2708 | 32.58 |
| P27825 | CAL-XorC-NE1 | LDNSLTC[+57.021464]GGAFIK | 2 | light | 698.3505 | y10 | 1053.54 | 50.6 |
| P27825 | CAL-XorC-NE1 | LDNSLTC[+57.021464]GGAFIK | 2 | light | 698.3505 | y9 | 966.5077 | 50.6 |
| P27825 | CAL-XorC-NE1 | LDNSLTC[+57.021464]GGAFIK | 2 | light | 698.3505 | y8 | 853.4237 | 50.6 |
| P27825 | CAL-XorC-NE1 | LDNSLTC[+57.021464]GGAFIK | 2 | light | 698.3505 | y6 | 592.3453 | 50.6 |
| P27825 | CAL-XorC-NE1 | LDNSLTC[+57.021464]GGAFIK | 2 | light | 698.3505 | y5 | 535.3239 | 50.6 |
| P27825 | CAL-XorC-NE1 | LDNSLTC[+57.021464]GGAFIK | 2 | light | 698.3505 | y3 | 407.2653 | 50.6 |
| P27825 | CAL-XorC-NE1 | LDNSLTC[+57.021464]GGAFIK | 2 | heavy | 702.3576 | y10 | 1061.554 | 50.6 |
| P27825 | CAL-XorC-NE1 | LDNSLTC[+57.021464]GGAFIK | 2 | heavy | 702.3576 | y9 | 974.5219 | 50.6 |
| P27825 | CAL-XorC-NE1 | LDNSLTC[+57.021464]GGAFIK | 2 | heavy | 702.3576 | y8 | 861.4378 | 50.6 |
| P27825 | CAL-XorC-NE1 | LDNSLTC[+57.021464]GGAFIK | 2 | heavy | 702.3576 | y6 | 600.3595 | 50.6 |

| | | | | | | | | |
|---|---|---|---|---|---|---|---|---|
| P27825 | CAL-XorC-NE1 | LDNSLTC[+57.021464]GGAFIK | 2 | heavy | 702.3576 | y5 | 543.3381 | 50.6 |
| P27825 | CAL-XorC-NE1 | LDNSLTC[+57.021464]GGAFIK | 2 | heavy | 702.3576 | y3 | 415.2795 | 50.6 |
| P40557 | EPS1 | VALVLPN[+203.079373]K | 2 | light | 528.8186 | y7 | 957.5615 | 42.29 |
| P40557 | EPS1 | VALVLPN[+203.079373]K | 2 | light | 528.8186 | y7 | 754.4822 | 42.29 |
| P40557 | EPS1 | VALVLPN[+203.079373]K | 2 | light | 528.8186 | y6 | 886.5244 | 42.29 |
| P40557 | EPS1 | VALVLPN[+203.079373]K | 2 | light | 528.8186 | y6 | 683.445 | 42.29 |
| P40557 | EPS1 | VALVLPN[+203.079373]K | 2 | light | 528.8186 | y5 | 570.361 | 42.29 |
| P40557 | EPS1 | VALVLPN[+203.079373]K | 2 | light | 528.8186 | y4 | 471.2926 | 42.29 |
| P40557 | EPS1 | VALVLPN[+203.079373]K | 2 | light | 528.8186 | y3 | 358.2085 | 42.29 |
| P40557 | EPS1 | VALVLPN[+203.079373]K | 2 | light | 528.8186 | b3 | 284.1969 | 42.29 |
| P40557 | EPS1 | VALVLPN[+203.079373]K | 2 | heavy | 532.8257 | y7 | 965.5757 | 42.29 |
| P40557 | EPS1 | VALVLPN[+203.079373]K | 2 | heavy | 532.8257 | y7 | 762.4964 | 42.29 |
| P40557 | EPS1 | VALVLPN[+203.079373]K | 2 | heavy | 532.8257 | y6 | 894.5386 | 42.29 |
| P40557 | EPS1 | VALVLPN[+203.079373]K | 2 | heavy | 532.8257 | y6 | 691.4592 | 42.29 |
| P40557 | EPS1 | VALVLPN[+203.079373]K | 2 | heavy | 532.8257 | y5 | 578.3752 | 42.29 |
| P40557 | EPS1 | VALVLPN[+203.079373]K | 2 | heavy | 532.8257 | y4 | 479.3068 | 42.29 |
| P40557 | EPS1 | VALVLPN[+203.079373]K | 2 | heavy | 532.8257 | y3 | 366.2227 | 42.29 |
| P40557 | EPS1 | VALVLPN[+203.079373]K | 2 | heavy | 532.8257 | b3 | 284.1969 | 42.29 |
| P40557 | EPS1 | FPN[+203.079373]ITEGELEK | 2 | light | 740.3643 | y10 | 1129.574 | 49.21 |
| P40557 | EPS1 | FPN[+203.079373]ITEGELEK | 2 | light | 740.3643 | y9 | 1235.6 | 49.21 |
| P40557 | EPS1 | FPN[+203.079373]ITEGELEK | 2 | light | 740.3643 | y9 | 1032.521 | 49.21 |
| P40557 | EPS1 | FPN[+203.079373]ITEGELEK | 2 | light | 740.3643 | y8 | 918.4779 | 49.21 |
| P40557 | EPS1 | FPN[+203.079373]ITEGELEK | 2 | light | 740.3643 | y7 | 805.3938 | 49.21 |
| P40557 | EPS1 | FPN[+203.079373]ITEGELEK | 2 | light | 740.3643 | y6 | 704.3461 | 49.21 |
| P40557 | EPS1 | FPN[+203.079373]ITEGELEK | 2 | light | 740.3643 | y5 | 575.3035 | 49.21 |
| P40557 | EPS1 | FPN[+203.079373]ITEGELEK | 2 | light | 740.3643 | y4 | 518.2821 | 49.21 |
| P40557 | EPS1 | FPN[+203.079373]ITEGELEK | 2 | light | 740.3643 | y3 | 389.2395 | 49.21 |
| P40557 | EPS1 | FPN[+203.079373]ITEGELEK | 2 | heavy | 744.3714 | y10 | 1137.588 | 49.21 |
| P40557 | EPS1 | FPN[+203.079373]ITEGELEK | 2 | heavy | 744.3714 | y9 | 1243.614 | 49.21 |
| P40557 | EPS1 | FPN[+203.079373]ITEGELEK | 2 | heavy | 744.3714 | y9 | 1040.535 | 49.21 |
| P40557 | EPS1 | FPN[+203.079373]ITEGELEK | 2 | heavy | 744.3714 | y8 | 926.4921 | 49.21 |

| | | | | | | | | |
|---|---|---|---|---|---|---|---|---|
| P40557 | EPS1 | FPN[+203.079373]ITEGELEK | 2 | heavy | 744.3714 | y7 | 813.408 | 49.21 |
| P40557 | EPS1 | FPN[+203.079373]ITEGELEK | 2 | heavy | 744.3714 | y6 | 712.3603 | 49.21 |
| P40557 | EPS1 | FPN[+203.079373]ITEGELEK | 2 | heavy | 744.3714 | y5 | 583.3177 | 49.21 |
| P40557 | EPS1 | FPN[+203.079373]ITEGELEK | 2 | heavy | 744.3714 | y4 | 526.2963 | 49.21 |
| P40557 | EPS1 | FPN[+203.079373]ITEGELEK | 2 | heavy | 744.3714 | y3 | 397.2537 | 49.21 |
| P40557 | EPS1 | EAFVSLNIPSK | 2 | light | 602.8322 | y9 | 1004.578 | 56.63 |
| P40557 | EPS1 | EAFVSLNIPSK | 2 | light | 602.8322 | y8 | 857.5091 | 56.63 |
| P40557 | EPS1 | EAFVSLNIPSK | 2 | light | 602.8322 | y7 | 758.4407 | 56.63 |
| P40557 | EPS1 | EAFVSLNIPSK | 2 | light | 602.8322 | y6 | 671.4087 | 56.63 |
| P40557 | EPS1 | EAFVSLNIPSK | 2 | light | 602.8322 | y5 | 558.3246 | 56.63 |
| P40557 | EPS1 | EAFVSLNIPSK | 2 | light | 602.8322 | y4 | 444.2817 | 56.63 |
| P40557 | EPS1 | EAFVSLNIPSK | 2 | light | 602.8322 | y3 | 331.1976 | 56.63 |
| P40557 | EPS1 | EAFVSLNIPSK | 2 | heavy | 606.8393 | y9 | 1012.592 | 56.63 |
| P40557 | EPS1 | EAFVSLNIPSK | 2 | heavy | 606.8393 | y8 | 865.5233 | 56.63 |
| P40557 | EPS1 | EAFVSLNIPSK | 2 | heavy | 606.8393 | y7 | 766.4549 | 56.63 |
| P40557 | EPS1 | EAFVSLNIPSK | 2 | heavy | 606.8393 | y6 | 679.4229 | 56.63 |
| P40557 | EPS1 | EAFVSLNIPSK | 2 | heavy | 606.8393 | y5 | 566.3388 | 56.63 |
| P40557 | EPS1 | EAFVSLNIPSK | 2 | heavy | 606.8393 | y4 | 452.2959 | 56.63 |
| P40557 | EPS1 | EAFVSLNIPSK | 2 | heavy | 606.8393 | y3 | 339.2118 | 56.63 |
| P40557 | EPS1 | NIDAIMDWVK | 2 | light | 602.8052 | y8 | 977.4761 | 37.01 |
| P40557 | EPS1 | NIDAIMDWVK | 2 | light | 602.8052 | y7 | 862.4491 | 37.01 |
| P40557 | EPS1 | NIDAIMDWVK | 2 | light | 602.8052 | y4 | 547.2875 | 37.01 |
| P40557 | EPS1 | NIDAIMDWVK | 2 | light | 602.8052 | y3 | 432.2605 | 37.01 |
| P40557 | EPS1 | NIDAIMDWVK | 2 | heavy | 606.8123 | y8 | 985.4903 | 37.01 |
| P40557 | EPS1 | NIDAIMDWVK | 2 | heavy | 606.8123 | y7 | 870.4633 | 37.01 |
| P40557 | EPS1 | NIDAIMDWVK | 2 | heavy | 606.8123 | y4 | 555.3017 | 37.01 |
| P40557 | EPS1 | NIDAIMDWVK | 2 | heavy | 606.8123 | y3 | 440.2747 | 37.01 |
| P52911 | EXG2 | FASYYAN[+203.079373]DTITVK | 2 | light | 848.4092 | y10 | 1390.674 | 46.19 |
| P52911 | EXG2 | FASYYAN[+203.079373]DTITVK | 2 | light | 848.4092 | y10 | 1187.594 | 46.19 |
| P52911 | EXG2 | FASYYAN[+203.079373]DTITVK | 2 | light | 848.4092 | y9 | 1227.61 | 46.19 |
| P52911 | EXG2 | FASYYAN[+203.079373]DTITVK | 2 | light | 848.4092 | y9 | 1024.531 | 46.19 |
| P52911 | EXG2 | FASYYAN[+203.079373]DTITVK | 2 | light | 848.4092 | y8 | 861.4676 | 46.19 |

| | | | | | | | | |
|---|---|---|---|---|---|---|---|---|
| P52911 | EXG2 | FASYYAN[+203.079373]DTITVK | 2 | light | 848.4092 | y7 | 993.5099 | 46.19 |
| P52911 | EXG2 | FASYYAN[+203.079373]DTITVK | 2 | light | 848.4092 | y7 | 790.4305 | 46.19 |
| P52911 | EXG2 | FASYYAN[+203.079373]DTITVK | 2 | light | 848.4092 | y5 | 561.3606 | 46.19 |
| P52911 | EXG2 | FASYYAN[+203.079373]DTITVK | 2 | light | 848.4092 | b3 | 306.1448 | 46.19 |
| P52911 | EXG2 | FASYYAN[+203.079373]DTITVK | 2 | heavy | 852.4163 | y10 | 1398.688 | 46.19 |
| P52911 | EXG2 | FASYYAN[+203.079373]DTITVK | 2 | heavy | 852.4163 | y10 | 1195.608 | 46.19 |
| P52911 | EXG2 | FASYYAN[+203.079373]DTITVK | 2 | heavy | 852.4163 | y9 | 1235.625 | 46.19 |
| P52911 | EXG2 | FASYYAN[+203.079373]DTITVK | 2 | heavy | 852.4163 | y9 | 1032.545 | 46.19 |
| P52911 | EXG2 | FASYYAN[+203.079373]DTITVK | 2 | heavy | 852.4163 | y8 | 869.4818 | 46.19 |
| P52911 | EXG2 | FASYYAN[+203.079373]DTITVK | 2 | heavy | 852.4163 | y7 | 1001.524 | 46.19 |
| P52911 | EXG2 | FASYYAN[+203.079373]DTITVK | 2 | heavy | 852.4163 | y7 | 798.4447 | 46.19 |
| P52911 | EXG2 | FASYYAN[+203.079373]DTITVK | 2 | heavy | 852.4163 | y5 | 569.3748 | 46.19 |
| P52911 | EXG2 | FASYYAN[+203.079373]DTITVK | 2 | heavy | 852.4163 | b3 | 306.1448 | 46.19 |
| P52911 | EXG2 | NLYIDN[+203.079373]ITFND-PYVSDGLQLK | 2 | light | 1323.153 | y11 | 1234.631 | 77.4 |
| P52911 | EXG2 | NLYIDN[+203.079373]ITFND-PYVSDGLQLK | 2 | light | 1323.153 | y10 | 1119.604 | 77.4 |
| P52911 | EXG2 | NLYIDN[+203.079373]ITFND-PYVSDGLQLK | 2 | light | 1323.153 | y8 | 859.4884 | 77.4 |
| P52911 | EXG2 | NLYIDN[+203.079373]ITFND-PYVSDGLQLK | 2 | light | 1323.153 | y7 | 760.4199 | 77.4 |
| P52911 | EXG2 | NLYIDN[+203.079373]ITFND-PYVSDGLQLK | 2 | light | 1323.153 | y6 | 673.3879 | 77.4 |
| P52911 | EXG2 | NLYIDN[+203.079373]ITFND-PYVSDGLQLK | 2 | light | 1323.153 | y5 | 558.361 | 77.4 |
| P52911 | EXG2 | NLYIDN[+203.079373]ITFND-PYVSDGLQLK | 2 | light | 1323.153 | b3 | 391.1976 | 77.4 |
| P52911 | EXG2 | NLYIDN[+203.079373]ITFND-PYVSDGLQLK | 2 | heavy | 1327.16 | y11 | 1242.646 | 77.4 |
| P52911 | EXG2 | NLYIDN[+203.079373]ITFND-PYVSDGLQLK | 2 | heavy | 1327.16 | y10 | 1127.619 | 77.4 |
| P52911 | EXG2 | NLYIDN[+203.079373]ITFND-PYVSDGLQLK | 2 | heavy | 1327.16 | y8 | 867.5026 | 77.4 |
| P52911 | EXG2 | NLYIDN[+203.079373]ITFND-PYVSDGLQLK | 2 | heavy | 1327.16 | y7 | 768.4341 | 77.4 |
| P52911 | EXG2 | NLYIDN[+203.079373]ITFND-PYVSDGLQLK | 2 | heavy | 1327.16 | y6 | 681.4021 | 77.4 |
| P52911 | EXG2 | NLYIDN[+203.079373]ITFND-PYVSDGLQLK | 2 | heavy | 1327.16 | y5 | 566.3752 | 77.4 |
| P52911 | EXG2 | NLYIDN[+203.079373]ITFND-PYVSDGLQLK | 2 | heavy | 1327.16 | b3 | 391.1976 | 77.4 |
| P52911 | EXG2 | IPIGYWAWK | 2 | light | 567.3108 | y8 | 1020.53 | 69.19 |
| P52911 | EXG2 | IPIGYWAWK | 2 | light | 567.3108 | y7 | 923.4774 | 69.19 |
| P52911 | EXG2 | IPIGYWAWK | 2 | light | 567.3108 | y6 | 810.3933 | 69.19 |
| P52911 | EXG2 | IPIGYWAWK | 2 | light | 567.3108 | y4 | 590.3085 | 69.19 |
| P52911 | EXG2 | IPIGYWAWK | 2 | light | 567.3108 | y3 | 404.2292 | 69.19 |

| | | | | | | | | |
|---|---|---|---|---|---|---|---|---|
| P52911 | EXG2 | IPIGYWAWK | 2 | light | 567.3108 | y2 | 333.1921 | 69.19 |
| P52911 | EXG2 | IPIGYWAWK | 2 | light | 567.3108 | y8 | 510.7687 | 69.19 |
| P52911 | EXG2 | IPIGYWAWK | 2 | heavy | 571.3179 | y8 | 1028.544 | 69.19 |
| P52911 | EXG2 | IPIGYWAWK | 2 | heavy | 571.3179 | y7 | 931.4916 | 69.19 |
| P52911 | EXG2 | IPIGYWAWK | 2 | heavy | 571.3179 | y6 | 818.4075 | 69.19 |
| P52911 | EXG2 | IPIGYWAWK | 2 | heavy | 571.3179 | y4 | 598.3227 | 69.19 |
| P52911 | EXG2 | IPIGYWAWK | 2 | heavy | 571.3179 | y3 | 412.2434 | 69.19 |
| P52911 | EXG2 | IPIGYWAWK | 2 | heavy | 571.3179 | y2 | 341.2063 | 69.19 |
| P52911 | EXG2 | IPIGYWAWK | 2 | heavy | 571.3179 | y8 | 514.7758 | 69.19 |
| P52911 | EXG2 | ILYGDLGWLR | 2 | light | 603.3375 | y9 | 1092.584 | 72.66 |
| P52911 | EXG2 | ILYGDLGWLR | 2 | light | 603.3375 | y8 | 979.4996 | 72.66 |
| P52911 | EXG2 | ILYGDLGWLR | 2 | light | 603.3375 | y7 | 816.4363 | 72.66 |
| P52911 | EXG2 | ILYGDLGWLR | 2 | light | 603.3375 | y6 | 759.4148 | 72.66 |
| P52911 | EXG2 | ILYGDLGWLR | 2 | light | 603.3375 | y5 | 644.3879 | 72.66 |
| P52911 | EXG2 | ILYGDLGWLR | 2 | light | 603.3375 | y4 | 531.3038 | 72.66 |
| P52911 | EXG2 | ILYGDLGWLR | 2 | heavy | 606.3476 | y9 | 1098.604 | 72.66 |
| P52911 | EXG2 | ILYGDLGWLR | 2 | heavy | 606.3476 | y8 | 985.5197 | 72.66 |
| P52911 | EXG2 | ILYGDLGWLR | 2 | heavy | 606.3476 | y7 | 822.4564 | 72.66 |
| P52911 | EXG2 | ILYGDLGWLR | 2 | heavy | 606.3476 | y6 | 765.4349 | 72.66 |
| P52911 | EXG2 | ILYGDLGWLR | 2 | heavy | 606.3476 | y5 | 650.408 | 72.66 |
| P52911 | EXG2 | ILYGDLGWLR | 2 | heavy | 606.3476 | y4 | 537.3239 | 72.66 |
| P23797 | GPI12 | VRELN[+203.079373]ESAALLLH-NER | 3 | light | 689.6992 | y10 | 1123.622 | 43.25 |
| P23797 | GPI12 | VRELN[+203.079373]ESAALLLH-NER | 3 | light | 689.6992 | y9 | 1036.59 | 43.25 |
| P23797 | GPI12 | VRELN[+203.079373]ESAALLLH-NER | 3 | light | 689.6992 | y8 | 965.5527 | 43.25 |
| P23797 | GPI12 | VRELN[+203.079373]ESAALLLH-NER | 3 | light | 689.6992 | y7 | 894.5156 | 43.25 |
| P23797 | GPI12 | VRELN[+203.079373]ESAALLLH-NER | 3 | light | 689.6992 | y6 | 781.4315 | 43.25 |
| P23797 | GPI12 | VRELN[+203.079373]ESAALLLH-NER | 3 | light | 689.6992 | y5 | 668.3474 | 43.25 |
| P23797 | GPI12 | VRELN[+203.079373]ESAALLLH-NER | 3 | light | 689.6992 | y4 | 555.2634 | 43.25 |
| P23797 | GPI12 | VRELN[+203.079373]ESAALLLH-NER | 3 | light | 689.6992 | b4 | 498.3035 | 43.25 |
| P23797 | GPI12 | VRELN[+203.079373]ESAALLLH-NER | 3 | light | 689.6992 | b5 | 815.4258 | 43.25 |
| P23797 | GPI12 | VRELN[+203.079373]ESAALLLH-NER | 3 | light | 689.6992 | b5 | 612.3464 | 43.25 |
| P23797 | GPI12 | VRELN[+203.079373]ESAALLLH-NER | 3 | light | 689.6992 | b6 | 944.4684 | 43.25 |

| | | | | | | | | |
|---|---|---|---|---|---|---|---|---|
| P23797 | GPI12 | VRELN[+203.079373]ESAALLLH-NER | 3 | heavy | 693.7126 | y10 | 1129.642 | 43.25 |
| P23797 | GPI12 | VRELN[+203.079373]ESAALLLH-NER | 3 | heavy | 693.7126 | y9 | 1042.61 | 43.25 |
| P23797 | GPI12 | VRELN[+203.079373]ESAALLLH-NER | 3 | heavy | 693.7126 | y8 | 971.5728 | 43.25 |
| P23797 | GPI12 | VRELN[+203.079373]ESAALLLH-NER | 3 | heavy | 693.7126 | y7 | 900.5357 | 43.25 |
| P23797 | GPI12 | VRELN[+203.079373]ESAALLLH-NER | 3 | heavy | 693.7126 | y6 | 787.4516 | 43.25 |
| P23797 | GPI12 | VRELN[+203.079373]ESAALLLH-NER | 3 | heavy | 693.7126 | y5 | 674.3676 | 43.25 |
| P23797 | GPI12 | VRELN[+203.079373]ESAALLLH-NER | 3 | heavy | 693.7126 | y4 | 561.2835 | 43.25 |
| P23797 | GPI12 | VRELN[+203.079373]ESAALLLH-NER | 3 | heavy | 693.7126 | b4 | 504.3236 | 43.25 |
| P23797 | GPI12 | VRELN[+203.079373]ESAALLLH-NER | 3 | heavy | 693.7126 | b5 | 821.4459 | 43.25 |
| P23797 | GPI12 | VRELN[+203.079373]ESAALLLH-NER | 3 | heavy | 693.7126 | b5 | 618.3665 | 43.25 |
| P23797 | GPI12 | VRELN[+203.079373]ESAALLLH-NER | 3 | heavy | 693.7126 | b6 | 950.4885 | 43.25 |
| P23797 | GPI12 | TVPFNIIC[+57.021464]LSK | 2 | light | 646.3576 | y9 | 1091.592 | 70.09 |
| P23797 | GPI12 | TVPFNIIC[+57.021464]LSK | 2 | light | 646.3576 | y8 | 994.539 | 70.09 |
| P23797 | GPI12 | TVPFNIIC[+57.021464]LSK | 2 | light | 646.3576 | y7 | 847.4706 | 70.09 |
| P23797 | GPI12 | TVPFNIIC[+57.021464]LSK | 2 | light | 646.3576 | y6 | 733.4277 | 70.09 |
| P23797 | GPI12 | TVPFNIIC[+57.021464]LSK | 2 | light | 646.3576 | y5 | 620.3436 | 70.09 |
| P23797 | GPI12 | TVPFNIIC[+57.021464]LSK | 2 | light | 646.3576 | y4 | 507.2595 | 70.09 |
| P23797 | GPI12 | TVPFNIIC[+57.021464]LSK | 2 | light | 646.3576 | y3 | 347.2289 | 70.09 |
| P23797 | GPI12 | TVPFNIIC[+57.021464]LSK | 2 | heavy | 650.3647 | y9 | 1099.606 | 70.09 |
| P23797 | GPI12 | TVPFNIIC[+57.021464]LSK | 2 | heavy | 650.3647 | y8 | 1002.553 | 70.09 |
| P23797 | GPI12 | TVPFNIIC[+57.021464]LSK | 2 | heavy | 650.3647 | y7 | 855.4848 | 70.09 |
| P23797 | GPI12 | TVPFNIIC[+57.021464]LSK | 2 | heavy | 650.3647 | y6 | 741.4419 | 70.09 |
| P23797 | GPI12 | TVPFNIIC[+57.021464]LSK | 2 | heavy | 650.3647 | y5 | 628.3578 | 70.09 |
| P23797 | GPI12 | TVPFNIIC[+57.021464]LSK | 2 | heavy | 650.3647 | y4 | 515.2737 | 70.09 |
| P23797 | GPI12 | TVPFNIIC[+57.021464]LSK | 2 | heavy | 650.3647 | y3 | 355.2431 | 70.09 |
| P23797 | GPI12 | GNAEGLGETR | 2 | light | 502.2438 | y8 | 832.4159 | 18.46 |
| P23797 | GPI12 | GNAEGLGETR | 2 | light | 502.2438 | y7 | 761.3788 | 18.46 |
| P23797 | GPI12 | GNAEGLGETR | 2 | light | 502.2438 | y6 | 632.3362 | 18.46 |
| P23797 | GPI12 | GNAEGLGETR | 2 | light | 502.2438 | y5 | 575.3148 | 18.46 |
| P23797 | GPI12 | GNAEGLGETR | 2 | light | 502.2438 | y4 | 462.2307 | 18.46 |
| P23797 | GPI12 | GNAEGLGETR | 2 | light | 502.2438 | y3 | 405.2092 | 18.46 |
| P23797 | GPI12 | GNAEGLGETR | 2 | heavy | 505.2539 | y8 | 838.4361 | 18.46 |

| | | | | | | | | |
|---|---|---|---|---|---|---|---|---|
| P23797 | GPI12 | GNAEGLGETR | 2 | heavy | 505.2539 | y7 | 767.3989 | 18.46 |
| P23797 | GPI12 | GNAEGLGETR | 2 | heavy | 505.2539 | y6 | 638.3563 | 18.46 |
| P23797 | GPI12 | GNAEGLGETR | 2 | heavy | 505.2539 | y5 | 581.3349 | 18.46 |
| P23797 | GPI12 | GNAEGLGETR | 2 | heavy | 505.2539 | y4 | 468.2508 | 18.46 |
| P23797 | GPI12 | GNAEGLGETR | 2 | heavy | 505.2539 | y3 | 411.2294 | 18.46 |
| P40345 | PDAT | SSSEDALNN[+203.079373]NTDTYGNFIR | 2 | light | 1161.012 | y10 | 1200.564 | 49.1 |
| P40345 | PDAT | SSSEDALNN[+203.079373]NTDTYGNFIR | 2 | light | 1161.012 | y9 | 1086.521 | 49.1 |
| P40345 | PDAT | SSSEDALNN[+203.079373]NTDTYGNFIR | 2 | light | 1161.012 | y8 | 985.4738 | 49.1 |
| P40345 | PDAT | SSSEDALNN[+203.079373]NTDTYGNFIR | 2 | light | 1161.012 | y7 | 870.4468 | 49.1 |
| P40345 | PDAT | SSSEDALNN[+203.079373]NTDTYGNFIR | 2 | light | 1161.012 | y6 | 769.3991 | 49.1 |
| P40345 | PDAT | SSSEDALNN[+203.079373]NTDTYGNFIR | 2 | light | 1161.012 | y5 | 606.3358 | 49.1 |
| P40345 | PDAT | SSSEDALNN[+203.079373]NTDTYGNFIR | 2 | heavy | 1164.022 | y10 | 1206.585 | 49.1 |
| P40345 | PDAT | SSSEDALNN[+203.079373]NTDTYGNFIR | 2 | heavy | 1164.022 | y9 | 1092.542 | 49.1 |
| P40345 | PDAT | SSSEDALNN[+203.079373]NTDTYGNFIR | 2 | heavy | 1164.022 | y8 | 991.4939 | 49.1 |
| P40345 | PDAT | SSSEDALNN[+203.079373]NTDTYGNFIR | 2 | heavy | 1164.022 | y7 | 876.467 | 49.1 |
| P40345 | PDAT | SSSEDALNN[+203.079373]NTDTYGNFIR | 2 | heavy | 1164.022 | y6 | 775.4193 | 49.1 |
| P40345 | PDAT | SSSEDALNN[+203.079373]NTDTYGNFIR | 2 | heavy | 1164.022 | y5 | 612.356 | 49.1 |
| P40345 | PDAT | MLQTWGGIPSMLPK | 2 | light | 779.9096 | y11 | 1186.629 | 74.32 |
| P40345 | PDAT | MLQTWGGIPSMLPK | 2 | light | 779.9096 | y10 | 1085.581 | 74.32 |
| P40345 | PDAT | MLQTWGGIPSMLPK | 2 | light | 779.9096 | y9 | 899.5019 | 74.32 |
| P40345 | PDAT | MLQTWGGIPSMLPK | 2 | light | 779.9096 | y8 | 842.4804 | 74.32 |
| P40345 | PDAT | MLQTWGGIPSMLPK | 2 | light | 779.9096 | y6 | 672.3749 | 74.32 |
| P40345 | PDAT | MLQTWGGIPSMLPK | 2 | light | 779.9096 | y2 | 244.1656 | 74.32 |
| P40345 | PDAT | MLQTWGGIPSMLPK | 2 | heavy | 783.9167 | y11 | 1194.643 | 74.32 |
| P40345 | PDAT | MLQTWGGIPSMLPK | 2 | heavy | 783.9167 | y10 | 1093.595 | 74.32 |
| P40345 | PDAT | MLQTWGGIPSMLPK | 2 | heavy | 783.9167 | y9 | 907.5161 | 74.32 |
| P40345 | PDAT | MLQTWGGIPSMLPK | 2 | heavy | 783.9167 | y8 | 850.4946 | 74.32 |
| P40345 | PDAT | MLQTWGGIPSMLPK | 2 | heavy | 783.9167 | y6 | 680.3891 | 74.32 |
| P40345 | PDAT | MLQTWGGIPSMLPK | 2 | heavy | 783.9167 | y2 | 252.1798 | 74.32 |
| P40345 | PDAT | IASGNGDLVEPR | 2 | light | 614.32 | y8 | 899.4581 | 32.82 |
| P40345 | PDAT | IASGNGDLVEPR | 2 | light | 614.32 | y7 | 785.4152 | 32.82 |
| P40345 | PDAT | IASGNGDLVEPR | 2 | light | 614.32 | y6 | 728.3937 | 32.82 |

| | | | | | | | | |
|---|---|---|---|---|---|---|---|---|
| P40345 | PDAT | IASGNGDLVEPR | 2 | light | 614.32 | y4 | 500.2827 | 32.82 |
| P40345 | PDAT | IASGNGDLVEPR | 2 | heavy | 617.3301 | y8 | 905.4783 | 32.82 |
| P40345 | PDAT | IASGNGDLVEPR | 2 | heavy | 617.3301 | y7 | 791.4353 | 32.82 |
| P40345 | PDAT | IASGNGDLVEPR | 2 | heavy | 617.3301 | y6 | 734.4139 | 32.82 |
| P40345 | PDAT | IASGNGDLVEPR | 2 | heavy | 617.3301 | y4 | 506.3029 | 32.82 |
| P38244 | M28P1orPFF1 | SILFQQQDPFN[+203.079373]ESSR | 2 | light | 999.9738 | y8 | 1154.496 | 58.13 |
| P38244 | M28P1orPFF1 | SILFQQQDPFN[+203.079373]ESSR | 2 | light | 999.9738 | y7 | 1039.469 | 58.13 |
| P38244 | M28P1orPFF1 | SILFQQQDPFN[+203.079373]ESSR | 2 | light | 999.9738 | y7 | 836.3897 | 58.13 |
| P38244 | M28P1orPFF1 | SILFQQQDPFN[+203.079373]ESSR | 2 | light | 999.9738 | y3 | 349.183 | 58.13 |
| P38244 | M28P1orPFF1 | SILFQQQDPFN[+203.079373]ESSR | 2 | light | 999.9738 | b3 | 314.2074 | 58.13 |
| P38244 | M28P1orPFF1 | SILFQQQDPFN[+203.079373]ESSR | 2 | heavy | 1002.984 | y8 | 1160.516 | 58.13 |
| P38244 | M28P1orPFF1 | SILFQQQDPFN[+203.079373]ESSR | 2 | heavy | 1002.984 | y7 | 1045.489 | 58.13 |
| P38244 | M28P1orPFF1 | SILFQQQDPFN[+203.079373]ESSR | 2 | heavy | 1002.984 | y7 | 842.4098 | 58.13 |
| P38244 | M28P1orPFF1 | SILFQQQDPFN[+203.079373]ESSR | 2 | heavy | 1002.984 | y3 | 355.2031 | 58.13 |
| P38244 | M28P1orPFF1 | SILFQQQDPFN[+203.079373]ESSR | 2 | heavy | 1002.984 | b3 | 314.2074 | 58.13 |
| P38244 | M28P1orPFF1 | GSNSMEEGLSTR | 2 | light | 634.2828 | y9 | 1009.462 | 26.97 |
| P38244 | M28P1orPFF1 | GSNSMEEGLSTR | 2 | light | 634.2828 | y8 | 922.4299 | 26.97 |
| P38244 | M28P1orPFF1 | GSNSMEEGLSTR | 2 | light | 634.2828 | y7 | 791.3894 | 26.97 |
| P38244 | M28P1orPFF1 | GSNSMEEGLSTR | 2 | light | 634.2828 | y6 | 662.3468 | 26.97 |
| P38244 | M28P1orPFF1 | GSNSMEEGLSTR | 2 | light | 634.2828 | y5 | 533.3042 | 26.97 |
| P38244 | M28P1orPFF1 | GSNSMEEGLSTR | 2 | heavy | 637.2929 | y9 | 1015.482 | 26.97 |
| P38244 | M28P1orPFF1 | GSNSMEEGLSTR | 2 | heavy | 637.2929 | y8 | 928.45 | 26.97 |
| P38244 | M28P1orPFF1 | GSNSMEEGLSTR | 2 | heavy | 637.2929 | y7 | 797.4095 | 26.97 |

| | | | | | | | | |
|---|---|---|---|---|---|---|---|---|
| P38244 | M28P1orPFF1 | GSNSMEEGLSTR | 2 | heavy | 637.2929 | y6 | 668.3669 | 26.97 |
| P38244 | M28P1orPFF1 | GSNSMEEGLSTR | 2 | heavy | 637.2929 | y5 | 539.3243 | 26.97 |
| P38244 | M28P1orPFF1 | FSQNIDLSQGNAASVHVLGR | 2 | light | 1057.043 | y11 | 1080.591 | 51.4 |
| P38244 | M28P1orPFF1 | FSQNIDLSQGNAASVHVLGR | 2 | light | 1057.043 | y10 | 1023.569 | 51.4 |
| P38244 | M28P1orPFF1 | FSQNIDLSQGNAASVHVLGR | 2 | light | 1057.043 | y9 | 909.5265 | 51.4 |
| P38244 | M28P1orPFF1 | FSQNIDLSQGNAASVHVLGR | 2 | light | 1057.043 | y8 | 838.4894 | 51.4 |
| P38244 | M28P1orPFF1 | FSQNIDLSQGNAASVHVLGR | 2 | light | 1057.043 | y7 | 767.4522 | 51.4 |
| P38244 | M28P1orPFF1 | FSQNIDLSQGNAASVHVLGR | 2 | light | 1057.043 | y6 | 680.4202 | 51.4 |
| P38244 | M28P1orPFF1 | FSQNIDLSQGNAASVHVLGR | 2 | light | 1057.043 | y5 | 581.3518 | 51.4 |
| P38244 | M28P1orPFF1 | FSQNIDLSQGNAASVHVLGR | 2 | light | 1057.043 | y4 | 444.2929 | 51.4 |
| P38244 | M28P1orPFF1 | FSQNIDLSQGNAASVHVLGR | 2 | heavy | 1060.053 | y11 | 1086.611 | 51.4 |
| P38244 | M28P1orPFF1 | FSQNIDLSQGNAASVHVLGR | 2 | heavy | 1060.053 | y10 | 1029.59 | 51.4 |
| P38244 | M28P1orPFF1 | FSQNIDLSQGNAASVHVLGR | 2 | heavy | 1060.053 | y9 | 915.5466 | 51.4 |
| P38244 | M28P1orPFF1 | FSQNIDLSQGNAASVHVLGR | 2 | heavy | 1060.053 | y8 | 844.5095 | 51.4 |
| P38244 | M28P1orPFF1 | FSQNIDLSQGNAASVHVLGR | 2 | heavy | 1060.053 | y7 | 773.4724 | 51.4 |
| P38244 | M28P1orPFF1 | FSQNIDLSQGNAASVHVLGR | 2 | heavy | 1060.053 | y6 | 686.4403 | 51.4 |
| P38244 | M28P1orPFF1 | FSQNIDLSQGNAASVHVLGR | 2 | heavy | 1060.053 | y5 | 587.3719 | 51.4 |
| P38244 | M28P1orPFF1 | FSQNIDLSQGNAASVHVLGR | 2 | heavy | 1060.053 | y4 | 450.313 | 51.4 |
| P00729 | CBPY | VRN[+203.079373]WTASITDEVA-GEVK | 3 | light | 693.3515 | y9 | 947.468 | 55.5 |
| P00729 | CBPY | VRN[+203.079373]WTASITDEVA-GEVK | 3 | light | 693.3515 | y8 | 846.4203 | 55.5 |
| P00729 | CBPY | VRN[+203.079373]WTASITDEVA-GEVK | 3 | light | 693.3515 | y6 | 602.3508 | 55.5 |
| P00729 | CBPY | VRN[+203.079373]WTASITDEVA-GEVK | 3 | light | 693.3515 | b6 | 931.4632 | 55.5 |
| P00729 | CBPY | VRN[+203.079373]WTASITDEVA-GEVK | 3 | light | 693.3515 | b7 | 1018.495 | 55.5 |

| | | | | | | | | |
|---|---|---|---|---|---|---|---|---|
| P00729 | CBPY | VRN[+203.079373]WTASITDEVA-GEVK | 3 | heavy | 698.0296 | y9 | 955.4822 | 55.5 |
| P00729 | CBPY | VRN[+203.079373]WTASITDEVA-GEVK | 3 | heavy | 698.0296 | y8 | 854.4345 | 55.5 |
| P00729 | CBPY | VRN[+203.079373]WTASITDEVA-GEVK | 3 | heavy | 698.0296 | y6 | 610.365 | 55.5 |
| P00729 | CBPY | VRN[+203.079373]WTASITDEVA-GEVK | 3 | heavy | 698.0296 | b6 | 937.4833 | 55.5 |
| P00729 | CBPY | VRN[+203.079373]WTASITDEVA-GEVK | 3 | heavy | 698.0296 | b7 | 1024.515 | 55.5 |
| P00729 | CBPY | ILGIDPN[+203.079373]VTQYT-GYLDVEDEDK | 2 | light | 1350.642 | y11 | 1283.564 | 74.69 |
| P00729 | CBPY | ILGIDPN[+203.079373]VTQYT-GYLDVEDEDK | 2 | light | 1350.642 | y10 | 1182.516 | 74.69 |
| P00729 | CBPY | ILGIDPN[+203.079373]VTQYT-GYLDVEDEDK | 2 | light | 1350.642 | y7 | 849.3472 | 74.69 |
| P00729 | CBPY | ILGIDPN[+203.079373]VTQYT-GYLDVEDEDK | 2 | light | 1350.642 | y6 | 734.3203 | 74.69 |
| P00729 | CBPY | ILGIDPN[+203.079373]VTQYT-GYLDVEDEDK | 2 | light | 1350.642 | y4 | 506.2093 | 74.69 |
| P00729 | CBPY | ILGIDPN[+203.079373]VTQYT-GYLDVEDEDK | 2 | light | 1350.642 | b2 | 227.1754 | 74.69 |
| P00729 | CBPY | ILGIDPN[+203.079373]VTQYT-GYLDVEDEDK | 2 | light | 1350.642 | b3 | 284.1969 | 74.69 |
| P00729 | CBPY | ILGIDPN[+203.079373]VTQYT-GYLDVEDEDK | 2 | heavy | 1354.649 | y11 | 1291.578 | 74.69 |
| P00729 | CBPY | ILGIDPN[+203.079373]VTQYT-GYLDVEDEDK | 2 | heavy | 1354.649 | y10 | 1190.53 | 74.69 |
| P00729 | CBPY | ILGIDPN[+203.079373]VTQYT-GYLDVEDEDK | 2 | heavy | 1354.649 | y7 | 857.3614 | 74.69 |
| P00729 | CBPY | ILGIDPN[+203.079373]VTQYT-GYLDVEDEDK | 2 | heavy | 1354.649 | y6 | 742.3345 | 74.69 |
| P00729 | CBPY | ILGIDPN[+203.079373]VTQYT-GYLDVEDEDK | 2 | heavy | 1354.649 | y4 | 514.2235 | 74.69 |
| P00729 | CBPY | ILGIDPN[+203.079373]VTQYT-GYLDVEDEDK | 2 | heavy | 1354.649 | b2 | 227.1754 | 74.69 |
| P00729 | CBPY | ILGIDPN[+203.079373]VTQYT-GYLDVEDEDK | 2 | heavy | 1354.649 | b3 | 284.1969 | 74.69 |
| P00729 | CBPY | AWTDVLPWK | 2 | light | 558.2978 | y7 | 858.472 | 72.08 |
| P00729 | CBPY | AWTDVLPWK | 2 | light | 558.2978 | y6 | 757.4243 | 72.08 |
| P00729 | CBPY | AWTDVLPWK | 2 | light | 558.2978 | y5 | 642.3974 | 72.08 |
| P00729 | CBPY | AWTDVLPWK | 2 | light | 558.2978 | y4 | 543.3289 | 72.08 |
| P00729 | CBPY | AWTDVLPWK | 2 | light | 558.2978 | y3 | 430.2449 | 72.08 |
| P00729 | CBPY | AWTDVLPWK | 2 | heavy | 562.3049 | y7 | 866.4862 | 72.08 |
| P00729 | CBPY | AWTDVLPWK | 2 | heavy | 562.3049 | y6 | 765.4385 | 72.08 |
| P00729 | CBPY | AWTDVLPWK | 2 | heavy | 562.3049 | y5 | 650.4116 | 72.08 |
| P00729 | CBPY | AWTDVLPWK | 2 | heavy | 562.3049 | y4 | 551.3431 | 72.08 |
| P00729 | CBPY | AWTDVLPWK | 2 | heavy | 562.3049 | y3 | 438.2591 | 72.08 |
| P00729 | CBPY | DFIC[+57.021464]NWLGNK | 2 | light | 633.8004 | y8 | 1004.498 | 67.2 |
| P00729 | CBPY | DFIC[+57.021464]NWLGNK | 2 | light | 633.8004 | y7 | 891.4141 | 67.2 |
| P00729 | CBPY | DFIC[+57.021464]NWLGNK | 2 | light | 633.8004 | y6 | 731.3835 | 67.2 |

| | | | | | | | | |
|---|---|---|---|---|---|---|---|---|
| P00729 | CBPY | DFIC[+57.021464]NWLGNK | 2 | light | 633.8004 | y5 | 617.3406 | 67.2 |
| P00729 | CBPY | DFIC[+57.021464]NWLGNK | 2 | light | 633.8004 | y4 | 431.2613 | 67.2 |
| P00729 | CBPY | DFIC[+57.021464]NWLGNK | 2 | light | 633.8004 | y3 | 318.1772 | 67.2 |
| P00729 | CBPY | DFIC[+57.021464]NWLGNK | 2 | heavy | 637.8075 | y8 | 1012.512 | 67.2 |
| P00729 | CBPY | DFIC[+57.021464]NWLGNK | 2 | heavy | 637.8075 | y7 | 899.4283 | 67.2 |
| P00729 | CBPY | DFIC[+57.021464]NWLGNK | 2 | heavy | 637.8075 | y6 | 739.3977 | 67.2 |
| P00729 | CBPY | DFIC[+57.021464]NWLGNK | 2 | heavy | 637.8075 | y5 | 625.3548 | 67.2 |
| P00729 | CBPY | DFIC[+57.021464]NWLGNK | 2 | heavy | 637.8075 | y4 | 439.2755 | 67.2 |
| P00729 | CBPY | DFIC[+57.021464]NWLGNK | 2 | heavy | 637.8075 | y3 | 326.1914 | 67.2 |
| P38875 | GPI16 | SYASDIGAPLFN[+203.079373]STEK | 2 | light | 951.952 | y11 | 1379.705 | 56.29 |
| P38875 | GPI16 | SYASDIGAPLFN[+203.079373]STEK | 2 | light | 951.952 | y11 | 1176.626 | 56.29 |
| P38875 | GPI16 | SYASDIGAPLFN[+203.079373]STEK | 2 | light | 951.952 | y10 | 1266.621 | 56.29 |
| P38875 | GPI16 | SYASDIGAPLFN[+203.079373]STEK | 2 | light | 951.952 | y10 | 1063.542 | 56.29 |
| P38875 | GPI16 | SYASDIGAPLFN[+203.079373]STEK | 2 | light | 951.952 | y9 | 1209.6 | 56.29 |
| P38875 | GPI16 | SYASDIGAPLFN[+203.079373]STEK | 2 | light | 951.952 | y9 | 1006.52 | 56.29 |
| P38875 | GPI16 | SYASDIGAPLFN[+203.079373]STEK | 2 | light | 951.952 | y8 | 1138.563 | 56.29 |
| P38875 | GPI16 | SYASDIGAPLFN[+203.079373]STEK | 2 | light | 951.952 | y8 | 935.4833 | 56.29 |
| P38875 | GPI16 | SYASDIGAPLFN[+203.079373]STEK | 2 | light | 951.952 | y6 | 928.4258 | 56.29 |
| P38875 | GPI16 | SYASDIGAPLFN[+203.079373]STEK | 2 | light | 951.952 | y6 | 725.3464 | 56.29 |
| P38875 | GPI16 | SYASDIGAPLFN[+203.079373]STEK | 2 | light | 951.952 | y5 | 781.3574 | 56.29 |
| P38875 | GPI16 | SYASDIGAPLFN[+203.079373]STEK | 2 | light | 951.952 | b2 | 251.1026 | 56.29 |
| P38875 | GPI16 | SYASDIGAPLFN[+203.079373]STEK | 2 | light | 951.952 | b3 | 322.1397 | 56.29 |
| P38875 | GPI16 | SYASDIGAPLFN[+203.079373]STEK | 2 | heavy | 955.9591 | y11 | 1387.719 | 56.29 |
| P38875 | GPI16 | SYASDIGAPLFN[+203.079373]STEK | 2 | heavy | 955.9591 | y11 | 1184.64 | 56.29 |
| P38875 | GPI16 | SYASDIGAPLFN[+203.079373]STEK | 2 | heavy | 955.9591 | y10 | 1274.635 | 56.29 |
| P38875 | GPI16 | SYASDIGAPLFN[+203.079373]STEK | 2 | heavy | 955.9591 | y10 | 1071.556 | 56.29 |
| P38875 | GPI16 | SYASDIGAPLFN[+203.079373]STEK | 2 | heavy | 955.9591 | y9 | 1217.614 | 56.29 |
| P38875 | GPI16 | SYASDIGAPLFN[+203.079373]STEK | 2 | heavy | 955.9591 | y9 | 1014.535 | 56.29 |
| P38875 | GPI16 | SYASDIGAPLFN[+203.079373]STEK | 2 | heavy | 955.9591 | y8 | 1146.577 | 56.29 |
| P38875 | GPI16 | SYASDIGAPLFN[+203.079373]STEK | 2 | heavy | 955.9591 | y8 | 943.4975 | 56.29 |
| P38875 | GPI16 | SYASDIGAPLFN[+203.079373]STEK | 2 | heavy | 955.9591 | y6 | 936.44 | 56.29 |
| P38875 | GPI16 | SYASDIGAPLFN[+203.079373]STEK | 2 | heavy | 955.9591 | y6 | 733.3606 | 56.29 |

| | | | | | | | | |
|---|---|---|---|---|---|---|---|---|
| P38875 | GPI16 | SYASDIGAPLFN[+203.079373]STEK | 2 | heavy | 955.9591 | y5 | 789.3716 | 56.29 |
| P38875 | GPI16 | SYASDIGAPLFN[+203.079373]STEK | 2 | heavy | 955.9591 | b2 | 251.1026 | 56.29 |
| P38875 | GPI16 | SYASDIGAPLFN[+203.079373]STEK | 2 | heavy | 955.9591 | b3 | 322.1397 | 56.29 |
| P38875 | GPI16 | AIPPLLESTATR | 2 | light | 634.8641 | y10 | 1084.6 | 54.76 |
| P38875 | GPI16 | AIPPLLESTATR | 2 | light | 634.8641 | y9 | 987.5469 | 54.76 |
| P38875 | GPI16 | AIPPLLESTATR | 2 | light | 634.8641 | y8 | 890.4942 | 54.76 |
| P38875 | GPI16 | AIPPLLESTATR | 2 | light | 634.8641 | y7 | 777.4101 | 54.76 |
| P38875 | GPI16 | AIPPLLESTATR | 2 | light | 634.8641 | y6 | 664.326 | 54.76 |
| P38875 | GPI16 | AIPPLLESTATR | 2 | light | 634.8641 | y5 | 535.2835 | 54.76 |
| P38875 | GPI16 | AIPPLLESTATR | 2 | heavy | 637.8741 | y10 | 1090.62 | 54.76 |
| P38875 | GPI16 | AIPPLLESTATR | 2 | heavy | 637.8741 | y9 | 993.5671 | 54.76 |
| P38875 | GPI16 | AIPPLLESTATR | 2 | heavy | 637.8741 | y8 | 896.5143 | 54.76 |
| P38875 | GPI16 | AIPPLLESTATR | 2 | heavy | 637.8741 | y7 | 783.4302 | 54.76 |
| P38875 | GPI16 | AIPPLLESTATR | 2 | heavy | 637.8741 | y6 | 670.3462 | 54.76 |
| P38875 | GPI16 | AIPPLLESTATR | 2 | heavy | 637.8741 | y5 | 541.3036 | 54.76 |
| P38875 | GPI16 | VTPIVPVPIHVSR | 3 | light | 471.9574 | y9 | 1003.605 | 57.06 |
| P38875 | GPI16 | VTPIVPVPIHVSR | 3 | light | 471.9574 | y8 | 904.5363 | 57.06 |
| P38875 | GPI16 | VTPIVPVPIHVSR | 3 | light | 471.9574 | y7 | 807.4835 | 57.06 |
| P38875 | GPI16 | VTPIVPVPIHVSR | 3 | light | 471.9574 | y6 | 708.4151 | 57.06 |
| P38875 | GPI16 | VTPIVPVPIHVSR | 3 | light | 471.9574 | y5 | 611.3624 | 57.06 |
| P38875 | GPI16 | VTPIVPVPIHVSR | 3 | light | 471.9574 | y4 | 498.2783 | 57.06 |
| P38875 | GPI16 | VTPIVPVPIHVSR | 3 | light | 471.9574 | y3 | 361.2194 | 57.06 |
| P38875 | GPI16 | VTPIVPVPIHVSR | 3 | heavy | 473.9641 | y9 | 1009.625 | 57.06 |
| P38875 | GPI16 | VTPIVPVPIHVSR | 3 | heavy | 473.9641 | y8 | 910.5564 | 57.06 |
| P38875 | GPI16 | VTPIVPVPIHVSR | 3 | heavy | 473.9641 | y7 | 813.5037 | 57.06 |
| P38875 | GPI16 | VTPIVPVPIHVSR | 3 | heavy | 473.9641 | y6 | 714.4353 | 57.06 |
| P38875 | GPI16 | VTPIVPVPIHVSR | 3 | heavy | 473.9641 | y5 | 617.3825 | 57.06 |
| P38875 | GPI16 | VTPIVPVPIHVSR | 3 | heavy | 473.9641 | y4 | 504.2984 | 57.06 |
| P38875 | GPI16 | VTPIVPVPIHVSR | 3 | heavy | 473.9641 | y3 | 367.2395 | 57.06 |
| P17967 | PDI | NSDVN[+203.079373]NSIDYEGPR | 2 | light | 891.8925 | y9 | 1050.485 | 32.47 |
| P17967 | PDI | NSDVN[+203.079373]NSIDYEGPR | 2 | light | 891.8925 | y8 | 936.4421 | 32.47 |
| P17967 | PDI | NSDVN[+203.079373]NSIDYEGPR | 2 | light | 891.8925 | y6 | 736.326 | 32.47 |

| | | | | | | | | |
|---|---|---|---|---|---|---|---|---|
| P17967 | PDI | NSDVN[+203.079373]NSIDYEGPR | 2 | light | 891.8925 | y5 | 621.2991 | 32.47 |
| P17967 | PDI | NSDVN[+203.079373]NSIDYEGPR | 2 | light | 891.8925 | y4 | 458.2358 | 32.47 |
| P17967 | PDI | NSDVN[+203.079373]NSIDYEGPR | 2 | light | 891.8925 | y3 | 329.1932 | 32.47 |
| P17967 | PDI | NSDVN[+203.079373]NSIDYEGPR | 2 | light | 891.8925 | b4 | 416.1776 | 32.47 |
| P17967 | PDI | NSDVN[+203.079373]NSIDYEGPR | 2 | light | 891.8925 | b5 | 530.2205 | 32.47 |
| P17967 | PDI | NSDVN[+203.079373]NSIDYEGPR | 2 | light | 891.8925 | b6 | 644.2634 | 32.47 |
| P17967 | PDI | NSDVN[+203.079373]NSIDYEGPR | 2 | heavy | 894.9025 | y9 | 1056.505 | 32.47 |
| P17967 | PDI | NSDVN[+203.079373]NSIDYEGPR | 2 | heavy | 894.9025 | y8 | 942.4623 | 32.47 |
| P17967 | PDI | NSDVN[+203.079373]NSIDYEGPR | 2 | heavy | 894.9025 | y6 | 742.3462 | 32.47 |
| P17967 | PDI | NSDVN[+203.079373]NSIDYEGPR | 2 | heavy | 894.9025 | y5 | 627.3192 | 32.47 |
| P17967 | PDI | NSDVN[+203.079373]NSIDYEGPR | 2 | heavy | 894.9025 | y4 | 464.2559 | 32.47 |
| P17967 | PDI | NSDVN[+203.079373]NSIDYEGPR | 2 | heavy | 894.9025 | y3 | 335.2133 | 32.47 |
| P17967 | PDI | NSDVN[+203.079373]NSIDYEGPR | 2 | heavy | 894.9025 | b4 | 416.1776 | 32.47 |
| P17967 | PDI | NSDVN[+203.079373]NSIDYEGPR | 2 | heavy | 894.9025 | b5 | 530.2205 | 32.47 |
| P17967 | PDI | NSDVN[+203.079373]NSIDYEGPR | 2 | heavy | 894.9025 | b6 | 644.2634 | 32.47 |
| P17967 | PDI | IDADFN[+203.079373]ATFYSM[+15.994915]ANK | 2 | light | 963.9249 | y8 | 977.4397 | 51.23 |
| P17967 | PDI | IDADFN[+203.079373]ATFYSM[+15.994915]ANK | 2 | light | 963.9249 | y7 | 876.392 | 51.23 |
| P17967 | PDI | IDADFN[+203.079373]ATFYSM[+15.994915]ANK | 2 | light | 963.9249 | y7 | 812.3937 | 51.23 |
| P17967 | PDI | IDADFN[+203.079373]ATFYSM[+15.994915]ANK | 2 | light | 963.9249 | y6 | 729.3236 | 51.23 |
| P17967 | PDI | IDADFN[+203.079373]ATFYSM[+15.994915]ANK | 2 | light | 963.9249 | y6 | 665.3253 | 51.23 |
| P17967 | PDI | IDADFN[+203.079373]ATFYSM[+15.994915]ANK | 2 | heavy | 967.932 | y8 | 985.4539 | 51.23 |
| P17967 | PDI | IDADFN[+203.079373]ATFYSM[+15.994915]ANK | 2 | heavy | 967.932 | y7 | 884.4062 | 51.23 |
| P17967 | PDI | IDADFN[+203.079373]ATFYSM[+15.994915]ANK | 2 | heavy | 967.932 | y7 | 820.4079 | 51.23 |
| P17967 | PDI | IDADFN[+203.079373]ATFYSM[+15.994915]ANK | 2 | heavy | 967.932 | y6 | 737.3378 | 51.23 |
| P17967 | PDI | IDADFN[+203.079373]ATFYSM[+15.994915]ANK | 2 | heavy | 967.932 | y6 | 673.3395 | 51.23 |
| P17967 | PDI | QSQPAVAVVADLPAYLAN[+203.079373]ETFVTPVIVQSGK | 3 | light | 1139.271 | y12 | 1275.731 | 99.45 |
| P17967 | PDI | QSQPAVAVVADLPAYLAN[+203.079373]ETFVTPVIVQSGK | 3 | light | 1139.271 | y11 | 1174.683 | 99.45 |
| P17967 | PDI | QSQPAVAVVADLPAYLAN[+203.079373]ETFVTPVIVQSGK | 3 | light | 1139.271 | y10 | 1027.615 | 99.45 |
| P17967 | PDI | QSQPAVAVVADLPAYLAN[+203.079373]ETFVTPVIVQSGK | 3 | light | 1139.271 | y9 | 928.5462 | 99.45 |
| P17967 | PDI | QSQPAVAVVADLPAYLAN[+203.079373]ETFVTPVIVQSGK | 3 | light | 1139.271 | y8 | 827.4985 | 99.45 |
| P17967 | PDI | QSQPAVAVVADLPAYLAN[+203.079373]ETFVTPVIVQSGK | 3 | light | 1139.271 | y5 | 518.2933 | 99.45 |
| P17967 | PDI | QSQPAVAVVADLPAYLAN[+203.079373]ETFVTPVIVQSGK | 3 | light | 1139.271 | y4 | 419.2249 | 99.45 |

| | | | | | | | | |
|---|---|---|---|---|---|---|---|---|
| P17967 | PDI | QSQPAVAVVADLPAYLAN[+203.079373]ETFVTPVIVQSGK | 3 | heavy | 1141.943 | y12 | 1283.745 | 99.45 |
| P17967 | PDI | QSQPAVAVVADLPAYLAN[+203.079373]ETFVTPVIVQSGK | 3 | heavy | 1141.943 | y11 | 1182.697 | 99.45 |
| P17967 | PDI | QSQPAVAVVADLPAYLAN[+203.079373]ETFVTPVIVQSGK | 3 | heavy | 1141.943 | y10 | 1035.629 | 99.45 |
| P17967 | PDI | QSQPAVAVVADLPAYLAN[+203.079373]ETFVTPVIVQSGK | 3 | heavy | 1141.943 | y9 | 936.5604 | 99.45 |
| P17967 | PDI | QSQPAVAVVADLPAYLAN[+203.079373]ETFVTPVIVQSGK | 3 | heavy | 1141.943 | y8 | 835.5127 | 99.45 |
| P17967 | PDI | QSQPAVAVVADLPAYLAN[+203.079373]ETFVTPVIVQSGK | 3 | heavy | 1141.943 | y5 | 526.3075 | 99.45 |
| P17967 | PDI | QSQPAVAVVADLPAYLAN[+203.079373]ETFVTPVIVQSGK | 3 | heavy | 1141.943 | y4 | 427.2391 | 99.45 |
| P17967 | PDI | N[+203.079373]ITLAQIDC[+57.021464]TENQDLC[+57.021464]M[+15.994915]EHNIPGFPSLK | 3 | light | 1159.871 | y9 | 972.5513 | 80.16 |
| P17967 | PDI | N[+203.079373]ITLAQIDC[+57.021464]TENQDLC[+57.021464]M[+15.994915]EHNIPGFPSLK | 3 | light | 1159.871 | y8 | 858.5084 | 80.16 |
| P17967 | PDI | N[+203.079373]ITLAQIDC[+57.021464]TENQDLC[+57.021464]M[+15.994915]EHNIPGFPSLK | 3 | light | 1159.871 | y7 | 745.4243 | 80.16 |
| P17967 | PDI | N[+203.079373]ITLAQIDC[+57.021464]TENQDLC[+57.021464]M[+15.994915]EHNIPGFPSLK | 3 | light | 1159.871 | y4 | 444.2817 | 80.16 |
| P17967 | PDI | N[+203.079373]ITLAQIDC[+57.021464]TENQDLC[+57.021464]M[+15.994915]EHNIPGFPSLK | 3 | light | 1159.871 | b2 | 228.1343 | 80.16 |
| P17967 | PDI | N[+203.079373]ITLAQIDC[+57.021464]TENQDLC[+57.021464]M[+15.994915]EHNIPGFPSLK | 3 | heavy | 1162.542 | y9 | 980.5655 | 80.16 |
| P17967 | PDI | N[+203.079373]ITLAQIDC[+57.021464]TENQDLC[+57.021464]M[+15.994915]EHNIPGFPSLK | 3 | heavy | 1162.542 | y8 | 866.5226 | 80.16 |
| P17967 | PDI | N[+203.079373]ITLAQIDC[+57.021464]TENQDLC[+57.021464]M[+15.994915]EHNIPGFPSLK | 3 | heavy | 1162.542 | y7 | 753.4385 | 80.16 |
| P17967 | PDI | N[+203.079373]ITLAQIDC[+57.021464]TENQDLC[+57.021464]M[+15.994915]EHNIPGFPSLK | 3 | heavy | 1162.542 | y4 | 452.2959 | 80.16 |
| P17967 | PDI | N[+203.079373]ITLAQIDC[+57.021464]TENQDLC[+57.021464]M[+15.994915]EHNIPGFPSLK | 3 | heavy | 1162.542 | b2 | 228.1343 | 80.16 |
| P17967 | PDI | TAEAIVQFMIK | 2 | light | 625.8443 | y9 | 1078.597 | 69.05 |
| P17967 | PDI | TAEAIVQFMIK | 2 | light | 625.8443 | y8 | 949.5539 | 69.05 |
| P17967 | PDI | TAEAIVQFMIK | 2 | light | 625.8443 | y7 | 878.5168 | 69.05 |
| P17967 | PDI | TAEAIVQFMIK | 2 | light | 625.8443 | y6 | 765.4328 | 69.05 |
| P17967 | PDI | TAEAIVQFMIK | 2 | light | 625.8443 | y4 | 538.3058 | 69.05 |
| P17967 | PDI | TAEAIVQFMIK | 2 | heavy | 629.8514 | y9 | 1086.611 | 69.05 |
| P17967 | PDI | TAEAIVQFMIK | 2 | heavy | 629.8514 | y8 | 957.5681 | 69.05 |
| P17967 | PDI | TAEAIVQFMIK | 2 | heavy | 629.8514 | y7 | 886.531 | 69.05 |
| P17967 | PDI | TAEAIVQFMIK | 2 | heavy | 629.8514 | y6 | 773.447 | 69.05 |
| P17967 | PDI | TAEAIVQFMIK | 2 | heavy | 629.8514 | y4 | 546.32 | 69.05 |

| | | | | | | | | |
|---|---|---|---|---|---|---|---|---|
| P17967 | PDI | GLMNFVSIDAR | 2 | light | 611.8161 | y9 | 1052.519 | 65.66 |
| P17967 | PDI | GLMNFVSIDAR | 2 | light | 611.8161 | y7 | 807.4359 | 65.66 |
| P17967 | PDI | GLMNFVSIDAR | 2 | light | 611.8161 | y6 | 660.3675 | 65.66 |
| P17967 | PDI | GLMNFVSIDAR | 2 | light | 611.8161 | y5 | 561.2991 | 65.66 |
| P17967 | PDI | GLMNFVSIDAR | 2 | light | 611.8161 | y3 | 361.183 | 65.66 |
| P17967 | PDI | GLMNFVSIDAR | 2 | light | 611.8161 | y2 | 246.1561 | 65.66 |
| P17967 | PDI | GLMNFVSIDAR | 2 | heavy | 614.8261 | y9 | 1058.539 | 65.66 |
| P17967 | PDI | GLMNFVSIDAR | 2 | heavy | 614.8261 | y7 | 813.4561 | 65.66 |
| P17967 | PDI | GLMNFVSIDAR | 2 | heavy | 614.8261 | y6 | 666.3876 | 65.66 |
| P17967 | PDI | GLMNFVSIDAR | 2 | heavy | 614.8261 | y5 | 567.3192 | 65.66 |
| P17967 | PDI | GLMNFVSIDAR | 2 | heavy | 614.8261 | y3 | 367.2031 | 65.66 |
| P17967 | PDI | GLMNFVSIDAR | 2 | heavy | 614.8261 | y2 | 252.1762 | 65.66 |
| P17967 | PDI | LAPTYQELADTYAN[+203.079373]ATSDVLIAK | 3 | light | 891.1165 | y11 | 1305.69 | 74.33 |
| P17967 | PDI | LAPTYQELADTYAN[+203.079373]ATSDVLIAK | 3 | light | 891.1165 | y11 | 1102.61 | 74.33 |
| P17967 | PDI | LAPTYQELADTYAN[+203.079373]ATSDVLIAK | 3 | light | 891.1165 | y10 | 1234.653 | 74.33 |
| P17967 | PDI | LAPTYQELADTYAN[+203.079373]ATSDVLIAK | 3 | light | 891.1165 | y10 | 1031.573 | 74.33 |
| P17967 | PDI | LAPTYQELADTYAN[+203.079373]ATSDVLIAK | 3 | light | 891.1165 | y9 | 917.5302 | 74.33 |
| P17967 | PDI | LAPTYQELADTYAN[+203.079373]ATSDVLIAK | 3 | light | 891.1165 | y8 | 846.4931 | 74.33 |
| P17967 | PDI | LAPTYQELADTYAN[+203.079373]ATSDVLIAK | 3 | light | 891.1165 | y7 | 745.4454 | 74.33 |
| P17967 | PDI | LAPTYQELADTYAN[+203.079373]ATSDVLIAK | 3 | light | 891.1165 | y4 | 444.318 | 74.33 |
| P17967 | PDI | LAPTYQELADTYAN[+203.079373]ATSDVLIAK | 3 | light | 891.1165 | y3 | 331.234 | 74.33 |
| P17967 | PDI | LAPTYQELADTYAN[+203.079373]ATSDVLIAK | 3 | light | 891.1165 | y2 | 218.1499 | 74.33 |
| P17967 | PDI | LAPTYQELADTYAN[+203.079373]ATSDVLIAK | 3 | light | 891.1165 | b7 | 803.3934 | 74.33 |
| P17967 | PDI | LAPTYQELADTYAN[+203.079373]ATSDVLIAK | 3 | heavy | 893.7879 | y11 | 1313.704 | 74.33 |
| P17967 | PDI | LAPTYQELADTYAN[+203.079373]ATSDVLIAK | 3 | heavy | 893.7879 | y11 | 1110.624 | 74.33 |
| P17967 | PDI | LAPTYQELADTYAN[+203.079373]ATSDVLIAK | 3 | heavy | 893.7879 | y10 | 1242.667 | 74.33 |
| P17967 | PDI | LAPTYQELADTYAN[+203.079373]ATSDVLIAK | 3 | heavy | 893.7879 | y10 | 1039.587 | 74.33 |
| P17967 | PDI | LAPTYQELADTYAN[+203.079373]ATSDVLIAK | 3 | heavy | 893.7879 | y9 | 925.5444 | 74.33 |
| P17967 | PDI | LAPTYQELADTYAN[+203.079373]ATSDVLIAK | 3 | heavy | 893.7879 | y8 | 854.5073 | 74.33 |
| P17967 | PDI | LAPTYQELADTYAN[+203.079373]ATSDVLIAK | 3 | heavy | 893.7879 | y7 | 753.4596 | 74.33 |
| P17967 | PDI | LAPTYQELADTYAN[+203.079373]ATSDVLIAK | 3 | heavy | 893.7879 | y4 | 452.3322 | 74.33 |
| P17967 | PDI | LAPTYQELADTYAN[+203.079373]ATSDVLIAK | 3 | heavy | 893.7879 | y3 | 339.2482 | 74.33 |

| | | | | | | | | |
|---|---|---|---|---|---|---|---|---|
| P17967 | PDI | LAPTYQELADTYAN[+203.079373]ATSDVLIAK | 3 | heavy | 893.7879 | y2 | 226.1641 | 74.33 |
| P17967 | PDI | LAPTYQELADTYAN[+203.079373]ATSDVLIAK | 3 | heavy | 893.7879 | b7 | 803.3934 | 74.33 |
| Q03691 | ROT1 | YN[+203.079373]QTETFK | 2 | light | 617.2853 | y7 | 1070.5 | 21.88 |
| Q03691 | ROT1 | YN[+203.079373]QTETFK | 2 | light | 617.2853 | y7 | 867.4207 | 21.88 |
| Q03691 | ROT1 | YN[+203.079373]QTETFK | 2 | light | 617.2853 | y6 | 753.3777 | 21.88 |
| Q03691 | ROT1 | YN[+203.079373]QTETFK | 2 | light | 617.2853 | y5 | 625.3192 | 21.88 |
| Q03691 | ROT1 | YN[+203.079373]QTETFK | 2 | light | 617.2853 | y4 | 524.2715 | 21.88 |
| Q03691 | ROT1 | YN[+203.079373]QTETFK | 2 | light | 617.2853 | y3 | 395.2289 | 21.88 |
| Q03691 | ROT1 | YN[+203.079373]QTETFK | 2 | light | 617.2853 | b2 | 481.1929 | 21.88 |
| Q03691 | ROT1 | YN[+203.079373]QTETFK | 2 | light | 617.2853 | b2 | 278.1135 | 21.88 |
| Q03691 | ROT1 | YN[+203.079373]QTETFK | 2 | light | 617.2853 | b4 | 507.2198 | 21.88 |
| Q03691 | ROT1 | YN[+203.079373]QTETFK | 2 | heavy | 621.2924 | y7 | 1078.514 | 21.88 |
| Q03691 | ROT1 | YN[+203.079373]QTETFK | 2 | heavy | 621.2924 | y7 | 875.4349 | 21.88 |
| Q03691 | ROT1 | YN[+203.079373]QTETFK | 2 | heavy | 621.2924 | y6 | 761.3919 | 21.88 |
| Q03691 | ROT1 | YN[+203.079373]QTETFK | 2 | heavy | 621.2924 | y5 | 633.3334 | 21.88 |
| Q03691 | ROT1 | YN[+203.079373]QTETFK | 2 | heavy | 621.2924 | y4 | 532.2857 | 21.88 |
| Q03691 | ROT1 | YN[+203.079373]QTETFK | 2 | heavy | 621.2924 | y3 | 403.2431 | 21.88 |
| Q03691 | ROT1 | YN[+203.079373]QTETFK | 2 | heavy | 621.2924 | b2 | 481.1929 | 21.88 |
| Q03691 | ROT1 | YN[+203.079373]QTETFK | 2 | heavy | 621.2924 | b2 | 278.1135 | 21.88 |
| Q03691 | ROT1 | YN[+203.079373]QTETFK | 2 | heavy | 621.2924 | b4 | 507.2198 | 21.88 |
| Q03691 | ROT1 | EDESNSIYGTWSSK | 2 | light | 801.8495 | y11 | 1229.58 | 43.73 |
| Q03691 | ROT1 | EDESNSIYGTWSSK | 2 | light | 801.8495 | y9 | 1028.505 | 43.73 |
| Q03691 | ROT1 | EDESNSIYGTWSSK | 2 | light | 801.8495 | y8 | 941.4727 | 43.73 |
| Q03691 | ROT1 | EDESNSIYGTWSSK | 2 | light | 801.8495 | y7 | 828.3886 | 43.73 |
| Q03691 | ROT1 | EDESNSIYGTWSSK | 2 | light | 801.8495 | y6 | 665.3253 | 43.73 |
| Q03691 | ROT1 | EDESNSIYGTWSSK | 2 | light | 801.8495 | y4 | 507.2562 | 43.73 |
| Q03691 | ROT1 | EDESNSIYGTWSSK | 2 | light | 801.8495 | y3 | 321.1769 | 43.73 |
| Q03691 | ROT1 | EDESNSIYGTWSSK | 2 | heavy | 805.8566 | y11 | 1237.594 | 43.73 |
| Q03691 | ROT1 | EDESNSIYGTWSSK | 2 | heavy | 805.8566 | y9 | 1036.519 | 43.73 |
| Q03691 | ROT1 | EDESNSIYGTWSSK | 2 | heavy | 805.8566 | y8 | 949.4869 | 43.73 |
| Q03691 | ROT1 | EDESNSIYGTWSSK | 2 | heavy | 805.8566 | y7 | 836.4028 | 43.73 |
| Q03691 | ROT1 | EDESNSIYGTWSSK | 2 | heavy | 805.8566 | y6 | 673.3395 | 43.73 |

| | | | | | | | | |
|---|---|---|---|---|---|---|---|---|
| Q03691 | ROT1 | EDESNSIYGTWSSK | 2 | heavy | 805.8566 | y4 | 515.2704 | 43.73 |
| Q03691 | ROT1 | EDESNSIYGTWSSK | 2 | heavy | 805.8566 | y3 | 329.1911 | 43.73 |
| Q03691 | ROT1 | QLFSDPC[+57.021464]NDDGVSTYSR | 2 | light | 980.9207 | y9 | 999.4378 | 46.58 |
| Q03691 | ROT1 | QLFSDPC[+57.021464]NDDGVSTYSR | 2 | light | 980.9207 | y8 | 884.4108 | 46.58 |
| Q03691 | ROT1 | QLFSDPC[+57.021464]NDDGVSTYSR | 2 | light | 980.9207 | y7 | 769.3839 | 46.58 |
| Q03691 | ROT1 | QLFSDPC[+57.021464]NDDGVSTYSR | 2 | light | 980.9207 | y5 | 613.294 | 46.58 |
| Q03691 | ROT1 | QLFSDPC[+57.021464]NDDGVSTYSR | 2 | light | 980.9207 | y3 | 425.2143 | 46.58 |
| Q03691 | ROT1 | QLFSDPC[+57.021464]NDDGVSTYSR | 2 | heavy | 983.9308 | y9 | 1005.458 | 46.58 |
| Q03691 | ROT1 | QLFSDPC[+57.021464]NDDGVSTYSR | 2 | heavy | 983.9308 | y8 | 890.431 | 46.58 |
| Q03691 | ROT1 | QLFSDPC[+57.021464]NDDGVSTYSR | 2 | heavy | 983.9308 | y7 | 775.404 | 46.58 |
| Q03691 | ROT1 | QLFSDPC[+57.021464]NDDGVSTYSR | 2 | heavy | 983.9308 | y5 | 619.3141 | 46.58 |
| Q03691 | ROT1 | QLFSDPC[+57.021464]NDDGVSTYSR | 2 | heavy | 983.9308 | y3 | 431.2344 | 46.58 |
| P40533 | TED1 | DNYWIEYETN[+203.079373]TTHPWR | 3 | light | 776.6783 | y8 | 1215.575 | 57.73 |
| P40533 | TED1 | DNYWIEYETN[+203.079373]TTHPWR | 3 | light | 776.6783 | y7 | 1114.528 | 57.73 |
| P40533 | TED1 | DNYWIEYETN[+203.079373]TTHPWR | 3 | light | 776.6783 | y7 | 911.4482 | 57.73 |
| P40533 | TED1 | DNYWIEYETN[+203.079373]TTHPWR | 3 | light | 776.6783 | y6 | 797.4053 | 57.73 |
| P40533 | TED1 | DNYWIEYETN[+203.079373]TTHPWR | 3 | light | 776.6783 | y3 | 458.251 | 57.73 |
| P40533 | TED1 | DNYWIEYETN[+203.079373]TTHPWR | 3 | light | 776.6783 | b2 | 230.0771 | 57.73 |
| P40533 | TED1 | DNYWIEYETN[+203.079373]TTHPWR | 3 | light | 776.6783 | b3 | 393.1405 | 57.73 |
| P40533 | TED1 | DNYWIEYETN[+203.079373]TTHPWR | 3 | heavy | 778.685 | y8 | 1221.595 | 57.73 |
| P40533 | TED1 | DNYWIEYETN[+203.079373]TTHPWR | 3 | heavy | 778.685 | y7 | 1120.548 | 57.73 |
| P40533 | TED1 | DNYWIEYETN[+203.079373]TTHPWR | 3 | heavy | 778.685 | y7 | 917.4684 | 57.73 |
| P40533 | TED1 | DNYWIEYETN[+203.079373]TTHPWR | 3 | heavy | 778.685 | y6 | 803.4254 | 57.73 |
| P40533 | TED1 | DNYWIEYETN[+203.079373]TTHPWR | 3 | heavy | 778.685 | y3 | 464.2712 | 57.73 |
| P40533 | TED1 | DNYWIEYETN[+203.079373]TTHPWR | 3 | heavy | 778.685 | b2 | 230.0771 | 57.73 |
| P40533 | TED1 | DNYWIEYETN[+203.079373]TTHPWR | 3 | heavy | 778.685 | b3 | 393.1405 | 57.73 |
| P40533 | TED1 | NIESDVFVK | 2 | light | 525.7769 | y7 | 823.4196 | 44.91 |
| P40533 | TED1 | NIESDVFVK | 2 | light | 525.7769 | y6 | 694.377 | 44.91 |
| P40533 | TED1 | NIESDVFVK | 2 | light | 525.7769 | y5 | 607.345 | 44.91 |
| P40533 | TED1 | NIESDVFVK | 2 | light | 525.7769 | y4 | 492.318 | 44.91 |
| P40533 | TED1 | NIESDVFVK | 2 | light | 525.7769 | y3 | 393.2496 | 44.91 |
| P40533 | TED1 | NIESDVFVK | 2 | light | 525.7769 | y5 | 304.1761 | 44.91 |

| | | | | | | | | |
|---|---|---|---|---|---|---|---|---|
| P40533 | TED1 | NIESDVFVK | 2 | heavy | 529.784 | y7 | 831.4338 | 44.91 |
| P40533 | TED1 | NIESDVFVK | 2 | heavy | 529.784 | y6 | 702.3912 | 44.91 |
| P40533 | TED1 | NIESDVFVK | 2 | heavy | 529.784 | y5 | 615.3592 | 44.91 |
| P40533 | TED1 | NIESDVFVK | 2 | heavy | 529.784 | y4 | 500.3322 | 44.91 |
| P40533 | TED1 | NIESDVFVK | 2 | heavy | 529.784 | y3 | 401.2638 | 44.91 |
| P40533 | TED1 | NIESDVFVK | 2 | heavy | 529.784 | y5 | 308.1832 | 44.91 |
| P40533 | TED1 | EGLC[+57.021464]VDGPDTR | 2 | light | 609.7746 | y8 | 919.3938 | 32.48 |
| P40533 | TED1 | EGLC[+57.021464]VDGPDTR | 2 | light | 609.7746 | y7 | 759.3632 | 32.48 |
| P40533 | TED1 | EGLC[+57.021464]VDGPDTR | 2 | light | 609.7746 | y6 | 660.2947 | 32.48 |
| P40533 | TED1 | EGLC[+57.021464]VDGPDTR | 2 | light | 609.7746 | y5 | 545.2678 | 32.48 |
| P40533 | TED1 | EGLC[+57.021464]VDGPDTR | 2 | light | 609.7746 | y4 | 488.2463 | 32.48 |
| P40533 | TED1 | EGLC[+57.021464]VDGPDTR | 2 | light | 609.7746 | y3 | 391.1936 | 32.48 |
| P40533 | TED1 | EGLC[+57.021464]VDGPDTR | 2 | heavy | 612.7847 | y8 | 925.4139 | 32.48 |
| P40533 | TED1 | EGLC[+57.021464]VDGPDTR | 2 | heavy | 612.7847 | y7 | 765.3833 | 32.48 |
| P40533 | TED1 | EGLC[+57.021464]VDGPDTR | 2 | heavy | 612.7847 | y6 | 666.3149 | 32.48 |
| P40533 | TED1 | EGLC[+57.021464]VDGPDTR | 2 | heavy | 612.7847 | y5 | 551.2879 | 32.48 |
| P40533 | TED1 | EGLC[+57.021464]VDGPDTR | 2 | heavy | 612.7847 | y4 | 494.2665 | 32.48 |
| P40533 | TED1 | EGLC[+57.021464]VDGPDTR | 2 | heavy | 612.7847 | y3 | 397.2137 | 32.48 |
| P43611 | OSW7 | NLDDLN[+203.079373]TTVNEQLVFLDSK | 2 | light | 1191.09 | y10 | 1192.621 | 70.74 |
| P43611 | OSW7 | NLDDLN[+203.079373]TTVNEQLVFLDSK | 2 | light | 1191.09 | y9 | 1078.578 | 70.74 |
| P43611 | OSW7 | NLDDLN[+203.079373]TTVNEQLVFLDSK | 2 | light | 1191.09 | y8 | 949.5353 | 70.74 |
| P43611 | OSW7 | NLDDLN[+203.079373]TTVNEQLVFLDSK | 2 | light | 1191.09 | y7 | 821.4767 | 70.74 |
| P43611 | OSW7 | NLDDLN[+203.079373]TTVNEQLVFLDSK | 2 | light | 1191.09 | y6 | 708.3927 | 70.74 |
| P43611 | OSW7 | NLDDLN[+203.079373]TTVNEQLVFLDSK | 2 | light | 1191.09 | y5 | 609.3243 | 70.74 |
| P43611 | OSW7 | NLDDLN[+203.079373]TTVNEQLVFLDSK | 2 | light | 1191.09 | y4 | 462.2558 | 70.74 |
| P43611 | OSW7 | NLDDLN[+203.079373]TTVNEQLVFLDSK | 2 | light | 1191.09 | y3 | 349.1718 | 70.74 |
| P43611 | OSW7 | NLDDLN[+203.079373]TTVNEQLVFLDSK | 2 | light | 1191.09 | y2 | 234.1448 | 70.74 |
| P43611 | OSW7 | NLDDLN[+203.079373]TTVNEQLVFLDSK | 2 | heavy | 1195.097 | y10 | 1200.635 | 70.74 |
| P43611 | OSW7 | NLDDLN[+203.079373]TTVNEQLVFLDSK | 2 | heavy | 1195.097 | y9 | 1086.592 | 70.74 |
| P43611 | OSW7 | NLDDLN[+203.079373]TTVNEQLVFLDSK | 2 | heavy | 1195.097 | y8 | 957.5495 | 70.74 |
| P43611 | OSW7 | NLDDLN[+203.079373]TTVNEQLVFLDSK | 2 | heavy | 1195.097 | y7 | 829.4909 | 70.74 |
| P43611 | OSW7 | NLDDLN[+203.079373]TTVNEQLVFLDSK | 2 | heavy | 1195.097 | y6 | 716.4069 | 70.74 |

| | | | | | | | | |
|---|---|---|---|---|---|---|---|---|
| P43611 | OSW7 | NLDDLN[+203.079373]TTVNEQLV FLDSK | 2 | heavy | 1195.097 | y5 | 617.3385 | 70.74 |
| P43611 | OSW7 | NLDDLN[+203.079373]TTVNEQLV FLDSK | 2 | heavy | 1195.097 | y4 | 470.27 | 70.74 |
| P43611 | OSW7 | NLDDLN[+203.079373]TTVNEQLV FLDSK | 2 | heavy | 1195.097 | y3 | 357.186 | 70.74 |
| P43611 | OSW7 | NLDDLN[+203.079373]TTVNEQLV FLDSK | 2 | heavy | 1195.097 | y2 | 242.159 | 70.74 |
| P43611 | OSW7 | SSITSILK | 2 | light | 424.758 | y7 | 761.4767 | 45.69 |
| P43611 | OSW7 | SSITSILK | 2 | light | 424.758 | y6 | 674.4447 | 45.69 |
| P43611 | OSW7 | SSITSILK | 2 | light | 424.758 | y5 | 561.3606 | 45.69 |
| P43611 | OSW7 | SSITSILK | 2 | light | 424.758 | y4 | 460.313 | 45.69 |
| P43611 | OSW7 | SSITSILK | 2 | light | 424.758 | y3 | 373.2809 | 45.69 |
| P43611 | OSW7 | SSITSILK | 2 | light | 424.758 | b2 | 175.0713 | 45.69 |
| P43611 | OSW7 | SSITSILK | 2 | heavy | 428.7651 | y7 | 769.4909 | 45.69 |
| P43611 | OSW7 | SSITSILK | 2 | heavy | 428.7651 | y6 | 682.4589 | 45.69 |
| P43611 | OSW7 | SSITSILK | 2 | heavy | 428.7651 | y5 | 569.3748 | 45.69 |
| P43611 | OSW7 | SSITSILK | 2 | heavy | 428.7651 | y4 | 468.3272 | 45.69 |
| P43611 | OSW7 | SSITSILK | 2 | heavy | 428.7651 | y3 | 381.2951 | 45.69 |
| P43611 | OSW7 | SSITSILK | 2 | heavy | 428.7651 | b2 | 175.0713 | 45.69 |
| P36051 | MCD4 | HLDQLFHN[+203.079373]STLN[+203.079373]STLDYEIR | 4 | light | 706.3424 | y8 | 996.4997 | 56.31 |
| P36051 | MCD4 | HLDQLFHN[+203.079373]STLN[+203.079373]STLDYEIR | 4 | light | 706.3424 | y7 | 909.4676 | 56.31 |
| P36051 | MCD4 | HLDQLFHN[+203.079373]STLN[+203.079373]STLDYEIR | 4 | light | 706.3424 | y6 | 808.4199 | 56.31 |
| P36051 | MCD4 | HLDQLFHN[+203.079373]STLN[+203.079373]STLDYEIR | 4 | light | 706.3424 | y5 | 695.3359 | 56.31 |
| P36051 | MCD4 | HLDQLFHN[+203.079373]STLN[+203.079373]STLDYEIR | 4 | light | 706.3424 | y4 | 580.3089 | 56.31 |
| P36051 | MCD4 | HLDQLFHN[+203.079373]STLN[+203.079373]STLDYEIR | 4 | light | 706.3424 | y2 | 288.203 | 56.31 |
| P36051 | MCD4 | HLDQLFHN[+203.079373]STLN[+203.079373]STLDYEIR | 4 | light | 706.3424 | y12 | 706.3568 | 56.31 |
| P36051 | MCD4 | HLDQLFHN[+203.079373]STLN[+203.079373]STLDYEIR | 4 | heavy | 707.8475 | y8 | 1002.52 | 56.31 |
| P36051 | MCD4 | HLDQLFHN[+203.079373]STLN[+203.079373]STLDYEIR | 4 | heavy | 707.8475 | y7 | 915.4878 | 56.31 |
| P36051 | MCD4 | HLDQLFHN[+203.079373]STLN[+203.079373]STLDYEIR | 4 | heavy | 707.8475 | y6 | 814.4401 | 56.31 |
| P36051 | MCD4 | HLDQLFHN[+203.079373]STLN[+203.079373]STLDYEIR | 4 | heavy | 707.8475 | y5 | 701.356 | 56.31 |
| P36051 | MCD4 | HLDQLFHN[+203.079373]STLN[+203.079373]STLDYEIR | 4 | heavy | 707.8475 | y4 | 586.3291 | 56.31 |
| P36051 | MCD4 | HLDQLFHN[+203.079373]STLN[+203.079373]STLDYEIR | 4 | heavy | 707.8475 | y2 | 294.2231 | 56.31 |
| P36051 | MCD4 | HLDQLFHN[+203.079373]STLN[+203.079373]STLDYEIR | 4 | heavy | 707.8475 | y12 | 709.3669 | 56.31 |
| P36051 | MCD4 | TEFLAPFIR | 2 | light | 547.3057 | y7 | 863.5138 | 70.44 |
| P36051 | MCD4 | TEFLAPFIR | 2 | light | 547.3057 | y6 | 716.4454 | 70.44 |

| | | | | | | | | |
|---|---|---|---|---|---|---|---|---|
| P36051 | MCD4 | TEFLAPFIR | 2 | light | 547.3057 | y5 | 603.3613 | 70.44 |
| P36051 | MCD4 | TEFLAPFIR | 2 | light | 547.3057 | y4 | 532.3242 | 70.44 |
| P36051 | MCD4 | TEFLAPFIR | 2 | light | 547.3057 | b2 | 231.0975 | 70.44 |
| P36051 | MCD4 | TEFLAPFIR | 2 | light | 547.3057 | b3 | 378.166 | 70.44 |
| P36051 | MCD4 | TEFLAPFIR | 2 | heavy | 550.3157 | y7 | 869.5339 | 70.44 |
| P36051 | MCD4 | TEFLAPFIR | 2 | heavy | 550.3157 | y6 | 722.4655 | 70.44 |
| P36051 | MCD4 | TEFLAPFIR | 2 | heavy | 550.3157 | y5 | 609.3814 | 70.44 |
| P36051 | MCD4 | TEFLAPFIR | 2 | heavy | 550.3157 | y4 | 538.3443 | 70.44 |
| P36051 | MCD4 | TEFLAPFIR | 2 | heavy | 550.3157 | b2 | 231.0975 | 70.44 |
| P36051 | MCD4 | TEFLAPFIR | 2 | heavy | 550.3157 | b3 | 378.166 | 70.44 |
| P36051 | MCD4 | YIDDQIPILIDK | 2 | light | 723.398 | y7 | 811.5288 | 67.84 |
| P36051 | MCD4 | YIDDQIPILIDK | 2 | light | 723.398 | y6 | 698.4447 | 67.84 |
| P36051 | MCD4 | YIDDQIPILIDK | 2 | light | 723.398 | y4 | 488.3079 | 67.84 |
| P36051 | MCD4 | YIDDQIPILIDK | 2 | light | 723.398 | y3 | 375.2238 | 67.84 |
| P36051 | MCD4 | YIDDQIPILIDK | 2 | light | 723.398 | y2 | 262.1397 | 67.84 |
| P36051 | MCD4 | YIDDQIPILIDK | 2 | light | 723.398 | b2 | 277.1547 | 67.84 |
| P36051 | MCD4 | YIDDQIPILIDK | 2 | heavy | 727.405 | y7 | 819.543 | 67.84 |
| P36051 | MCD4 | YIDDQIPILIDK | 2 | heavy | 727.405 | y6 | 706.4589 | 67.84 |
| P36051 | MCD4 | YIDDQIPILIDK | 2 | heavy | 727.405 | y4 | 496.3221 | 67.84 |
| P36051 | MCD4 | YIDDQIPILIDK | 2 | heavy | 727.405 | y3 | 383.238 | 67.84 |
| P36051 | MCD4 | YIDDQIPILIDK | 2 | heavy | 727.405 | y2 | 270.1539 | 67.84 |
| P36051 | MCD4 | YIDDQIPILIDK | 2 | heavy | 727.405 | b2 | 277.1547 | 67.84 |
| Q03281 | HEH2 | SN[+203.079373]NTNYIYR | 2 | light | 674.3124 | y7 | 943.4632 | 26.9 |
| Q03281 | HEH2 | SN[+203.079373]NTNYIYR | 2 | light | 674.3124 | y6 | 829.4203 | 26.9 |
| Q03281 | HEH2 | SN[+203.079373]NTNYIYR | 2 | light | 674.3124 | y5 | 728.3726 | 26.9 |
| Q03281 | HEH2 | SN[+203.079373]NTNYIYR | 2 | light | 674.3124 | y4 | 614.3297 | 26.9 |
| Q03281 | HEH2 | SN[+203.079373]NTNYIYR | 2 | light | 674.3124 | y3 | 451.2663 | 26.9 |
| Q03281 | HEH2 | SN[+203.079373]NTNYIYR | 2 | light | 674.3124 | y2 | 338.1823 | 26.9 |
| Q03281 | HEH2 | SN[+203.079373]NTNYIYR | 2 | heavy | 677.3225 | y7 | 949.4833 | 26.9 |
| Q03281 | HEH2 | SN[+203.079373]NTNYIYR | 2 | heavy | 677.3225 | y6 | 835.4404 | 26.9 |
| Q03281 | HEH2 | SN[+203.079373]NTNYIYR | 2 | heavy | 677.3225 | y5 | 734.3927 | 26.9 |
| Q03281 | HEH2 | SN[+203.079373]NTNYIYR | 2 | heavy | 677.3225 | y4 | 620.3498 | 26.9 |

| | | | | | | | | |
|---|---|---|---|---|---|---|---|---|
| Q03281 | HEH2 | SN[+203.079373]NTNYIYR | 2 | heavy | 677.3225 | y3 | 457.2865 | 26.9 |
| Q03281 | HEH2 | SN[+203.079373]NTNYIYR | 2 | heavy | 677.3225 | y2 | 344.2024 | 26.9 |
| Q03281 | HEH2 | ATLLSDIPNIK | 2 | light | 592.8479 | y9 | 1012.604 | 62.6 |
| Q03281 | HEH2 | ATLLSDIPNIK | 2 | light | 592.8479 | y8 | 899.5197 | 62.6 |
| Q03281 | HEH2 | ATLLSDIPNIK | 2 | light | 592.8479 | y7 | 786.4356 | 62.6 |
| Q03281 | HEH2 | ATLLSDIPNIK | 2 | light | 592.8479 | y6 | 699.4036 | 62.6 |
| Q03281 | HEH2 | ATLLSDIPNIK | 2 | light | 592.8479 | y5 | 584.3766 | 62.6 |
| Q03281 | HEH2 | ATLLSDIPNIK | 2 | light | 592.8479 | y4 | 471.2926 | 62.6 |
| Q03281 | HEH2 | ATLLSDIPNIK | 2 | light | 592.8479 | y3 | 374.2398 | 62.6 |
| Q03281 | HEH2 | ATLLSDIPNIK | 2 | heavy | 596.855 | y9 | 1020.618 | 62.6 |
| Q03281 | HEH2 | ATLLSDIPNIK | 2 | heavy | 596.855 | y8 | 907.5339 | 62.6 |
| Q03281 | HEH2 | ATLLSDIPNIK | 2 | heavy | 596.855 | y7 | 794.4498 | 62.6 |
| Q03281 | HEH2 | ATLLSDIPNIK | 2 | heavy | 596.855 | y6 | 707.4178 | 62.6 |
| Q03281 | HEH2 | ATLLSDIPNIK | 2 | heavy | 596.855 | y5 | 592.3908 | 62.6 |
| Q03281 | HEH2 | ATLLSDIPNIK | 2 | heavy | 596.855 | y4 | 479.3068 | 62.6 |
| Q03281 | HEH2 | ATLLSDIPNIK | 2 | heavy | 596.855 | y3 | 382.254 | 62.6 |
| Q03281 | HEH2 | SSILETYGIIPFPK | 2 | light | 782.9347 | y11 | 1277.714 | 80.69 |
| Q03281 | HEH2 | SSILETYGIIPFPK | 2 | light | 782.9347 | y10 | 1164.63 | 80.69 |
| Q03281 | HEH2 | SSILETYGIIPFPK | 2 | light | 782.9347 | y9 | 1035.587 | 80.69 |
| Q03281 | HEH2 | SSILETYGIIPFPK | 2 | light | 782.9347 | y8 | 934.5397 | 80.69 |
| Q03281 | HEH2 | SSILETYGIIPFPK | 2 | light | 782.9347 | y7 | 771.4763 | 80.69 |
| Q03281 | HEH2 | SSILETYGIIPFPK | 2 | light | 782.9347 | y5 | 601.3708 | 80.69 |
| Q03281 | HEH2 | SSILETYGIIPFPK | 2 | light | 782.9347 | y4 | 488.2867 | 80.69 |
| Q03281 | HEH2 | SSILETYGIIPFPK | 2 | light | 782.9347 | y3 | 391.234 | 80.69 |
| Q03281 | HEH2 | SSILETYGIIPFPK | 2 | heavy | 786.9418 | y11 | 1285.728 | 80.69 |
| Q03281 | HEH2 | SSILETYGIIPFPK | 2 | heavy | 786.9418 | y10 | 1172.644 | 80.69 |
| Q03281 | HEH2 | SSILETYGIIPFPK | 2 | heavy | 786.9418 | y9 | 1043.602 | 80.69 |
| Q03281 | HEH2 | SSILETYGIIPFPK | 2 | heavy | 786.9418 | y8 | 942.5539 | 80.69 |
| Q03281 | HEH2 | SSILETYGIIPFPK | 2 | heavy | 786.9418 | y7 | 779.4905 | 80.69 |
| Q03281 | HEH2 | SSILETYGIIPFPK | 2 | heavy | 786.9418 | y5 | 609.385 | 80.69 |
| Q03281 | HEH2 | SSILETYGIIPFPK | 2 | heavy | 786.9418 | y4 | 496.3009 | 80.69 |
| Q03281 | HEH2 | SSILETYGIIPFPK | 2 | heavy | 786.9418 | y3 | 399.2482 | 80.69 |

| | | | | | | | | |
|---|---|---|---|---|---|---|---|---|
| P38993 | FET3 | NGVNYAFFNN[+203.079373]ITY-TAPK | 2 | light | 1069.015 | y11 | 1315.668 | 68.06 |
| P38993 | FET3 | NGVNYAFFNN[+203.079373]ITY-TAPK | 2 | light | 1069.015 | y10 | 1371.679 | 68.06 |
| P38993 | FET3 | NGVNYAFFNN[+203.079373]ITY-TAPK | 2 | light | 1069.015 | y10 | 1168.6 | 68.06 |
| P38993 | FET3 | NGVNYAFFNN[+203.079373]ITY-TAPK | 2 | light | 1069.015 | y9 | 1224.611 | 68.06 |
| P38993 | FET3 | NGVNYAFFNN[+203.079373]ITY-TAPK | 2 | light | 1069.015 | y9 | 1021.531 | 68.06 |
| P38993 | FET3 | NGVNYAFFNN[+203.079373]ITY-TAPK | 2 | light | 1069.015 | y8 | 1110.568 | 68.06 |
| P38993 | FET3 | NGVNYAFFNN[+203.079373]ITY-TAPK | 2 | light | 1069.015 | y8 | 907.4884 | 68.06 |
| P38993 | FET3 | NGVNYAFFNN[+203.079373]ITY-TAPK | 2 | light | 1069.015 | y7 | 793.4454 | 68.06 |
| P38993 | FET3 | NGVNYAFFNN[+203.079373]ITY-TAPK | 2 | light | 1069.015 | y6 | 680.3614 | 68.06 |
| P38993 | FET3 | NGVNYAFFNN[+203.079373]ITY-TAPK | 2 | light | 1069.015 | y4 | 416.2504 | 68.06 |
| P38993 | FET3 | NGVNYAFFNN[+203.079373]ITY-TAPK | 2 | light | 1069.015 | y2 | 244.1656 | 68.06 |
| P38993 | FET3 | NGVNYAFFNN[+203.079373]ITY-TAPK | 2 | heavy | 1073.023 | y11 | 1323.682 | 68.06 |
| P38993 | FET3 | NGVNYAFFNN[+203.079373]ITY-TAPK | 2 | heavy | 1073.023 | y10 | 1379.693 | 68.06 |
| P38993 | FET3 | NGVNYAFFNN[+203.079373]ITY-TAPK | 2 | heavy | 1073.023 | y10 | 1176.614 | 68.06 |
| P38993 | FET3 | NGVNYAFFNN[+203.079373]ITY-TAPK | 2 | heavy | 1073.023 | y9 | 1232.625 | 68.06 |
| P38993 | FET3 | NGVNYAFFNN[+203.079373]ITY-TAPK | 2 | heavy | 1073.023 | y9 | 1029.545 | 68.06 |
| P38993 | FET3 | NGVNYAFFNN[+203.079373]ITY-TAPK | 2 | heavy | 1073.023 | y8 | 1118.582 | 68.06 |
| P38993 | FET3 | NGVNYAFFNN[+203.079373]ITY-TAPK | 2 | heavy | 1073.023 | y8 | 915.5026 | 68.06 |
| P38993 | FET3 | NGVNYAFFNN[+203.079373]ITY-TAPK | 2 | heavy | 1073.023 | y7 | 801.4596 | 68.06 |
| P38993 | FET3 | NGVNYAFFNN[+203.079373]ITY-TAPK | 2 | heavy | 1073.023 | y6 | 688.3756 | 68.06 |
| P38993 | FET3 | NGVNYAFFNN[+203.079373]ITY-TAPK | 2 | heavy | 1073.023 | y4 | 424.2646 | 68.06 |
| P38993 | FET3 | NGVNYAFFNN[+203.079373]ITY-TAPK | 2 | heavy | 1073.023 | y2 | 252.1798 | 68.06 |
| P38993 | FET3 | N[+203.079373]VTDM[+15.994915]LYITVAQR | 2 | light | 871.9351 | y11 | 1262.674 | 54.11 |
| P38993 | FET3 | N[+203.079373]VTDM[+15.994915]LYITVAQR | 2 | light | 871.9351 | y10 | 1225.625 | 54.11 |
| P38993 | FET3 | N[+203.079373]VTDM[+15.994915]LYITVAQR | 2 | light | 871.9351 | y10 | 1161.626 | 54.11 |
| P38993 | FET3 | N[+203.079373]VTDM[+15.994915]LYITVAQR | 2 | light | 871.9351 | y9 | 1110.598 | 54.11 |
| P38993 | FET3 | N[+203.079373]VTDM[+15.994915]LYITVAQR | 2 | light | 871.9351 | y9 | 1046.599 | 54.11 |
| P38993 | FET3 | N[+203.079373]VTDM[+15.994915]LYITVAQR | 2 | light | 871.9351 | y8 | 963.5622 | 54.11 |
| P38993 | FET3 | N[+203.079373]VTDM[+15.994915]LYITVAQR | 2 | light | 871.9351 | y7 | 850.4781 | 54.11 |
| P38993 | FET3 | N[+203.079373]VTDM[+15.994915]LYITVAQR | 2 | light | 871.9351 | y6 | 687.4148 | 54.11 |
| P38993 | FET3 | N[+203.079373]VTDM[+15.994915]LYITVAQR | 2 | light | 871.9351 | y5 | 574.3307 | 54.11 |
| P38993 | FET3 | N[+203.079373]VTDM[+15.994915]LYITVAQR | 2 | light | 871.9351 | y3 | 374.2146 | 54.11 |

| | | | | | | | | |
|---|---|---|---|---|---|---|---|---|
| P38993 | FET3 | N[+203.079373]VTDM[+15.994915]LYITVAQR | 2 | heavy | 874.9452 | y11 | 1268.694 | 54.11 |
| P38993 | FET3 | N[+203.079373]VTDM[+15.994915]LYITVAQR | 2 | heavy | 874.9452 | y10 | 1231.645 | 54.11 |
| P38993 | FET3 | N[+203.079373]VTDM[+15.994915]LYITVAQR | 2 | heavy | 874.9452 | y10 | 1167.646 | 54.11 |
| P38993 | FET3 | N[+203.079373]VTDM[+15.994915]LYITVAQR | 2 | heavy | 874.9452 | y9 | 1116.618 | 54.11 |
| P38993 | FET3 | N[+203.079373]VTDM[+15.994915]LYITVAQR | 2 | heavy | 874.9452 | y9 | 1052.619 | 54.11 |
| P38993 | FET3 | N[+203.079373]VTDM[+15.994915]LYITVAQR | 2 | heavy | 874.9452 | y8 | 969.5823 | 54.11 |
| P38993 | FET3 | N[+203.079373]VTDM[+15.994915]LYITVAQR | 2 | heavy | 874.9452 | y7 | 856.4983 | 54.11 |
| P38993 | FET3 | N[+203.079373]VTDM[+15.994915]LYITVAQR | 2 | heavy | 874.9452 | y6 | 693.4349 | 54.11 |
| P38993 | FET3 | N[+203.079373]VTDM[+15.994915]LYITVAQR | 2 | heavy | 874.9452 | y5 | 580.3509 | 54.11 |
| P38993 | FET3 | N[+203.079373]VTDM[+15.994915]LYITVAQR | 2 | heavy | 874.9452 | y3 | 380.2348 | 54.11 |
| P38993 | FET3 | DLHVDPEVLLNEVDENEER | 2 | light | 1132.537 | y5 | 676.2897 | 74.49 |
| P38993 | FET3 | DLHVDPEVLLNEVDENEER | 2 | light | 1132.537 | y2 | 304.1615 | 74.49 |
| P38993 | FET3 | DLHVDPEVLLNEVDENEER | 2 | light | 1132.537 | b3 | 366.1772 | 74.49 |
| P38993 | FET3 | DLHVDPEVLLNEVDENEER | 2 | light | 1132.537 | b4 | 465.2456 | 74.49 |
| P38993 | FET3 | DLHVDPEVLLNEVDENEER | 2 | light | 1132.537 | b5 | 580.2726 | 74.49 |
| P38993 | FET3 | DLHVDPEVLLNEVDENEER | 2 | heavy | 1135.548 | y5 | 682.3098 | 74.49 |
| P38993 | FET3 | DLHVDPEVLLNEVDENEER | 2 | heavy | 1135.548 | y2 | 310.1817 | 74.49 |
| P38993 | FET3 | DLHVDPEVLLNEVDENEER | 2 | heavy | 1135.548 | b3 | 366.1772 | 74.49 |
| P38993 | FET3 | DLHVDPEVLLNEVDENEER | 2 | heavy | 1135.548 | b4 | 465.2456 | 74.49 |
| P38993 | FET3 | DLHVDPEVLLNEVDENEER | 2 | heavy | 1135.548 | b5 | 580.2726 | 74.49 |
| P43561 | FET5 | YAFFNN[+203.079373]ITYVTPK | 2 | light | 890.9433 | y9 | 1049.563 | 63.27 |
| P43561 | FET5 | YAFFNN[+203.079373]ITYVTPK | 2 | light | 890.9433 | y8 | 935.5197 | 63.27 |
| P43561 | FET5 | YAFFNN[+203.079373]ITYVTPK | 2 | light | 890.9433 | y6 | 708.3927 | 63.27 |
| P43561 | FET5 | YAFFNN[+203.079373]ITYVTPK | 2 | light | 890.9433 | y5 | 607.345 | 63.27 |
| P43561 | FET5 | YAFFNN[+203.079373]ITYVTPK | 2 | light | 890.9433 | y3 | 345.2132 | 63.27 |
| P43561 | FET5 | YAFFNN[+203.079373]ITYVTPK | 2 | heavy | 894.9504 | y9 | 1057.577 | 63.27 |
| P43561 | FET5 | YAFFNN[+203.079373]ITYVTPK | 2 | heavy | 894.9504 | y8 | 943.5339 | 63.27 |
| P43561 | FET5 | YAFFNN[+203.079373]ITYVTPK | 2 | heavy | 894.9504 | y6 | 716.4069 | 63.27 |
| P43561 | FET5 | YAFFNN[+203.079373]ITYVTPK | 2 | heavy | 894.9504 | y5 | 615.3592 | 63.27 |
| P43561 | FET5 | YAFFNN[+203.079373]ITYVTPK | 2 | heavy | 894.9504 | y3 | 353.2274 | 63.27 |
| P43561 | FET5 | LN[+203.079373]YTASWV-TANPDGLHEK | 3 | light | 740.3587 | y11 | 1180.596 | 50.7 |
| P43561 | FET5 | LN[+203.079373]YTASWV-TANPDGLHEK | 3 | light | 740.3587 | y10 | 1081.527 | 50.7 |

| | | | | | | | | |
|---|---|---|---|---|---|---|---|---|
| P43561 | FET5 | LN[+203.079373]YTASWV-TANPDGLHEK | 3 | light | 740.3587 | y9 | 980.4796 | 50.7 |
| P43561 | FET5 | LN[+203.079373]YTASWV-TANPDGLHEK | 3 | light | 740.3587 | y8 | 909.4425 | 50.7 |
| P43561 | FET5 | LN[+203.079373]YTASWV-TANPDGLHEK | 3 | light | 740.3587 | y7 | 795.3995 | 50.7 |
| P43561 | FET5 | LN[+203.079373]YTASWV-TANPDGLHEK | 3 | light | 740.3587 | y5 | 583.3198 | 50.7 |
| P43561 | FET5 | LN[+203.079373]YTASWV-TANPDGLHEK | 3 | light | 740.3587 | y3 | 413.2143 | 50.7 |
| P43561 | FET5 | LN[+203.079373]YTASWV-TANPDGLHEK | 3 | light | 740.3587 | b3 | 391.1976 | 50.7 |
| P43561 | FET5 | LN[+203.079373]YTASWV-TANPDGLHEK | 3 | heavy | 743.0301 | y11 | 1188.61 | 50.7 |
| P43561 | FET5 | LN[+203.079373]YTASWV-TANPDGLHEK | 3 | heavy | 743.0301 | y10 | 1089.541 | 50.7 |
| P43561 | FET5 | LN[+203.079373]YTASWV-TANPDGLHEK | 3 | heavy | 743.0301 | y9 | 988.4938 | 50.7 |
| P43561 | FET5 | LN[+203.079373]YTASWV-TANPDGLHEK | 3 | heavy | 743.0301 | y8 | 917.4567 | 50.7 |
| P43561 | FET5 | LN[+203.079373]YTASWV-TANPDGLHEK | 3 | heavy | 743.0301 | y7 | 803.4137 | 50.7 |
| P43561 | FET5 | LN[+203.079373]YTASWV-TANPDGLHEK | 3 | heavy | 743.0301 | y5 | 591.334 | 50.7 |
| P43561 | FET5 | LN[+203.079373]YTASWV-TANPDGLHEK | 3 | heavy | 743.0301 | y3 | 421.2285 | 50.7 |
| P43561 | FET5 | LN[+203.079373]YTASWV-TANPDGLHEK | 3 | heavy | 743.0301 | b3 | 391.1976 | 50.7 |
| P43561 | FET5 | VPTLTTLLTSGK | 2 | light | 615.8688 | y10 | 1034.609 | 65.87 |
| P43561 | FET5 | VPTLTTLLTSGK | 2 | light | 615.8688 | y9 | 933.5615 | 65.87 |
| P43561 | FET5 | VPTLTTLLTSGK | 2 | light | 615.8688 | y8 | 820.4775 | 65.87 |
| P43561 | FET5 | VPTLTTLLTSGK | 2 | light | 615.8688 | y7 | 719.4298 | 65.87 |
| P43561 | FET5 | VPTLTTLLTSGK | 2 | light | 615.8688 | y5 | 505.298 | 65.87 |
| P43561 | FET5 | VPTLTTLLTSGK | 2 | light | 615.8688 | y4 | 392.214 | 65.87 |
| P43561 | FET5 | VPTLTTLLTSGK | 2 | heavy | 619.8759 | y10 | 1042.623 | 65.87 |
| P43561 | FET5 | VPTLTTLLTSGK | 2 | heavy | 619.8759 | y9 | 941.5757 | 65.87 |
| P43561 | FET5 | VPTLTTLLTSGK | 2 | heavy | 619.8759 | y8 | 828.4917 | 65.87 |
| P43561 | FET5 | VPTLTTLLTSGK | 2 | heavy | 619.8759 | y7 | 727.444 | 65.87 |
| P43561 | FET5 | VPTLTTLLTSGK | 2 | heavy | 619.8759 | y5 | 513.3122 | 65.87 |
| P43561 | FET5 | VPTLTTLLTSGK | 2 | heavy | 619.8759 | y4 | 400.2282 | 65.87 |
| P38843 | CHS7 | THLILSN[+203.079373]STI-IHDFDPLNLNVGVLPR | 3 | light | 1034.559 | y11 | 1191.721 | 81.84 |
| P38843 | CHS7 | THLILSN[+203.079373]STI-IHDFDPLNLNVGVLPR | 3 | light | 1034.559 | y10 | 1094.668 | 81.84 |
| P38843 | CHS7 | THLILSN[+203.079373]STI-IHDFDPLNLNVGVLPR | 3 | light | 1034.559 | y9 | 981.584 | 81.84 |
| P38843 | CHS7 | THLILSN[+203.079373]STI-IHDFDPLNLNVGVLPR | 3 | light | 1034.559 | y8 | 867.5411 | 81.84 |
| P38843 | CHS7 | THLILSN[+203.079373]STI-IHDFDPLNLNVGVLPR | 3 | light | 1034.559 | y7 | 754.457 | 81.84 |
| P38843 | CHS7 | THLILSN[+203.079373]STI-IHDFDPLNLNVGVLPR | 3 | light | 1034.559 | y6 | 640.4141 | 81.84 |

| | | | | | | | | |
|---|---|---|---|---|---|---|---|---|
| P38843 | CHS7 | THLILSN[+203.079373]STI-IHDFDPLNLNVGVLPR | 3 | light | 1034.559 | y5 | 541.3457 | 81.84 |
| P38843 | CHS7 | THLILSN[+203.079373]STI-IHDFDPLNLNVGVLPR | 3 | light | 1034.559 | b3 | 352.1979 | 81.84 |
| P38843 | CHS7 | THLILSN[+203.079373]STI-IHDFDPLNLNVGVLPR | 3 | light | 1034.559 | b4 | 465.282 | 81.84 |
| P38843 | CHS7 | THLILSN[+203.079373]STI-IHDFDPLNLNVGVLPR | 3 | heavy | 1036.566 | y11 | 1197.741 | 81.84 |
| P38843 | CHS7 | THLILSN[+203.079373]STI-IHDFDPLNLNVGVLPR | 3 | heavy | 1036.566 | y10 | 1100.688 | 81.84 |
| P38843 | CHS7 | THLILSN[+203.079373]STI-IHDFDPLNLNVGVLPR | 3 | heavy | 1036.566 | y9 | 987.6041 | 81.84 |
| P38843 | CHS7 | THLILSN[+203.079373]STI-IHDFDPLNLNVGVLPR | 3 | heavy | 1036.566 | y8 | 873.5612 | 81.84 |
| P38843 | CHS7 | THLILSN[+203.079373]STI-IHDFDPLNLNVGVLPR | 3 | heavy | 1036.566 | y7 | 760.4771 | 81.84 |
| P38843 | CHS7 | THLILSN[+203.079373]STI-IHDFDPLNLNVGVLPR | 3 | heavy | 1036.566 | y6 | 646.4342 | 81.84 |
| P38843 | CHS7 | THLILSN[+203.079373]STI-IHDFDPLNLNVGVLPR | 3 | heavy | 1036.566 | y5 | 547.3658 | 81.84 |
| P38843 | CHS7 | THLILSN[+203.079373]STI-IHDFDPLNLNVGVLPR | 3 | heavy | 1036.566 | b3 | 352.1979 | 81.84 |
| P38843 | CHS7 | THLILSN[+203.079373]STI-IHDFDPLNLNVGVLPR | 3 | heavy | 1036.566 | b4 | 465.282 | 81.84 |
| P38843 | CHS7 | TPLPLC[+57.021464]SVIK | 2 | light | 564.3283 | y8 | 929.5489 | 59.02 |
| P38843 | CHS7 | TPLPLC[+57.021464]SVIK | 2 | light | 564.3283 | y7 | 816.4648 | 59.02 |
| P38843 | CHS7 | TPLPLC[+57.021464]SVIK | 2 | light | 564.3283 | y5 | 606.328 | 59.02 |
| P38843 | CHS7 | TPLPLC[+57.021464]SVIK | 2 | light | 564.3283 | y4 | 446.2973 | 59.02 |
| P38843 | CHS7 | TPLPLC[+57.021464]SVIK | 2 | light | 564.3283 | b3 | 312.1918 | 59.02 |
| P38843 | CHS7 | TPLPLC[+57.021464]SVIK | 2 | heavy | 568.3354 | y8 | 937.5631 | 59.02 |
| P38843 | CHS7 | TPLPLC[+57.021464]SVIK | 2 | heavy | 568.3354 | y7 | 824.479 | 59.02 |
| P38843 | CHS7 | TPLPLC[+57.021464]SVIK | 2 | heavy | 568.3354 | y5 | 614.3422 | 59.02 |
| P38843 | CHS7 | TPLPLC[+57.021464]SVIK | 2 | heavy | 568.3354 | y4 | 454.3115 | 59.02 |
| P38843 | CHS7 | TPLPLC[+57.021464]SVIK | 2 | heavy | 568.3354 | b3 | 312.1918 | 59.02 |
| P27810 | KTR1 | N[+203.079373]VTSALVSGTTK | 2 | light | 690.8645 | y8 | 776.4512 | 34.11 |
| P27810 | KTR1 | N[+203.079373]VTSALVSGTTK | 2 | light | 690.8645 | y7 | 705.4141 | 34.11 |
| P27810 | KTR1 | N[+203.079373]VTSALVSGTTK | 2 | light | 690.8645 | y6 | 592.3301 | 34.11 |
| P27810 | KTR1 | N[+203.079373]VTSALVSGTTK | 2 | light | 690.8645 | y5 | 493.2617 | 34.11 |
| P27810 | KTR1 | N[+203.079373]VTSALVSGTTK | 2 | light | 690.8645 | y4 | 406.2296 | 34.11 |
| P27810 | KTR1 | N[+203.079373]VTSALVSGTTK | 2 | heavy | 694.8716 | y8 | 784.4654 | 34.11 |
| P27810 | KTR1 | N[+203.079373]VTSALVSGTTK | 2 | heavy | 694.8716 | y7 | 713.4283 | 34.11 |
| P27810 | KTR1 | N[+203.079373]VTSALVSGTTK | 2 | heavy | 694.8716 | y6 | 600.3443 | 34.11 |
| P27810 | KTR1 | N[+203.079373]VTSALVSGTTK | 2 | heavy | 694.8716 | y5 | 501.2759 | 34.11 |
| P27810 | KTR1 | N[+203.079373]VTSALVSGTTK | 2 | heavy | 694.8716 | y4 | 414.2438 | 34.11 |

| | | | | | | | | |
|---|---|---|---|---|---|---|---|---|
| P27810 | KTR1 | SPAYSAYFDYLDR | 2 | light | 784.3568 | y9 | 1149.521 | 65.34 |
| P27810 | KTR1 | SPAYSAYFDYLDR | 2 | light | 784.3568 | y8 | 1062.489 | 65.34 |
| P27810 | KTR1 | SPAYSAYFDYLDR | 2 | light | 784.3568 | y7 | 991.452 | 65.34 |
| P27810 | KTR1 | SPAYSAYFDYLDR | 2 | light | 784.3568 | y6 | 828.3886 | 65.34 |
| P27810 | KTR1 | SPAYSAYFDYLDR | 2 | light | 784.3568 | y5 | 681.3202 | 65.34 |
| P27810 | KTR1 | SPAYSAYFDYLDR | 2 | light | 784.3568 | y4 | 566.2933 | 65.34 |
| P27810 | KTR1 | SPAYSAYFDYLDR | 2 | light | 784.3568 | y3 | 403.23 | 65.34 |
| P27810 | KTR1 | SPAYSAYFDYLDR | 2 | heavy | 787.3669 | y9 | 1155.541 | 65.34 |
| P27810 | KTR1 | SPAYSAYFDYLDR | 2 | heavy | 787.3669 | y8 | 1068.509 | 65.34 |
| P27810 | KTR1 | SPAYSAYFDYLDR | 2 | heavy | 787.3669 | y7 | 997.4721 | 65.34 |
| P27810 | KTR1 | SPAYSAYFDYLDR | 2 | heavy | 787.3669 | y6 | 834.4088 | 65.34 |
| P27810 | KTR1 | SPAYSAYFDYLDR | 2 | heavy | 787.3669 | y5 | 687.3404 | 65.34 |
| P27810 | KTR1 | SPAYSAYFDYLDR | 2 | heavy | 787.3669 | y4 | 572.3134 | 65.34 |
| P27810 | KTR1 | SPAYSAYFDYLDR | 2 | heavy | 787.3669 | y3 | 409.2501 | 65.34 |
| P27810 | KTR1 | EHWSFPEWIDEEK | 3 | light | 577.9265 | y8 | 1045.484 | 65.68 |
| P27810 | KTR1 | EHWSFPEWIDEEK | 3 | light | 577.9265 | y7 | 948.4309 | 65.68 |
| P27810 | KTR1 | EHWSFPEWIDEEK | 3 | light | 577.9265 | y6 | 819.3883 | 65.68 |
| P27810 | KTR1 | EHWSFPEWIDEEK | 3 | light | 577.9265 | y5 | 633.309 | 65.68 |
| P27810 | KTR1 | EHWSFPEWIDEEK | 3 | light | 577.9265 | y4 | 520.2249 | 65.68 |
| P27810 | KTR1 | EHWSFPEWIDEEK | 3 | heavy | 580.5979 | y8 | 1053.498 | 65.68 |
| P27810 | KTR1 | EHWSFPEWIDEEK | 3 | heavy | 580.5979 | y7 | 956.4451 | 65.68 |
| P27810 | KTR1 | EHWSFPEWIDEEK | 3 | heavy | 580.5979 | y6 | 827.4025 | 65.68 |
| P27810 | KTR1 | EHWSFPEWIDEEK | 3 | heavy | 580.5979 | y5 | 641.3232 | 65.68 |
| P27810 | KTR1 | EHWSFPEWIDEEK | 3 | heavy | 580.5979 | y4 | 528.2391 | 65.68 |
| Q03103 | ERO1 | YTIENIN[+203.079373]STK | 2 | light | 693.3434 | y9 | 1019.537 | 32.08 |
| Q03103 | ERO1 | YTIENIN[+203.079373]STK | 2 | light | 693.3434 | y8 | 1121.568 | 32.08 |
| Q03103 | ERO1 | YTIENIN[+203.079373]STK | 2 | light | 693.3434 | y8 | 918.4891 | 32.08 |
| Q03103 | ERO1 | YTIENIN[+203.079373]STK | 2 | light | 693.3434 | y7 | 1008.484 | 32.08 |
| Q03103 | ERO1 | YTIENIN[+203.079373]STK | 2 | light | 693.3434 | y7 | 805.405 | 32.08 |
| Q03103 | ERO1 | YTIENIN[+203.079373]STK | 2 | light | 693.3434 | y6 | 879.4418 | 32.08 |
| Q03103 | ERO1 | YTIENIN[+203.079373]STK | 2 | light | 693.3434 | y5 | 765.3989 | 32.08 |
| Q03103 | ERO1 | YTIENIN[+203.079373]STK | 2 | light | 693.3434 | y5 | 562.3195 | 32.08 |

| | | | | | | | | |
|---|---|---|---|---|---|---|---|---|
| Q03103 | ERO1 | YTIENIN[+203.079373]STK | 2 | light | 693.3434 | y4 | 652.3148 | 32.08 |
| Q03103 | ERO1 | YTIENIN[+203.079373]STK | 2 | light | 693.3434 | y4 | 449.2354 | 32.08 |
| Q03103 | ERO1 | YTIENIN[+203.079373]STK | 2 | light | 693.3434 | y3 | 335.1925 | 32.08 |
| Q03103 | ERO1 | YTIENIN[+203.079373]STK | 2 | heavy | 697.3505 | y9 | 1027.551 | 32.08 |
| Q03103 | ERO1 | YTIENIN[+203.079373]STK | 2 | heavy | 697.3505 | y8 | 1129.583 | 32.08 |
| Q03103 | ERO1 | YTIENIN[+203.079373]STK | 2 | heavy | 697.3505 | y8 | 926.5033 | 32.08 |
| Q03103 | ERO1 | YTIENIN[+203.079373]STK | 2 | heavy | 697.3505 | y7 | 1016.499 | 32.08 |
| Q03103 | ERO1 | YTIENIN[+203.079373]STK | 2 | heavy | 697.3505 | y7 | 813.4192 | 32.08 |
| Q03103 | ERO1 | YTIENIN[+203.079373]STK | 2 | heavy | 697.3505 | y6 | 887.456 | 32.08 |
| Q03103 | ERO1 | YTIENIN[+203.079373]STK | 2 | heavy | 697.3505 | y5 | 773.4131 | 32.08 |
| Q03103 | ERO1 | YTIENIN[+203.079373]STK | 2 | heavy | 697.3505 | y5 | 570.3337 | 32.08 |
| Q03103 | ERO1 | YTIENIN[+203.079373]STK | 2 | heavy | 697.3505 | y4 | 660.329 | 32.08 |
| Q03103 | ERO1 | YTIENIN[+203.079373]STK | 2 | heavy | 697.3505 | y4 | 457.2496 | 32.08 |
| Q03103 | ERO1 | YTIENIN[+203.079373]STK | 2 | heavy | 697.3505 | y3 | 343.2067 | 32.08 |
| Q03103 | ERO1 | WEPNLDLFMAR | 2 | light | 696.3425 | y8 | 979.503 | 78.02 |
| Q03103 | ERO1 | WEPNLDLFMAR | 2 | light | 696.3425 | y7 | 865.46 | 78.02 |
| Q03103 | ERO1 | WEPNLDLFMAR | 2 | light | 696.3425 | y6 | 752.376 | 78.02 |
| Q03103 | ERO1 | WEPNLDLFMAR | 2 | light | 696.3425 | y5 | 637.349 | 78.02 |
| Q03103 | ERO1 | WEPNLDLFMAR | 2 | light | 696.3425 | y4 | 524.265 | 78.02 |
| Q03103 | ERO1 | WEPNLDLFMAR | 2 | heavy | 699.3525 | y8 | 985.5231 | 78.02 |
| Q03103 | ERO1 | WEPNLDLFMAR | 2 | heavy | 699.3525 | y7 | 871.4802 | 78.02 |
| Q03103 | ERO1 | WEPNLDLFMAR | 2 | heavy | 699.3525 | y6 | 758.3961 | 78.02 |
| Q03103 | ERO1 | WEPNLDLFMAR | 2 | heavy | 699.3525 | y5 | 643.3692 | 78.02 |
| Q03103 | ERO1 | WEPNLDLFMAR | 2 | heavy | 699.3525 | y4 | 530.2851 | 78.02 |
| Q03103 | ERO1 | NAVLIDLTANPER | 2 | light | 713.3884 | y11 | 1240.69 | 56.77 |
| Q03103 | ERO1 | NAVLIDLTANPER | 2 | light | 713.3884 | y10 | 1141.621 | 56.77 |
| Q03103 | ERO1 | NAVLIDLTANPER | 2 | light | 713.3884 | y9 | 1028.537 | 56.77 |
| Q03103 | ERO1 | NAVLIDLTANPER | 2 | light | 713.3884 | y8 | 915.453 | 56.77 |
| Q03103 | ERO1 | NAVLIDLTANPER | 2 | light | 713.3884 | y7 | 800.4261 | 56.77 |
| Q03103 | ERO1 | NAVLIDLTANPER | 2 | light | 713.3884 | y6 | 687.342 | 56.77 |
| Q03103 | ERO1 | NAVLIDLTANPER | 2 | light | 713.3884 | y5 | 586.2944 | 56.77 |
| Q03103 | ERO1 | NAVLIDLTANPER | 2 | light | 713.3884 | y4 | 515.2572 | 56.77 |

| | | | | | | | | |
|---|---|---|---|---|---|---|---|---|
| Q03103 | ERO1 | NAVLIDLTANPER | 2 | heavy | 716.3985 | y11 | 1246.71 | 56.77 |
| Q03103 | ERO1 | NAVLIDLTANPER | 2 | heavy | 716.3985 | y10 | 1147.641 | 56.77 |
| Q03103 | ERO1 | NAVLIDLTANPER | 2 | heavy | 716.3985 | y9 | 1034.557 | 56.77 |
| Q03103 | ERO1 | NAVLIDLTANPER | 2 | heavy | 716.3985 | y8 | 921.4732 | 56.77 |
| Q03103 | ERO1 | NAVLIDLTANPER | 2 | heavy | 716.3985 | y7 | 806.4462 | 56.77 |
| Q03103 | ERO1 | NAVLIDLTANPER | 2 | heavy | 716.3985 | y6 | 693.3622 | 56.77 |
| Q03103 | ERO1 | NAVLIDLTANPER | 2 | heavy | 716.3985 | y5 | 592.3145 | 56.77 |
| Q03103 | ERO1 | NAVLIDLTANPER | 2 | heavy | 716.3985 | y4 | 521.2774 | 56.77 |
