## Supplementary Table VII for "Functional analysis of Ost3p and Ost6p containing yeast oligosaccharyltransferase*s*"

**Supplement Table VII: Glycosylation occupancy analysis results**

**R=60000**

| Peptide sequence | | 63 N | ± | 63 NTM | ± | 36 N | ± | 36 NTM | ± | OST3 TM1-4 | ± | OST3 TM2-4 | ± | OST6 TM1-4 | ± | OST6 TM2-4 | ± | OST3 | ± | OST6 | ± | DKO | ± |
|---|---|---|---|---|---|---|---|---|---|---|---|---|---|---|---|---|---|---|---|---|---|---|---|
| VRELNE-SAALLLHNER | GPI12_N110 | 13.8 | 3.1 | 12.2 | 2.0 | 12.9 | 1.5 | 23.9 | 1.1 | 6.9 | 0.7 | 13.1 | 3.5 | 4.8 | 0.9 | 3.5 | 0.3 | 71.2 | 4.7 | 7.4 | 3.6 | 4.2 | 1.5 |
| LSNESELYDVFTR | LHS1_N458 | 42.5 | 4.4 | 26.0 | 4.3 | 9.3 | 4.2 | 12.8 | 1.6 | 22.7 | 3.3 | 33.9 | 2.4 | 6.5 | 1.0 | 6.5 | 1.0 | 78.1 | 8.3 | 17.8 | 4.2 | 6.5 | 0.3 |
| IPTELVSENGTK | HMG2_N150 | 36.2 | 10.7 | 17.9 | 6.4 | 27.9 | 5.9 | 43.7 | 1.4 | 17.5 | 6.0 | 25.5 | 3.8 | 28.5 | 0.9 | 6.6 | 0.1 | 84.1 | 3.9 | 17.4 | 0.6 | 8.5 | 0.4 |
| SNNTNYIYR | HEH2_N520 | 46.4 | 9.9 | 63.5 | 6.7 | 89.5 | 7.1 | 96.9 | 1.3 | 23.3 | 3.4 | 26.8 | 3.5 | 87.0 | 3.0 | 23.0 | 10.1 | 44.0 | 4.4 | 111.5 | 6.6 | 10.2 | 2.6 |
| NITLAQIDC-TENQDLCMEH-NIPGFPSLK | PDI_N82 | 33.2 | 0.0 | 11.7 | 1.6 | 32.6 | 13.6 | 15.6 | 6.7 | 17.3 | 16.2 | 65.7 | 16.5 | 18.6 | 4.4 | 19.5 | 8.3 | 21.9 | 7.8 | 54.9 | 29.1 | 10.9 | 5.5 |
| SILFQQQDPF-NESSR | PFF1_N121 | 47.1 | 1.0 | 24.6 | 1.4 | 44.6 | 4.6 | 64.5 | 5.5 | 24.8 | 3.1 | 36.8 | 3.4 | 24.9 | 5.0 | 19.2 | 6.6 | 88.3 | 0.2 | 61.3 | 5.9 | 11.0 | 0.9 |
| NSDVNNSI-DYEGPR | PDI_N117 | 46.1 | 1.5 | 33.3 | 0.6 | 22.9 | 3.8 | 41.1 | 2.5 | 23.4 | 0.2 | 24.1 | 1.1 | 13.2 | 2.5 | 24.2 | 1.4 | 95.0 | 4.8 | 21.4 | 2.0 | 11.6 | 0.7 |
| DNY-WIEYETNTTHPWR | TED1_N266 | 66.1 | 2.6 | 56.5 | 7.8 | 61.7 | 6.7 | 70.7 | 3.0 | 45.5 | 1.9 | 58.1 | 7.4 | 37.1 | 3.9 | 53.5 | 3.4 | 87.0 | 2.9 | 69.8 | 3.3 | 21.0 | 1.1 |
| YTIENINSTK | ERO1_458 | 52.4 | 3.0 | 27.6 | 5.8 | 51.9 | 5.6 | 79.0 | 9.1 | 52.2 | 8.0 | 39.2 | 3.8 | 71.4 | 6.6 | 35.7 | 0.3 | 78.4 | 10.5 | 10.3 | 7.4 | 27.0 | 0.6 |
| LAPTYQELAD-TYANATSDVLI-AK | PDI_N425 | 61.0 | 2.3 | 40.1 | 2.6 | 45.7 | 3.9 | 74.8 | 7.7 | 57.4 | 2.2 | 65.5 | 9.3 | 41.4 | 7.1 | 71.6 | 3.3 | 95.5 | 18.4 | 52.5 | 4.5 | 27.6 | 4.8 |
| NVTDMLYIT-VAQR | FET3_N244 | 47.5 | 21.9 | 35.1 | 8.0 | 57.8 | 9.1 | 64.1 | 1.4 | 52.8 | 3.6 | 55.3 | 8.2 | 16.8 | 8.7 | 58.4 | 4.7 | 127.6 | 21.7 | 60.5 | 3.4 | 28.5 | 2.3 |
| NLDDLNTTVNEQLVFLDSK | OSW7_N297 | 55.4 | 4.9 | 37.6 | 1.3 | 42.8 | 6.7 | 52.6 | 1.6 | 35.4 | 4.4 | 37.9 | 6.1 | 41.8 | 7.1 | 40.3 | 2.1 | 64.9 | 9.5 | 48.2 | 1.4 | 28.6 | 4.1 |
| GPAYANISS-TESVCTVV-GAVPGQ-DYVLGDDFIR | PDR5_N734 | 53.6 | 0.1 | 67.5 | 11.6 | 84.2 | 1.2 | 90.8 | 1.6 | 65.0 | 3.1 | 71.1 | 3.8 | 83.3 | 3.9 | 77.0 | 9.0 | 78.0 | 10.0 | 93.0 | 7.9 | 35.3 | 3.6 |
| NVTSALVSGTTK | KTR1_N120 | 57.4 | 3.3 | 64.0 | 12.3 | 92.8 | 10.5 | 95.7 | 4.4 | 66.6 | 0.9 | 71.6 | 3.5 | 55.0 | 4.2 | 72.3 | 3.9 | 79.6 | 10.2 | 61.2 | 3.6 | 37.3 | 2.4 |
| NSSLYADIYDNK | RAX2_N640 | 89.6 | 43.7 | 32.8 | 10.1 | 64.4 | 8.9 | 59.9 | 8.7 | 69.3 | 5.7 | 81.7 | 7.9 | 67.2 | 4.2 | 64.0 | 3.6 | 71.8 | 16.4 | 80.2 | 17.9 | 38.1 | 9.2 |
| VALVLPNK | EPS1_N264 | 53.9 | 7.0 | 80.6 | 16.6 | 67.2 | 4.4 | 72.5 | 1.8 | 78.0 | 8.8 | 69.4 | 9.6 | 37.7 | 3.1 | 35.7 | 3.8 | 73.7 | 3.0 | 38.3 | 2.8 | 38.5 | 2.2 |
| FSSNETLAI-VYSH-NAPLNQVVNLR | OST1_N217 | 56.7 | 1.4 | 53.0 | 0 | 61.6 | 0.6 | 67.7 | 1.5 | 63.1 | 9.1 | 66.7 | 1.8 | 51.0 | 2.3 | 48.0 | 0.9 | 97.9 | 4.8 | 60.8 | 2.7 | 40.6 | 0.9 |

| | | | | | | | | | | | | | | | | | | | | | | | |
|---|---|---|---|---|---|---|---|---|---|---|---|---|---|---|---|---|---|---|---|---|---|---|---|
| IISFNLSDAETGK | APE3_N150 | 71.3 | 3.6 | 83.1 | 4.1 | 72.8 | 4.6 | 86.7 | 2.9 | 64.7 | 9.4 | 84.5 | 7.9 | 46.8 | 5.6 | 42.3 | 5.3 | 100.3 | 8.3 | 79.0 | 5.3 | 40.9 | 2.7 |
| STAYSL-FANDSDSK | MKC7_N286 | 47.6 | 3.0 | 76.1 | 12.7 | 46.1 | 0.1 | 74.7 | 2.8 | 61.5 | 19.1 | 74.6 | 4.4 | 36.1 | 2.3 | 27.0 | 5.4 | 57.8 | 0.3 | 44.9 | 8.1 | 43.4 | 2.2 |
| FPNITEGELEK | EPS1_N299 | 72.1 | 9.7 | 68.1 | 7.6 | 86.7 | 5.1 | 79.8 | 7.8 | 66.6 | 3.8 | 64.2 | 6.6 | 56.5 | 2.5 | 54.5 | 2.0 | 78.8 | 5.6 | 59.7 | 3.7 | 43.9 | 1.0 |
| NSSSIGYYDL-PAIWLLNDHIAR | YJR1_N219 | 68.5 | 2.8 | 59.6 | 2.5 | 68.6 | 0.4 | 69.6 | 5.0 | 69.6 | 1.7 | 78.7 | 5.5 | 59.3 | 5.0 | 57.3 | 0.7 | 85.7 | 8.9 | 72.2 | 4.7 | 45.0 | 3.1 |
| NLYIDNITFND-PYVSDGLQLK | EXG2_N157 | 69.6 | 6.7 | 65.6 | 5.2 | 34.8 | 2.3 | 47.2 | 3.0 | 56.8 | 5.8 | 62.8 | 5.3 | 36.3 | 3.1 | 23.6 | 0.6 | 86.4 | 6.7 | 67.9 | 4.3 | 47.2 | 1.0 |
| ILGIDPNVTQYT-GYLDVEDEDK | CPY_N124 | 57.4 | 1.3 | 82.6 | 2.2 | 62.8 | 1.6 | 75.3 | 4.8 | 70.0 | 4.7 | 70.6 | 5.4 | 43.7 | 5.0 | 39.3 | 2.4 | 84.2 | 6.9 | 69.0 | 1.0 | 49.8 | 2.9 |
| ILNSAVNMTTIT-PEQLK | YLR413_N429 | 78.8 | 0.2 | 67.3 | 3.3 | 63.0 | 2.1 | 74.0 | 1.9 | 76.2 | 3.2 | 78.8 | 4.4 | 51.6 | 2.1 | 52.7 | 8.5 | 96.8 | 3.8 | 57.1 | 1.3 | 53.7 | 5.3 |
| LNFSIPQR | MNN5_N136 | 84.5 | 3.2 | 85.3 | 2.0 | 66.8 | 5.8 | 86.2 | 5.2 | 75.1 | 4.5 | 78.7 | 8.8 | 47.5 | 1.8 | 54.6 | 12.8 | 98.3 | 21.5 | 69.0 | 8.1 | 53.9 | 3.2 |
| FYWVQGNTT-GIPNAGDETR | SUR7_N47 | 83.0 | 14.3 | 80.3 | 8.0 | 76.4 | 2.4 | 97.7 | 11.1 | 86.0 | 5.1 | 90.6 | 11.4 | 85.5 | 8.4 | 80.6 | 8.9 | 84.0 | 23.8 | 77.4 | 3.0 | 56.7 | 1.0 |
| DAGFNIS-LADVWGR | PLB1_N215 | 195.0 | 17.8 | 179.4 | 20.2 | 142.5 | 7.0 | 180.7 | 26.6 | 85.7 | 8.5 | 85.7 | 1.8 | 76.0 | 0.1 | 83.5 | 8.1 | 206.5 | 18.3 | 148.7 | 2.7 | 57.7 | 1.1 |
| LEYL-DINSTSTTVDL-YDK | WBP1_N60 | 87.7 | 16.6 | 69.3 | 4.4 | 70.8 | 10.0 | 82.4 | 6.4 | 81.8 | 5.1 | 86.1 | 4.9 | 52.6 | 7.8 | 50.0 | 3.1 | 101.2 | 15.0 | 62.0 | 0.8 | 58.0 | 3.6 |
| VRNWTASITDE-VAGEVK | CPY_N479 | 66.8 | 2.1 | 79.3 | 7.9 | 66.0 | 2.8 | 69.8 | 1.9 | 68.2 | 4.8 | 72.1 | 3.3 | 59.6 | 2.8 | 51.6 | 1.2 | 75.9 | 5.2 | 88.1 | 5.0 | 58.6 | 2.5 |
| SDAGFNISLSDL-WAR | PLB2_N217 | 146.6 | 28.2 | 128.6 | 99.6 | 172.7 | 83.6 | 129.4 | 63.5 | 69.4 | 24.2 | 31.4 | 6.9 | 114.2 | 31.7 | 39.9 | 16.5 | 105.6 | 44.1 | 134.4 | 9.9 | 60.0 | 7.6 |
| GLPSWSENET-DIEYLKPGTSYR | PMT2_N403 | 86.2 | 0.1 | 85.0 | 2.7 | 28.8 | 0.7 | 32.8 | 4.4 | 80.6 | 5.0 | 84.1 | 6.0 | 30.6 | 3.1 | 32.4 | 2.3 | 92.7 | 1.5 | 29.0 | 1.8 | 60.2 | 3.7 |
| LNYTASWV-TANPDGLHEK | FET5_N24 | 88.2 | 8.0 | 78.5 | 3.0 | 89.2 | 2.7 | 94.8 | 1.7 | 70.1 | 3.8 | 70.6 | 2.7 | 83.6 | 4.7 | 68.5 | 2.4 | 94.0 | 1.6 | 86.7 | 3.8 | 66.9 | 2.0 |
| IKVDDLNA-TAWDLYR | APE3_N85 | 92.3 | 14.4 | 92.9 | 7.3 | 100.8 | 22.9 | 107.0 | 7.0 | 84.7 | 17.2 | 95.4 | 10.7 | 90.6 | 5.1 | 70.5 | 12.5 | 107.0 | 14.2 | 77.1 | 13.5 | 70.5 | 5.3 |
| LANYSTP-DYGHPTR | APE3_N96 | 77.6 | 7.4 | 77.1 | 4.1 | 78.6 | 9.7 | 93.8 | 6.0 | 84.7 | 14.1 | 96.9 | 11.8 | 78.0 | 5.5 | 73.9 | 7.5 | 90.7 | 6.5 | 88.0 | 3.6 | 71.1 | 4.1 |
| EIGPETSSHGL-VYYSNNTYIQLE-DASDDTR | RAX2_N88 | 86.6 | 1.3 | 75.0 | 1.7 | 65.6 | 4.7 | 77.5 | 14.6 | 87.3 | 11.9 | 84.7 | 25.8 | 72.8 | 2.3 | 54.6 | 8.0 | 93.1 | 5.6 | 66.8 | 4.4 | 75.8 | 7.2 |
| YAFFNNI-TYVTPK | FET5_N364 | 88.8 | 1.4 | 75.3 | 2.9 | 83.8 | 8.2 | 75.0 | 2.8 | 87.7 | 6.1 | 92.4 | 12.1 | 85.3 | 14.6 | 84.3 | 3.3 | 73.7 | 12.0 | 79.9 | 12.1 | 79.2 | 7.2 |
| LTLSPSGNDSET-QYYTTGEFILPDR | WBP1_N332 | 91.4 | 16.1 | 86.5 | 7.5 | 84.6 | 10.2 | 92.3 | 3.0 | 96.5 | 6.6 | 99.3 | 6.9 | 84.7 | 4.2 | 82.2 | 8.4 | 94.8 | 4.1 | 82.5 | 4.5 | 79.3 | 2.2 |
| SFANTTA-FALSPPVDGFVGK | APE3_N162 | 86.0 | 3.4 | 85.2 | 7.4 | 89.9 | 7.1 | 96.3 | 9.0 | 90.3 | 2.5 | 92.7 | 6.3 | 96.3 | 3.6 | 90.1 | 4.3 | 95.1 | 7.1 | 86.7 | 4.8 | 81.4 | 5.8 |

| | | | | | | | | | | | | | | | | | | | | | | | |
|---|---|---|---|---|---|---|---|---|---|---|---|---|---|---|---|---|---|---|---|---|---|---|---|
| THLILSNSTI-IHDFDPLNLNVGVLPR | CHS7_N31 | 82.3 | 4.4 | 83.3 | 2.5 | 98.4 | 3.3 | 88.1 | 5.5 | 90.7 | 6.4 | 82.0 | 8.1 | 93.8 | 3.1 | 87.3 | 4.2 | 88.9 | 3.1 | 91.0 | 4.2 | 82.2 | 3.2 |
| TTLVDNNTWNNTHIAIVGK | STT3_N539 | 81.2 | 8.7 | 90.1 | 7.5 | 82.9 | 3.0 | 80.3 | 9.3 | 96.2 | 4.5 | 89.3 | 9.9 | 78.4 | 4.9 | 64.5 | 14.2 | 78.6 | 11.4 | 77.1 | 10.8 | 83.5 | 9.0 |
| IDADFNATFYS-MANK | PDI_N174 | 72.3 | 5.1 | 72.8 | 9.2 | 90.6 | 23.2 | 127.0 | 34.0 | 71.7 | 2.3 | 95.3 | 23.2 | 92.7 | 9.3 | 84.6 | 8.1 | 106.6 | 7.5 | 125.6 | 15.1 | 83.6 | 3.3 |
| NGVNYAFFNNI-TYTAPK | FET3_N359 | 62.8 | 18.9 | 78.3 | 23.5 | 73.4 | 16.5 | 74.0 | 6.3 | 94.6 | 6.0 | 99.4 | 39.3 | 55.6 | 17.0 | 94.5 | 4.1 | 123.3 | 62.8 | 96.6 | 12.2 | 85.3 | 3.7 |
| VQTVGGAIEVTGNFSTLDLSSLK | ECM33_N304 | 95.2 | 7.3 | 110.1 | 2.6 | 95.2 | 5.0 | 93.0 | 11.5 | 107.6 | 9.3 | 116.0 | 3.1 | 103.2 | 13.7 | 73.3 | 15.0 | 116.5 | 14.1 | 81.3 | 1.4 | 87.9 | 0.9 |
| SYASDIGAPLFN-STEK | GPI16_N184 | 88.7 | 7.2 | 89.8 | 6.5 | 97.1 | 2.6 | 95.2 | 7.3 | 103.1 | 3.8 | 105.8 | 6.7 | 97.2 | 5.9 | 90.5 | 15.9 | 96.1 | 2.7 | 102.1 | 1.1 | 91.6 | 5.3 |
| LLGLNSGFSNSTILQETLNSK | ERG3_N40 | 77.5 | 1.0 | 89.8 | 6.6 | 97.8 | 0.2 | 88.1 | 1.3 | 91.4 | 9.7 | 92.0 | 11.1 | 102.5 | 1.0 | 91.9 | 1.1 | 87.3 | 6.0 | 101.1 | 2.9 | 94.7 | 2.6 |
| FSNNGTFFETEE-PIVETK | SEC66_N12 | 82.9 | 1.3 | 86.7 | 1.5 | 80.4 | 3.7 | 86.7 | 4.6 | 86.9 | 2.4 | 83.9 | 2.4 | 83.8 | 5.3 | 83.1 | 2.1 | 91.4 | 6.4 | 85.4 | 6.1 | 94.9 | 0.6 |
| NISNTPPTSD-PEK | GPI13_N411 | 100.4 | 10.0 | 98.7 | 7.9 | 89.9 | 5.6 | 94.1 | 2.9 | 95.2 | 10.3 | 102.8 | 14.5 | 100.2 | 5.6 | 97.7 | 4.0 | 102.7 | 7.7 | 91.3 | 5.4 | 96.2 | 7.0 |
| YNQTETFK | ROT1_N139 | 110.6 | 5.9 | 100.5 | 1.2 | 108.6 | 4.0 | 103.1 | 2.3 | 102.0 | 5.1 | 99.9 | 7.4 | 101.6 | 5.0 | 93.8 | 6.1 | 96.9 | 4.4 | 101.3 | 0.7 | 104.9 | 5.7 |
| FFYSNNGSQ-FYIR | GAS1_N40 | 97.8 | 6.7 | 104.1 | 3.6 | 85.9 | 5.5 | 93.9 | 6.1 | 104.0 | 6.0 | 95.2 | 5.3 | 88.7 | 1.4 | 93.9 | 4.6 | 101.5 | 17.2 | 71.6 | 4.4 | 105.1 | 5.5 |
| SSSE-DALNNNTD-TYGNFIR | PDAT_N439 | 132.2 | 12.3 | 120.4 | 4.5 | 129.6 | 3.0 | 125.8 | 10.1 | 113.4 | 11.7 | 107.9 | 17.5 | 116.0 | 1.2 | 117.5 | 6.0 | 119.6 | 18.2 | 109.1 | 6.4 | 106.7 | 1.7 |
| NQTIQGDVH-GITK | RAX2_N677 | 97.1 | 0.4 | 118.6 | 4.9 | 118.4 | 1.9 | 107.8 | 4.8 | 110.2 | 7.6 | 102.5 | 7.5 | 102.4 | 1.0 | 107.0 | 2.3 | 98.3 | 6.6 | 104.8 | 11.9 | 111.0 | 6.4 |
| YTGNQTYVD-WAEK | DCW1_N203 | 145.0 | 46.8 | 102.8 | 3.7 | 112.9 | 3.3 | 113.6 | 20.3 | 115.1 | 15.2 | 112.5 | 15.7 | 96.0 | 10.0 | 111.4 | 44.2 | 116.6 | 5.9 | 112.9 | 5.3 | 112.4 | 23.6 |
| NGTSAYVYTSS-SEFLAK | CRH2_N310 | 90.7 | 18.9 | 104.2 | 10.3 | 90.0 | 3.2 | 113.0 | 14.9 | 103.3 | 9.2 | 100.7 | 17.5 | 95.6 | 13.0 | 99.1 | 9.1 | 108.5 | 38.5 | 81.3 | 21.9 | 112.6 | 48.2 |
| SIVNPGGSNLTY-TIER | PLB2_N193 | 86.3 | 3.2 | 102.4 | 9.8 | 102.7 | 1.9 | 97.0 | 4.1 | 95.1 | 12.8 | 35.5 | 4.4 | 59.0 | 49.6 | 77.7 | 40.0 | 98.3 | 25.8 | 93.7 | 13.8 | 114.9 | 0.8 |
| NVTEAQIIGNK | CNE1_N416 | 106.6 | 21.0 | 112.5 | 5.6 | 111.9 | 11.2 | 147.6 | 28.4 | 109.1 | 9.2 | 96.1 | 13.3 | 146.8 | 1.3 | 105.1 | 22.8 | 103.9 | 26.1 | 90.2 | 12.6 | 117.8 | 33.0 |
| FASYYAND-TITVK | EXG2_N50 | 97.5 | 7.1 | 125.0 | 10.3 | 127.7 | 13.6 | 108.2 | 8.3 | 123.0 | 19.4 | 113.9 | 19.8 | 111.7 | 7.9 | 66.1 | 3.6 | 92.3 | 7.5 | 89.2 | 1.2 | 139.3 | 3.6 |
